# Computational Analysis of Protein Stability and Allosteric Interaction Networks in Distinct Conformational Forms of the SARS-CoV-2 Spike D614G Mutant: Reconciling Functional Mechanisms through Allosteric Model of Spike Regulation

**DOI:** 10.1101/2021.01.26.428331

**Authors:** Gennady M. Verkhivker, Steve Agajanian, Denis Oztas, Grace Gupta

## Abstract

Structural and biochemical studies SARS-CoV-2 spike mutants with the enhanced infectivity have attracted significant attention and offered several mechanisms to explain the experimental data. The development of a unified view and a working model which is consistent with the diverse experimental data is an important focal point of the current work. In this study, we used an integrative computational approach to examine molecular mechanisms underlying functional effects of the D614G mutation by exploring atomistic modeling of the SARS-CoV-2 spike proteins as allosteric regulatory machines. We combined coarse-grained simulations, protein stability and dynamic fluctuation communication analysis along with network-based community analysis to simulate structures of the native and mutant SARS-CoV-2 spike proteins in different functional states. The results demonstrated that the D614 position anchors a key regulatory cluster that dictates functional transitions between open and closed states. Using molecular simulations and mutational sensitivity analysis of the SARS-CoV-2 spike proteins we showed that the D614G mutation can improve stability of the spike protein in both closed and open forms, but shifting thermodynamic preferences towards the open mutant form. The results offer support to the reduced shedding mechanism of S1 domain as a driver of the increased infectivity triggered by the D614G mutation. Through distance fluctuations communication analysis, we probed stability and allosteric communication propensities of protein residues in the native and mutant SARS-CoV-2 spike proteins, providing evidence that the D614G mutation can enhance long-range signaling of the allosteric spike engine. By employing network community analysis of the SARS-CoV-2 spike proteins, our results revealed that the D614G mutation can promote the increased number of stable communities and allosteric hub centers in the open form by reorganizing and enhancing the stability of the S1-S2 inter-domain interactions and restricting mobility of the S1 regions. This study provides atomistic-based view of the allosteric interactions and communications in the SARS-CoV-2 spike proteins, suggesting that the D614G mutation can exert its primary effect through allosterically induced changes on stability and communications in the residue interaction networks.

## Introduction

Understanding of the molecular principles driving the coronavirus disease 2019 (COVID-19) associated with the severe acute respiratory syndrome (SARS)^1–5^ has been at the focal point of biomedical research since the start of the pandemic a year ago. SARS-CoV-2 infection is transmitted when the viral spike (S) glycoprotein binds to the host cell receptor, leading to the entry of S protein into host cells and membrane fusion.^6–8^ The full-length SARS-CoV-2 S protein consists of two main domains, amino (N)-terminal S1 subunit and carboxyl (C)-terminal S2 subunit. The subunit S1 is involved in the interactions with the host receptor and includes an N-terminal domain (NTD), the receptor-binding domain (RBD), and two structurally conserved subdomains (SD1 and SD2). Structural and biochemical studies have shown that the mechanism of virus infection may involve spontaneous conformational transformations of the SARS-CoV-2 S protein between a spectrum of closed and receptor-accessible open forms, where RBD continuously switches between “down” and “up” positions where the latter can promote binding with the host receptor ACE2.^9–11^ The S1 subunit is characterized by variant regions, particularly in the receptor binding motif (RBM) interacting with the host receptor ACE2 enzyme. The S1 regions also include C-terminal domain 1/CTD1 (or SD1) and C-terminal domain 2/CTD2 (or SD2) domains. Genomic studies established that the S2 subunit is an evolutionary conserved fusion machinery that contains upstream helix (UH), an N-terminal hydrophobic fusion peptide (FP), fusion peptide proximal region (FPPR), heptad repeat 1 (HR1), central helix region (CH), connector domain (CD), heptad repeat 2 (HR2), transmembrane domain (TM) and cytoplasmic tail (CT).^12^ The S1 regions are structurally situated above the S2 subunit^13–17^ and serve as dynamic protective shield of the fusion machine. Upon proteolytic activation at the S1/S2 and dissociation of S1 from S2, a cascade of tectonic structural rearrangements in the S2 subunit is initiated to mediate the fusion of the viral and cellular membranes.^18, 19^ The rapidly evolving body of biophysical studies and cryo-EM structures of the SARS-CoV-2 S proteins characterized distinct conformational arrangements of the S protein trimers in the prefusion form that are manifested by a dynamic equilibrium between the closed (“RBD-down”) and the receptor-accessible open (“RBD-up”) form required for the S protein fusion to the viral membrane.^20–29^ The recent cryo-EM structure of the SARS-CoV-2 S trimer demonstrated a population-shift between a spectrum of closed states that included a structurally rigid closed form and more dynamic closed states preceding a transition to the fully open S conformation.^30^ Protein engineering and structural studies also showed how disulfide bonds and proline mutations can modulate stability of the SARS-CoV-2 S trimer^31^ and lead to the thermodynamic shifts between the closed-down conformation and the open form exposed to binding with the ACE2 host receptor.^32–34^ The cryo-EM structures and biophysical tomography tools characterized the structures of the SARS-CoV-2 S trimers in situ on the virion surface and confirmed a population shift between different functional states, showing that conformational transitions can proceed through an obligatory intermediate in which all three RBD domains are in the closed conformations and are oriented towards the viral particle membrane.^35, 36^ Cryo-EM structural studies also mapped a mechanism of conformational events associated with ACE2 binding, showing that the compact closed form of the SARS-CoV-2 S protein becomes weakened after furin cleavage between the S1 and S2 domains, leading to the increased population of partially open states and followed by ACE2 recognition that accelerates conformational transformations to a fully open and ACE2-bound form priming the protein for fusion activation.^37^ Cryo-EM structures of SARS-CoV-2 S protein in the presence and absence of ACE2 receptor suggested a pH-dependent switch that mediates conformational switching of RBD regions.^38^ This study demonstrated that pH-dependent refolding region (residues 824-858) at the interdomain interface displayed dramatic structural rearrangements and mediated coordinated movements of the entire trimer, giving rise to a single 1 RBD-up conformation at pH 5.5 while all-down closed conformation was favorable at lower pH. ^38^

SARS-CoV-2 S mutants with the enhanced infectivity profile including D614G mutational variant have attracted an enormous attention in the scientific community following the evidence of the mutation enrichment via epidemiological surveillance, resulting in proliferation of experimental data and a considerable variety of the proposed mechanisms explaining functional observations.^39–41^ The latest biochemical studies provided a compelling evidence of a phenotypic advantage and the enhanced infectivity conferred by the D614G mutation.^42^ The recent structural and biochemical studies suggested that the D614G mutation can act by shifting the population of the SARS-CoV-2 S trimer from the closed form (53% of the equilibrium) in the native spike protein to a widely-open topology of the “up” protomers in the D614G mutant with 36% of the population adopting a single open protomer, 39% with two open protomers and 20% with all three protomers in the open conformation.^43^ The cryo-EM structures of the S-D614 and S-G614 ectodomains and the structure of the cleaved S-G614 ectodomain detailed subtle effects of the D614G mutation, showing the increased population of the 1-RBD-up open form over the closed state in the S-GSAS/D614G structure.^44^

This study also demonstrated that the S-D614G mutant could modulate conformational population of the S protein and result in the increased furin cleavage efficiency of the S ectodomain. In addition, these experiments suggested that the D614G mutation in the SD2 domain can induce allosteric effect leading to coordinated movements and structural shifts between the up and down RBD conformations.^44^ Negative stain electron microscopy revealed the higher 84% percentage of the 1-up RBD conformation in the S-G614 protein, suggesting the increased epitope exposure as a mechanism of enhanced vulnerability to neutralization.^45^ The reported retroviruses pseudotyped with S-G614 showed a markedly greater infectivity than the S-D614 protein that was correlated with a reduced S1 shedding, greater stability of the S-G614 mutant and more significant incorporation of the S protein into the pseudovirion.^46^ In addition, it was confirmed that the D614G mutation does not produce greater binding affinity of S protein for ACE2 neither makes it more resistant to neutralization.^46^ This evidence offered an alternative mechanism to the previously proposed functional scenario in which D614G mutation would promote rather than limit shedding of the S1 domain.^47^ Consistent with the reduced shedding mechanism induced by the D614G mutation, the reported cryo-EM structures of a full-length S-G614 trimer featuring three distinct prefusion conformations provided a mechanistic explanation for the increased stability of the highly infective mutant.^48^

According to this study, D614G may promote ordering of the partly disordered loop located near the furin cleavage site that strengthens the inter-domain interactions between the NTD and CTD1 regions and enhances the inter-protomer contacts and stability of the mutated S protein, thereby inhibiting a premature dissociation of the S1 subunit which eventually leads to the increased number of functional spikes and stronger infectivity.^48^ Structure-based protein design and cryo-EM structure determination established that both D614G and D614N mutations can result in the increased fusogenicity and stability which can be explained by a decrease in a premature shedding of the S1 domain.^49^ The cryo-EM structure revealed a stable closed mutant conformation, suggesting that D614G/N mutations can attenuate the repulsive charge interactions at the interface between S1 and S2 providing tighter packing of the head domains against S2.^49^

Nonetheless, a consensus view on the exact mechanism underlying the functional effects and increased infectivity of S-D614G spike mutant is yet to be established, owing to often conflicting experimental evidence and the proposed mutually exclusive scenarios underpinning molecular consequences of the D614G mutation on structure, dynamics and energetics of virus entry.^42^ The recent illumining review highlighted several prevalent mechanisms actively debated in the field offered to explain diverse experimental data, including D614G-induced modulation of cleavage efficiency of S protein; “openness” scenario advocating mutation-induced shift to the open states favorable for RBD-ACE2 interaction; “density” hypothesis suggesting a more efficient S incorporation into the virion; and “stability” mechanism that implicates mutation-induced enhancement in the association and stability of prefusion spike trimers as a driving force of greater infectivity.^42^

The growing body of computational modeling studies investigating dynamics and molecular mechanisms of S-D614G mutational variant produced interesting but often inconsistent data that fit different mechanisms. The development of a more unified view and a working theoretical model which can explain the diverse experimental observations is an important focal point of the current work. Several studies suggested that the D614G mutation may affect the intra-protomer energetics and favor “1-up” open conformation by eliciting specific interaction changes in the CTD1 and FP regions.^50, 51^ Microsecond all-atom molecular dynamics (MD) simulations recently probed the effects of the D614G mutation, suggesting that mutational variant favors an open conformation in which S-G614 protein maintains the symmetry in the number of persistent contacts between the three protomers.^50^ By applying a differential contact analysis, this study argue that contacts in CTD1-CTD2 (528-685) and FP-FPPR (816-911) regions may be responsible for conformational shift of the S-G614 mutant towards a partially open state. Coarse-grained normal mode analyses combined with Markov model and computation of transition probabilities characterized the dynamics of the S protein and mutational variants, predicting the increase in the open state occupancy for the more infectious D614G mutation due to the increased flexibility of the closed state and the enhanced rigidification of the open spike form.^52^ Computer-based mutagenesis and energy analysis of the thermodynamic stability of the S-D614 and S-G614 proteins in the closed and partially open conformations showed that local interactions near D614 position are energetically frustrated and may create an unfavorable environment that is stabilized in the S-G614 mutant through strengthening of the inter-protomer association between S1 and S2 regions.^53^ Using time-independent component analysis (tICA) and protein graph connectivity network, another computational study identified the hotspot residues that may exhibit long-distance coupling with the RBD opening, showing that the D614G could exert allosteric effect on the flexibility of the RBD regions.^54^ Structure-based physical model showed that the D614G mutation may induce a packing defect in S1 that promotes closer association and stronger interactions with S2 subunit, thereby supporting the reduced shedding hypothesis.^55^ Computational modeling and MD simulations have been instrumental in predicting dynamics and function of SARS-CoV-2 glycoproteins.^56–64^ Our recent study examined molecular basis of the SARS-CoV-2 binding with ACE2 enzyme showing that coevolution and conformational dynamics conspire to drive cooperative binding interactions and signal transmission.^62^ Using protein contact networks and perturbation response scanning allosteric sites on the SARS-CoV-2 spike protein were proposed.^63^ Molecular simulations and network modeling approaches were used on our most recent investigation to present evidence that the SARS-CoV-2 spike protein can function as an allosteric regulatory engine that fluctuates between dynamically distinct functional states.^64^

In this study, we used an integrative computational approach to examine molecular mechanisms underlying the functional effects of the D614G mutation in different states of the SARS-CoV-2 S protein. We combined coarse-grained (CG) simulations and atomistic reconstruction of dynamics trajectories with dynamic fluctuation communication analysis, modeling of the residue interaction networks and network community analysis to simulate cryo-EM structures of the SARS-CoV-2 D614 and G614 proteins corresponding to the S-RRAR and S-GSAS furin-cleavage deficient constructs. We quantify the effect of mutations on functional dynamics and stability of the SARS-CoV-2 S proteins and demonstrate that D614 site may anchor a key regulatory hinge cluster that dictates functional transitions between open and closed states. Through distance fluctuations communication analysis of the atomistic conformational ensembles, we probe allosteric communication residue preferences in the S-D614 and S-G614 proteins, providing evidence that the D614G mutation can favor open state to restore allosteric signaling function of the spike engine. By employing network-based community analysis as proxy for characterization of protein stability and allosteric interactions, we show that the D614G mutation can partially rearrange the network organization and promote the larger number of stable communities in the open form by enhancing the S1-S2 inter-domain interactions. This study provides a novel insight into the molecular mechanisms underlying the effect of D614G mutation by examining SARS-CoV-2 S protein as an allosteric regulatory machine. The results support the reduced shedding hypothesis of the D614G mutant function suggesting that mutation can have a moderate effect on local dynamics and interactions, while exerting its primary effect through allosterically induced changes on stability and communications in the residue interaction networks.

## Materials and Methods

### Coarse-Grained Simulations and Elastic Models

We employed coarse-grained (CG) CABS model^65–71^ for simulations of the cryo-EM structures of the SARS-CoV-2 S D614 and D614G mutant ectodomains with the native K986 and V987 residues. More specifically CG-CABS simulations were performed for the cryo-EM structure of SARS-CoV-2 S-GSAS/D614 in the closed-all down state (pdb id 7KDG), S-GSAS/D614 in the 1 RBD-up open state (pdb id 7KDH), SARS-CoV-2 S-GSAS/G614 mutant ectodomain in the closed form (pdb id 7KDK), SARS-CoV-2 S-GSAS/G614 mutant in the 1 RBD-up open state (pdb id 7KDL) as well as furin cleaved SARS-CoV-2 spike mutant S-RRAR/G614 in the closed form (pdb id 7KDI) and 1 RBD-up open state (pdb id 7KDJ) (Table 1, Figure 1). In the CG-CABS model, the amino acid residues are represented by Cα, Cβ, the center of mass of side chains and another pseudoatom placed in the center of the Cα-Cα pseudo-bond.^65–67^ We employed CABS-flex approach that efficiently combines a high-resolution coarse-grained model and efficient search protocol capable of accurately reproducing all-atom MD simulation trajectories and dynamic profiles of large biomolecules on a long time scale.^69–71^ The sampling scheme of the CABS model used in our study is based on Monte Carlo replica-exchange dynamics and involves a sequence of local moves of individual amino acids in the protein structure as well as moves of small fragments.^65–67^

**Figure 1.**
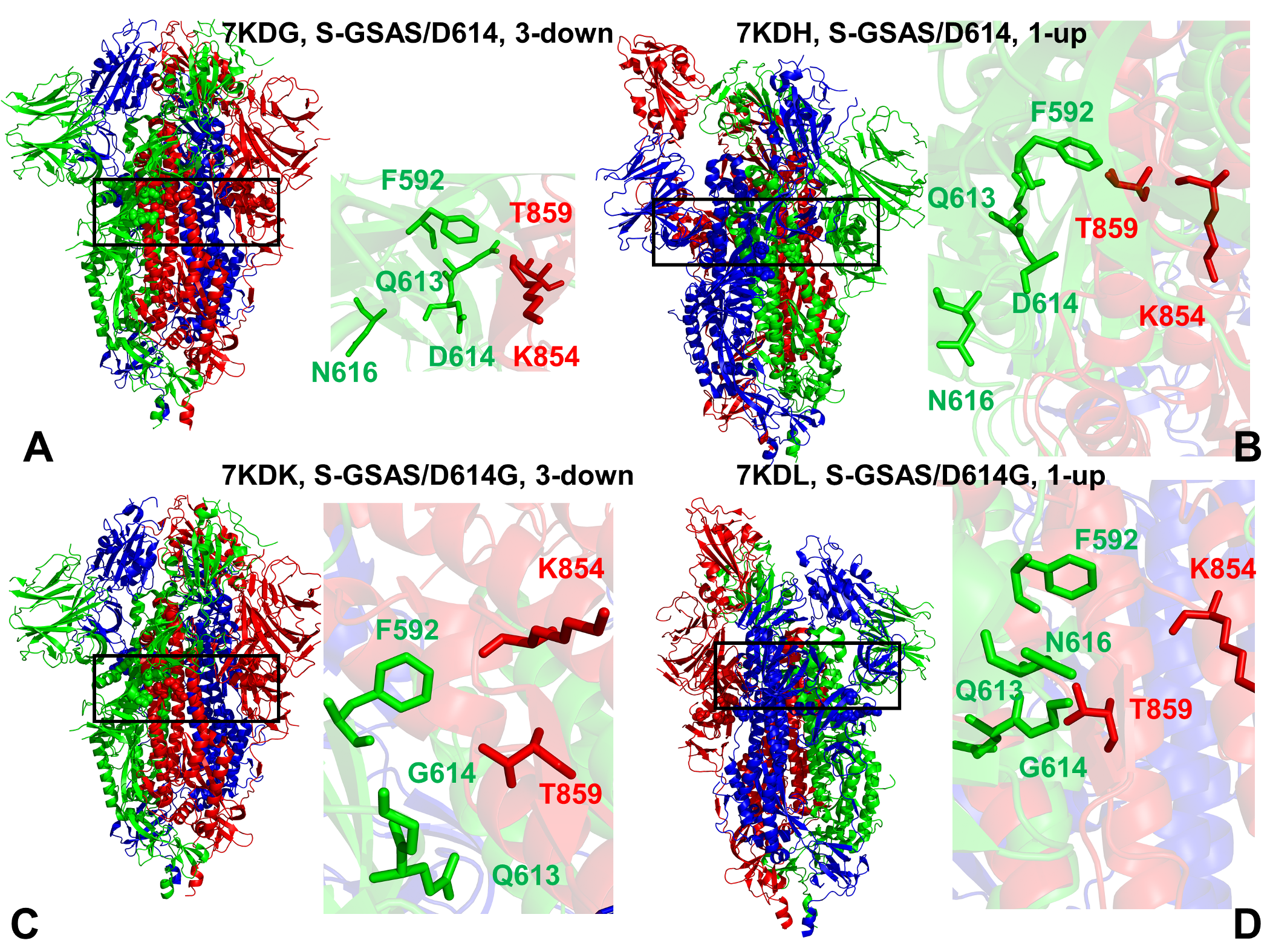
Cryo-EM structures of the SARS-CoV-2 spike (S) protein trimer structures used in this study. (A) The cryo-EM structure of SARS-CoV-2 S-GSAS/D614 in the closed-all down state (pdb id 7KDG). The structure is shown in ribbons with protomers A,B,C are colored in green, red and blue respectively. A close-up of the key residues and interactions near D614 position. D614, Q613, N616 of protomer A (green sticks) and K854, T859 of the adjacent protomer B are shown in red sticks. (B) S-GSAS/D614 in the 1 RBD-up open states (pdb id 7KDH). A close-up of the key residues and interactions near D614 position is shown and residues are annotated as in panel (A). (C) The cryo-EM structure of SARS-CoV-2 S-GSAS/G614 mutant ectodomain in the closed form (pdb id 7KDK). A close-up of the key residues and interactions near D614 position is shown and residues are annotated as in panel (A). (D) The cryo-EM structure the SARS-CoV-2 S-GSAS/G614 mutant in the 1 RBD-up open states (pdb id 7KDL). A close-up of the key residues and interactions near D614 position is shown and residues are annotated as in panel (A).

**Table 1.** . Structures of SARS-CoV2 spike protein structures examined in this study.

| **PDB code** | **Description** | **RBDs position** |
| --- | --- | --- |
| 7KDG | SARS-CoV-2 Spike Protein Trimer (S-GSAS) D614 | 3 down |
| 7KDH | SARS-CoV-2 Spike Protein Trimer (S-GSAS) D614 | 1 up |
| 7KDK | SARS-CoV-2 Spike Protein Trimer (S-GSAS) G614 | 3 down |
| 7KDL | SARS-CoV-2 Spike Protein Trimer (S-GSAS) G614 | 1 up |
| 7KDI | SARS-CoV-2 Spike Protein Trimer (S-RRAR) G614 | 3 down |
| 7KDJ | SARS-CoV-2 Spike Protein Trimer (S-RRAR) G614 | 1 up |

CABS-flex standalone package dynamics implemented as a Python 2.7 object-oriented package was used for fast simulations of protein structures.^69–71^ In the CABS-flex package we also MODELLER-based reconstruction of generated models and simulation trajectories to all-atom representation. The default settings were applied in which soft native-like restraints are imposed only on pairs of residues fulfilling the following conditions : the distance between their *C*^α^ atoms was smaller than 8 Å, and both residues belong to the same secondary structure elements. A total of 1,000 independent CG-CABS simulations were performed for each of the studied systems. In each simulation, the total number of cycles was set to 10,000 and the number of cycles between trajectory frames was 100.

All structures were obtained from the Protein Data Bank.^72, 73^ Protein residues in the cryo-EM structures were inspected for missing residues and protons that were initially added and assigned according to the WHATIF program web interface.^74, 75^ The structures were further parsed through the Protein Preparation Wizard (Schrödinger, LLC, New York, NY) and were subjected to the check of bond order, assignment and adjustment of ionization states, removal of crystallographic water molecules and co-factors, capping of the termini, assignment of partial charges, and addition of possible missing atoms and side chains that were not assigned in the initial processing with the WHATIF program.

The cryo-EM structures of the SARS-CoV-2 S proteins contained a number of missing loops of various lengths that were dispersed throughout the structure and required a rigorous template-based loop modeling and reconstruction. The modeled missing loops were located in the NTD regions (residues 1-26, 70-79,144-164,173-185,246-262, 294-304), RBD regions (residues 469-488) CTD2 regions (residues 621-640), the cleavage site at the S1/S2 boundary (residues 677-689) and residues in the S2 subunit 828-853.

The missing loops in the RBD regions (residues 445-446,469-488) were reconstructed by template-based modeling using the cryo-EM structure of human ACE2 in the presence of the neutral amino acid transporter B^0^AT1 complexed with SARS-CoV-2 RBD protein (pdb id 6M17) that includes the complete RBD region as a template. The missing loops in the studied cryo-EM structures of the SARS-CoV-2 S protein were reconstructed and optimized using template-based loop prediction approaches ModLoop,^76^ ArchPRED server^77^ and further confirmed by FALC (Fragment Assembly and Loop Closure) program.^78^ The side chain rotamers were refined and optimized by SCWRL4 tool.^79^ The conformational ensembles were also subjected to all-atom reconstruction using PULCHRA method^80^ and CG2AA tool^81^ to produce atomistic models of simulation trajectories. The protein structures were then optimized using atomic-level energy minimization with a composite physics and knowledge-based force fields as implemented in the 3Drefine method.^82^. The atomistic structures from simulation trajectories were further elaborated by adding N-acetyl glycosamine (NAG) glycan residues and optimized. The glycosylated microenvironment for atomistic models of the simulation trajectories was mimicked by using the structurally resolved glycan conformations for 22 most occupied N-glycans in each as determined in the cryo-EM structures of the SARS-CoV-2 spike S trimer in the closed state (K986P/V987P,) (pdb id 6VXX) and open state (pdb id 6VYB),^16^ and the cryo-EM structure SARS-CoV-2 spike trimer (K986P/V987P) in the open state (pdb id 6VSB).^17^

Principal component analysis (PCA) of simulation trajectories was done using elastic network models (ENM) analysis.^83^ Two elastic network models: Gaussian network model (GNM)^83–85^ and Anisotropic network model (ANM) approaches^86^ were used to compute the amplitudes of isotropic thermal motions and directionality of anisotropic motions. The functional dynamics analysis was conducted using the GNM in which protein structure is reduced to a network of *N* residue nodes identified by *Cα* atoms and the fluctuations of each node are assumed to be isotropic and Gaussian. Conformational mobility profiles in the essential space of low frequency modes were obtained using DynOmics server^85^ and ANM server.^86^

### Mutational Sensitivity Analysis and Alanine Scanning

To compute protein stability changes in the SARS-CoV-2 S structures, we conducted a systematic alanine scanning of protein residues in the SARS-CoV-2 trimer mutants as well as mutational sensitivity analysis at the mutational site for both SARS-CoV-2 S-D614 and SARS-CoV-2 S-G614 structures. Two different approaches were used. Alanine scanning of protein residues was performed using FoldX approach.^87–90^ and BeAtMuSiC approach.^91–93^ If a free energy change between a mutant and the wild type (WT) proteins ΔΔG= ΔG (MT)-ΔG (WT) > 0, the mutation is destabilizing, while when ΔΔG <0 the respective mutation is stabilizing. BeAtMuSiC approach is based on statistical potentials describing the pairwise inter-residue distances, backbone torsion angles and solvent accessibilities, and considers the effect of the mutation on the strength of the interactions at the interface and on the overall stability of the complex.^91–93^ We leveraged rapid calculations based on statistical potentials to compute the ensemble-averaged alanine scanning computations and mutational sensitivity analysis at D614 and G614 positions using equilibrium samples from reconstructed simulation trajectories.

### Distance Fluctuations Analysis of Protein Stability and Allosteric Propensities

Using a protein mechanics-based approach^94–97^ we probed residue stability and communication propensities with the aid of distance fluctuation analysis of the simulation trajectories for the studied SARS-CoV-2 S proteins. The fluctuations of the mean distance between a given residue and all other residues in the ensemble were converted into distance fluctuation stability and communication indexes that measure the energy cost of the residue deformation during simulations. Our previous studies^98, 99^ and related models^100, 101^ adapted this metric to characterize allosteric communication propensities of protein residues through their mechanical properties since mean square deformations of a given residue with respect to the rest of the protein are related to the inter-residue communication strength.

We computed the fluctuations of the mean distance between each atom within a given residue and the atoms that belong to the remaining residues of the protein. The high values of distance fluctuation indexes are associated with residues that display small fluctuations in their distances to all other residues, while small values of this stability parameter would point to more flexible sites that experience large deviations of their inter-residue distances. In our model, the distance fluctuation stability index for each residue is calculated by averaging the distances between the residues over the simulation trajectory using the following expression:

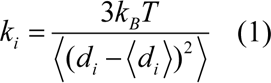

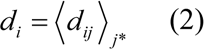

*d_ij_* is the instantaneous distance between residue *i* and residue *j*, *k_B_* is the Boltzmann constant, T =300K. 〈 〉 denotes an average taken over the MD simulation trajectory and d = 〈*d_ij_*〉_j_* is *^i ij^ j*\*the average distance from residue *i* to all other atoms *j* in the protein (the sum over *j*_*_ implies the exclusion of the atoms that belong to the residue *i*). The interactions between the *C*_α_ atom of residue *i* and the *C*_α_ atom of the neighboring residues *i* -1 and *i* +1 are excluded in the calculation since the corresponding distances are nearly constant. The inverse of these fluctuations yields an effective force constant *ki* that describes the ease of moving an atom with respect to the protein structure.

### Dynamic-Based Modeling of Residue Interaction Network and Community Analysis

A graph-based representation of protein structures^102, 103^ is used to represent residues as network nodes and the inter-residue edges to describe residue interactions. The details of network construction were described in our previous studies.^104, 105^ We constructed the residue interaction networks using both dynamic correlations^103^ and coevolutionary residue couplings^99^ that provide complementary descriptions of the allosteric interactions and yield robust network signatures of long-range couplings and communications. The ensemble of shortest paths is determined from matrix of communication distances by the Floyd-Warshall algorithm.^106^

Network graph calculations were performed using the python package NetworkX.^107^

Using the constructed protein structure networks, we computed the residue-based betweenness parameter. The short path betweenness centrality of residue *i* is defined to be the sum of the fraction of shortest paths between all pairs of residues that pass through residue *i* :

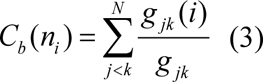

where *g _jk_* denotes the number of shortest geodesics paths connecting *j* and *k,* and *g _jk_* (*i*) is the number of shortest paths between residues *j* and *k* passing through the node *n_i_* .

The Girvan-Newman algorithm^108–110^ is used to identify local communities. An improvement of Girvan-Newman method was implemented where all highest betweenness edges are removed at each step of the protocol. The algorithmic details of this modified scheme were presented in our recent study.^111, 112^ The network parameters were computed using the python package NetworkX^107^ and Cytoscape package for network analysis.^113, 114^ A community-based analysis and modularity assessment of allosteric interaction networks is based on the notion that groups of residues that form local interacting communities are expected to be highly correlated and can switch their conformational states cooperatively. As a result, long-range allosteric interaction signals can be transmitted through a hierarchical chain of local communities on the dynamic interaction networks. The community analysis and a comparative evaluation of the communities in the SARS-CoV-2 S-D614 and S-G614 structures are used as proxy for measuring global protein stability and communication efficiency in performing allosteric functions. This analysis provides also a global metric to measure changes in connectivity and interaction between subdomains and inter-protomer association in the SARS-CoV-2 S structures.^115^

## Results and Discussion

### Conformational Mobility and Functional Dynamics Profiles of the SARS-CoV-2 Spike Proteins: The D614G Mutation Induces Subtle Perturbations of Collective Motions

We employed multiple CG-CABS simulations followed by atomistic reconstruction and refinement to provide a comparative analysis of the dynamic landscapes characteristic of the major functional states of the SARS-CoV-2 S trimer. While all-atom MD simulations with the explicit inclusion of the glycosylation shield could provide a rigorous assessment of conformational landscape of the SARS-CoV-2 S proteins, such direct simulations remain to be technically challenging and computationally demanding due to the gigantic size of a complete SARS-CoV-2 S system embedded onto the membrane.^56–59^ To partially overcome these computational challenges, we combined CG-CABS simulations with atomistic reconstruction and additional optimization by adding the glycosylated microenvironment for 22 most occupied N-glycan sites as described in Materials and Methods.

Using CABS-CG simulations, we examined how the D614G mutation could affect the interdomain contacts and global dynamic profiles of the closed and partially open states (Figure 2). The presented analysis characterized the intrinsic dynamics of the SARS-CoV-2 S proteins in the native and D614G mutant states, particularly probing the propensity of the S-D614G mutant protein for altering dynamic signatures and facilitating movements towards the open conformation. The conformational dynamics profiles of the SARS-CoV-2 S-GSAS/D614 (Figure 2A,B) and SARS-CoV-2 S-GSAS/G614 (Figure 2C,D) showed a more flexible S1 subunit and very stable S2 subunit in both closed and 1-up open states. As expected, the thermal fluctuations of both S1 and S2 regions were smaller in the closed state with RMSF < 1.0 Å.

**Figure 2.**
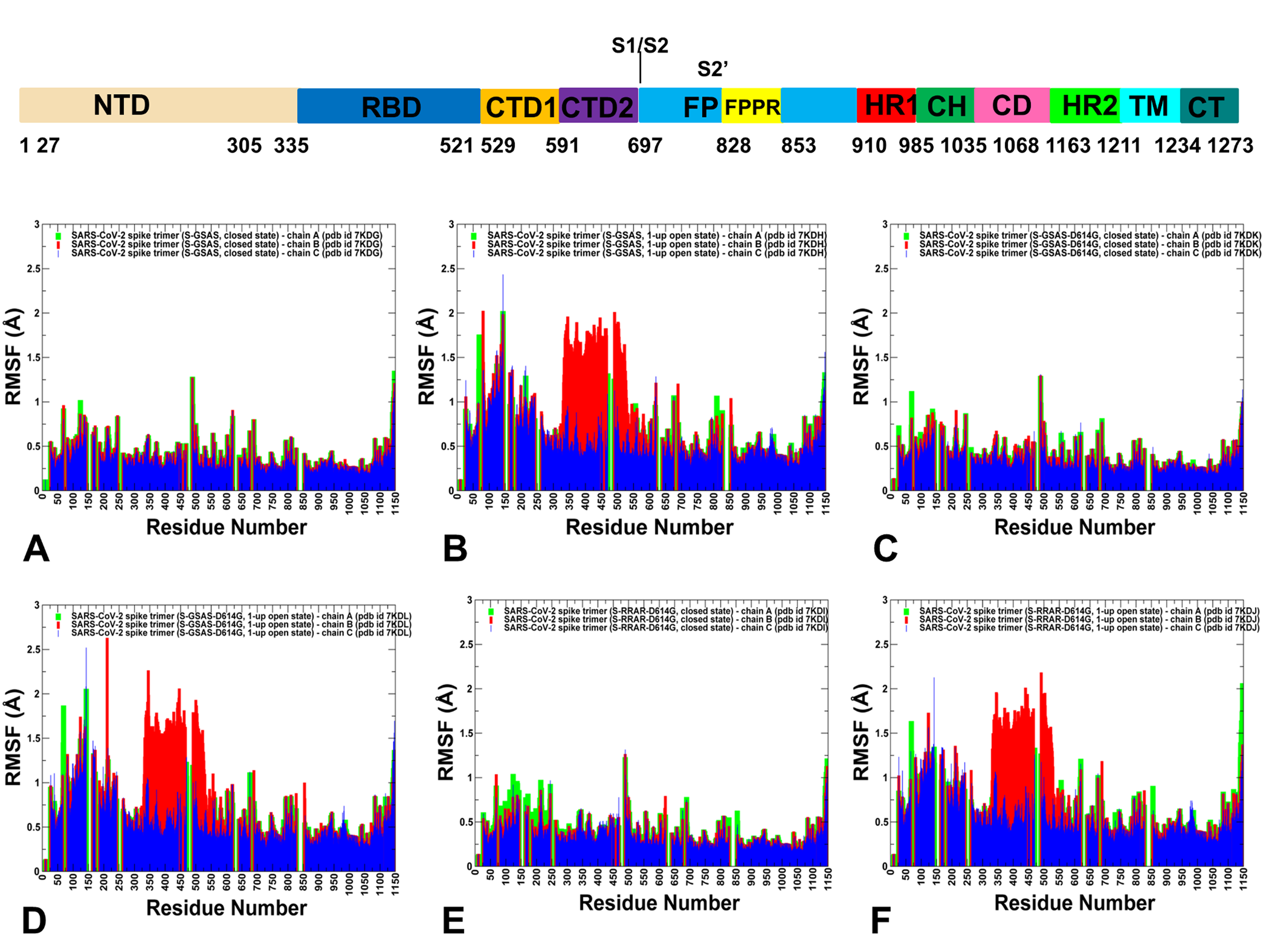
CABS-GG conformational dynamics of the SARS-CoV-2 spike (S) protein trimer mutants. A schematic representation of domain organization and residue range for the full-length SARS-CoV-2 S protein is shown above conformational dynamics profiles. The S1 regions include NTD (residues 14-306), RBD (residues 331-528), CTD1 (residues 528-591), CTD2 (residues 592-686), UH (residues 736-781), HR1 (residues 910-985), CH (residues 986-1035), CD (residues 1035-1068), HR2 (residues 1069-1163). (A) The root mean square fluctuations (RMSF) profiles obtained from CABS-CG simulations of the cryo-EM structure of SARS-CoV-2 S-GSAS/D614 in the closed-all down state (pdb id 7KDG). (B) The RMSF profiles for S-GSAS/D614 in the 1 RBD-up open state (pdb id 7KDH). (C) The RMSF profiles for the cryo-EM structure of SARS-CoV-2 S-GSAS/G614 mutant ectodomain in the closed form (pdb id 7KDK). (D) The RMSF profiles for the cryo-EM structure the SARS-CoV-2 S-GSAS/G614 mutant in the 1 RBD-up open states (pdb id 7KDL). (E) The RMSF profiles for the cryo-EM structure of SARS-CoV-2 S-RRAR/G614 mutant in the closed form (pdb id 7KDI). (F) The RMSF profiles for the cryo-EM structure of SARS-CoV-2 S-RRAR/G614 mutant in the open form (pdb id 7KDJ). The profiles for protomer chains A,B and C are shown in green, red and blue bars respectively.

In the closed state, the RBD (residues 331-528) and CTD1 (residues 528-591) corresponded to the most stable regions in the S1 subunit, while UH (residues 736-781) and CH (residues 986-1035) were the most stable regions in the S2 subunit (Figure 2A). Only marginally larger fluctuations were seen in the CTD2 region (residues 592-686) that connects S1 and S2 subunits. Our analysis showed that the conformational dynamics profiles were generally similar for the closed forms of both the SARS-CoV-2 S-GSAS/D614 (Figure 2A) and S-GSAS/G614 mutant (Figure 2C). Molecular simulations did not detect any radical changes in the dynamics profile of the D614G mutant, indicating that the effect of mutation on the conformational dynamics could be subtle and amount to a number of small local changes dispersed throughout the protein structure. This suggests that a more detailed and granular analysis of the residue interaction networks can be warranted to systematically explore and quantify the residue-based changes in protein stability. CABS-CG dynamics revealed a moderate increase in the conformational mobility of both D614 and D614G open states of the SARS-CoV-2 S protein (Figure 2B, D). The single up protomer in both the S-D614 S-G614 forms can undergo larger fluctuations up to RMSF =2.0-2.3 Å but the overall dynamic profile remained remarkably similar in the native and mutant open forms. Of some notice were moderately greater residue displacements in the NTD regions of the S-D614 and S-G614 open structures, including the NTDs of the two closed-down protomers (Figure 2B, D). Similar conformational dynamics profiles were obtained for the S-RRAR/G414 structures in the closed and open states (Figure 2E, F). Several previous studies suggested that the molecular basis of the increased infectivity for the D614G mutant may be explained via “openness” hypothesis^50–52^ according to which D614G modification can alter conformational dynamics signatures of the S protein and promote switching to the open conformation, positioning a single or multiple RBDs to make contacts with the ACE2 receptor. Our results showed no evidence of substantial alteration of the dynamic signatures caused by the D614G mutation, showing similar conformational mobility profiles for the open states of the S-D614 and S-G614 proteins.

To provide a more detailed analysis of the intra- and inter-protomer contacts in the closed and open states, we computed the ensemble-averaged numbers of these contacts for the SARS-CoV-2 S-GSAS/D614 and SARS-CoV-2 S-GSAS/G614 structures (Tables S1-S6) using a contacts-based Prodigy approach for prediction of binding affinity in protein–protein complexes and distance threshold of 5.5 Å to define an inter-molecular contact .^116, 117^ This analysis showed that the number of inter-protomer contacts is consistently greater in the closed state of the S-D614 and S-G614 proteins as compared to the partially open state. According to this assessment, D614 and Q613 residues can anchor the intra-protomer clusters with V597, S596 and T315 of the same protomer, while establishing contacts with T859, L861 and S735 residues of the adjacent protomers (Figure 1, Tables S1-S6). It is worth noting that in agreement with several computational studies advocating for the “openness” mechanism^50^ we also found a preferentially symmetric nature of the inter-protomer contacts in the closed state, while the number of inter-protomer contacts can be considerably reduced in the partially open form of S-D614 conformation (Tables S1,S2). These changes could become less pronounced in the S-G614 form, but the total number of the inter-protomer contacts is consistently reduced in the open state regardless of the mutational status of D614 position (Tables S3-S6). A close inspection of the local dynamics profiles revealed only minor flexibility changes near the mutational site, where the loss of favorable interactions with T859 of the adjacent protomer may be compensated through the intra-protomer contacts especially for the up protomer in the open state (Figure 2). It was proposed that a more flexible closed state would favor the opening of the S-D614 spike protein, while a more rigid open state of D614G would shift the conformational equilibrium towards the open state.^52^ Our results revealed small and largely synchronous dynamic changes in the closed and open forms of the D614G mutant, showing no indication of a dramatic alteration of the dynamic signatures to suggest an obvious trigger for the dynamic preferences of the D614G mutant towards partially open form as was proposed in a computational study.^52^ Based on these preliminary results, we suggested a hypothesis that the D614G substitution may cause a cascade of small and subtle changes in both the intra- and inter-protomer interactions that may be difficult to directly discern from conformational dynamics analysis. Consequently, a more rigorous analysis of collective motions and dynamic characterization of rearrangements in the residue interaction networks for the S-D614 and S-G614 proteins could be warranted to provide a more quantitative insight into the mechanism.

The experimental studies indicated that the D614G mutational site is located in the immobilized structural region of the SD2 domain where local environment of D614 combined with β strand formed by residues 311–319 may correspond to a hinge center governing motions of NTD and RBD, as well as isolating the motions in S1 from the S2 subunit.^44^ To identify hinge sites and characterize collective motions in the SARS-CoV-2 S-D614 and SARS-CoV-2 S-G614 structures, we performed PCA of trajectories derived from CABS-CG simulations and also determined the essential slow modes using ENM analysis.^85, 86^

The reported ENM-based functional dynamics profiles were averaged over the first three major low frequency modes (Figure 3A,B). The conserved hinge regions in the closed and open forms of the S-D614 protein that can regulate the inter-domain movements between RBD and NTD as well as the relative motions of S1 and S2 regions are localized at F318, S591, F592, V539, L570, I572 and Y855 residues (Figure 3C,D). Strikingly, these sites are located in the close proximity of the Q613 and D614 residues and could form several interaction clusters in both the closed and open states of the SARS-CoV-2 S-D614 and SARS-CoV-2 S-G614 structures (Figure 3C,D). Notably, L570, I572, Y855, I856, and S591 hinge residues are situated near the inter-domain SD1-S2 interfaces and could act collectively as regulatory switch centers governing the population shifts between closed and open forms. The central finding of this analysis is the unique role that D614 and Q613 sites could play in anchoring and integrating major hinge residues in regulatory interacting centers that may coordinate global motions in the SARS-CoV-2 S protein and govern allosteric structural changes between the closed and open forms (Figure3). The shape of the essential mode profiles is mainly determined by the topology of the protein fold and remained conserved between the closed forms of S-GSAS/D614 structure (Figure 3A) and S-GSAS/G614 structure (Figure 4A), highlighting the key role of these hinge clusters in modulating NTD/RBD displacements and functional motions of the S1 subunit with respect to the largely immobilized S2 subunit. The slow mode shapes in the closed forms of S-GSAS/D614 and S-GSAS/G614 reflected a general symmetrical nature of the closed S trimer as the essential mobility profiles are largely identical for all protomers showing minor movements of the RBD regions and moderate displacements of the NTDs.

**Figure 3.**
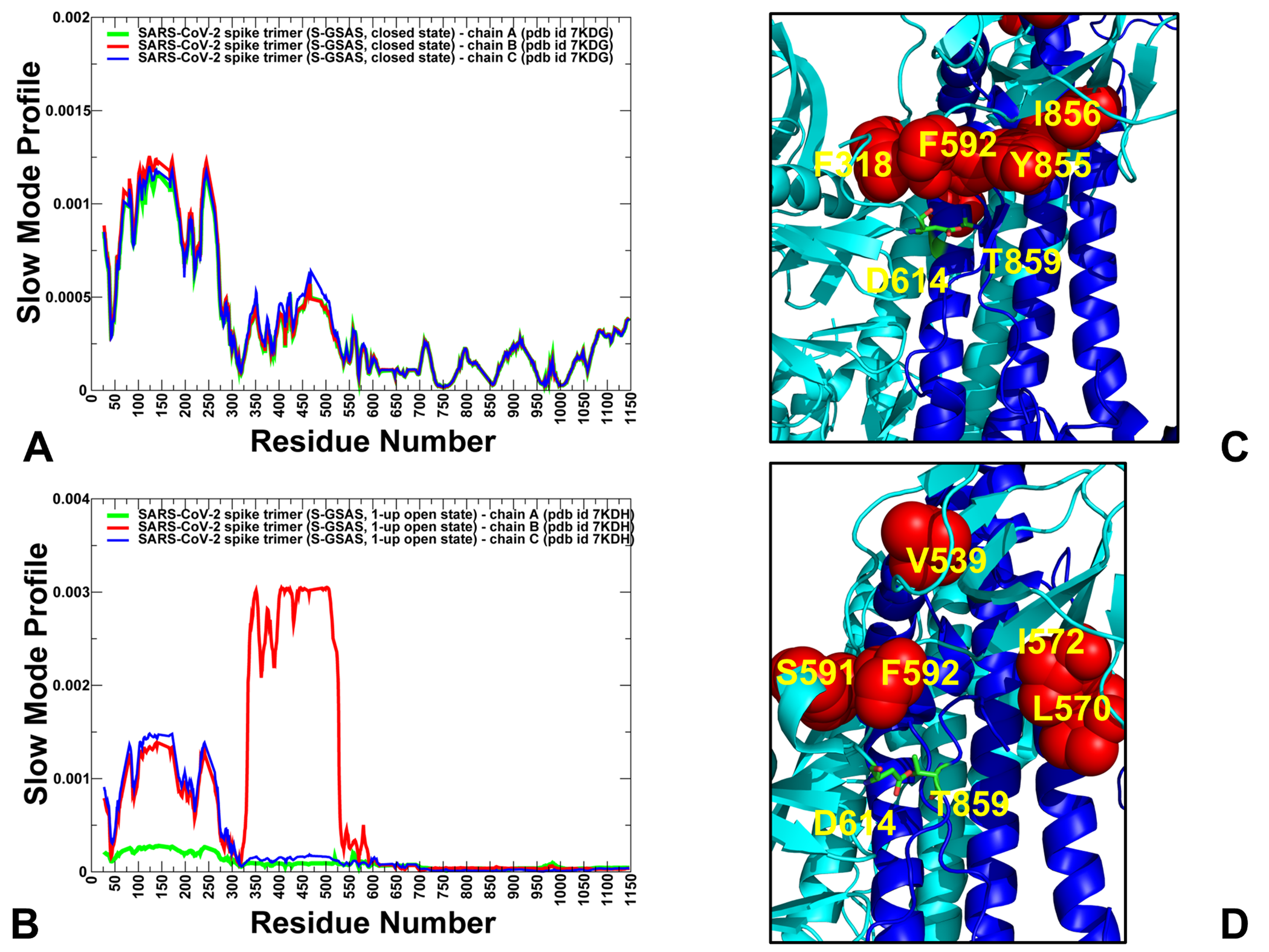
Collective dynamics of the SARS-Cov-2 S-GSAS/D614 structures in the closed and open forms. The mean square displacements in functional motions are averaged over the three lowest frequency modes. (A) The essential mobility profiles for the SARS-Cov-2 S-GSAS/D614 structure in the closed form (pdb id 7KDG). (B) The slow mode profile for the SARS-Cov-2 S-GSAS/D614 structure in the 1-up open state (pdb id 7KDH). The profiles for protomer chains A,B and C are shown in green, red and blue lines respectively. (C,D) Structural maps of the hinge region clusters in the SARS-CoV-2 S-GSAS/DS614 structure. The major hinge sites are shown in red spheres and mapped onto a closed-down protomer of SARS-CoV-2 S-GSAS/DS614 structure (pdb id 7KDG). The key hydrogen bonding between D614 (protomer A) and T589 (protomer B) is shown with D614 and T589 in atom-colored sticks. The close proximity of the salt bridge to the network of hinge centers is highlighted.

**Figure 4.**
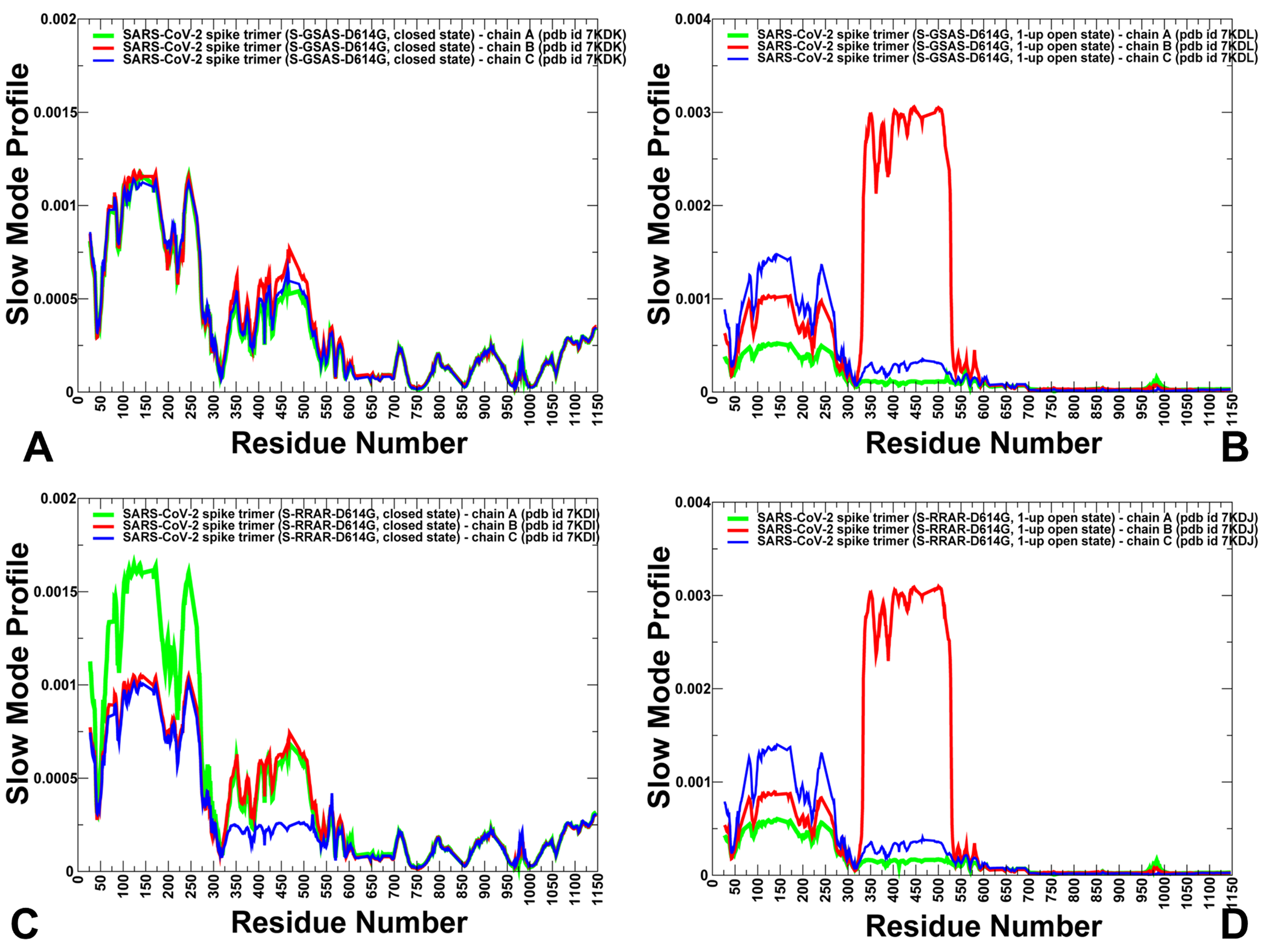
Collective dynamics of the SARS-Cov-2 S-GSAS/G614 mutant structures in the closed and open forms. The mean square displacements in functional motions are averaged over the three lowest frequency modes. (A) The essential mobility profiles for the SARS-CoV-2 S-GSAS/G614 structure in the closed form (pdb id 7KDK). (B) The slow mode profile for the SARS-Cov-2 S-GSAS/G614 structure in the 1-up open state (pdb id 7KDL). (C) The essential mobility profiles for the SARS-CoV-2 S-RRAR/G614 structure in the closed form (pdb id 7KDI). (D) The slow mode profile for the SARS-CoV-2 S-RRAR/G614 structure in the 1-up open state (pdb id 7KDJ). The profiles for protomer chains A,B and C are shown in green, red and blue lines respectively.

Interestingly, the functional dynamics of the closed form for the cleaved S-RRAR/G614 mutant structure revealed breaking of symmetry between protomers, showing the larger and asynchronous displacements of the NTD and RBD regions for two protomers while highlighting a partial immobilization of the S1-RBD and S2 regions for the third protomer (Figure 4C). This observation indicated that the D614G mutant may perturb hinge clusters and alter functional movements in the closed S-RRARS trimer, potentially priming protomers in the S protein for transition to the open form. These findings showed an agreement with previous studies suggesting the increase in the open state preferences for the D614G mutation due to the increased flexibility of the closed state and the enhanced rigidification of the open form.^52^ Indeed, while local conformational mobility profiles are similar for the native S-D614 and mutant S-G614 proteins, the subtle differences may emerge in the functional dynamics patterns that pointed to the increased global mobility of the S1 regions in the closed state. Accordingly, the D614G mutation may partly destabilize collective motions in the closed form and consequently alter the stability of the residue interaction networks in the S-RRARS trimer. The significance of these observations stems from the fact that local structural and interaction changes near the D614 position could modulate stability of key hinge clusters and thereby incur a significant effect on functional dynamics and allosteric transitions between closed and open forms. At the same time, we observed that slow mode profiles for the open form in the S-GSAS/D614 (Figure 3B), S-GSAS/G614 (Figure 4B) and S-RRAR/G614 mutants (Figure 4D) are similar, featuring movements of the single RBD-up protomer and moderate displacements of the NTD regions. Hence, we conclude D614G mutation may exert its dynamic effect through preferential modulation of collective movements in the closed form and promoting transition to the open forms rather than altering dynamic signatures of the open spike states.

### Mutational Sensitivity Analysis in the SARS-CoV-2 Spike Trimers Reveal D614G-Induced Differential Stabilization of the Closed and Open States

Previous mutagenesis analysis suggested that the D614G mutant may improve the stability of the S protein by strengthening the inter-protomer association between S1 and S2 regions.^53^ We employed the equilibrium ensembles generated by CABS-CG simulations of the SARS-CoV-structures to perform alanine scanning of the protein residues and mutational sensitivity analysis of the S-D614 and S-G614 proteins at the mutational site (Figure 5). The primary objective of this energetic analysis was to further test the shedding hypothesis suggesting that D614G may lead to stabilization of the S protein and block a premature dissociation of the S1 subunit, leading to the increased number of functional spikes and stronger infectivity.^48^ The important questions addressed in this analysis are (a) whether the D614G mutation can incur a similar or differential effect on the closed and open forms of the S protein and (b) if there are differences in the D614G-induced energetic effects in the S-GSAS and S-RRAR structures.

**Figure 5.**
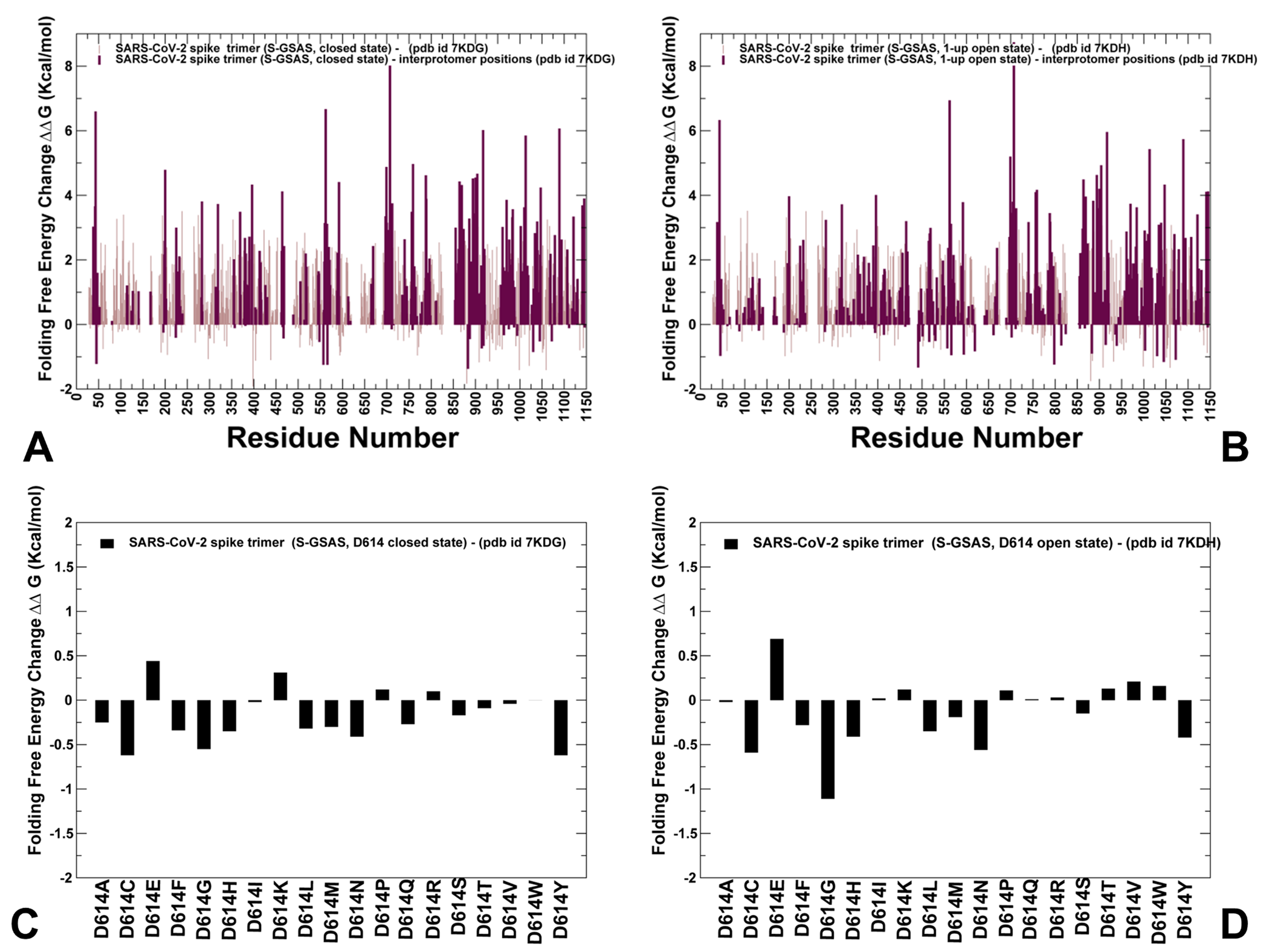
Alanine scanning of the protein residues and mutational sensitivity analysis of D614 site in the SARS-CoV-2 S trimers. (A) The residue-based protein stability changes upon alanine mutations in the closed form of the S-GSAS/D614 protein (pdb id 7KDG). (B) The residue-based protein stability changes upon alanine mutations in the open form of the S-GSAS/D614 protein (pdb id 7KDH). The protein stability changes upon alanine mutations are shown in light brown bars and the respective stability changes for residues involved in the inter-protomer contacts are shown in maroon-colored bars. The computed values were averaged over samples from molecular simulations. (C) Mutational sensitivity scanning of D614 position in the closed form of the S-GSAS/D614 protein (pdb id 7KDG). (D) Mutational sensitivity scanning of D614 position in the open form of the S-GSAS/D614 protein (pdb id 7KDH). The protein stability changes are shown in black filled bars.

Through FoldX-based alanine profiling of the protein residues in the SARS-CoV-2 S-D614 and S-G614 structures, we probed the hypothesis that hinge centers may correspond to the energetic hotspots. We also examined and highlighted stability of the residues involved in the inter-protomer contacts. The alanine scanning across the studied SARS-CoV-2 S structures revealed the presence of multiple energetic hotspots that are broadly distributed and generally are organized into local clusters (Figure 5). As may be expected, the majority of the energetic hotspot clusters were observed near the positions involved in both the intra- and inter-protomer contacts, including the RBD residues and CTD1 domain (residues 529-591). Most of the energetic hotspot clusters are localized in the rigid S2 subunit regions, including the UH regions (residues 736-781), HR1 (residues 910-985), and CH regions (residues 986-1035).

Notably, a number of the energetic hotspots were aligned with hinge positions F318, S591, F592, L570, I572, F759, and I788 where alanine mutations produced considerable destabilization changes (>3 kcal/mol). To gain a more detailed characterization of mutational sensitivity at the D614 position, we conducted a full scanning profile of the SARS-CoV-2 S-D614 and SARS-CoV-2 S-G614 structures in both closed and open states (Figure 5C,D). Interestingly, the mutational sensitivity profile of the SARS-CoV-2 S-D614 in the closed form showed moderate stabilization changes for majority of substitutions except D614E and D614K (Figure 5C). Interestingly, the D614C, D614G, and D614N mutations result in an appreciable and similar stabilization of the closed form. More considerable stabilization changes of ∼1.2 kcal/mol were induced by the D614G mutation in the open form (Figure 5D). Notably, the D614G mutation caused the largest energetic change as compared to other substitutions. Interestingly, we also observed that the D614G and D614N mutations could lead to the favorable and comparable stabilization changes in both the closed and open forms of the S-GSAS/D614 protein (Figure 5C,D). These findings are consistent with the latest differential scanning fluorimetry studies showing that the D614G and D614N induced changes have a very similar effect, leading to a considerable improvement of thermal stability which may be explained by a decrease in premature shedding of S1 domain.^49^

In line with these findings, mutational sensitivity analysis of the SARS-CoV-2 G614 structures revealed a significant destabilization effect of substitutions in both the closed and open forms (Figure 6). The results particularly highlighted the largest changes induced by the G614D and G614E modifications in the closed (Figure 6A,C) and open forms (Figure 6B,D). Of interest is the fact that destablization changes induced by the G614D and G614E modifications become more pronounced in the S-RRAR/G614 structures (Figure 6C,D) Hence, our data suggested that the local energetic gains caused by D614G can cause stabilization of the S trimer and stronger interactions between S1 and S2 domains that could also manifest in the improved communications in the residue interaction networks in the mutant variant. These results also offer support to the experimental observations that the enhanced stability of the S-D614G mutant may be linked with the mechanism of the reduced S1 shedding.^49^ Importantly, the differential stabilization of the S-G614 closed and open forms showing the energetic preferences of the mutant towards the open state may help to partly reconcile the “openness” and “S1-shedding” mechanism underpinning the D614G effects.

**Figure 6.**
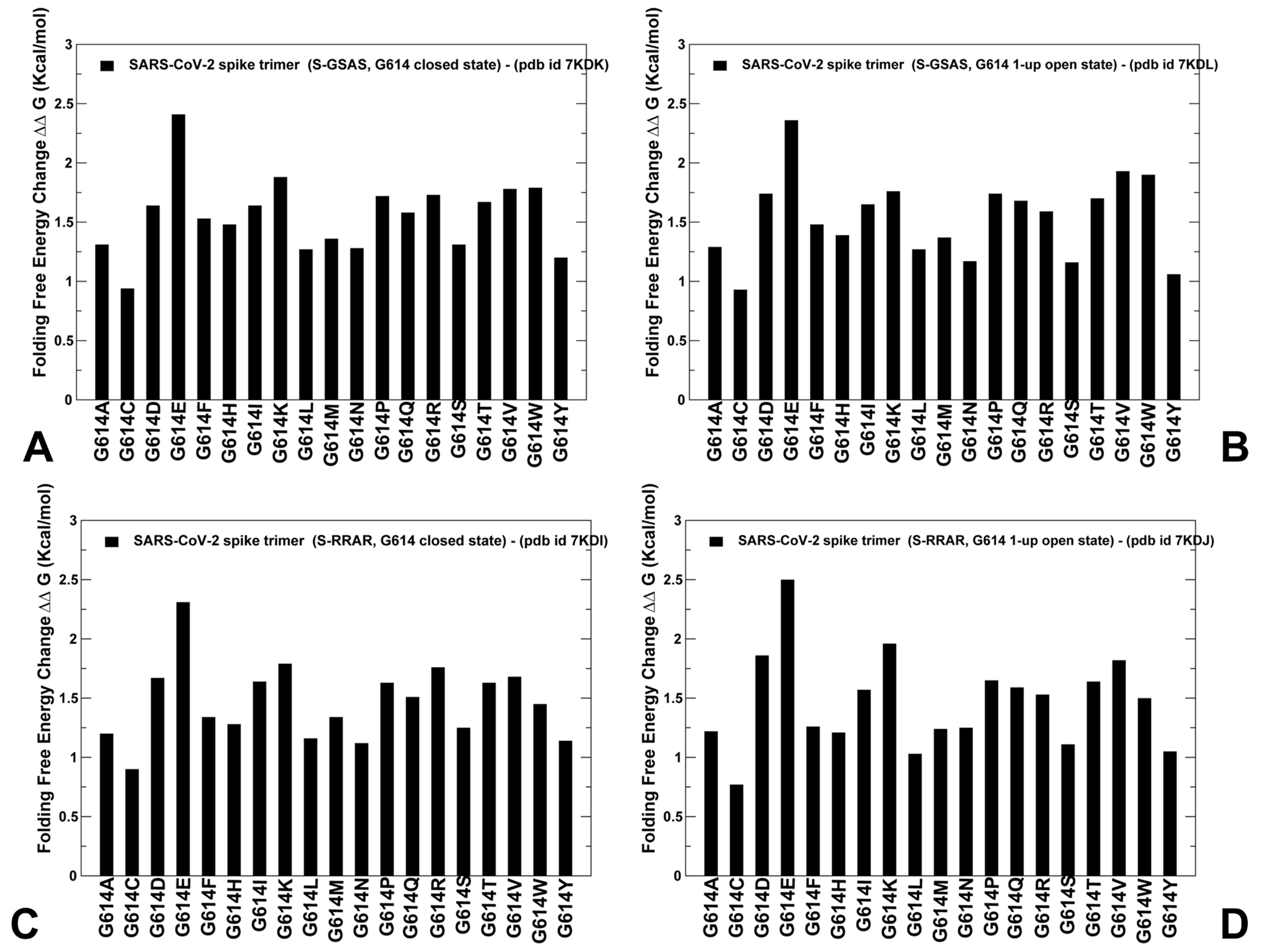
Mutational sensitivity analysis of D614G mutated site in the SARS-CoV-2 S trimers. (A) Mutational sensitivity scanning of G614 position in the closed form of the S-GSAS/G614 protein (pdb id 7KDK). (B) Mutational sensitivity scanning of G614 position in the open form of the S-GSAS/G614 protein (pdb id 7KDL). (C) Mutational sensitivity scanning of G614 position in the closed form of the S-RRAR/G614 protein (pdb id 7KDI). (D) Mutational sensitivity scanning of G614 position in the open form of the S-RRAR/G614 protein (pdb id 7KDJ). The protein stability changes are shown in black filled bars.

### Distance Fluctuation Analysis of the SARS-CoV-2 S Structures Quantifies D614G-Induced Modulation of Stability and Allosteric Communication Propensities

We employed distance fluctuations analysis of the conformational ensembles to probe structural stability and allosteric communication propensities of the SARS-CoV-2 native and D614G mutant forms. The residue-based distance fluctuation indexes measure the energy cost of the dynamic residue deformations and could serve as a robust metric for assessment of stability and allosteric propensities of protein residues.^98, 99^ Structural rigidity of functionally important regions may be reflected in their high fluctuation indexes associated with small fluctuations in the average inter-residue distances. In this model, dynamically correlated residues whose effective distances fluctuate with low or moderate intensity are expected to communicate with the higher efficiency than the residues that experience large fluctuations. According to our studies^98, 99^ the residues with high distance fluctuation indexes may also correspond to structurally stable regulatory positions implicated in coordination of allosteric communication, and abrupt changes between maxima and minima in the profile may point to boundaries between structurally rigid and flexible regions signaling presence of the inter-domain hinge sites.

By using this analysis, we identified allosteric hotspots that are responsible for modulation of stability and allosteric changes. The distance fluctuation profile of the SARS-CoV-2 S-D614 closed form featured a number of high peaks that were distributed in the S1 and S2 subunits (Figure 7A). In particular, a strong peak was seen in the S1-RBD region (residues312-320) that corresponds to the key component of the regulatory hinge site and located in the close structural proximity of the D614 position. Notably, the largest peak was aligned with cluster of residues anchored by the Q613/D614 sites that also included S596, V597 and I598 positions (Figure 7A). Other notable peaks corresponded to residues in the S2 regions including cluster 948-LQDVV-952 in the HR1 region (residues 910-985). According to our latest study of the SARS-CoV-2 S mutant trimers, this hydrophobic center is coupled with the Y855/I856 conformational switch that mediate couplings between the S2 subunit and the RBD regions.^118^ Interestingly, the HR1 region (residues 934-940) is also known to be targeted by naturally occurring mutations in SARS-CoV-2 protein.^119, 120^

**Figure 7.**
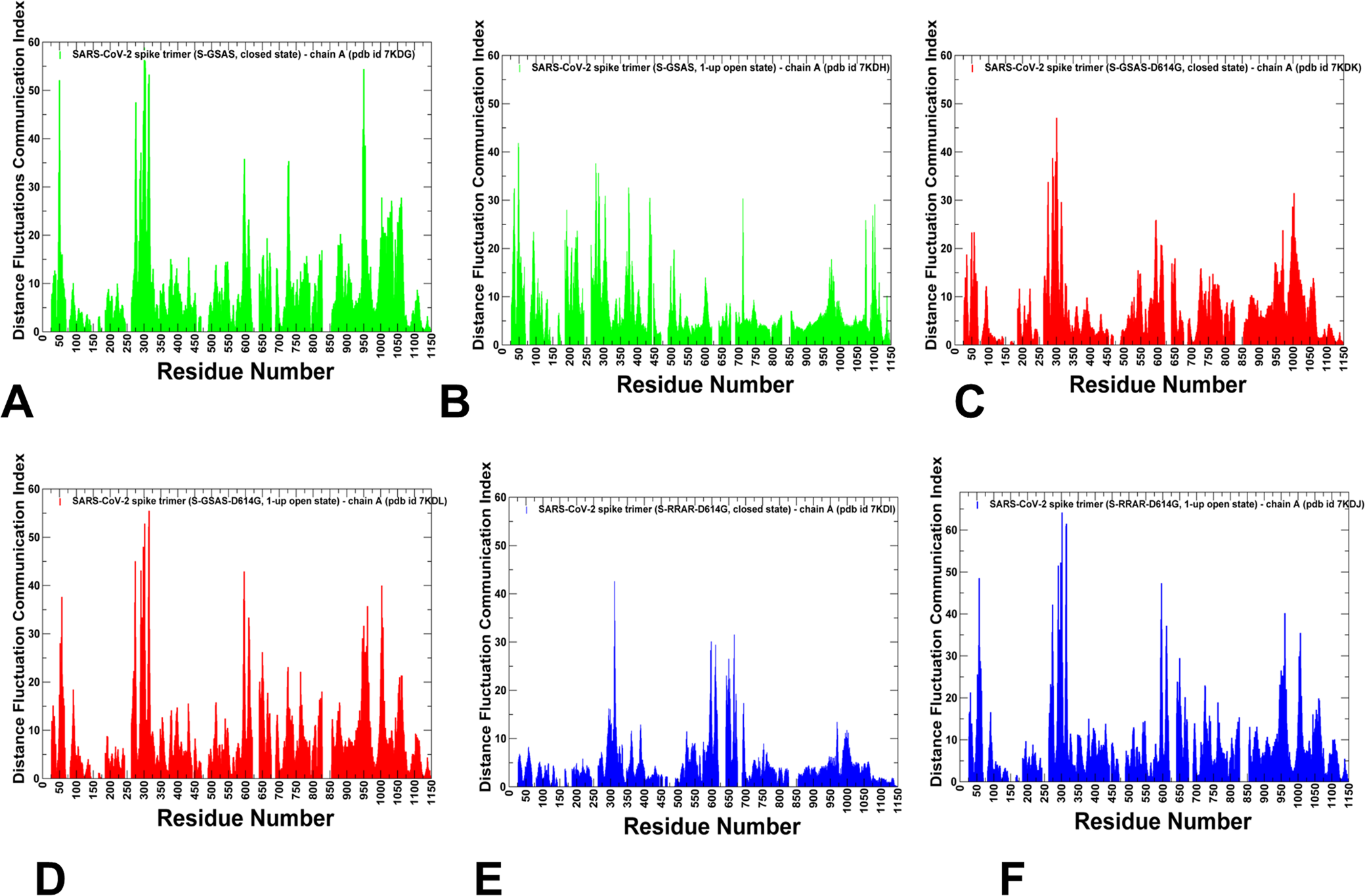
The distance fluctuations analysis of the conformational ensembles in the SARS-CoV-2 S proteins (A) the distance fluctuation communication profile for the S-GSAS/D614 in the closed-all down state (pdb id 7KDG). (B) The distance fluctuation communication profile for the S-GSAS/D614 in the 1 RBD-up open states (pdb id 7KDH). The profiles in A and B panels are shown in green bars. (C) The distance fluctuation communication profile for the S-GSAS/G614 mutant ectodomain in the closed form (pdb id 7KDK). (D) The distance fluctuation communication profile for the S-GSAS/G614 mutant in the 1 RBD-up open states (pdb id 7KDL). The profiles in panels C and D are shown in red bars. (E) The distance fluctuation communication profile for the S-RRAR/G614 mutant in the closed form (pdb id 7KDI). (F) The distance fluctuation communication profile for the S-RRAR/G614 mutant in the open form (pdb id 7KDJ). The profiles in panels E and F are shown in blue bars.

In the open state of the S-D614 spike protein, we observed notable changes in the distance fluctuation profile, particularly indicating the loss of appreciable peaks near the D614 cluster (Figure 7B). Structural mapping of the distance fluctuation profiles further highlighted these changes (Figure 8A,B). Indeed, the density of communication hotspots markedly changed in the open form of S-D614, featuring clusters in the S1 regions but revealing a conspicuous lack of the communication hotspots near the S1-S2 inter-domain regions (Figure 8B). Accordingly, based on this model, the allosteric interaction network in the native S-D614 protein may be stronger in the closed form than in the partially open state. Consistent with the energetic analysis, we observed that the distribution seen in the SARS-CoV-2 S-D614 closed form (Figure 7A) can be partially altered in the S-G614 closed states (Figure 7C,E).

**Figure 8.**
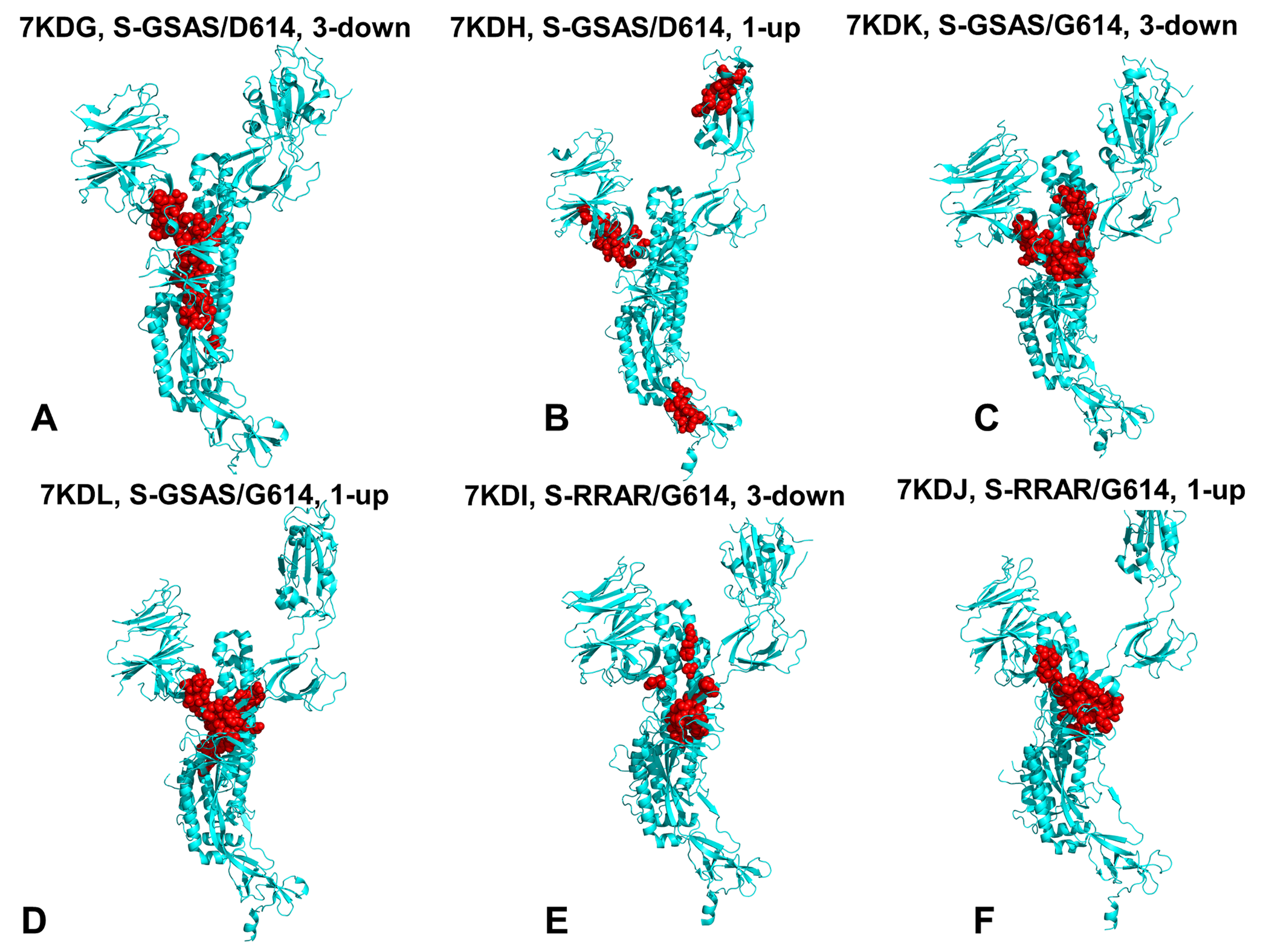
Structural mapping of the distance fluctuations profile hotspots in the SARS-CoV-2 S proteins. (A) Structural mapping of stable communication hotspots for the S-GSAS/D614 in the closed-all down state (pdb id 7KDG). (B) Structural mapping of stable communication hotspots for the S-GSAS/D614 in the 1 RBD-up open states (pdb id 7KDH). (C) Structural mapping of stable communication hotspots in the closed form of S-GSAS/D614 (pdb id 7KDK). (D) Structural mapping of stable communication hotspots for the S-GSAS/G614 mutant in the 1 RBD-up open states (pdb id 7KDL). (E) Structural mapping of stable communication hotspots for the S-RRAR/G614 mutant in the closed form (pdb id 7KDI). (F) Structural mapping of stable communication hotspots for the S-RRAR/G614 mutant in the open form (pdb id 7KDJ). The hotspots are shown in red spheres. The mapping is projected onto a single protomer shown in cyan ribbons.

Indeed, in the closed form of the D614G mutant strong peaks remained in the S1-RBD hinge region (residues 312-320) and near the D614G local cluster, while the density of hotspots in the S2 regions was weakened. Structural mapping illustrated these subtle adjustments (Figure 8A,C), pointing to the increased consolidation of communication hotspots near S1-S2 interface in the S-G614 mutant. These observations suggested that allosteric couplings between S1 and S2 subunits may be partly reorganized in the S-G614 mutant to strengthen stability and communication at the S1-S2 interface. Of special importance are the distribution profiles of the SARS-CoV-2 S-G614 mutant in the open state featuring multiple pronounced peaks in the S1 and S2 regions as well as at the S1-S2 interface (Figure 7D,F) Strikingly, the S-G614 mutant profiles in the open state revealed a strong peak aligned almost precisely with the mutational site (Figure 7D,F). This suggested that the D614G mutation may strengthen the stability and enhance allosteric communication propensities in this region, which could arguably lead to the stronger S1-S2 inter-domain associations and the improved allosteric signaling in the open state. Structural analysis of the distribution peaks (Figure 8D,F) also pointed to a preferential consolidation of the S-G614 communication hotspots in the CTD1 and CTD1-CTD2 regions (528-685). To summarize, the distribution of allosteric hotspots in the S-D614 and S-G614 mutant structures revealed a partial reorganization, showing minor changes between the closed states but more significant alterations in the open states of the S-G614 mutant. The observed reorganization and consolidation of the communication hotspots in the CTD1-CTD2 regions of the S-G614 mutant may promote the greater stability and efficient allosteric couplings between S1 and S2 regions. We argue that D614G mutation-induced modulation of stability and allosteric propensities could therefore limit S1 shedding and favor acquisition of the open state to restore and further optimize allosteric signaling of the SARS-CoV-2 S machinery.

### Network Modeling and Community Analysis Suggest D614G-Induced Reorganization of the Residue Interaction Networks and Improved Allosteric Signaling in the Open States

Mechanistic network-based models allow for a quantitative analysis of allosteric molecular events in which conformational landscapes of protein systems can be remodeled by various perturbations such as mutations, ligand binding, or interactions with other proteins. Using this framework, the residue interaction networks in the SARS-CoV-2 spike trimer structures were built using a graph-based representation of protein structures in which residue nodes are interconnected through both dynamic^103^ and coevolutionary correlations.^99, 104, 105^ Using community decomposition, the residue interaction networks were divided into local interaction modules in which residues are densely interconnected and highly correlated, while the local communities are weakly coupled through long-range allosteric couplings. A community-based model of allosteric communications is based on the notion that groups of residues that form local interacting communities are correlated and switch their conformational states cooperatively. Using network-centric description of residue interactions, we computed the aggregate number of stable local communities and stable hubs in the closed and open forms of the S-D614 and S-G614 mutants (Figure 9).

**Figure 9.**
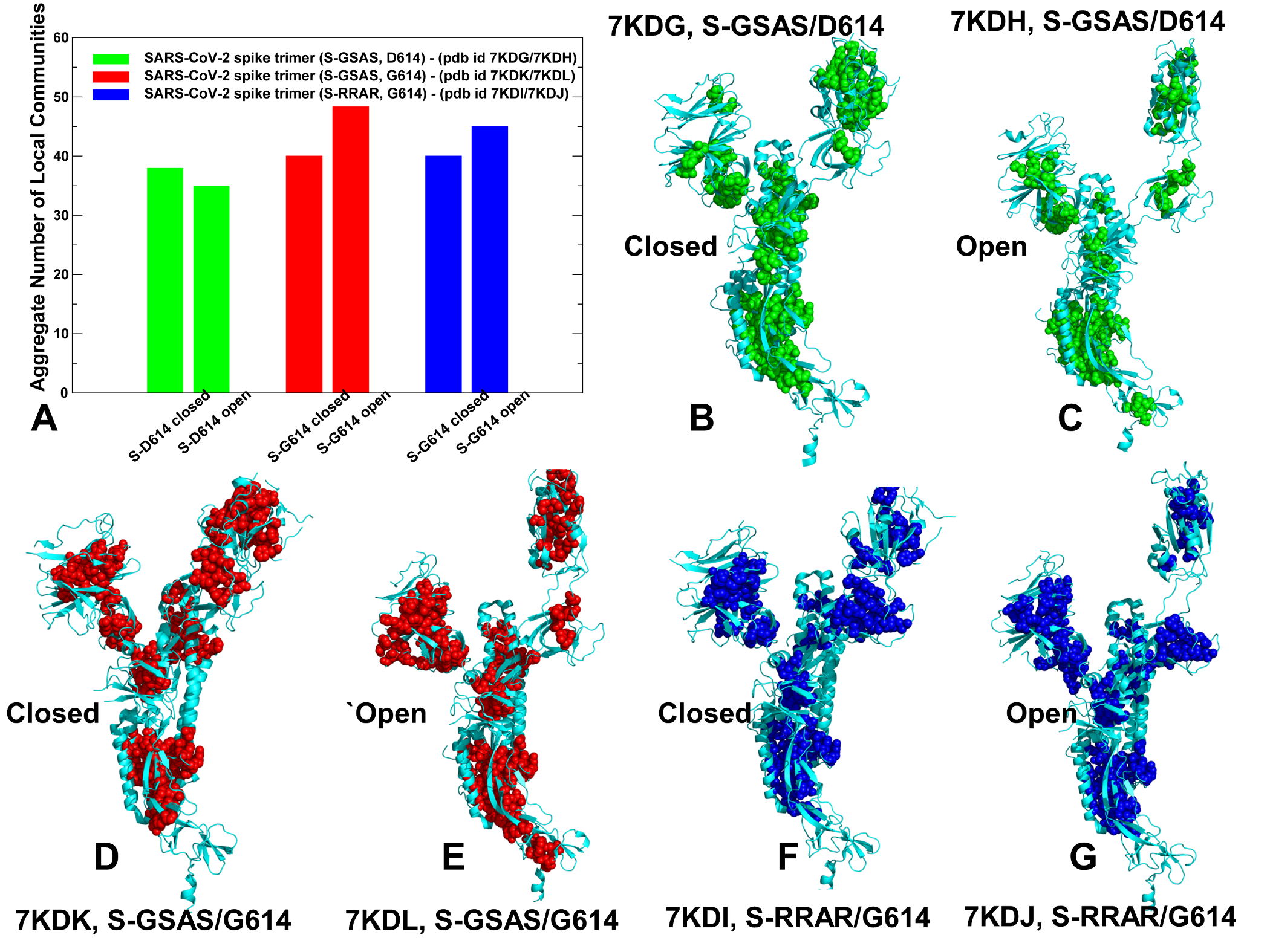
Community analysis and structural community maps in the SARS-CoV-2 S-D614 and S-D614G mutant structures. (A) The quantitative evaluation of the aggregate number of local communities in closed and open forms of S-GSAS/D614 (shown in green bars), closed and open forms of S-GSAS/G614 (red bars), and S-RRAR/G614 (blue bars). (B) Structural mapping of local communities is projected onto a single protomer for the S-GSAS/D614 in the closed-all down state (pdb id 7KDG). (C) Structural mapping of local communities for the S-GSAS/D614 in the open state (pdb id 7KDH). The communities are depicted in green spheres for panels B and C. (D) Structural mapping of local communities in the closed form of S-GSAS/G614 (pdb id 7KDK). (E) Structural mapping of local communities for the S-GSAS/G614 in the open state (pdb id 7KDL). The communities are shown in red spheres for panels D and E. (F) Structural mapping of local communities for the S-RRAR/G614 mutant in the closed form (pdb id 7KDI). (G) Structural mapping of local communities for the S-RRAR/G614 in the open form (pdb id 7KDJ). The communities are in blue spheres for panels F and G. The mapping is projected onto a single protomer shown in cyan ribbons.

In this model, we explored the community analysis as a network proxy for stability assessment. The number of communities for the S-D614 protein was greater in the closed form (Figure 9A). In some contrast, the network analysis of the S-G614 mutant showed a subtle redistribution in the number and allocation of communities as the open state of the mutant harbored more communities and this difference was more pronounced in the S-RRAR/G614 mutant (Figure 9A). Interestingly, the total number of local communities is moderately increased in both closed and open states of the S-G614 mutant. In network terms, this indicated the increased stability of the residue interaction networks in the mutant structures (Figure 9). These results are consistent with the latest experimental data that demonstrated the improved stability of the D614G mutant as compared to the S-D614 protein allowing for reduction in a premature shedding of S1 domain.^49^ A more detailed inspection of local communities in the closed and partially open conformational states of the native S-D614 protein provided some important insight into the proposed mechanisms. We found that not only the total number of local communities is greater in the closed S-D614 state, but also there are important changes in the communities situated in the RBD-CTD1, RBD-CTD2 and CTD1-CTD2 regions (Figure 9A-C). The local modules in these regions that are unique to the closed form included Q314-S596-Q613, I693-V656-Y660, F543-L598-V576 and I666-L650-I670-T645 clusters (Supporting Information, Table S7). In the partially open form, the state-specific communities in these regions included P579-P330-N544, F541-L552-I587 and R328-F543-P579 (Supporting Information, Table S7). Structural mapping of local communities in the S-D614 states illustrated subtle differences in the distribution and density of stable modules (Figure 9B,C). Characteristically, the key community in the closed form of the S-D614 protein is anchored by Q613, which is the immediate neighbor of D614, forming a tight stable cluster with Q314 in the NTD and S596 in CTD2 (Figure 9B,C). A general comparison of structural maps indicated the better connectivity of local communities in the closed form of S-D614 forming a broad network linking the S2 regions with the NTD and RBD regions (Figure 9B). In addition, a number of unique communities are localized in the CTD2 region (I693-V656-Y660 and I666-L650-I670-T645), suggesting the stronger S1-S2 interfacial interactions and tighter packing between S1 and S2 domains in the closed form of the S-D614 protein.

Notably, we also detected the larger number of communities in the NTD and RBD S1 regions in the closed form, indicating that the allosteric interaction network in the S1 and S1-S2 regions is stronger in the closed form, ensuring efficient signaling between S1 and S2 regions in protecting all-down closed form in the S-D614 protein. A somewhat weaker inter-connectivity of local communities near S1-S2 interfaces and in the NTD/RBD regions was observed in the open state of S-D614. These findings suggested that the stability and efficiency of allosteric communication in the residue interaction network could favor the closed-down form of the S-D614 spike protein. Importantly, the community analysis revealed the increased number of stable modules for the S-G614 mutant in both closed and open forms (Figure 9A, Supporting Information, Tables S9,S10). Both forms of S-G614 mutant featured the appreciable number of communities in the CTD1 and CTD2 regions involved in stabilization of the S1-S2 interfaces (Figure 9D,E). These inter-domain local communities bridging S1 and S2 regions included T315-T299-V597, R328-S530-Q580, Q314-S596-Q613, I664-I312-I598, L611-P6645-L650, and F898-F802-F797. The stabilizing communities in the S1-S2 interfaces appeared to strengthen stability of both closed and open S-G614 forms which is consistent with the experimentally observed increased in protein stability of the D614G mutant.^49^ However, the balance in the number of communities may be shifted towards the open state of the S-G614 mutant, showing a more significant increase of local stable modules and promoting the preferential stabilization of the open mutant form. The key community near mutational site Q314-S596-Q613 is uniquely present only in the open form of the S-G614 mutant, likely pointing to state-specific rearrangements of stable interactions induced by the D614G mutation. The important subtle rearrangements were found in the local communities localized in the N2R linker region (residues 306–334) that connects the NTD and RBD regions stacking against the CTD1 and CTD2 domains. In the favorable closed form of S-D614 protein, these modules include I326-V534-V539, R328-L533-D578, R328-S530-Q580, T299-T315-V597 and Q314-S596-Q613 (Figure 9B). Interestingly, the open form of the S-G614 mutant featured some of these modules (T299-T315-V597 and Q314-S596-Q613) and several additional N2R communities unique to this state (R328-F543-P579, P579-P330-N543, F338-F342-L368) that strengthen connections of the N2R linker with the NTD and RBD regions. This local subnetwork may be instrumental in bridging the individual domains in the S1 subunit and ensure a more efficient inter-connectivity between S2 regions, N2R and S1-RBD regions in the S-G614 mutant. The observed difference in the total number of local modules favoring the open mutant state is largely due emerging communities in the NTD and RBD regions (Figure 9D,E). Indeed, while the open form of the S-D614 spike protein is characterized by mobile NTDs that could provide flexible access to the ACE2 receptor, the increased stabilization of the NTD and RBD regions in the S-G614 mutant may accompany strengthening of the S1-S2 interdomain interactions and could contribute to a decrease in premature shedding of S1 domain. Hence, D614G mutation may also exert allosteric effect by partly immobilizing the distal NTD and RBD regions. These findings support the previously suggested notion that the D614G mutation in the SD2 domain could allosterically strengthen stability of the distal NTD and RBD regions in the open state^44, 45^ and therefore potentially promote exposure to the host receptor and greater infectivity.

These findings provide an interesting explanation supporting the reduced shedding hypothesis. Indeed, it has been previously speculated that the loss of hydrogen bonding interactions between D614 in S1 and T859 in S2 caused by D614G mutation may promote rather than limit shedding of the S1 domain.^47^ It was also indicated that D614-T859 protomer-protomer hydrogen bonding may be of critical importance for the integrity and stability of the trimer, and that the D614G mutation could weaken the interaction between the S1 and S2 units, facilitating the shedding of S1 from membrane-bound S2. Our results are more consistent with an alternative explanation and latest experimental data^46^ by showing that the D614G mutation may strengthen the stability of the local community Q314-S596-Q613 and promote hydrogen bonding interactions between Q613 and T859 of adjacent protomer afforded by the local backbone flexibility at the mutational site. The presented results also partly reconciled several scenarios offered to explain functional effects of the D614G mutation. Indeed, the community network analysis suggested that preferential stabilization of the open mutant form can be determined by two main factors : (a) through strengthening of the S1-S2 interactions and (b) by reducing functional movements of the NTD and RBD regions exposed to binding with the host receptor. Accordingly, the D614G mutation may exert its effect through allosteric stabilization of the S1-S2 interfaces limiting shedding of the S1 domain and by reducing flexibility and enhancing thermodynamic preferences of the open state that is central aspect of the “openness”-based mechanistic scenario.

We also determined the distribution of local hubs in the SARS-CoV-2 S-D614 and S-G614 mutant structures using this analysis as proxy for identifying hotspots of allosteric communication in the protein system. The hub centers are instrumental in regulation of allosteric couplings and signal transmission in the protein structure network and are often used to characterize the inter-community connectivity and mediating centers of allosteric interaction networks. The evaluation of the total number of state-specific hubs revealed a picture consistent with the community analysis, showing that the number of unique hubs in the open forms of the S-G614 mutant is moderately greater than in the closed mutant form (Figure 10A). Notably, the closed and open forms of the S-D614 and S-G614 structures can share a number of conserved hub positions that are largely determined by the topology of the spike trimer (Figure 10B,C). Interestingly, these common hubs are very similar to the conserved network hubs reported for cryo-EM structures of the SARS-CoV-2 spike S trimer in the closed state (K986P/V987P,) (pdb id 6VXX) and open state (pdb id 6VYB).^115^ Indeed, among conserved hub positions there are a number of residues proximal or directly aligned with the hinge sites shared by closed and open forms including T315, F318, R319, R328, V539, F592, and Y612 located near the D614 mutational site (Figure 10B,C). Interestingly, several of these common hubs (T315, F318, R319, and R328) belong to the functionally important for communication N2R linker region (residues 306–334) that connects the NTD and RBD regions with the CTD1 and CTD2 domains. Structural maps of common hubs in the S-D614 and S-G614 proteins showed that a significant number of these hotspots occupy conserved regions of the S2 subunit, particularly UH region (residues 736-781), HR1 (residues 910-985), and CH (residues 986-1035) (Figure 10B,C). This is consistent with structural and evolutional conservation of these regions that remain largely preserved in their native positions in the prefusion conformations.

**Figure 10.**
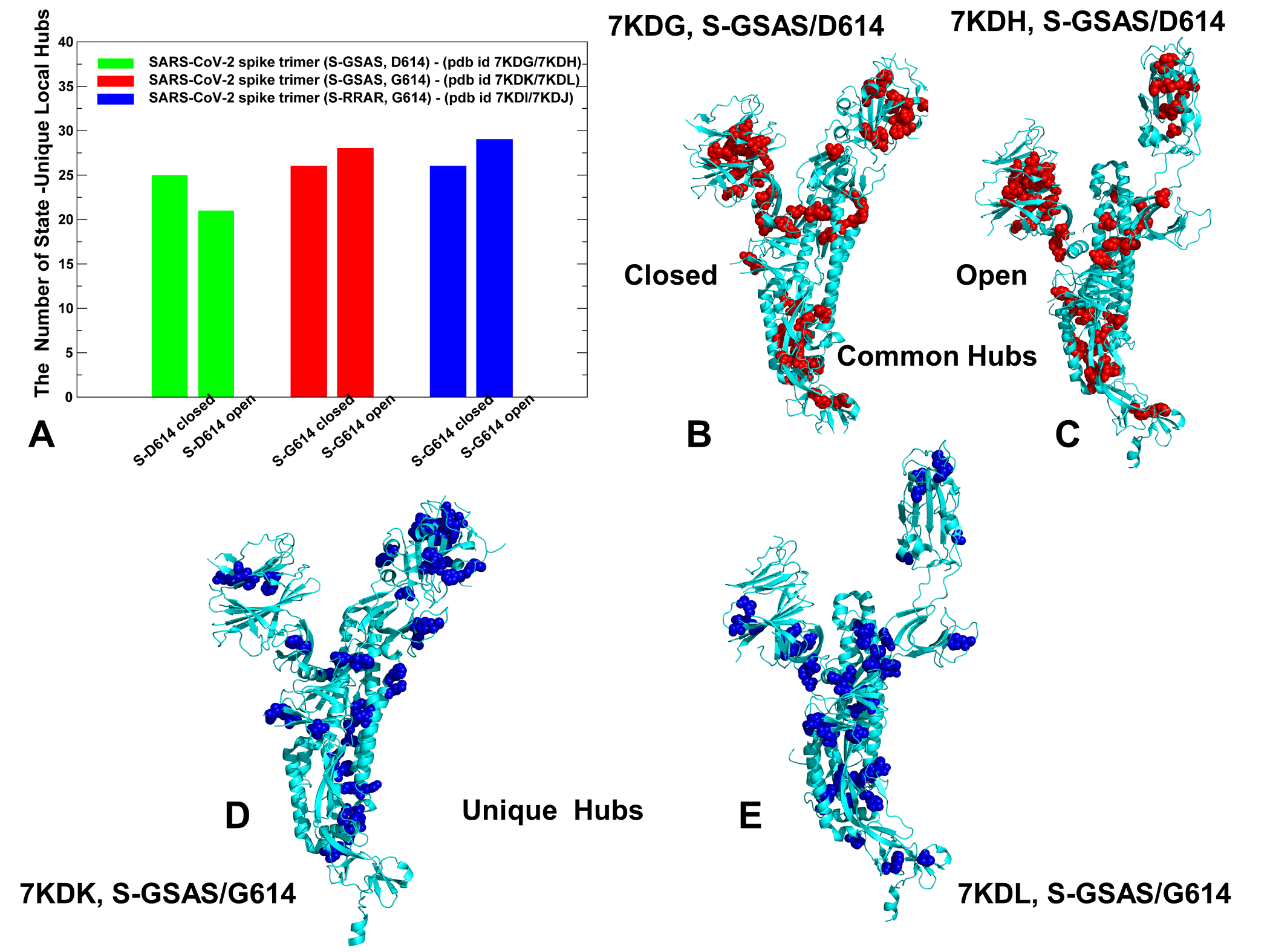
Network hub analysis of the SARS-CoV-2 S-D614 and S-D614G mutant structures. (A) The quantitative evaluation of the number of unique hubs in closed and open forms of S-GSAS/D614 (shown in green bars), closed and open forms of S-GSAS/G614 (red bars), and S-RRAR/G614 (blue bars). Structural mapping of common communities shared by closed and open states is projected onto a single protomer for the S-GSAS/D614 in the closed-all down state (B) and S-GSAS/D614 in the open state (C). The hubs are shown as red spheres. (D) Structural mapping of unique in the closed form of S-GSAS/G614 (pdb id 7KDK). (E) Structural mapping of unique hubs for the S-GSAS/G614 in the open state (pdb id 7KDL). The hubs are shown in blue spheres. The mapping is projected onto a single protomer shown in cyan ribbons.

By projecting state-specific hubs for the S-G614 mutant (Figure 10D,E) the differences between the closed and partially open states could be further highlighted, particularly showing a consolidation of new hotspots and a stronger connectivity of these mediating centers near the S1-S2 interface regions for the open form. Among unique hotspots that are specific for the closed mutant state are CTD1 residues N544, F559, R567 and CTD2 residues Y695,Y674 whereas in the open mutant state a number of unique new hubs were detected in the CTD2 region (V597, Q675, Q690, and Q672). These new unique hubs at the S1-S2 interfacial regions in combination with the common hubs in the N2R linker region (T315, F318, R319, and R328) form a subnetwork that stabilizes connections between the NTD and RBD regions and the CTD1 and CTD2 domains. Similar to other network-centric analyses of the SARS-CoV-2 S trimers^115^ we observed some variance in the distribution of hubs in different protomers, confirming that there are elements of the inherent asymmetry in the residue interaction networks across subunits.

To conclude, dynamic network modeling and community analysis of the S-D614 and S-G614 proteins revealed that D614G mutation can induce a partial rearrangement of the residue interaction networks and promote the larger number of stable communities in both the closed and open forms by enhancing the S1-S2 inter-domain interactions. Furthermore, the network analysis suggested a differential stabilization of the S-G614 mutant, favoring the open form in which strengthened allosteric couplings between mutational sites, S1-S2 regions and NTD/RBD regions in S1 could contribute to a decrease in premature shedding of S1 domain. One of the overarching objectives of our computational investigation is to examine functional mechanisms of D614G mutation from the point of view of allosteric regulation and reconcile multiple conflicting scenarios underlying D614G effects. By comparing our results with the growing body of experimental data, we suggested that an allosteric regulatory model may help to explain how the D614G mutation exerts its effect. The proposed allosteric model of SARS-CoV-2 S regulation and functions may be also useful to address questions concerning the effect of mutations on evading antibody binding and immune resistance. Although recent studies indicated that D614G variant did not itself drive escape from antibody binding, it was found that D614G can remarkably potentiate escape mutations at some positions in certain patients, supporting an allosteric mechanism of action triggered by this mutation on dynamics and function in remote regions exposed to interactions with antibodies.^121^ In light of these newly emerging experimental data, the proposed allosteric model of the D614G mechanism may provide a feasible theoretical and modeling framework to examine the role of combinations of mutations and further clarify the reasons underlying the increasing global dominance of the D614G variant. Our results argue that the D614G mutation may exert its global impact on other sites and amplify the effect of combined mutations by acting as an important mediating center governing regulation of the SARS-CoV-2 allosteric machine. Targeting of the predicted mediating centers of allosteric interactions and communication hubs in the SARS-CoV-2 S proteins with small molecules could also help to probe molecular mechanisms underlying infectivity variants while simultaneously facilitating discovery of novel variant-specific allosteric modulators and inhibitors of the SARS-CoV-2 functions.

## Conclusions

We combined several simulation-based approaches with dynamic network modeling and community analysis to quantify the effect of D614G mutation on dynamics, stability and network organization of the SARS-CoV-2 S proteins. The results of this study provide a novel insight into the molecular mechanisms underlying the effect of D614G mutation by examining SARS-CoV-2 S protein as an allosteric regulatory machine. By examining the dynamic and network properties of the SARS-CoV-2 S trimer proteins, we characterized the distribution of allosteric hotspots in the S-D614 and S-G614 mutant structures revealing consolidation of the communication hotspots in the CTD1-CTD2 regions of the S-G614 mutant that may promote the greater stability and efficient allosteric couplings between S1 and S2 regions. Using mutational sensitivity analysis of the SARS-CoV-2 S-D614 and S-G614 proteins we showed that D614G can improve stability of the spike protein, offering support to the reduced shedding mechanism. Through distance fluctuations communication analysis, we also probed stability and distribution of allosteric hotspots providing evidence that D614G mutation can favor the open state to enhance allosteric signaling of the spike engine.

A more detailed network analysis and systematic characterization of stable local communities in multiple states of the S-D614 and S-G614 proteins, we demonstrated that D614G mutation can increase protein stability and strengthen the allosteric interaction networks in both closed and open forms. The results indicated that D614G mutation-induced modulation of stability and allosteric propensities could stabilize the open mutant state by effectively restoring allosteric signaling of the native closed state. This study provides support to the reduced shedding hypothesis suggesting that D614G mutation can exert its primary effect through allosterically induced changes on stability and long-range communications in the residue interaction networks. We argue that considering functions of the SARS-CoV-2 pike proteins through the prism of an allosterically regulated machine may prove to be useful in uncovering functional mechanisms and rationalizing the growing body of diverse experimental data via allosteric models underpinning signaling events. These models can ultimately link the allosteric mechanisms of proteins to their functional role in regulatory processes and signaling events.^122–124^

## SUPPORTING INFORMATION

Supporting information contains Tables S1-S6 that characterizes the inter-protomer contacts in the SARS-CoV-2 S-D614 and SARS-CoV-2 S-G614 structures in the closed and open states. Table S7 details the local interacting communities of the protomer A in the structure of SARS-CoV-2 S-GSAS/D614 in the closed state (pdb id 7KDG). Table S8 describes the local interacting communities of the protomer A in the structure of SARS-CoV-2 S-GSAS/D614 in the open state (pdb id 7KDH). Table S9 details the local interacting communities of the protomer A in the structure of SARS-CoV-2 S-GSAS/G614 in the closed state (pdb id 7KDK). Table S10 lists the local interacting communities of the protomer A in the structure of SARS-CoV-2 S-GSAS/G614 in the open state (pdb id 7KDG). Table S11 details the local interacting communities of the protomer A in the structure of SARS-CoV-2 S-RRAR/G614 in the closed state (pdb id 7KDI). Table S12 unveils the local interacting communities of the protomer A in the structure of SARS-CoV-2 S-RRAR/G614 in the open state (pdb id 7KDJ).

## AUTHOR INFORMATION

* **Corresponding Author** Phone: 714-516-4586; Fax: 714-532-6048;

The authors declare no competing financial interest.

## Acknowledgment

This work was partly supported by institutional funding from Chapman University. The author acknowledges support by the Kay Family Foundation Grant A20-0032.

## ABBREVIATIONS

SARS: Severe Acute Respiratory Syndrome
RBD: Receptor Binding Domain
ACE2: Angiotensin-Converting Enzyme 2 (ACE2)
NTD: N-terminal domain
RBD: receptor-binding domain
CTD1: C-terminal domain 1
CTD2: C-terminal domain 2
FP: fusion peptide
FPPR: fusion peptide proximal region
HR1: heptad repeat 1
CH: central helix region
CD: connector domain
HR2: heptad repeat 2
TM: transmembrane anchor
CT: cytoplasmic tail

## References

1. Li, Q.; Guan, X.; Wu, P.; Wang, X.; Zhou, L.; Tong, Y.; Ren, R.; Leung, K. S. M.; Lau, E. H. Y.; Wong, J. Y.; Xing, X.; Xiang, N.; Wu, Y.; Li, C.; Chen, Q.; Li, D.; Liu, T.; Zhao, J.; Liu, M.; Tu, W.; Chen, C.; Jin, L.; Yang, R.; Wang, Q.; Zhou, S.; Wang, R.; Liu, H.; Luo, Y.; Liu, Y.; Shao, G.; Li, H.; Tao, Z.; Yang, Y.; Deng, Z.; Liu, B.; Ma, Z.; Zhang, Y.; Shi, G.; Lam, T. T. Y.; Wu, J. T.; Gao, G. F.; Cowling, B. J.; Yang, B.; Leung, G. M.; Feng, Z. Early transmission dynamics in Wuhan, China, of novel coronavirus-infected pneumonia. N. Engl. J. Med. 2020, 382, 1199–1207.

2. Wang, C.; Horby, P. W.; Hayden, F. G.; Gao, G. F. A novel coronavirus outbreak of global health concern. Lancet 2020, 395, 470–473.

3. Yi, Y.; Lagniton, P. N. P.; Ye, S.; Li, E.; Xu, R. H. COVID-19: what has been learned and to be learned about the novel coronavirus disease. Int. J. Biol. Sci. 2020, 16, 1753–1766.

4. Wu, A.; Peng, Y.; Huang, B.; Ding, X.; Wang, X.; Niu, P.; Meng, J.; Zhu, Z.; Zhang, Z.; Wang, J.; Sheng, J.; Quan, L.; Xia, Z.; Tan, W.; Cheng, G.; Jiang, T. Genome composition and divergence of the novel coronavirus (2019-nCoV) originating in China. Cell Host Microbe 2020, 27, 325–328.

5. Tai, W.; He, L.; Zhang, X.; Pu, J.; Voronin, D.; Jiang, S.; Zhou, Y.; Du, L. Characterization of the receptor-binding domain (RBD) of 2019 novel coronavirus: implication for development of RBD protein as a viral attachment inhibitor and vaccine. Cell. Mol. Immunol. 2020, 17, 613–620.

6. Hoffmann, M.; Kleine-Weber, H.; Schroeder, S.; Krüger, N.; Herrler, T.; Erichsen, S.; Schiergens, T. S.; Herrler, G.; Wu, N. H.; Nitsche, A.; Müller, M. A.; Drosten, C.; Pöhlmann, S.SARS-CoV-2 cell entry depends on ACE2 and TMPRSS2 and is blocked by a clinically proven protease inhibitor. Cell 2020, 181, 271–280.e8.

7. Lu, R.; Zhao, X.; Li, J.; Niu, P.; Yang, B.; Wu, H.; Wang, W.; Song, H.; Huang, B.; Zhu, N.; Bi, Y.; Ma, X.; Zhan, F.; Wang, L.; Hu, T.; Zhou, H.; Hu, Z.; Zhou, W.; Zhao, L.; Chen, J.; Meng, Y.; Wang, J.; Lin, Y.; Yuan, J.; Xie, Z.; Ma, J.; Liu, W. J.; Wang, D.; Xu, W.; Holmes, E. C.; Gao, G. F.; Wu, G.; Chen, W.; Shi, W.; Tan, W. Genomic characterisation and epidemiology of 2019 novel coronavirus: implications for virus origins and receptor binding. Lancet 2020, *395*, 565–574.

8. Duan, L.; Zheng, Q.; Zhang, H.; Niu, Y.; Lou, Y.; Wang, H. The SARS-CoV-2 spike glycoprotein biosynthesis, structure, function, and antigenicity: Implications for the design of spike-based vaccine immunogens. Front. Immunol. 2020, 11, 576622.

9. Wang, Q.; Zhang, Y.; Wu, L.; Niu, S.; Song, C.; Zhang, Z.; Lu, G.; Qiao, C.; Hu, Y.; Yuen, K. Y.; Zhou, H.; Yan, J.; Qi, J. Structural and functional basis of SARS-CoV-2 entry by using human ACE2. Cell 2020, 181, 894–904.e9.

10. Wan, Y.; Shang, J.; Graham, R.; Baric, R. S.; Li, F. Receptor recognition by the novel coronavirus from Wuhan: An analysis based on decade-long structural studies of SARS coronavirus. J. Virol. 2020, 94, e00127–20.

11. Shang, J.; Wan, Y.; Luo, C.; Ye, G.; Geng, Q.; Auerbach, A.; Li, F. Cell entry mechanisms of SARS-CoV-2. Proc. Natl. Acad. Sci. U. S. A. 2020, 117, 11727–11734.

12. Lu, R.; Zhao, X.; Li, J.; Niu, P.; Yang, B.; Wu, H.; Wang, W.; Song, H.; Huang, B.; Zhu, N.; Bi, Y.; Ma, X.; Zhan, F.; Wang, L.; Hu, T.; Zhou, H.; Hu, Z.; Zhou, W.; Zhao, L.; Chen, J.; Meng, Y.; Wang, J.; Lin, Y.; Yuan, J.; Xie, Z.; Ma, J.; Liu, W. J.; Wang, D.; Xu, W.; Holmes, E. C.; Gao, G. F.; Wu, G.; Chen, W.; Shi, W.; Tan, W. Genomic characterisation and epidemiology of 2019 novel coronavirus: implications for virus origins and receptor binding. Lancet 2020, *395*,565–574.

13. Wang, Q.; Zhang, Y.; Wu, L.; Niu, S.; Song, C.; Zhang, Z.; Lu, G.; Qiao, C.; Hu, Y.; Yuen, K. Y.; Zhou, H.; Yan, J.; Qi, J. Structural and functional basis of SARS-CoV-2 entry by using human ACE2. Cell 2020, 181, 894–904.

14. Wan, Y.; Shang, J.; Graham, R.; Baric, R. S.; Li, F. Receptor recognition by the novel coronavirus from Wuhan: An analysis based on decade-long structural studies of SARS coronavirus. J. Virol. 2020, 94, e00127–20.

15. Shang, J.; Wan, Y.; Luo, C.; Ye, G.; Geng, Q.; Auerbach, A.; Li, F. Cell entry mechanisms of SARS-CoV-2. Proc. Natl. Acad. Sci. U. S. A. 2020, 117, 11727–11734.

16. Walls, A. C.; Park, Y. J.; Tortorici, M. A.; Wall, A.; McGuire, A. T.; Veesler, D. Structure, Function, and Antigenicity of the SARS-CoV-2 Spike Glycoprotein. Cell 2020, 181, 281–292.

17. Wrapp, D.; Wang, N.; Corbett, K. S.; Goldsmith, J. A.; Hsieh, C. L.; Abiona, O.; Graham, B. S.; McLellan, J. S. Cryo-EM structure of the 2019-nCoV spike in the prefusion conformation. Science 2020, 367, 1260–1263.

18. Walls, A. C.; Tortorici, M. A.; Snijder, J.; Xiong, X.; Bosch, B. J.; Rey, F. A.; Veesler, D. Tectonic conformational changes of a coronavirus spike glycoprotein promote membrane fusion. Proc. Natl. Acad. Sci. U. S. A. 2017, 114, 11157–11162.

19. Fan, X.; Cao, D.; Kong, L.; Zhang, X. Cryo-EM analysis of the post-fusion structure of the SARS-CoV spike glycoprotein. Nat. Commun. 2020, 11, 3618.

20. Walls, A. C.; Tortorici, M. A.; Bosch, B. J.; Frenz, B.; Rottier, P. J. M.; DiMaio, F.; Rey, F. A.; Veesler, D. Cryo-electron microscopy structure of a coronavirus spike glycoprotein trimer. Nature 2016, 531, 114–117.

21. Gui, M.; Song, W.; Zhou, H.; Xu, J.; Chen, S.; Xiang, Y.; Wang, X. Cryo-electron microscopy structures of the SARS-CoV spike glycoprotein reveal a prerequisite conformational state for receptor binding. Cell Res. 2017, 27, 119–129.

22. Walls, A. C.; Xiong, X.; Park, Y. J.; Tortorici, M. A.; Snijder, J.; Quispe, J.; Cameroni, E.; Gopal, R.; Dai, M.; Lanzavecchia, A.; Zambon, M.; Rey, F. A.; Corti, D.; Veesler, D. Unexpected receptor functional mimicry elucidates activation of coronavirus fusion. Cell 2019, 176, 1026–1039.e15.

23. Kirchdoerfer, R. N.; Wang, N.; Pallesen, J.; Wrapp, D.; Turner, H. L.; Cottrell, C. A.; Corbett, K. S.; Graham, B. S.; McLellan, J. S.; Ward, A. B. Stabilized coronavirus spikes are resistant to conformational changes induced by receptor recognition or proteolysis. Sci. Rep. 2018, 8, 15701.

24. Walls, A. C.; Park, Y. J.; Tortorici, M. A.; Wall, A.; McGuire, A. T.; Veesler, D. Structure, Function, and Antigenicity of the SARS-CoV-2 Spike Glycoprotein. Cell 2020, 181, 281–292.e6.

25. Wrapp, D.; Wang, N.; Corbett, K. S.; Goldsmith, J. A.; Hsieh, C. L.; Abiona, O.; Graham, B. S.; McLellan, J. S. Cryo-EM structure of the 2019-nCoV spike in the prefusion conformation. Science 2020, 367, 1260–1263.

26. Cai, Y.; Zhang, J.; Xiao, T.; Peng, H.; Sterling, S. M.; Walsh, R. M., Jr.; Rawson, S.; Rits-Volloch, S.; Chen, B. Distinct conformational states of SARS-CoV-2 spike protein. Science 2020, 369, 1586–1592.

27. Lan, J.; Ge, J.; Yu, J.; Shan, S.; Zhou, H.; Fan, S.; Zhang, Q.; Shi, X.; Wang, Q.; Zhang, L.; Wang, X. Structure of the SARS-CoV-2 spike receptor-binding domain bound to the ACE2 receptor. Nature 2020, 581, 215–220.

28. Shang, J.; Ye, G.; Shi, K.; Wan, Y.; Luo, C.; Aihara, H.; Geng, Q.; Auerbach, A.; Li, F. Structural basis of receptor recognition by SARS-CoV-2. Nature 2020, 581, 221–224.

29. Tortorici, M. A.; Veesler, D. Structural insights into coronavirus entry. Adv. Virus Res. 2019, 105, 93–116.

30. Cai, Y.; Zhang, J.; Xiao, T.; Peng, H.; Sterling, S. M.; Walsh, R. M., Jr.; Rawson, S.; Rits-Volloch, S.; Chen, B. Distinct conformational states of SARS-CoV-2 spike protein. Science 2020, 369, 1586–1592.

31. Hsieh, C. L.; Goldsmith, J. A.; Schaub, J. M.; DiVenere, A. M.; Kuo, H. C.; Javanmardi, K.; Le, K. C.; Wrapp, D.; Lee, A. G.; Liu, Y.; Chou, C. W.; Byrne, P. O.; Hjorth, C. K.; Johnson, N. V.; Ludes-Meyers, J.; Nguyen, A. W.; Park, J.; Wang, N.; Amengor, D.; Lavinder, J. J.; Ippolito, G. C.; Maynard, J. A.; Finkelstein, I. J.; McLellan, J. S. Structure-based design of prefusion-stabilized SARS-CoV-2 spikes. Science 2020, 369, 1501–1505.

32. Henderson, R.; Edwards, R. J.; Mansouri, K.; Janowska, K.; Stalls, V.; Gobeil, S. M. C.; Kopp, M.; Li, D.; Parks, R.; Hsu, A. L.; Borgnia, M. J.; Haynes, B. F.; Acharya, P. Controlling the SARS-CoV-2 spike glycoprotein conformation. Nat. Struct. Mol. Biol. 2020, 27, 925–933.

33. McCallum, M.; Walls, A. C.; Bowen, J. E.; Corti, D.; Veesler, D. Structure-guided covalent stabilization of coronavirus spike glycoprotein trimers in the closed conformation. Nat. Struct. Mol. Biol. 2020, 27, 942–949.

34. Xiong, X.; Qu, K.; Ciazynska, K. A.; Hosmillo, M.; Carter, A. P.; Ebrahimi, S.; Ke, Z.; Scheres, S. H. W.; Bergamaschi, L.; Grice, G. L.; Zhang, Y.; Nathan, J. A.; Baker, S.; James, L. C.; Baxendale, H. E.; Goodfellow, I.; Doffinger, R.; Briggs, J. A. G. A thermostable, closed SARS-CoV-2 spike protein trimer. Nat. Struct. Mol. Biol. 2020, 27, 934–941.

35. Turoňová, B.; Sikora, M.; Schürmann, C.; Hagen, W. J. H.; Welsch, S.; Blanc, F. E. C.; von Bülow, S.; Gecht, M.; Bagola, K.; Hörner, C.; van Zandbergen, G.; Landry, J.; de Azevedo, N. T. D.; Mosalaganti, S.; Schwarz, A.; Covino, R.; Mühlebach, M. D.; Hummer, G.; Krijnse Locker, J.; Beck, M. In situ structural analysis of SARS-CoV-2 spike reveals flexibility mediated by three hinges. Science 2020, 370, 203–208.

36. Lu, M.; Uchil, P. D.; Li, W.; Zheng, D.; Terry, D. S.; Gorman, J.; Shi, W.; Zhang, B.; Zhou, T.; Ding, S.; Gasser, R.; Prevost, J.; Beaudoin-Bussieres, G.; Anand, S. P.; Laumaea, A.; Grover, J. R.; Lihong, L.; Ho, D. D.; Mascola, J.; Finzi, A.; Kwong, P. D.; Blanchard, S. C.; Mothes, W. Real-time conformational dynamics of SARS-CoV-2 spikes on virus particles. bioRxiv 2020, doi: 10.1101/2020.09.10.286948.

37. Benton, D. J.; Wrobel, A. G.; Xu, P.; Roustan, C.; Martin, S. R.; Rosenthal, P. B.; Skehel, J. J.; Gamblin, S. J. Receptor binding and priming of the spike protein of SARS-CoV-2 for membrane fusion. Nature 2020, doi: 10.1038/s41586-020-2772-0.

38. Zhou, T.; Tsybovsky, Y.; Gorman, J.; Rapp, M.; Cerutti, G.; Chuang, G. Y.; Katsamba, P. S.; Sampson, J. M.; Schön, A.; Bimela, J.; Boyington, J. C.; Nazzari, A.; Olia, A. S.; Shi, W.; Sastry, M.; Stephens, T.; Stuckey, J.; Teng, I. T.; Wang, P.; Wang, S.; Zhang, B.; Friesner, R. A.; Ho, D. D.; Mascola, J. R.; Shapiro, L.; Kwong, P. D., Cryo-EM Structures of SARS-CoV-2 Spike without and with ACE2 Reveal a pH-Dependent Switch to Mediate Endosomal Positioning of Receptor-Binding Domains. Cell Host Microbe 2020, 28 (6), 867–879.e5.

39. Korber, B.; Fischer, W. M.; Gnanakaran, S.; Yoon, H.; Theiler, J.; Abfalterer, W.; Hengartner, N.; Giorgi, E. E.; Bhattacharya, T.; Foley, B.; Hastie, K. M.; Parker, M. D.; Partridge, D. G.; Evans, C. M.; Freeman, T. M.; de Silva, T. I.; McDanal, C.; Perez, L. G.; Tang, H.; Moon-Walker, A.; Whelan, S. P.; LaBranche, C. C.; Saphire, E. O.; Montefiori, D. C. Tracking changes in SARS-CoV-2 Spike: Evidence that D614G increases infectivity of the COVID-19 virus. Cell 2020, 182, 812–827.e19.

40. Plante, J. A.; Liu, Y.; Liu, J.; Xia, H.; Johnson, B. A.; Lokugamage, K. G.; Zhang, X.; Muruato, A. E.; Zou, J.; Fontes-Garfias, C. R.; Mirchandani, D.; Scharton, D.; Bilello, J. P.; Ku, Z.; An, Z.; Kalveram, B.; Freiberg, A. N.; Menachery, V. D.; Xie, X.; Plante, K. S.; Weaver, S. C.; Shi, P. Y. Spike mutation D614G alters SARS-CoV-2 fitness. Nature 2020, doi:10.1038/s41586-020-2895-3.

41. Hou, Y. J.; Chiba, S.; Halfmann, P.; Ehre, C.; Kuroda, M.; Dinnon, K. H., 3rd; Leist, S. R.; Schäfer, A.; Nakajima, N.; Takahashi, K.; Lee, R. E.; Mascenik, T. M.; Graham, R.; Edwards, C. E.; Tse, L. V.; Okuda, K.; Markmann, A. J.; Bartelt, L.; de Silva, A.; Margolis, D. M.; Boucher, R. C.; Randell, S. H.; Suzuki, T.; Gralinski, L. E.; Kawaoka, Y.; Baric, R. S. SARS-CoV-2 D614G variant exhibits efficient replication ex vivo and transmission in vivo. Science 2020, doi: 10.1126/science.abe8499.

42. Jackson, C. B.; Zhang, L.; Farzan, M.; Choe, H., Functional importance of the D614G mutation in the SARS-CoV-2 spike protein. Biochem Biophys Res Commun 2020. doi:10.1016/j.bbrc.2020.11.026.

43. Yurkovetskiy, L.; Wang, X.; Pascal, K. E.; Tomkins-Tinch, C.; Nyalile, T. P.; Wang, Y.; Baum, A.; Diehl, W. E.; Dauphin, A.; Carbone, C.; Veinotte, K.; Egri, S. B.; Schaffner, S. F.; Lemieux, J. E.; Munro, J. B.; Rafique, A.; Barve, A.; Sabeti, P. C.; Kyratsous, C. A.; Dudkina, N. V.; Shen, K.; Luban, J. Structural and functional analysis of the D614G SARS-CoV-2 spike protein variant. Cell 2020, 183, 739–751.e8.

44. Gobeil, S. M.; Janowska, K.; McDowell, S.; Mansouri, K.; Parks, R.; Manne, K.; Stalls, V.; Kopp, M. F.; Henderson, R.; Edwards, R. J.; Haynes, B. F.; Acharya, P. D614G Mutation Alters SARS-CoV-2 Spike Conformation and Enhances Protease Cleavage at the S1/S2 Junction. Cell Rep 2021, 34, 108630.

45. Weissman, D.; Alameh, M. G.; de Silva, T.; Collini, P.; Hornsby, H.; Brown, R.; LaBranche, C. C.; Edwards, R. J.; Sutherland, L.; Santra, S.; Mansouri, K.; Gobeil, S.; McDanal, C.; Pardi, N.; Hengartner, N.; Lin, P. J. C.; Tam, Y.; Shaw, P. A.; Lewis, M. G.; Boesler, C.; Şahin, U.; Acharya, P.; Haynes, B. F.; Korber, B.; Montefiori, D. C. D614G Spike Mutation Increases SARS CoV-2 Susceptibility to Neutralization. Cell Host Microbe 2021, 29,23–31.

46. Zhang, L.; Jackson, C. B.; Mou, H.; Ojha, A.; Peng, H.; Quinlan, B. D.; Rangarajan, E. S.; Pan, A.; Vanderheiden, A.; Suthar, M. S.; Li, W.; Izard, T.; Rader, C.; Farzan, M.; Choe, H., SARS-CoV-2 spike-protein D614G mutation increases virion spike density and infectivity. Nat Commun 2020, 11, 6013.

47. Korber, B.; Fischer, W. M.; Gnanakaran, S.; Yoon, H.; Theiler, J.; Abfalterer, W.; Foley, B.; Giorgi, E. E.; Bhattacharya, T.; Parker, M. D.; Partridge, D. G.; Evans, C. M.; de Silva, T. I.; LaBranche, C. C.; Montefiori, D. Spike mutation pipeline reveals the emergence of a more transmissible form of SARS-CoV-2. bioRxiv 2020, doi: 10.1101/2020.04.29.069054.

48. Zhang, J.; Cai, Y.; Xiao, T.; Lu, J.; Peng, H.; Sterling, S. M.; Walsh, R. M.; Rits-Volloch, S.; Sliz, P.; Chen, B., Structural impact on SARS-CoV-2 spike protein by D614G substitution. bioRxiv 2020. doi: 10.1101/2020.10.13.337980.

49. Juraszek, J.; Rutten, L.; Blokland, S.; Bouchier, P.; Voorzaat, R.; Ritschel, T.; Bakkers, M. J. G.; Renault, L. L. R.; Langedijk, J. P. M. Stabilizing the closed SARS-CoV-2 spike trimer. Nat Commun 2021, 12, 244.

50. Mansbach, R. A.; Chakraborty, S.; Nguyen, K.; Montefiori, D.; Korber, B.; Gnanakaran, S. The SARS-CoV-2 spike variant D614G favors an open conformational state. bioRxiv 2020, doi: 10.1101/2020.07.26.219741.2007.2026.219741.

51. Xu, C.; Wang, Y.; Liu, C.; Zhang, C.; Han,W.; Hong, X.; Wang, Y.; Hong, Q.; Wang, S.; Zhao, Q.; Wang, Y.; Yang, Y.; Chen, K.; Zheng, W.; Kong, L.; Wang, F.; Zuo, Q.; Huang, Z., Cong, Y. Conformational dynamics of SARS-CoV-2 trimeric spike glycoprotein in complex with receptor ACE2 revealed by cryo-EM. bioRxiv 2020, doi: 10.1101/2020.06.30.177097.

52. Teruel, N.; Mailhot, O.; Najmanovich, R.J. Modeling conformational state dynamics and its role on infection for SARS-CoV-2 Spike protein variants. bioRxiv 2020, doi: https://doi.org/10.1101/2020.12.16.423118

53. Yazhini, A.; Prakash Sidhanta, D.S.; Srinivasan, N. D614G substitution enhances the stability of trimeric SARS-CoV-2 spike protein. bioRxiv 2020, doi: https://doi.org/10.1101/2020.11.02.364273.

54. Ray, D.; Le, L.; Andricioaei, I. Distant Residues Modulate Conformational Opening in SARS-CoV-2 Spike Protein. bioRxiv 2020, doi: https://doi.org/10.1101/2020.12.07.415596

55. Fernández, A., Structural Impact of Mutation D614G in SARS-CoV-2 Spike Protein: Enhanced Infectivity and Therapeutic Opportunity. ACS Med Chem Lett 2020, 11, 1667–1670.

56. Woo, H.; Park, S. J.; Choi, Y. K.; Park, T.; Tanveer, M.; Cao, Y.; Kern, N. R.; Lee, J.; Yeom, M. S.; Croll, T. I.; Seok, C.; Im, W. Developing a fully glycosylated full-length SARS-CoV-2 spike protein model in a viral membrane. J. Phys. Chem. B 2020, 124, 7128–7137.

57. Casalino, L.; Gaieb, Z.; Goldsmith, J.A.; Hjorth, C.K.; Dommer, A. C.; Harbison, A. M.; Fogarty, C. A.; Barros, E. P.; Taylor, B. C.; McLellan, J.S.; Fadda, E.; Amaro, R. E. Beyond shielding: The roles of glycans in the SARS-CoV-2 spike potein. ACS Cent. Sci. 2020, 6, 1722–1734.

58. Yu, A.; Pak, A. J.; He, P.; Monje-Galvan, V.; Casalino, L.; Gaieb, Z.; Dommer, A. C.; Amaro, R. E.; Voth, G. A. A multiscale coarse-grained model of the SARS-CoV-2 virion. Biophys J 2020, doi: 10.1016/j.bpj.2020.10.048.

59. Ghorbani, M.; Brooks, B. R.; Klauda, J. B. Exploring dynamics and network analysis of spike glycoprotein of SARS-COV-2. bioRxiv 2020, doi: 10.1101/2020.09.28.317206.

60. Wang, Y.; Liu, M.; Gao, J. Enhanced receptor binding of SARS-CoV-2 through networks of hydrogen-bonding and hydrophobic interactions. Proc. Natl. Acad. Sci. U. S. A. 2020, 117,13967–13974.

61. Ali, A.; Vijayan, R. Dynamics of the ACE2-SARS-CoV-2/SARS-CoV spike protein interface reveal unique mechanisms. Sci Rep 2020, 10, 14214.

62. Verkhivker, G.M. Coevolution, dynamics and allostery conspire in shaping cooperative binding and signal transmission of the SARS-CoV-2 spike protein with human angiotensin-converting enzyme 2. Int. J. Mol. Sci. 2020, 21, 8268.

63. Di Paola, L.; Hadi-Alijanvand, H.; Song, X.; Hu, G.; Giuliani, A. The discovery of a putative allosteric site in the SARS-CoV-2 spike protein using an integrated structural/dynamic approach. J. Proteome Res. 2020, 19, 4576–4586.

64. Verkhivker, G.M. Molecular simulations and network modeling reveal an allosteric signaling in the SARS-CoV-2 spike proteins. J. Proteome Res. 2020, 19, 4587–4608.

65. Kolinski, A. Protein modeling and structure prediction with a reduced representation. Acta Biochim. Pol. 2004, 51, 349–371.

66. Kmiecik, S.; Gront, D.; Kolinski, M.; Wieteska, L.; Dawid, A.E.; Kolinski, A. Coarse-grained protein models and their applications. Chem. Rev. 2016, 116, 7898–7936.

67. Kmiecik, S.; Kouza, M.; Badaczewska-Dawid, A.E.; Kloczkowski, A.; Kolinski, A. Modeling of protein structural flexibility and large-scale dynamics: Coarse-grained simulations and elastic network models. Int. J. Mol. Sci. 2018, 19, e3496.

68. Ciemny, M.P.; Badaczewska-Dawid, A.E.; Pikuzinska, M.; Kolinski, A.; Kmiecik, S. Modeling of disordered protein structures using monte carlo simulations and knowledge-based statistical force fields. Int. J. Mol. Sci. 2019, 20, e606.

69. Kurcinski, M.; Oleniecki, T.; Ciemny, M.P.; Kuriata, A.; Kolinski, A.; Kmiecik, S. CABS-flex standalone: A simulation environment for fast modeling of protein flexibility. Bioinformatics 2019, 35, 694–695.

70. Jamroz, M.; Orozco, M.; Kolinski, A.; Kmiecik, S. Consistent view of protein fluctuations from all-atom molecular dynamics and coarse-grained dynamics with knowledge-based force-field. J. Chem. Theory Comput. 2013, 9, 119–125.

71. Badaczewska-Dawid, A. E.; Kolinski, A.; Kmiecik, S. Protocols for fast simulations of protein structure flexibility using CABS-Flex and SURPASS. Methods Mol. Biol. 2020, 2165, 337–353.

72. Berman, H.M.; Westbrook, J.; Feng, Z.; Gilliland, G.; Bhat, T.N.; Weissig, H.; Shindyalov, I.N.; Bourne, P.E. The protein data nank. Nucleic Acids Res. 2000, 28, 235–242.

73. Rose, P. W.; Prlic, A.; Altunkaya, A.; Bi, C.; Bradley, A. R.; Christie, C. H.; Costanzo, L. D.; Duarte, J. M.; Dutta, S.; Feng, Z.; Green, R. K.; Goodsell, D. S.; Hudson, B.; Kalro, T.; Lowe, R.; Peisach, E.; Randle, C.; Rose, A. S.; Shao, C.; Tao, Y. P.; Valasatava, Y.; Voigt, M.; Westbrook, J. D.; Woo, J.; Yang, H.; Young, J. Y.; Zardecki, C.; Berman, H. M.; Burley, S. K. The RCSB protein data bank: integrative view of protein, gene and 3D structural information. Nucleic Acids Res. 2017, 45, D271–D281.

74. Hooft, R. W.; Sander, C.; Vriend, G. Positioning hydrogen atoms by optimizing hydrogen-bond networks in protein structures. Proteins 1996, 26, 363–376.

75. Hekkelman, M. L.; Te Beek, T. A.; Pettifer, S. R.; Thorne, D.; Attwood, T. K.; Vriend, G. WIWS: A protein structure bioinformatics web service collection. Nucleic Acids Res. 2010, 38, W719–W723.

76. Fiser, A.; Sali, A. ModLoop: Automated modeling of loops in protein structures. Bioinformatics 2003, 19, 2500–2501.

77. Fernandez-Fuentes, N.; Zhai, J.; Fiser, A. ArchPRED: A template based loop structure prediction server. Nucleic Acids Res. 2006, 34, W173–W176.

78. Ko, J.; Lee, D.; Park, H.; Coutsias, E. A.; Lee, J.; Seok, C. The FALC-Loop web server for protein loop modeling. Nucleic Acids Res. 2011, 39, W210–W214.

79. Krivov, G. G.; Shapovalov, M. V.; Dunbrack, R. L., Jr. Improved prediction of protein side-chain conformations with SCWRL4. Proteins 2009, 77, 778–795.

80. Rotkiewicz, P.; Skolnick, J. Fast procedure for reconstruction of full-atom protein models from reduced representations. J. Comput. Chem. 2008, 29, 1460–1465.

81. Lombardi, L. E.; Marti, M. A.; Capece, L. CG2AA: backmapping protein coarse-grained structures. Bioinformatics 2016, 32, 1235–1237.

82. Bhattacharya, D.; Nowotny, J.; Cao, R.; Cheng, J. 3Drefine: an interactive web server for efficient protein structure refinement. Nucleic Acids Res. 2016, 44, W406–W409.

83. Bahar, I.; Lezon, T. R.; Yang, L. W.; Eyal, E. Global dynamics of proteins: bridging between structure and function. Annu. Rev. Biophys. 2010, 39, 23–42.

84. Yang, L. W.; Rader, A. J.; Liu, X.; Jursa, C. J.; Chen, S. C.; Karimi, H. A.; Bahar, I. oGNM: online computation of structural dynamics using the gaussian network model. Nucleic Acids Res. 2006, 34, W24–W31.

85. Li, H.; Chang, Y. Y.; Lee, J. Y.; Bahar, I.; Yang, L. W. DynOmics: dynamics of structural proteome and beyond. Nucleic Acids Res. 2017, 45, W374–W380.

86. Eyal, E.; Lum, G.; Bahar, I. The anisotropic network model web server at 2015 (ANM 2.0). Bioinformatics 2015, 31, 1487–1489.

87. Guerois, R.; Nielsen, J.E.; Serrano, L. Predicting Changes in the Stability of Proteins and Protein Complexes: A Study of More than 1000 Mutations. J. Mol. Biol. 2002, 320, 369–387.

88. Tokuriki, N.; Stricher, F.; Schymkowitz, J.; Serrano, L.; Tawfik, D.S. The Stability Effects of Protein Mutations Appear to be Universally Distributed. J. Mol. Biol. 2007, 369, 1318–1332.

89. Schymkowitz, J.; Borg, J.; Stricher, F.; Nys, R.; Rousseau, F.; Serrano L. The FoldX Web Server: An Online Force Field. Nucleic Acids Res. 2005, 33, W382–W388.

90. Van Durme, J.; Delgado, J.; Stricher, F.; Serrano, L.; Schymkowitz, J.; Rousseau, F. A Graphical Interface for the FoldX Force Field. Bioinformatics 2011, 27, 1711–1712.

91. Dehouck, Y.; Kwasigroch, J. M.; Rooman, M.; Gilis, D. BeAtMuSiC: Prediction of changes in protein-protein binding affinity on mutations. Nucleic Acids Res. 2013, 41, W333–W339.

92. Dehouck, Y.; Gilis, D.; Rooman, M. A new generation of statistical potentials for proteins. Biophys. J. 2006, 90, 4010–4017.

93. Dehouck, Y.; Grosfils, A.; Folch, B.; Gilis, D.; Bogaerts, P.; Rooman, M. Fast and accurate predictions of protein stability changes upon mutations using statistical potentials and neural networks: PoPMuSiC-2.0. Bioinformatics 2009, 25, 2537–2543.

94. Navizet, I.; Cailliez, F.; Lavery, R., Probing protein mechanics: residue-level properties and their use in defining domains. Biophys. J. 2004, 87, 1426–1435.

95. Sacquin-Mora, S.; Lavery, R., Investigating the local flexibility of functional residues in hemoproteins. Biophys. J. 2006, 90, 2706–2717.

96. Sacquin-Mora, S.; Laforet, E.; Lavery, R., Locating the active sites of enzymes using mechanical properties. Proteins 2007, 67, 350–359.

97. Lavery, R.; Sacquin-Mora, S., Protein mechanics: a route from structure to function. J. Biosci. 2007, 32, 891–898.

98. Blacklock, K.; Verkhivker, G. M., Computational modeling of allosteric regulation in the hsp90 chaperones: a statistical ensemble analysis of protein structure networks and allosteric communications. PLoS Comput Biol 2014, 10, e1003679.

99. Stetz, G.; Verkhivker, G. M. Computational analysis of residue interaction networks and coevolutionary relationships in the Hsp70 chaperones: A community-hopping model of allosteric regulation and communication. Plos Comput. Biol. 2017, 13, e1005299.

100. Rader, A. J.; Brown, S. M., Correlating allostery with rigidity. Mol Biosyst 2011, 7, 464–471.

101. Rader, A.J.; Yennamalli, R.M.; Harter, A.K.; Sen, T.Z. A rigid network of long-range contacts increases thermostability in a mutant endoglucanase. J. Biomol. Struct. Dyn. 2012, 30, 628–637.

102. Vijayabaskar, M.S.; Vishveshwara, S. Interaction energy based protein structure networks. Biophys. J. 2010, 99, 3704–3715.

103. Sethi, A.; Eargle, J.; Black, A.A.; Luthey-Schulten, Z. Dynamical networks in tRNA:protein complexes. Proc. Natl. Acad. Sci. U.S.A. 2009, 106, 6620–6625.

104. Czemeres, J.; Buse, K.; Verkhivker, G. M. Atomistic simulations and network-based modeling of the Hsp90-Cdc37 chaperone binding with Cdk4 client protein: A mechanism of chaperoning kinase clients by exploiting weak spots of intrinsically dynamic kinase domains. PLoS One 2017, 12, e0190267.

105. Stetz, G.; Verkhivker, G. M. Dancing through life: Molecular dynamics simulations and network-centric modeling of allosteric mechanisms in Hsp70 and Hsp110 chaperone proteins. PLoS One 2015, 10, e0143752.61.

106. Floyd, R.W. Algorithm 97: Shortest path. Commun. A.C.M. 1962, 5, 345.

107. Hagberg, A.A.; Schult, D.A.; Swart, P.J. Exploring network structure, dynamics, and function using NetworkX, in : G. Varoquaux, T. Vaught, J. Millman (Eds.), Proceedings of the 7th Python in Science Conference (SciPy2008), Pasadena, 2008, pp. 11–15.

108. Girvan, M.; Newman, M. E. Community Structure in Social and Biological Networks. Proc. Natl. Acad. Sci. U. S. A. 2002, 99, 7821–7826.

109. Newman, M. E.; Girvan, M. Finding and Evaluating Community Structure in Networks. Phys. Rev. E Stat. Nonlin. Soft Matter Phys. 2004, 69, 026113.

110. Newman, M. E. Modularity and Community Structure in Networks. Proc. Natl. Acad. Sci. U. S. A. 2006, 103, 8577–8582.

111. Astl, L.; Verkhivker, G. M. Atomistic modeling of the ABL kinase regulation by allosteric modulators using structural perturbation analysis and community-based network reconstruction of allosteric communications. J. Chem. Theory Comput. 2019, 15, 3362–3380.

112. Astl, L.; Verkhivker, G.M. Dynamic view of allosteric regulation in the Hsp70 chaperones by J-Domain cochaperone and post-translational modifications: Computational analysis of Hsp70 mechanisms by exploring conformational landscapes and residue interaction networks. J. Chem. Inf. Model. 2020, 60, 1614–1631.

113. Shannon, P.; Markiel, A.; Ozier, O.; Baliga, N. S.; Wang, J. T.; Ramage, D.; Amin, N.; Schwikowski, B.; Ideker, T. Cytoscape: A software environment for integrated models of biomolecular interaction networks. Genome Res 2003, 13, 2498–2504.

114. Kovács, I.A.; Palotai, R.; Szalay, M.S.; Csermely, P. Community landscapes: an integrative approach to determine overlapping network module hierarchy, identify key nodes and predict network dynamics. PLoS One 2010, 5, e12528.

115. Halder, A.; Anto, A.; Subramanyan, V.; Bhattacharyya, M.; Vishveshwara, S., Surveying the Side-Chain Network Approach to Protein Structure and Dynamics: The SARS-CoV-2 Spike Protein as an Illustrative Case. Front Mol Biosci 2020, 7, 596945.

116. Vangone, A.; Bonvin, A. M., Contacts-based prediction of binding affinity in protein-protein complexes. Elife 2015, 4, e07454.

117. Xue, L. C.; Rodrigues, J. P.; Kastritis, P. L.; Bonvin, A. M.; Vangone, A., PRODIGY: a web server for predicting the binding affinity of protein-protein complexes. Bioinformatics 2016, 32, 3676–3678.

118. Verkhivker, G. M.; Di Paola, L., Dynamic Network Modeling of Allosteric Interactions and Communication Pathways in the SARS-CoV-2 Spike Trimer Mutants: Differential Modulation of Conformational Landscapes and Signal Transmission via Cascades of Regulatory Switches. J Phys Chem B 2021. doi: 10.1021/acs.jpcb.0c10637.

119. Li, Q.; Wu, J.; Nie, J.; Zhang, L.; Hao, H.; Liu, S.; Zhao, C.; Zhang, Q.; Liu, H.; Nie, L.; Qin, H.; Wang, M.; Lu, Q.; Li, X.; Sun, Q.; Liu, J.; Huang, W.; Wang, Y. The impact of mutations in SARS-CoV-2 spike on viral infectivity and antigenicity. Cell 2020, 182, 1284–1294.e9.

120. Laha, S.; Chakraborty, J.; Das, S.; Manna, S. K.; Biswas, S.; Chatterjee, R. Characterizations of SARS-CoV-2 mutational profile, spike protein stability and viral transmission. Infect. Genet. Evol. 2020, 85, 104445.

121. Garrett, M. E.; Galloway, J.; Chu, H. Y.; Itell, H. L.; Stoddard, C. I.; Wolf, C. R.; Logue, J. K.; McDonald, D.; Matsen, F. A.; Overbaugh, J., High resolution profiling of pathways of escape for SARS-CoV-2 spike-binding antibodies. bioRxiv 2020, doi: 10.1101/2020.11.16.385278.

122. Dokholyan, N. V. Controlling allosteric networks in proteins. Chem. Rev. 2016, 116, 6463–6487.

123. Tsai, C. J.; Nussinov, R. A unified view of “how allostery works”. PLoS Comput. Biol. 2014, 10, e1003394

124. Wodak, S. J.; Paci, E.; Dokholyan, N. V.; Berezovsky, I. N.; Horovitz, A.; Li, J.; Hilser, V. J.; Bahar, I.; Karanicolas, J.; Stock, G.; Hamm, P.; Stote, R. H.; Eberhardt, J.; Chebaro, Y.; Dejaegere, A.; Cecchini, M.; Changeux, J. P.; Bolhuis, P. G.; Vreede, J.; Faccioli, P.; Orioli, S.; Ravasio, R.; Yan, L.; Brito, C.; Wyart, M.; Gkeka, P.; Rivalta, I.; Palermo, G.; McCammon, J. A.; Panecka-Hofman, J.; Wade, R. C.; Di Pizio, A.; Niv, M. Y.; Nussinov, R.; Tsai, C. J.; Jang, H.; Padhorny, D.; Kozakov, D.; McLeish, T. Allostery in its many disguises: From theory to applications. Structure 2019, 27, 566–578.

125. Wodak, S. J.; Paci, E.; Dokholyan, N. V.; Berezovsky, I. N.; Horovitz, A.; Li, J.; Hilser, V. J.; Bahar, I.; Karanicolas, J.; Stock, G., et al. Allostery in its many disguises: From theory to applications. Structure 2019, 27, 566–578.

