## Supplemental Tables S1-S12 for "Computational Analysis of Protein Stability and Allosteric Interaction Networks in Distinct Conformational Forms of the SARS-CoV-2 Spike D614G Mutant: Reconciling Functional Mechanisms through Allosteric Model of Spike Regulation"

**Table S1.** The list of the inter-protomer contacts of the protomer A in the structure of SARS-CoV-2 S-GSAS/D614 in the closed state (pdb id 7KDG).

| **S protein residue name** | **S protein residue number** | **S protein chain** | **S protein residue name** | **S protein residue number** | **S protein chain** |
| --- | --- | --- | --- | --- | --- |
| PHE | 970 | A | GLN | 755 | B |
| ALA | 890 | A | VAL | 1068 | C |
| ASN | 919 | A | VAL | 1128 | C |
| MET | 900 | A | VAL | 1094 | C |
| MET | 740 | A | ARG | 319 | C |
| VAL | 705 | A | LEU | 894 | B |
| ILE | 569 | A | ASN | 960 | B |
| ASN | 907 | A | ARG | 1091 | C |
| GLN | 913 | A | GLY | 1093 | C |
| ARG | 466 | A | GLY | 232 | B |
| TYR | 369 | A | GLN | 414 | C |
| ALA | 1016 | A | ILE | 1013 | C |
| GLN | 779 | A | MET | 697 | C |
| SER | 758 | A | GLN | 965 | C |
| LEU | 864 | A | MET | 697 | C |
| ILE | 712 | A | PRO | 897 | B |
| PRO | 463 | A | GLY | 199 | B |
| ALA | 668 | A | PRO | 862 | B |
| GLY | 891 | A | LYS | 1045 | C |
| TYR | 707 | A | PHE | 898 | B |
| ALA | 890 | A | ASP | 1041 | C |
| LEU | 966 | A | ASP | 571 | C |
| ASN | 856 | A | THR | 572 | C |
| GLN | 613 | A | THR | 859 | B |
| SER | 758 | A | SER | 968 | C |
| ASN | 709 | A | PRO | 897 | B |
| SER | 45 | A | LYS | 557 | C |
| LEU | 546 | A | ASP | 979 | B |
| ILE | 670 | A | LEU | 864 | B |
| TYR | 505 | A | SER | 373 | B |
| ASN | 969 | A | GLN | 755 | B |
| VAL | 382 | A | LEU | 984 | B |
| THR | 1009 | A | ILE | 1013 | C |
| PRO | 897 | A | SER | 708 | C |
| PHE | 970 | A | GLY | 757 | B |
| LEU | 560 | A | GLY | 283 | B |
| TYR | 873 | A | LEU | 699 | C |
| GLY | 545 | A | ASN | 978 | B |
| ALA | 766 | A | GLN | 1010 | C |
| PHE | 855 | A | PHE | 592 | C |
| TYR | 1047 | A | ALA | 890 | B |
| PRO | 863 | A | GLY | 669 | C |
| GLU | 918 | A | GLY | 1124 | C |
| PHE | 855 | A | PRO | 589 | C |
| GLU | 988 | A | PRO | 384 | C |
| GLY | 889 | A | GLY | 1046 | C |
| ALA | 701 | A | GLN | 787 | B |
| TYR | 873 | A | GLY | 700 | C |
| VAL | 1068 | A | GLY | 891 | B |
| LYS | 386 | A | ARG | 983 | B |
| GLN | 1002 | A | TYR | 756 | B |
| GLN | 762 | A | GLN | 965 | C |
| LEU | 981 | A | LYS | 386 | C |
| LEU | 865 | A | MET | 697 | C |
| GLN | 1036 | A | VAL | 1040 | C |
| ALA | 892 | A | PRO | 1069 | C |
| GLY | 798 | A | ILE | 1130 | C |
| GLN | 314 | A | LEU | 861 | B |
| TYR | 873 | A | SER | 698 | C |
| PHE | 1089 | A | ASN | 914 | B |
| GLU | 773 | A | GLU | 1017 | C |
| GLN | 52 | A | LEU | 754 | B |
| ASP | 427 | A | VAL | 987 | C |
| ARG | 1019 | A | ALA | 1016 | C |
| LEU | 1145 | A | LEU | 1145 | B |
| TYR | 1047 | A | THR | 887 | B |
| ASP | 1041 | A | LEU | 1034 | B |
| GLY | 889 | A | ASP | 1041 | C |
| GLY | 999 | A | PHE | 759 | B |
| LEU | 390 | A | SER | 982 | B |
| PRO | 1079 | A | TYR | 917 | B |
| GLN | 872 | A | LEU | 699 | C |
| ALA | 372 | A | ASP | 405 | C |
| GLN | 913 | A | PRO | 1079 | C |
| GLU | 918 | A | VAL | 1128 | C |
| PHE | 43 | A | PHE | 559 | C |
| ASN | 703 | A | ILE | 788 | B |
| ASP | 1041 | A | GLY | 889 | B |
| SER | 884 | A | VAL | 705 | C |
| ASP | 571 | A | LEU | 966 | B |
| GLU | 465 | A | GLY | 232 | B |
| ILE | 1013 | A | ILE | 1013 | C |
| LEU | 864 | A | ILE | 670 | C |
| ALA | 668 | A | THR | 866 | B |
| PHE | 565 | A | LYS | 41 | B |
| GLY | 669 | A | PRO | 863 | B |
| SER | 735 | A | GLN | 314 | C |
| GLY | 757 | A | PHE | 970 | C |
| GLN | 895 | A | SER | 708 | C |
| VAL | 42 | A | GLY | 566 | C |
| LYS | 557 | A | PHE | 43 | B |
| GLY | 891 | A | GLY | 1046 | C |
| TYR | 873 | A | MET | 697 | C |
| ARG | 567 | A | PHE | 43 | B |
| GLN | 563 | A | PHE | 43 | B |
| PHE | 592 | A | LEU | 858 | B |
| ASP | 40 | A | PHE | 562 | C |
| THR | 167 | A | ARG | 466 | C |
| LEU | 518 | A | LYS | 41 | B |
| ARG | 567 | A | ASP | 979 | B |
| SER | 1123 | A | ASN | 914 | B |
| TYR | 904 | A | GLY | 1093 | C |
| ALA | 890 | A | TYR | 1067 | C |
| VAL | 1128 | A | GLU | 918 | B |
| ARG | 466 | A | THR | 167 | B |
| PHE | 970 | A | ASP | 994 | B |
| ILE | 1013 | A | ARG | 1019 | B |
| LEU | 894 | A | ARG | 1107 | C |
| ARG | 567 | A | VAL | 976 | B |
| LEU | 861 | A | GLN | 314 | C |
| PRO | 230 | A | TYR | 396 | C |
| TYR | 707 | A | ILE | 794 | B |
| TYR | 707 | A | ASP | 796 | B |
| GLU | 1072 | A | ALA | 892 | B |
| GLN | 895 | A | ILE | 712 | C |
| PRO | 897 | A | ASN | 709 | C |
| ASN | 1074 | A | GLN | 895 | B |
| ARG | 403 | A | ALA | 372 | B |
| MET | 900 | A | THR | 1077 | C |
| GLN | 564 | A | LYS | 41 | B |
| MET | 740 | A | PRO | 589 | C |
| GLY | 669 | A | THR | 866 | B |
| THR | 1009 | A | THR | 1009 | B |
| ASP | 405 | A | ALA | 372 | B |
| GLN | 957 | A | ARG | 765 | B |
| SER | 968 | A | GLY | 757 | B |
| TRP | 886 | A | TYR | 1047 | C |
| ILE | 666 | A | LEU | 864 | B |
| GLY | 971 | A | ASP | 994 | B |
| SER | 982 | A | GLY | 545 | C |
| TYR | 38 | A | GLN | 563 | C |
| ALA | 570 | A | SER | 975 | B |
| TYR | 38 | A | LEU | 560 | C |
| GLN | 755 | A | SER | 968 | C |
| PRO | 1069 | A | LEU | 894 | B |
| GLY | 283 | A | LEU | 560 | C |
| PHE | 562 | A | VAL | 42 | B |
| ILE | 468 | A | GLN | 115 | B |
| GLN | 563 | A | GLY | 283 | B |
| PHE | 559 | A | PHE | 43 | B |
| SER | 698 | A | TYR | 873 | B |
| TYR | 707 | A | ILE | 896 | B |
| ILE | 896 | A | ILE | 712 | C |
| LEU | 560 | A | PHE | 43 | B |
| ARG | 983 | A | VAL | 382 | C |
| ARG | 983 | A | LYS | 386 | C |
| PHE | 759 | A | SER | 1003 | C |
| THR | 912 | A | PHE | 1121 | C |
| THR | 883 | A | VAL | 705 | C |
| LYS | 41 | A | HIS | 519 | C |
| LYS | 202 | A | LEU | 518 | C |
| ILE | 233 | A | PHE | 464 | C |
| GLN | 784 | A | ASP | 1041 | C |
| VAL | 42 | A | PHE | 562 | C |
| LYS | 786 | A | GLY | 700 | C |
| LEU | 984 | A | TYR | 380 | C |
| GLU | 1017 | A | GLU | 773 | B |
| GLN | 1010 | A | LEU | 1012 | B |
| LYS | 986 | A | ASP | 427 | B |
| GLY | 1046 | A | GLY | 891 | B |
| LYS | 964 | A | SER | 758 | B |
| MET | 869 | A | GLY | 669 | C |
| ARG | 1091 | A | ASN | 907 | B |
| ILE | 569 | A | VAL | 963 | B |
| ILE | 896 | A | TYR | 707 | C |
| ASN | 907 | A | ARG | 1107 | C |
| SER | 968 | A | SER | 758 | B |
| MET | 697 | A | LEU | 864 | B |
| VAL | 987 | A | PRO | 412 | B |
| PHE | 562 | A | LYS | 41 | B |
| TYR | 396 | A | PRO | 230 | B |
| LEU | 865 | A | LEU | 699 | C |
| SER | 968 | A | ASP | 571 | C |
| VAL | 1008 | A | GLN | 1010 | C |
| GLU | 918 | A | VAL | 1129 | C |
| ARG | 1019 | A | ILE | 1013 | C |
| ARG | 983 | A | GLY | 381 | C |
| GLN | 913 | A | PHE | 1121 | C |
| TYR | 380 | A | LEU | 984 | B |
| ILE | 468 | A | LYS | 113 | B |
| GLY | 971 | A | TYR | 756 | B |
| GLN | 787 | A | ALA | 701 | C |
| ASN | 709 | A | ASP | 796 | B |
| TYR | 789 | A | ALA | 701 | C |
| ASN | 764 | A | GLN | 314 | C |
| ARG | 466 | A | GLN | 115 | B |
| LYS | 790 | A | SER | 704 | C |
| GLN | 613 | A | LEU | 861 | B |
| GLY | 880 | A | VAL | 705 | C |
| PHE | 592 | A | PHE | 855 | B |
| ALA | 647 | A | PRO | 862 | B |
| ILE | 882 | A | TYR | 707 | C |
| TYR | 789 | A | ASN | 703 | C |
| GLU | 516 | A | ASP | 228 | B |
| THR | 415 | A | TYR | 369 | B |
| ASP | 979 | A | ARG | 567 | C |
| GLN | 1002 | A | LEU | 1001 | B |
| SER | 375 | A | ARG | 408 | C |
| SER | 968 | A | GLN | 755 | B |
| VAL | 1128 | A | ASN | 919 | B |
| GLY | 566 | A | ARG | 44 | B |
| VAL | 1040 | A | SER | 1030 | B |
| MET | 697 | A | MET | 869 | B |
| GLY | 199 | A | PRO | 463 | C |
| ILE | 1013 | A | ALA | 1016 | B |
| TYR | 756 | A | GLN | 965 | C |
| LYS | 41 | A | PHE | 562 | C |
| ASN | 282 | A | LEU | 560 | C |
| ALA | 570 | A | SER | 967 | B |
| TYR | 707 | A | PRO | 897 | B |
| THR | 866 | A | GLY | 669 | C |
| MET | 740 | A | PHE | 592 | C |
| PHE | 43 | A | GLY | 566 | C |
| ASP | 568 | A | PHE | 855 | B |
| ASP | 1041 | A | SER | 1030 | B |
| ALA | 713 | A | ALA | 893 | B |
| GLN | 895 | A | ASN | 1074 | C |
| PHE | 1121 | A | ASN | 914 | B |
| TYR | 200 | A | ARG | 355 | C |
| ARG | 319 | A | ASP | 745 | B |
| GLY | 381 | A | ARG | 983 | B |
| ILE | 794 | A | TYR | 707 | C |
| ALA | 668 | A | LEU | 864 | B |
| PHE | 562 | A | ASP | 40 | B |
| GLN | 314 | A | THR | 768 | B |
| GLY | 381 | A | ILE | 973 | B |
| GLY | 381 | A | LEU | 984 | B |
| ASP | 571 | A | VAL | 976 | B |
| LEU | 1034 | A | VAL | 1040 | C |
| LEU | 1141 | A | LEU | 1141 | C |
| ASP | 571 | A | SER | 975 | B |
| GLN | 762 | A | GLN | 1010 | C |
| ASP | 1041 | A | ALA | 890 | B |
| ILE | 569 | A | VAL | 47 | B |
| ILE | 569 | A | LYS | 964 | B |
| TYR | 200 | A | SER | 514 | C |
| VAL | 1040 | A | GLU | 1031 | B |
| PRO | 862 | A | ALA | 647 | C |
| ASN | 703 | A | GLN | 787 | B |
| TYR | 789 | A | VAL | 705 | C |
| GLN | 762 | A | TYR | 1007 | C |
| GLY | 667 | A | PRO | 863 | B |
| ASP | 40 | A | HIS | 519 | C |
| ASP | 198 | A | ASP | 428 | C |
| THR | 961 | A | GLN | 762 | B |
| PHE | 464 | A | TYR | 200 | B |
| ASN | 856 | A | ALA | 570 | C |
| TYR | 756 | A | GLY | 971 | C |
| TYR | 1067 | A | ALA | 890 | B |
| GLY | 700 | A | ILE | 788 | B |
| ALA | 701 | A | ILE | 788 | B |
| SER | 982 | A | THR | 547 | C |
| SER | 373 | A | TYR | 505 | C |
| GLN | 755 | A | GLY | 971 | C |
| ASN | 317 | A | THR | 739 | B |
| THR | 739 | A | ARG | 319 | C |
| THR | 1009 | A | THR | 1009 | C |
| GLY | 548 | A | ASN | 978 | B |
| PHE | 1095 | A | MET | 900 | B |
| LYS | 417 | A | ASN | 370 | B |
| LYS | 41 | A | LEU | 518 | C |
| ILE | 233 | A | ARG | 466 | C |
| GLY | 669 | A | MET | 869 | B |
| MET | 869 | A | LEU | 699 | C |
| SER | 514 | A | TYR | 200 | B |
| PRO | 230 | A | ARG | 357 | C |
| ARG | 319 | A | THR | 739 | B |
| GLN | 965 | A | GLY | 757 | B |
| ASP | 427 | A | LYS | 986 | C |
| LYS | 558 | A | ASN | 282 | B |
| PRO | 589 | A | PHE | 855 | B |
| GLY | 667 | A | LEU | 864 | B |
| LEU | 984 | A | GLY | 381 | C |
| PHE | 759 | A | GLN | 965 | C |
| VAL | 987 | A | GLY | 413 | B |
| LEU | 1141 | A | GLU | 1144 | B |
| LYS | 786 | A | LYS | 1045 | C |
| GLY | 566 | A | PHE | 43 | B |
| THR | 961 | A | SER | 758 | B |
| LYS | 1045 | A | ALA | 890 | B |
| GLY | 283 | A | GLN | 563 | C |
| GLY | 232 | A | ARG | 466 | C |
| TYR | 707 | A | PHE | 797 | B |
| THR | 415 | A | ASP | 985 | C |
| LYS | 557 | A | SER | 45 | B |
| VAL | 42 | A | HIS | 519 | C |
| GLY | 283 | A | LYS | 558 | C |
| ASN | 914 | A | SER | 1123 | C |
| TYR | 505 | A | ALA | 372 | B |
| TRP | 886 | A | ARG | 1107 | C |
| LEU | 864 | A | CYS | 662 | C |
| LYS | 854 | A | ASP | 568 | C |
| ARG | 408 | A | TYR | 369 | B |
| GLU | 516 | A | LYS | 41 | B |
| GLY | 700 | A | LYS | 786 | B |
| ASP | 985 | A | SER | 383 | C |
| TYR | 904 | A | ARG | 1107 | C |
| LYS | 386 | A | ASP | 985 | B |
| THR | 696 | A | MET | 869 | B |
| ARG | 1039 | A | ARG | 1039 | B |
| PRO | 897 | A | SER | 711 | C |
| PHE | 43 | A | ARG | 567 | C |
| TYR | 917 | A | ILE | 1130 | C |
| MET | 900 | A | PHE | 1095 | C |
| PHE | 562 | A | TYR | 38 | B |
| ASN | 196 | A | LYS | 462 | C |
| PHE | 43 | A | PHE | 565 | C |
| GLU | 516 | A | TYR | 200 | B |
| VAL | 705 | A | GLN | 895 | B |
| ARG | 983 | A | LEU | 390 | C |
| PHE | 1121 | A | GLN | 913 | B |
| VAL | 382 | A | ARG | 983 | B |
| GLY | 413 | A | ASP | 985 | C |
| ASP | 994 | A | GLY | 971 | C |
| LYS | 462 | A | ASN | 196 | B |
| LYS | 1045 | A | GLY | 891 | B |
| GLU | 702 | A | TYR | 789 | B |
| ARG | 983 | A | LEU | 517 | C |
| SER | 711 | A | GLN | 895 | B |
| LEU | 560 | A | THR | 284 | B |
| ASP | 737 | A | ASN | 317 | C |
| VAL | 976 | A | ASP | 571 | C |
| ASP | 796 | A | SER | 708 | C |
| SER | 975 | A | ASP | 571 | C |
| ASP | 40 | A | GLN | 563 | C |
| ILE | 1013 | A | ILE | 1013 | B |
| LYS | 558 | A | GLY | 283 | B |
| SER | 711 | A | PRO | 897 | B |
| GLN | 895 | A | VAL | 705 | C |
| LEU | 560 | A | ASN | 282 | B |
| GLN | 787 | A | GLU | 702 | C |
| TYR | 396 | A | ASP | 228 | B |
| THR | 385 | A | THR | 415 | C |
| ALA | 701 | A | TYR | 789 | B |
| LYS | 790 | A | ASN | 703 | C |
| ALA | 890 | A | TYR | 1047 | C |
| ASP | 198 | A | PRO | 426 | C |
| LEU | 966 | A | ALA | 570 | C |
| ASN | 317 | A | ASP | 737 | B |
| PRO | 1069 | A | ALA | 892 | B |
| ASP | 228 | A | GLU | 516 | C |
| ARG | 1107 | A | TRP | 886 | B |
| ARG | 355 | A | TYR | 200 | B |
| GLU | 1111 | A | SER | 1123 | C |
| ARG | 1107 | A | ASN | 907 | B |
| GLU | 132 | A | ILE | 468 | C |
| ASN | 703 | A | TYR | 789 | B |
| LEU | 1012 | A | ILE | 1013 | C |
| GLU | 988 | A | CYS | 379 | C |
| PHE | 565 | A | PHE | 43 | B |
| ARG | 765 | A | GLN | 957 | C |
| SER | 383 | A | ARG | 983 | B |
| TYR | 369 | A | THR | 415 | C |
| THR | 768 | A | GLN | 314 | C |
| GLU | 1031 | A | PHE | 1042 | C |
| LYS | 854 | A | PHE | 592 | C |
| PRO | 1069 | A | GLY | 891 | B |
| THR | 859 | A | PHE | 592 | C |
| GLU | 702 | A | LYS | 790 | B |
| ALA | 713 | A | LEU | 894 | B |
| TYR | 756 | A | PHE | 970 | C |
| SER | 383 | A | GLU | 988 | B |
| ASN | 370 | A | LYS | 417 | C |
| GLU | 918 | A | PHE | 1089 | C |
| GLN | 563 | A | ARG | 44 | B |
| LEU | 390 | A | ARG | 983 | B |
| GLN | 965 | A | PHE | 759 | B |
| CYS | 671 | A | LEU | 864 | B |
| ASN | 914 | A | PHE | 1121 | C |
| PRO | 1079 | A | MET | 900 | B |
| LYS | 1038 | A | GLN | 1036 | B |
| GLU | 988 | A | VAL | 382 | C |
| ILE | 896 | A | SER | 711 | C |
| PHE | 759 | A | GLN | 1002 | C |
| LEU | 699 | A | ILE | 870 | B |
| GLN | 314 | A | SER | 735 | B |
| VAL | 705 | A | THR | 883 | B |
| PHE | 592 | A | THR | 859 | B |
| LYS | 854 | A | ILE | 569 | C |
| SER | 735 | A | GLN | 613 | C |
| VAL | 1040 | A | GLY | 889 | B |
| CYS | 662 | A | LEU | 864 | B |
| LEU | 864 | A | ALA | 668 | C |
| PRO | 792 | A | TYR | 707 | C |
| GLY | 757 | A | SER | 968 | C |
| ASP | 389 | A | SER | 982 | B |
| LYS | 386 | A | LEU | 984 | B |
| LEU | 865 | A | ALA | 668 | C |
| ARG | 1107 | A | LEU | 894 | B |
| LEU | 984 | A | SER | 383 | C |
| VAL | 1128 | A | TYR | 917 | B |
| ILE | 666 | A | PRO | 862 | B |
| LEU | 864 | A | ILE | 666 | C |
| MET | 869 | A | MET | 697 | C |
| PHE | 898 | A | TYR | 707 | C |
| LYS | 41 | A | GLN | 564 | C |
| GLU | 988 | A | LYS | 378 | C |
| ARG | 1019 | A | GLU | 1017 | C |
| ARG | 1091 | A | GLN | 913 | B |
| ILE | 973 | A | GLY | 381 | C |
| ASP | 198 | A | PRO | 463 | C |
| VAL | 42 | A | GLN | 563 | C |
| PHE | 759 | A | GLY | 999 | C |
| PRO | 463 | A | ASP | 198 | B |
| LEU | 1145 | A | GLU | 1144 | B |
| THR | 1077 | A | PRO | 897 | B |
| PHE | 759 | A | THR | 1006 | C |
| ARG | 1107 | A | TYR | 904 | B |
| VAL | 42 | A | PHE | 565 | C |
| LYS | 386 | A | SER | 982 | B |
| SER | 1030 | A | VAL | 1040 | C |
| PRO | 862 | A | GLY | 667 | C |
| PHE | 592 | A | MET | 740 | B |
| MET | 697 | A | LEU | 865 | B |
| ILE | 788 | A | GLY | 700 | C |
| GLU | 1072 | A | LEU | 894 | B |
| LEU | 699 | A | LEU | 865 | B |
| ASN | 370 | A | GLY | 416 | C |
| ILE | 714 | A | LEU | 894 | B |
| PRO | 230 | A | ARG | 355 | C |
| ASP | 198 | A | PHE | 464 | C |
| THR | 887 | A | TYR | 1047 | C |
| ASN | 856 | A | ASP | 568 | C |
| LEU | 894 | A | ALA | 713 | C |
| LYS | 790 | A | GLU | 702 | C |
| GLU | 1031 | A | ARG | 1039 | C |
| ILE | 1130 | A | GLN | 920 | B |
| ALA | 899 | A | TYR | 707 | C |
| GLN | 1005 | A | GLN | 1002 | C |
| GLU | 1144 | A | LEU | 1141 | C |
| PRO | 897 | A | ILE | 712 | C |
| ALA | 1016 | A | ARG | 1019 | B |
| VAL | 705 | A | TYR | 789 | B |
| ALA | 893 | A | ALA | 713 | C |
| ASN | 710 | A | PRO | 897 | B |
| VAL | 785 | A | LEU | 699 | C |
| LEU | 894 | A | GLU | 1072 | C |
| VAL | 705 | A | ALA | 879 | B |
| SER | 758 | A | LYS | 964 | C |
| ASN | 1108 | A | TRP | 886 | B |
| PHE | 970 | A | PHE | 759 | B |
| ALA | 890 | A | LYS | 1045 | C |
| GLN | 1010 | A | ALA | 766 | B |
| GLU | 702 | A | GLN | 787 | B |
| GLN | 1010 | A | GLN | 762 | B |
| SER | 1030 | A | ASP | 1041 | C |
| ARG | 355 | A | PRO | 230 | B |
| ILE | 569 | A | SER | 967 | B |
| GLN | 314 | A | ASN | 764 | B |
| TYR | 200 | A | PHE | 464 | C |
| GLN | 613 | A | SER | 735 | B |
| TYR | 707 | A | SER | 884 | B |
| ALA | 890 | A | PRO | 1069 | C |
| SER | 711 | A | ILE | 896 | B |
| ALA | 372 | A | TYR | 505 | C |
| LYS | 786 | A | ALA | 701 | C |
| ILE | 569 | A | LYS | 854 | B |
| ARG | 319 | A | MET | 740 | B |
| PHE | 1089 | A | TYR | 917 | B |
| THR | 385 | A | ASP | 985 | B |
| GLN | 913 | A | PRO | 1090 | C |
| THR | 1006 | A | GLN | 762 | B |
| THR | 883 | A | TYR | 707 | C |
| ILE | 1130 | A | LYS | 921 | B |
| ALA | 892 | A | GLU | 1072 | C |
| TYR | 789 | A | GLU | 702 | C |
| GLN | 755 | A | PHE | 970 | C |
| VAL | 963 | A | ALA | 570 | C |
| MET | 900 | A | ILE | 712 | C |
| GLU | 702 | A | ILE | 788 | B |
| ALA | 372 | A | ARG | 403 | C |
| ASN | 703 | A | LYS | 790 | B |
| GLN | 1005 | A | THR | 1006 | C |
| ASP | 985 | A | LYS | 386 | C |
| PHE | 1042 | A | GLU | 1031 | B |
| TYR | 917 | A | PHE | 1089 | C |
| GLY | 416 | A | TYR | 369 | B |
| GLN | 895 | A | ALA | 713 | C |
| TYR | 707 | A | PRO | 792 | B |
| ILE | 788 | A | GLU | 702 | C |
| ARG | 44 | A | GLN | 563 | C |
| THR | 547 | A | ASN | 978 | B |
| THR | 572 | A | ASN | 856 | B |
| GLU | 1031 | A | VAL | 1040 | C |
| PRO | 225 | A | PHE | 562 | C |
| SER | 383 | A | ASP | 985 | B |
| LYS | 378 | A | GLU | 988 | B |
| VAL | 987 | A | GLN | 414 | B |
| THR | 415 | A | THR | 385 | B |
| LYS | 786 | A | LEU | 699 | C |
| TYR | 917 | A | PRO | 1079 | C |
| GLN | 965 | A | SER | 758 | B |
| PHE | 759 | A | PHE | 970 | C |
| TYR | 1047 | A | TRP | 886 | B |
| LEU | 894 | A | ILE | 714 | C |
| PRO | 715 | A | LEU | 894 | B |
| GLN | 115 | A | ILE | 468 | C |
| LEU | 1001 | A | GLN | 1002 | C |
| TYR | 917 | A | VAL | 1129 | C |
| GLY | 1046 | A | GLY | 889 | B |
| LEU | 699 | A | VAL | 785 | B |
| MET | 900 | A | ALA | 1078 | C |
| LEU | 517 | A | ARG | 983 | B |
| PHE | 1089 | A | GLN | 913 | B |
| ALA | 570 | A | LEU | 966 | B |
| ASP | 796 | A | TYR | 707 | C |
| PHE | 464 | A | ASP | 198 | B |
| GLN | 563 | A | ASP | 40 | B |
| VAL | 1068 | A | ALA | 890 | B |
| ALA | 701 | A | LYS | 786 | B |
| ARG | 983 | A | SER | 383 | C |
| SER | 1123 | A | GLU | 1111 | B |
| ILE | 788 | A | ALA | 701 | C |
| VAL | 1040 | A | GLY | 1035 | B |
| TYR | 396 | A | TYR | 200 | B |
| LEU | 858 | A | PHE | 592 | C |
| ASP | 1041 | A | GLN | 784 | B |
| ASP | 796 | A | ASN | 709 | C |
| VAL | 976 | A | THR | 572 | C |
| ARG | 1039 | A | ARG | 1039 | C |
| TYR | 707 | A | ILE | 882 | B |
| TYR | 707 | A | GLN | 895 | B |
| HIS | 519 | A | VAL | 42 | B |
| ASN | 914 | A | PHE | 1089 | C |
| ARG | 319 | A | ASP | 737 | B |
| LEU | 984 | A | LYS | 386 | C |
| GLY | 891 | A | PRO | 1069 | C |
| SER | 884 | A | TYR | 707 | C |
| PRO | 1069 | A | ALA | 890 | B |
| VAL | 1129 | A | GLU | 918 | B |
| GLY | 566 | A | VAL | 42 | B |
| VAL | 705 | A | SER | 884 | B |
| ASP | 228 | A | LEU | 518 | C |
| ASP | 985 | A | THR | 415 | B |
| LYS | 1045 | A | LYS | 786 | B |
| GLY | 889 | A | VAL | 1040 | C |
| GLU | 988 | A | TYR | 380 | C |
| ALA | 570 | A | ASN | 856 | B |
| THR | 430 | A | ARG | 983 | B |
| GLN | 1002 | A | GLN | 1002 | C |
| TYR | 756 | A | GLN | 1002 | C |
| THR | 572 | A | VAL | 976 | B |
| LEU | 699 | A | ILE | 788 | B |
| ARG | 765 | A | THR | 961 | C |
| ILE | 788 | A | ASN | 703 | C |
| PRO | 863 | A | GLY | 667 | C |
| ASP | 228 | A | TYR | 396 | C |
| TRP | 353 | A | GLY | 232 | B |
| SER | 967 | A | ASP | 571 | C |
| ALA | 668 | A | LEU | 865 | B |
| THR | 1006 | A | GLN | 1005 | B |
| ILE | 1013 | A | THR | 1009 | B |
| ARG | 646 | A | PRO | 862 | B |
| SER | 708 | A | ASP | 796 | B |
| GLY | 971 | A | GLN | 755 | B |
| GLY | 232 | A | GLU | 465 | C |
| LEU | 861 | A | GLN | 613 | C |
| SER | 982 | A | LYS | 386 | C |
| PRO | 412 | A | VAL | 987 | C |
| LEU | 699 | A | GLN | 872 | B |
| ASP | 571 | A | SER | 967 | B |
| LEU | 984 | A | VAL | 382 | C |
| VAL | 1094 | A | TYR | 904 | B |
| LEU | 864 | A | GLY | 669 | C |
| LEU | 518 | A | LYS | 202 | B |
| THR | 859 | A | GLN | 613 | C |
| ASP | 568 | A | LYS | 854 | B |
| GLY | 232 | A | PHE | 464 | C |
| ILE | 712 | A | ILE | 896 | B |
| GLN | 895 | A | TYR | 707 | C |
| LEU | 865 | A | GLY | 669 | C |
| CYS | 379 | A | GLU | 988 | B |
| ARG | 466 | A | ILE | 233 | B |
| SER | 967 | A | ILE | 569 | C |
| GLY | 889 | A | LYS | 1045 | C |
| TYR | 200 | A | TYR | 396 | C |
| GLY | 1093 | A | TYR | 904 | B |
| THR | 1027 | A | ARG | 1039 | C |
| VAL | 705 | A | ALA | 893 | B |
| ALA | 570 | A | LYS | 964 | B |
| PHE | 970 | A | TYR | 756 | B |
| GLU | 224 | A | LEU | 560 | C |
| PRO | 897 | A | TYR | 707 | C |
| ARG | 1039 | A | GLU | 1031 | B |
| PHE | 562 | A | GLU | 224 | B |
| ILE | 231 | A | ARG | 466 | C |
| VAL | 382 | A | GLU | 988 | B |
| PRO | 862 | A | ILE | 666 | C |
| THR | 961 | A | ARG | 765 | B |
| GLU | 224 | A | PHE | 562 | C |
| ALA | 713 | A | ILE | 896 | B |
| LEU | 1145 | A | LEU | 1145 | C |
| PRO | 792 | A | VAL | 705 | C |
| GLN | 755 | A | ASN | 969 | C |
| GLY | 757 | A | GLN | 965 | C |
| VAL | 976 | A | ARG | 567 | C |
| GLN | 913 | A | ARG | 1091 | C |
| PHE | 855 | A | THR | 588 | C |
| SER | 1003 | A | PHE | 759 | B |
| THR | 1006 | A | PHE | 759 | B |
| PRO | 665 | A | LEU | 864 | B |
| GLY | 669 | A | LEU | 864 | B |
| PHE | 43 | A | LYS | 557 | C |
| GLY | 798 | A | TYR | 707 | C |
| PHE | 592 | A | LYS | 854 | B |
| SER | 975 | A | ALA | 570 | C |
| LYS | 113 | A | ILE | 468 | C |
| LEU | 894 | A | PRO | 715 | C |
| PHE | 43 | A | LYS | 558 | C |
| GLN | 787 | A | ASN | 703 | C |
| GLN | 1002 | A | GLN | 1002 | B |
| TYR | 789 | A | SER | 704 | C |
| ILE | 788 | A | LEU | 699 | C |
| THR | 866 | A | ALA | 668 | C |
| GLY | 1046 | A | ALA | 890 | B |
| GLY | 700 | A | TYR | 873 | B |
| ASP | 994 | A | PHE | 970 | C |
| VAL | 1129 | A | TYR | 917 | B |
| TRP | 886 | A | ASN | 1108 | C |
| PRO | 863 | A | ALA | 668 | C |
| ALA | 372 | A | GLU | 406 | C |
| TYR | 38 | A | PHE | 562 | C |
| PHE | 565 | A | VAL | 42 | B |
| ALA | 520 | A | LYS | 41 | B |
| GLN | 762 | A | THR | 961 | C |
| VAL | 705 | A | LYS | 790 | B |
| GLY | 667 | A | PRO | 862 | B |
| GLN | 1036 | A | LYS | 1038 | C |
| VAL | 987 | A | ASP | 427 | B |
| THR | 1077 | A | MET | 900 | B |
| ASP | 745 | A | ARG | 319 | C |
| SER | 1147 | A | LEU | 1145 | C |
| GLN | 965 | A | TYR | 756 | B |
| PHE | 797 | A | TYR | 707 | C |
| GLY | 413 | A | VAL | 987 | C |
| VAL | 705 | A | PRO | 792 | B |
| THR | 284 | A | LEU | 560 | C |
| VAL | 963 | A | ILE | 569 | C |
| ILE | 896 | A | SER | 708 | C |
| GLU | 918 | A | SER | 1123 | C |
| ALA | 879 | A | VAL | 705 | C |
| ARG | 567 | A | ARG | 44 | B |
| PRO | 862 | A | ARG | 646 | C |
| THR | 547 | A | SER | 982 | B |
| LYS | 558 | A | PHE | 43 | B |
| GLN | 965 | A | GLN | 762 | B |
| ALA | 570 | A | VAL | 963 | B |
| SER | 967 | A | ALA | 570 | C |
| ILE | 1013 | A | LEU | 1012 | B |
| ALA | 706 | A | GLN | 895 | B |
| TYR | 904 | A | VAL | 1094 | C |
| HIS | 519 | A | LYS | 41 | B |
| ILE | 1130 | A | TYR | 917 | B |
| LEU | 754 | A | GLN | 52 | C |
| GLN | 414 | A | TYR | 369 | B |
| LEU | 699 | A | LYS | 786 | B |
| PHE | 464 | A | ILE | 233 | B |
| PRO | 897 | A | ASN | 710 | C |
| TYR | 707 | A | ALA | 899 | B |
| GLN | 762 | A | THR | 1006 | C |
| GLY | 199 | A | PHE | 464 | C |
| ASP | 568 | A | ASN | 856 | B |
| ASN | 282 | A | LYS | 558 | C |
| LYS | 964 | A | ALA | 570 | C |
| LYS | 41 | A | GLN | 563 | C |
| TYR | 369 | A | GLY | 416 | C |
| ALA | 570 | A | VAL | 976 | B |
| LEU | 1141 | A | LEU | 1145 | B |
| ALA | 713 | A | GLN | 895 | B |
| VAL | 976 | A | ALA | 570 | C |
| GLU | 988 | A | SER | 383 | C |
| GLN | 895 | A | ALA | 706 | C |
| ARG | 466 | A | ILE | 231 | B |
| GLY | 1035 | A | VAL | 1040 | C |
| ASP | 737 | A | PHE | 592 | C |
| GLY | 545 | A | SER | 982 | B |
| ILE | 788 | A | SER | 704 | C |
| SER | 968 | A | TYR | 756 | B |
| TYR | 904 | A | ILE | 712 | C |
| PHE | 374 | A | ARG | 408 | C |
| PRO | 1090 | A | GLN | 913 | B |
| PHE | 43 | A | GLN | 563 | C |
| SER | 758 | A | THR | 961 | C |
| TYR | 917 | A | VAL | 1128 | C |
| ASP | 571 | A | SER | 968 | B |
| GLN | 913 | A | PHE | 1089 | C |
| SER | 704 | A | TYR | 789 | B |
| PHE | 592 | A | ASN | 856 | B |
| ASN | 978 | A | GLY | 548 | C |
| LEU | 1145 | A | LEU | 1141 | B |
| LYS | 790 | A | VAL | 705 | C |
| ASN | 960 | A | ILE | 569 | C |
| PHE | 464 | A | GLY | 199 | B |
| GLY | 1093 | A | GLN | 913 | B |
| TYR | 707 | A | THR | 883 | B |
| GLN | 1002 | A | GLN | 1005 | B |
| ALA | 1078 | A | MET | 900 | B |
| LEU | 864 | A | GLY | 667 | C |
| SER | 982 | A | ASP | 389 | C |
| LYS | 1045 | A | GLY | 889 | B |
| TYR | 917 | A | ALA | 1080 | C |
| MET | 697 | A | TYR | 873 | B |
| THR | 547 | A | LEU | 981 | B |
| GLN | 895 | A | SER | 711 | C |
| SER | 708 | A | GLN | 895 | B |
| SER | 704 | A | LYS | 790 | B |
| ASN | 856 | A | PHE | 592 | C |
| ALA | 893 | A | VAL | 705 | C |
| LEU | 981 | A | THR | 547 | C |
| TYR | 756 | A | SER | 968 | C |
| ARG | 1039 | A | THR | 1027 | B |
| MET | 900 | A | PRO | 1079 | C |
| SER | 704 | A | ILE | 788 | B |
| GLN | 1002 | A | PHE | 759 | B |
| LEU | 1012 | A | GLN | 1010 | C |
| GLY | 891 | A | VAL | 1068 | C |
| ARG | 44 | A | GLY | 566 | C |
| ASP | 737 | A | ARG | 319 | C |
| VAL | 705 | A | GLY | 880 | B |
| PHE | 562 | A | PRO | 225 | B |
| LEU | 699 | A | MET | 869 | B |
| VAL | 1094 | A | MET | 900 | B |
| TYR | 707 | A | GLY | 798 | B |
| VAL | 42 | A | ARG | 567 | C |
| LYS | 41 | A | ALA | 520 | C |
| LYS | 41 | A | PHE | 565 | C |
| SER | 708 | A | ILE | 896 | B |
| ILE | 712 | A | MET | 900 | B |
| ASP | 985 | A | THR | 385 | C |
| PHE | 592 | A | GLY | 857 | B |
| GLN | 115 | A | ARG | 466 | C |
| GLY | 669 | A | LEU | 865 | B |
| ILE | 1130 | A | GLY | 798 | B |
| SER | 383 | A | LEU | 984 | B |
| GLN | 787 | A | GLY | 700 | C |
| GLY | 857 | A | PHE | 592 | C |
| PRO | 862 | A | ALA | 668 | C |
| LEU | 864 | A | CYS | 671 | C |
| LEU | 560 | A | TYR | 38 | B |
| PHE | 464 | A | GLY | 232 | B |
| SER | 708 | A | PRO | 897 | B |
| ILE | 712 | A | TYR | 904 | B |
| TYR | 380 | A | GLU | 988 | B |
| GLN | 920 | A | ILE | 1130 | C |
| ASN | 978 | A | THR | 547 | C |
| PRO | 897 | A | THR | 1077 | C |
| ILE | 896 | A | ALA | 713 | C |
| PHE | 1089 | A | GLU | 918 | B |
| ARG | 567 | A | VAL | 42 | B |
| GLN | 563 | A | VAL | 42 | B |
| GLY | 232 | A | TRP | 353 | C |
| GLU | 1017 | A | ARG | 1019 | B |
| LEU | 1034 | A | ASP | 1041 | C |
| ASP | 428 | A | ASP | 198 | B |
| MET | 869 | A | THR | 696 | C |
| PHE | 43 | A | LEU | 560 | C |
| THR | 588 | A | PHE | 855 | B |
| TYR | 1007 | A | GLN | 762 | B |
| GLY | 1124 | A | GLU | 918 | B |
| PRO | 1079 | A | GLN | 913 | B |
| LEU | 1141 | A | LEU | 1141 | B |
| PHE | 1121 | A | THR | 912 | B |
| ILE | 468 | A | GLU | 132 | B |
| LEU | 864 | A | PRO | 665 | C |
| LYS | 386 | A | LEU | 981 | B |
| ALA | 668 | A | PRO | 863 | B |
| ASN | 978 | A | GLY | 545 | C |
| ILE | 712 | A | GLN | 895 | B |
| GLN | 563 | A | TYR | 38 | B |
| LEU | 894 | A | VAL | 705 | C |
| PRO | 384 | A | GLU | 988 | B |
| GLU | 1144 | A | LEU | 1145 | C |
| GLN | 563 | A | LYS | 41 | B |
| ARG | 983 | A | THR | 430 | C |
| VAL | 1040 | A | GLN | 1036 | B |
| LEU | 894 | A | PRO | 1069 | C |
| ALA | 890 | A | GLY | 1046 | C |
| GLY | 700 | A | GLN | 787 | B |
| SER | 982 | A | LEU | 390 | C |
| PRO | 426 | A | ASP | 198 | B |
| VAL | 47 | A | ILE | 569 | C |
| TYR | 200 | A | GLU | 516 | C |
| LEU | 1141 | A | LEU | 1145 | C |
| LEU | 699 | A | TYR | 873 | B |
| SER | 1123 | A | GLU | 918 | B |
| MET | 697 | A | GLN | 779 | B |
| LYS | 964 | A | ILE | 569 | C |
| ASP | 979 | A | LEU | 546 | C |
| ARG | 44 | A | ARG | 567 | C |
| LYS | 921 | A | ILE | 1130 | C |
| VAL | 1040 | A | LEU | 1034 | B |

**Table S2.** The list of the inter-protomer contacts of the protomer A in the structure of SARS-CoV-2 S-GSAS/D614 in the open state (pdb id 7KDH).

| **S protein residue name** | **S protein residue number** | **S protein chain** | **S protein residue name** | **S protein residue number** | **S protein chain** |
| --- | --- | --- | --- | --- | --- |
| GLN | 563 | A | ARG | 44 | B |
| ARG | 319 | A | MET | 740 | B |
| ALA | 570 | A | LEU | 966 | B |
| ARG | 44 | A | ASP | 571 | C |
| GLY | 381 | A | ARG | 983 | B |
| TYR | 873 | A | MET | 697 | C |
| ASP | 737 | A | ASN | 317 | C |
| ILE | 1130 | A | LYS | 921 | B |
| TYR | 200 | A | TYR | 396 | C |
| LEU | 984 | A | GLY | 381 | C |
| LEU | 699 | A | LEU | 865 | B |
| ASN | 703 | A | TYR | 789 | B |
| ARG | 1091 | A | ASP | 1118 | B |
| ASP | 40 | A | PHE | 562 | C |
| GLN | 784 | A | LYS | 1045 | C |
| LEU | 1034 | A | ASP | 1041 | C |
| VAL | 1094 | A | ILE | 896 | B |
| ILE | 1013 | A | LEU | 1012 | B |
| GLU | 1072 | A | ALA | 892 | B |
| GLN | 755 | A | PHE | 970 | C |
| LYS | 386 | A | LEU | 984 | B |
| SER | 967 | A | ILE | 569 | C |
| PHE | 43 | A | PHE | 559 | C |
| ARG | 995 | A | ASP | 994 | B |
| VAL | 976 | A | ARG | 567 | C |
| PHE | 1089 | A | GLN | 913 | B |
| TYR | 873 | A | LEU | 699 | C |
| GLN | 787 | A | GLY | 700 | C |
| TYR | 707 | A | PHE | 898 | B |
| LYS | 41 | A | PRO | 521 | C |
| GLY | 889 | A | VAL | 1040 | C |
| TYR | 756 | A | PHE | 970 | C |
| LEU | 894 | A | ALA | 713 | C |
| ASN | 978 | A | GLY | 545 | C |
| ARG | 1039 | A | THR | 1027 | B |
| PHE | 1121 | A | THR | 912 | B |
| THR | 1006 | A | GLN | 762 | B |
| PRO | 230 | A | ARG | 355 | C |
| ALA | 713 | A | LEU | 894 | B |
| ALA | 879 | A | VAL | 705 | C |
| LEU | 699 | A | GLN | 787 | B |
| GLN | 613 | A | THR | 859 | B |
| ASP | 745 | A | ARG | 319 | C |
| ALA | 701 | A | ILE | 788 | B |
| ASN | 703 | A | LYS | 790 | B |
| TYR | 756 | A | ASN | 969 | C |
| TYR | 38 | A | PHE | 562 | C |
| ILE | 714 | A | LEU | 894 | B |
| ASN | 165 | A | ILE | 468 | C |
| THR | 284 | A | LEU | 560 | C |
| ILE | 788 | A | ALA | 701 | C |
| VAL | 1068 | A | GLY | 891 | B |
| ASN | 907 | A | ARG | 1107 | C |
| MET | 900 | A | ILE | 712 | C |
| ASP | 979 | A | LEU | 546 | C |
| ILE | 896 | A | SER | 711 | C |
| PHE | 43 | A | GLY | 566 | C |
| GLY | 798 | A | ILE | 1130 | C |
| TYR | 904 | A | ARG | 1107 | C |
| PRO | 1069 | A | ALA | 892 | B |
| LEU | 1034 | A | LYS | 1045 | C |
| ASN | 394 | A | ASP | 228 | B |
| GLY | 1093 | A | GLN | 913 | B |
| GLN | 965 | A | GLY | 757 | B |
| PRO | 1069 | A | LEU | 894 | B |
| PHE | 888 | A | LYS | 1045 | C |
| ALA | 570 | A | ASN | 856 | B |
| TYR | 707 | A | ALA | 899 | B |
| LEU | 1141 | A | LEU | 1141 | C |
| THR | 866 | A | ALA | 668 | C |
| GLN | 784 | A | ASP | 1041 | C |
| VAL | 705 | A | TYR | 789 | B |
| VAL | 705 | A | ALA | 879 | B |
| GLY | 757 | A | PHE | 970 | C |
| SER | 1003 | A | PHE | 759 | B |
| LEU | 546 | A | ASP | 979 | B |
| LEU | 1141 | A | LEU | 1145 | C |
| THR | 588 | A | PHE | 855 | B |
| LEU | 894 | A | PRO | 715 | C |
| ALA | 570 | A | VAL | 963 | B |
| LEU | 865 | A | MET | 697 | C |
| THR | 1077 | A | MET | 900 | B |
| ILE | 1013 | A | ALA | 1016 | B |
| GLN | 895 | A | ILE | 712 | C |
| LYS | 1045 | A | GLN | 784 | B |
| LEU | 699 | A | TYR | 873 | B |
| GLY | 700 | A | GLN | 787 | B |
| LEU | 864 | A | ILE | 666 | C |
| PRO | 1079 | A | MET | 900 | B |
| GLU | 1031 | A | ARG | 1039 | C |
| ALA | 701 | A | TYR | 789 | B |
| GLU | 702 | A | GLN | 872 | B |
| ASP | 994 | A | PHE | 970 | C |
| PRO | 1069 | A | GLY | 891 | B |
| ALA | 701 | A | GLN | 787 | B |
| GLU | 918 | A | VAL | 1128 | C |
| GLN | 965 | A | PHE | 759 | B |
| GLY | 669 | A | LEU | 865 | B |
| LEU | 699 | A | ILE | 788 | B |
| LEU | 864 | A | ILE | 670 | C |
| SER | 968 | A | GLN | 755 | B |
| GLY | 667 | A | PRO | 862 | B |
| VAL | 976 | A | ALA | 570 | C |
| GLY | 757 | A | SER | 968 | C |
| SER | 383 | A | GLU | 988 | B |
| GLY | 798 | A | TYR | 707 | C |
| ILE | 896 | A | SER | 708 | C |
| ASP | 737 | A | PHE | 592 | C |
| LYS | 786 | A | GLY | 700 | C |
| SER | 708 | A | ILE | 896 | B |
| CYS | 379 | A | GLU | 988 | B |
| ASP | 571 | A | SER | 967 | B |
| GLU | 702 | A | LYS | 790 | B |
| GLY | 971 | A | ASP | 994 | B |
| GLN | 913 | A | GLY | 1093 | C |
| PHE | 1089 | A | GLU | 918 | B |
| THR | 961 | A | SER | 758 | B |
| PRO | 862 | A | GLY | 667 | C |
| SER | 968 | A | GLY | 757 | B |
| TYR | 789 | A | GLU | 702 | C |
| ALA | 647 | A | PRO | 862 | B |
| LEU | 517 | A | ARG | 983 | B |
| ARG | 44 | A | ARG | 567 | C |
| GLY | 232 | A | PHE | 464 | C |
| LEU | 865 | A | GLY | 669 | C |
| THR | 415 | A | ASP | 985 | C |
| PHE | 43 | A | LEU | 560 | C |
| THR | 1009 | A | THR | 1009 | B |
| ALA | 890 | A | LYS | 1045 | C |
| ALA | 520 | A | LYS | 41 | B |
| PRO | 897 | A | SER | 711 | C |
| GLN | 965 | A | TYR | 756 | B |
| GLY | 566 | A | PHE | 43 | B |
| ALA | 892 | A | PRO | 1069 | C |
| THR | 167 | A | ARG | 466 | C |
| LYS | 386 | A | ARG | 983 | B |
| GLU | 1017 | A | ARG | 1019 | B |
| GLN | 314 | A | SER | 735 | B |
| LEU | 894 | A | VAL | 705 | C |
| GLN | 787 | A | ALA | 701 | C |
| LEU | 1145 | A | LEU | 1145 | C |
| PHE | 1042 | A | GLU | 1031 | B |
| LEU | 546 | A | VAL | 976 | B |
| GLY | 971 | A | TYR | 756 | B |
| LEU | 1012 | A | ILE | 1013 | C |
| ASN | 978 | A | GLY | 548 | C |
| THR | 1027 | A | ARG | 1039 | C |
| VAL | 705 | A | THR | 791 | B |
| ASP | 796 | A | ASN | 709 | C |
| TYR | 904 | A | ILE | 712 | C |
| THR | 912 | A | PHE | 1121 | C |
| LEU | 1145 | A | SER | 1147 | B |
| GLN | 762 | A | GLN | 965 | C |
| LEU | 858 | A | PHE | 592 | C |
| THR | 883 | A | TYR | 707 | C |
| GLY | 667 | A | LEU | 864 | B |
| GLY | 667 | A | PRO | 863 | B |
| SER | 711 | A | PRO | 897 | B |
| VAL | 1129 | A | TYR | 917 | B |
| SER | 982 | A | THR | 547 | C |
| ARG | 44 | A | GLY | 566 | C |
| LEU | 560 | A | THR | 284 | B |
| THR | 572 | A | ASN | 856 | B |
| GLU | 1072 | A | LEU | 894 | B |
| ASN | 856 | A | ALA | 570 | C |
| THR | 430 | A | ARG | 983 | B |
| PRO | 862 | A | ALA | 647 | C |
| PRO | 1079 | A | GLN | 913 | B |
| ARG | 567 | A | PHE | 43 | B |
| THR | 1006 | A | PHE | 759 | B |
| ASN | 317 | A | GLY | 857 | B |
| ARG | 1091 | A | ASN | 907 | B |
| LYS | 1045 | A | ALA | 890 | B |
| THR | 385 | A | ASP | 985 | B |
| TYR | 707 | A | PRO | 897 | B |
| SER | 884 | A | VAL | 705 | C |
| PRO | 897 | A | THR | 1077 | C |
| LEU | 1001 | A | GLN | 1002 | C |
| ASP | 574 | A | PHE | 855 | B |
| ASP | 994 | A | GLY | 971 | C |
| LEU | 894 | A | PRO | 1069 | C |
| ASP | 1041 | A | LEU | 1034 | B |
| LYS | 558 | A | GLY | 283 | B |
| GLN | 787 | A | GLU | 702 | C |
| THR | 961 | A | GLN | 762 | B |
| ALA | 713 | A | GLN | 895 | B |
| PRO | 897 | A | ASN | 710 | C |
| ASN | 856 | A | ASP | 568 | C |
| VAL | 1094 | A | TYR | 904 | B |
| PRO | 230 | A | ARG | 466 | C |
| LEU | 1145 | A | GLU | 1144 | B |
| GLY | 381 | A | LEU | 984 | B |
| ARG | 983 | A | VAL | 382 | C |
| ASP | 1041 | A | SER | 1030 | B |
| LYS | 386 | A | ASP | 985 | B |
| PHE | 855 | A | PHE | 592 | C |
| ILE | 670 | A | LEU | 864 | B |
| ALA | 668 | A | MET | 869 | B |
| ASP | 571 | A | VAL | 976 | B |
| GLY | 413 | A | LYS | 986 | C |
| PHE | 456 | A | SER | 383 | B |
| ALA | 1078 | A | MET | 900 | B |
| TYR | 38 | A | GLN | 563 | C |
| GLN | 895 | A | SER | 711 | C |
| LYS | 854 | A | ASP | 568 | C |
| ARG | 1039 | A | GLU | 1031 | B |
| VAL | 42 | A | PHE | 562 | C |
| SER | 383 | A | ARG | 983 | B |
| THR | 739 | A | ASN | 317 | C |
| ASP | 198 | A | PRO | 463 | C |
| THR | 887 | A | TYR | 1047 | C |
| VAL | 860 | A | ARG | 646 | C |
| PHE | 970 | A | GLN | 755 | B |
| TYR | 1047 | A | THR | 887 | B |
| GLN | 563 | A | ASP | 40 | B |
| ALA | 1070 | A | ALA | 892 | B |
| ILE | 712 | A | GLN | 895 | B |
| THR | 961 | A | ARG | 765 | B |
| ILE | 1130 | A | GLY | 798 | B |
| TYR | 707 | A | PHE | 797 | B |
| GLN | 913 | A | PHE | 1121 | C |
| SER | 383 | A | ASP | 985 | B |
| ASP | 737 | A | ARG | 319 | C |
| PRO | 862 | A | ARG | 646 | C |
| GLY | 857 | A | PHE | 592 | C |
| PHE | 759 | A | GLY | 999 | C |
| ASN | 709 | A | ASP | 796 | B |
| GLN | 895 | A | ALA | 713 | C |
| THR | 385 | A | THR | 415 | C |
| PRO | 1069 | A | ALA | 890 | B |
| TYR | 756 | A | ARG | 995 | C |
| SER | 758 | A | LYS | 964 | C |
| GLY | 413 | A | VAL | 987 | C |
| PHE | 759 | A | PHE | 970 | C |
| LYS | 1045 | A | GLY | 889 | B |
| PRO | 792 | A | TYR | 707 | C |
| TYR | 904 | A | VAL | 1094 | C |
| LYS | 41 | A | ALA | 520 | C |
| LYS | 964 | A | ILE | 569 | C |
| ALA | 899 | A | TYR | 707 | C |
| GLY | 700 | A | ILE | 788 | B |
| GLY | 891 | A | LYS | 1045 | C |
| GLN | 563 | A | GLY | 283 | B |
| THR | 1006 | A | GLN | 1005 | B |
| GLN | 1010 | A | ALA | 766 | B |
| SER | 1123 | A | ASN | 914 | B |
| ASP | 985 | A | PRO | 384 | C |
| ILE | 973 | A | GLY | 381 | C |
| LEU | 864 | A | PRO | 665 | C |
| GLU | 918 | A | SER | 1123 | C |
| ASN | 856 | A | THR | 572 | C |
| GLY | 669 | A | LEU | 864 | B |
| TYR | 707 | A | ILE | 794 | B |
| ALA | 570 | A | SER | 967 | B |
| TYR | 707 | A | ASP | 796 | B |
| ALA | 575 | A | PHE | 43 | B |
| SER | 45 | A | ARG | 567 | C |
| ASN | 234 | A | ARG | 466 | C |
| ASN | 394 | A | PRO | 230 | B |
| GLY | 232 | A | ARG | 466 | C |
| LYS | 557 | A | SER | 45 | B |
| LEU | 1034 | A | VAL | 1040 | C |
| SER | 704 | A | THR | 791 | B |
| THR | 572 | A | VAL | 976 | B |
| TYR | 707 | A | ILE | 896 | B |
| GLU | 918 | A | GLY | 1124 | C |
| PHE | 1121 | A | ASN | 914 | B |
| GLY | 548 | A | ASN | 978 | B |
| ARG | 1091 | A | GLN | 913 | B |
| GLN | 895 | A | TYR | 707 | C |
| GLN | 564 | A | LYS | 41 | B |
| TYR | 904 | A | GLY | 1093 | C |
| TYR | 917 | A | ILE | 1130 | C |
| THR | 883 | A | VAL | 705 | C |
| PHE | 592 | A | GLY | 857 | B |
| ILE | 569 | A | VAL | 47 | B |
| ASP | 571 | A | SER | 975 | B |
| TYR | 1047 | A | TRP | 886 | B |
| GLU | 988 | A | SER | 383 | C |
| ILE | 1130 | A | TYR | 917 | B |
| LYS | 41 | A | GLN | 564 | C |
| MET | 869 | A | GLY | 669 | C |
| PHE | 565 | A | VAL | 42 | B |
| VAL | 705 | A | GLN | 895 | B |
| GLN | 762 | A | THR | 961 | C |
| PHE | 562 | A | LYS | 41 | B |
| PRO | 589 | A | MET | 740 | B |
| ASP | 228 | A | TYR | 396 | C |
| GLN | 762 | A | SER | 1003 | C |
| LEU | 981 | A | LYS | 386 | C |
| PRO | 1140 | A | LEU | 1141 | C |
| LEU | 984 | A | VAL | 382 | C |
| LEU | 1145 | A | LEU | 1145 | B |
| THR | 1009 | A | ILE | 1013 | C |
| TYR | 756 | A | GLN | 965 | C |
| LEU | 894 | A | GLU | 1072 | C |
| SER | 982 | A | GLY | 545 | C |
| PRO | 384 | A | GLU | 988 | B |
| PHE | 759 | A | GLN | 1002 | C |
| GLN | 762 | A | GLN | 1010 | C |
| LYS | 41 | A | PHE | 562 | C |
| ALA | 570 | A | VAL | 976 | B |
| ILE | 712 | A | MET | 900 | B |
| MET | 740 | A | ARG | 319 | C |
| ILE | 666 | A | PRO | 862 | B |
| PRO | 665 | A | LEU | 864 | B |
| GLN | 563 | A | TYR | 279 | B |
| GLY | 757 | A | GLN | 965 | C |
| GLY | 891 | A | VAL | 1068 | C |
| ASP | 427 | A | GLU | 990 | C |
| GLU | 516 | A | ASP | 228 | B |
| VAL | 42 | A | GLY | 566 | C |
| SER | 1030 | A | PHE | 1042 | C |
| ARG | 983 | A | SER | 383 | C |
| GLN | 564 | A | VAL | 42 | B |
| GLN | 563 | A | VAL | 42 | B |
| VAL | 1068 | A | ALA | 890 | B |
| ALA | 1080 | A | TYR | 917 | B |
| THR | 547 | A | ASN | 978 | B |
| GLY | 889 | A | ASP | 1041 | C |
| LYS | 557 | A | PHE | 43 | B |
| TYR | 369 | A | GLN | 414 | C |
| ASP | 979 | A | THR | 547 | C |
| TYR | 707 | A | THR | 883 | B |
| GLU | 918 | A | PHE | 1089 | C |
| TYR | 756 | A | GLY | 971 | C |
| ARG | 1019 | A | GLU | 1017 | C |
| LYS | 790 | A | GLU | 702 | C |
| GLY | 700 | A | LYS | 786 | B |
| CYS | 671 | A | LEU | 864 | B |
| LYS | 786 | A | ALA | 701 | C |
| TYR | 1007 | A | GLN | 762 | B |
| GLN | 314 | A | THR | 768 | B |
| ILE | 712 | A | ILE | 896 | B |
| LEU | 560 | A | PHE | 43 | B |
| SER | 1123 | A | GLU | 1111 | B |
| VAL | 705 | A | PRO | 792 | B |
| LYS | 790 | A | SER | 704 | C |
| TYR | 707 | A | GLY | 798 | B |
| ALA | 706 | A | GLN | 895 | B |
| MET | 869 | A | MET | 697 | C |
| ARG | 44 | A | GLN | 563 | C |
| ASN | 969 | A | GLN | 755 | B |
| GLY | 199 | A | PHE | 464 | C |
| ARG | 765 | A | THR | 961 | C |
| GLU | 224 | A | PHE | 562 | C |
| GLU | 1031 | A | PHE | 1042 | C |
| PRO | 863 | A | GLY | 669 | C |
| PHE | 43 | A | LYS | 558 | C |
| ASP | 796 | A | SER | 708 | C |
| GLU | 773 | A | GLU | 1017 | C |
| ALA | 713 | A | ILE | 896 | B |
| ARG | 1107 | A | GLN | 913 | B |
| GLY | 999 | A | PHE | 759 | B |
| ALA | 701 | A | LYS | 786 | B |
| ASN | 317 | A | MET | 740 | B |
| GLN | 1010 | A | LEU | 1012 | B |
| PHE | 855 | A | ASP | 568 | C |
| ASP | 1041 | A | GLY | 889 | B |
| GLN | 115 | A | ARG | 466 | C |
| ASN | 710 | A | PRO | 897 | B |
| GLN | 913 | A | PRO | 1090 | C |
| GLY | 1124 | A | GLU | 918 | B |
| VAL | 785 | A | LEU | 699 | C |
| ASP | 796 | A | TYR | 707 | C |
| TYR | 789 | A | SER | 704 | C |
| GLN | 957 | A | ARG | 765 | B |
| ALA | 972 | A | GLN | 755 | B |
| LEU | 699 | A | LYS | 786 | B |
| GLN | 1036 | A | LYS | 1038 | C |
| TYR | 369 | A | ARG | 408 | C |
| ALA | 890 | A | VAL | 1040 | C |
| TYR | 200 | A | ARG | 355 | C |
| ASP | 1041 | A | GLN | 784 | B |
| PHE | 1089 | A | TYR | 917 | B |
| ASP | 994 | A | ARG | 995 | C |
| GLN | 895 | A | SER | 708 | C |
| PHE | 565 | A | PHE | 43 | B |
| LEU | 966 | A | ALA | 570 | C |
| GLN | 913 | A | PHE | 1089 | C |
| TYR | 789 | A | VAL | 705 | C |
| LEU | 864 | A | ALA | 668 | C |
| ILE | 794 | A | TYR | 707 | C |
| GLU | 1111 | A | SER | 1123 | C |
| ILE | 870 | A | LEU | 699 | C |
| PRO | 897 | A | SER | 708 | C |
| SER | 704 | A | TYR | 789 | B |
| GLN | 755 | A | ASN | 969 | C |
| GLN | 563 | A | TYR | 38 | B |
| PHE | 898 | A | TYR | 707 | C |
| ARG | 1039 | A | ARG | 1039 | C |
| ALA | 890 | A | GLY | 1046 | C |
| VAL | 976 | A | THR | 572 | C |
| ARG | 983 | A | GLY | 381 | C |
| LYS | 790 | A | VAL | 705 | C |
| LYS | 41 | A | GLN | 563 | C |
| ARG | 567 | A | ARG | 44 | B |
| LEU | 560 | A | TYR | 38 | B |
| GLN | 913 | A | ARG | 1091 | C |
| PRO | 412 | A | VAL | 987 | C |
| SER | 45 | A | LYS | 557 | C |
| GLY | 1046 | A | ALA | 890 | B |
| VAL | 1122 | A | ASN | 914 | B |
| LYS | 964 | A | SER | 758 | B |
| TYR | 369 | A | THR | 415 | C |
| ILE | 712 | A | PRO | 897 | B |
| LEU | 560 | A | GLY | 283 | B |
| MET | 697 | A | MET | 869 | B |
| LEU | 984 | A | TYR | 380 | C |
| THR | 547 | A | SER | 982 | B |
| GLN | 755 | A | SER | 968 | C |
| ASP | 1118 | A | ARG | 1091 | C |
| LEU | 984 | A | THR | 385 | C |
| GLU | 1092 | A | GLN | 913 | B |
| VAL | 382 | A | GLU | 988 | B |
| VAL | 1040 | A | SER | 1030 | B |
| TYR | 756 | A | SER | 968 | C |
| PHE | 855 | A | PRO | 589 | C |
| LEU | 865 | A | ALA | 668 | C |
| LEU | 864 | A | GLY | 667 | C |
| GLN | 1002 | A | THR | 998 | B |
| SER | 982 | A | LEU | 390 | C |
| TRP | 886 | A | ASN | 1108 | C |
| ASP | 40 | A | HIS | 519 | C |
| GLY | 1046 | A | GLY | 891 | B |
| THR | 167 | A | ILE | 468 | C |
| VAL | 42 | A | GLN | 563 | C |
| THR | 1009 | A | THR | 1009 | C |
| GLY | 413 | A | ASP | 985 | C |
| SER | 1030 | A | VAL | 1040 | C |
| ASP | 198 | A | PHE | 464 | C |
| TYR | 1047 | A | ALA | 890 | B |
| ALA | 668 | A | PRO | 863 | B |
| TYR | 707 | A | PRO | 792 | B |
| ALA | 668 | A | THR | 866 | B |
| GLU | 516 | A | TYR | 200 | B |
| ASN | 1108 | A | TRP | 886 | B |
| VAL | 1094 | A | MET | 900 | B |
| PRO | 897 | A | ILE | 712 | C |
| GLN | 755 | A | GLY | 971 | C |
| PHE | 43 | A | LYS | 557 | C |
| GLN | 314 | A | ASN | 764 | B |
| GLU | 1017 | A | GLU | 773 | B |
| TYR | 200 | A | PHE | 464 | C |
| MET | 900 | A | ALA | 1078 | C |
| ALA | 372 | A | ARG | 403 | C |
| SER | 968 | A | SER | 758 | B |
| GLY | 891 | A | PRO | 1069 | C |
| MET | 697 | A | TYR | 873 | B |
| TRP | 886 | A | TYR | 1047 | C |
| VAL | 42 | A | PHE | 565 | C |
| PRO | 230 | A | TYR | 396 | C |
| GLY | 669 | A | THR | 866 | B |
| TYR | 917 | A | VAL | 1128 | C |
| ALA | 668 | A | LEU | 864 | B |
| ARG | 1107 | A | TRP | 886 | B |
| ASN | 914 | A | SER | 1123 | C |
| PHE | 592 | A | PHE | 855 | B |
| ASP | 427 | A | VAL | 987 | C |
| PRO | 1090 | A | GLN | 913 | B |
| MET | 869 | A | ALA | 668 | C |
| THR | 547 | A | ASP | 979 | B |
| GLU | 1144 | A | LEU | 1141 | C |
| ALA | 893 | A | VAL | 705 | C |
| MET | 697 | A | LEU | 865 | B |
| TYR | 789 | A | ALA | 701 | C |
| PHE | 970 | A | TYR | 756 | B |
| GLY | 566 | A | VAL | 42 | B |
| SER | 982 | A | LEU | 546 | C |
| GLY | 889 | A | GLY | 1046 | C |
| SER | 708 | A | GLN | 895 | B |
| ASN | 282 | A | LEU | 560 | C |
| SER | 968 | A | TYR | 756 | B |
| TYR | 707 | A | GLN | 895 | B |
| ASP | 985 | A | THR | 385 | C |
| TYR | 707 | A | ILE | 882 | B |
| VAL | 1068 | A | ALA | 892 | B |
| SER | 967 | A | ASP | 571 | C |
| CYS | 662 | A | LEU | 864 | B |
| MET | 900 | A | THR | 1077 | C |
| SER | 982 | A | LYS | 386 | C |
| ILE | 231 | A | ARG | 466 | C |
| GLN | 787 | A | ASN | 703 | C |
| VAL | 705 | A | THR | 883 | B |
| ARG | 983 | A | LEU | 390 | C |
| GLY | 283 | A | LEU | 560 | C |
| PRO | 521 | A | LYS | 41 | B |
| PRO | 897 | A | TYR | 707 | C |
| THR | 859 | A | PHE | 592 | C |
| ILE | 1130 | A | GLN | 920 | B |
| GLU | 1072 | A | ALA | 893 | B |
| HIS | 519 | A | VAL | 42 | B |
| ARG | 567 | A | VAL | 976 | B |
| GLY | 232 | A | GLU | 465 | C |
| ARG | 646 | A | THR | 866 | B |
| ASN | 978 | A | THR | 547 | C |
| GLY | 1093 | A | TYR | 904 | B |
| LYS | 776 | A | LYS | 947 | C |
| GLY | 1046 | A | GLY | 889 | B |
| ARG | 1107 | A | TYR | 904 | B |
| SER | 708 | A | PRO | 897 | B |
| ASN | 703 | A | ILE | 788 | B |
| SER | 758 | A | THR | 961 | C |
| GLN | 1005 | A | GLN | 1002 | C |
| VAL | 963 | A | ALA | 570 | C |
| LEU | 699 | A | VAL | 785 | B |
| ARG | 765 | A | GLN | 957 | C |
| TYR | 369 | A | GLY | 416 | C |
| ILE | 788 | A | ASN | 703 | C |
| LYS | 790 | A | ASN | 703 | C |
| LYS | 386 | A | LEU | 981 | B |
| TYR | 396 | A | PRO | 230 | B |
| MET | 900 | A | PRO | 1079 | C |
| VAL | 705 | A | LYS | 790 | B |
| GLN | 895 | A | ASN | 1074 | C |
| PRO | 1079 | A | TYR | 917 | B |
| GLN | 920 | A | ILE | 1130 | C |
| GLY | 700 | A | TYR | 873 | B |
| VAL | 47 | A | ILE | 569 | C |
| VAL | 1128 | A | TYR | 917 | B |
| GLY | 880 | A | VAL | 705 | C |
| PHE | 970 | A | PHE | 759 | B |
| PHE | 562 | A | TYR | 38 | B |
| MET | 740 | A | PHE | 592 | C |
| PHE | 43 | A | PHE | 565 | C |
| THR | 739 | A | ARG | 319 | C |
| PHE | 592 | A | MET | 740 | B |
| LEU | 894 | A | ILE | 714 | C |
| ARG | 983 | A | LEU | 517 | C |
| ILE | 569 | A | LYS | 964 | B |
| THR | 696 | A | MET | 869 | B |
| HIS | 519 | A | ASP | 40 | B |
| ALA | 890 | A | VAL | 1068 | C |
| ILE | 788 | A | GLY | 700 | C |
| ALA | 668 | A | LEU | 865 | B |
| GLN | 115 | A | SER | 469 | C |
| PHE | 562 | A | PRO | 225 | B |
| VAL | 705 | A | ALA | 893 | B |
| VAL | 1040 | A | GLU | 1031 | B |
| GLN | 563 | A | PHE | 43 | B |
| ILE | 1013 | A | ILE | 1013 | B |
| GLN | 314 | A | LEU | 861 | B |
| PHE | 797 | A | TYR | 707 | C |
| TYR | 1067 | A | ALA | 890 | B |
| GLN | 895 | A | ALA | 706 | C |
| ARG | 1039 | A | ARG | 1039 | B |
| VAL | 1040 | A | GLY | 1035 | B |
| SER | 758 | A | SER | 968 | C |
| GLY | 283 | A | GLN | 563 | C |
| LYS | 964 | A | ALA | 570 | C |
| LEU | 865 | A | LEU | 699 | C |
| ILE | 788 | A | LEU | 699 | C |
| MET | 900 | A | PHE | 1095 | C |
| ILE | 882 | A | TYR | 707 | C |
| ASP | 198 | A | PRO | 426 | C |
| VAL | 382 | A | LEU | 984 | B |
| LEU | 1141 | A | LEU | 1141 | B |
| SER | 1003 | A | GLN | 762 | B |
| VAL | 705 | A | LEU | 894 | B |
| ILE | 896 | A | ILE | 712 | C |
| LYS | 41 | A | HIS | 519 | C |
| PRO | 225 | A | PHE | 562 | C |
| PHE | 759 | A | GLN | 965 | C |
| ALA | 520 | A | ASP | 228 | B |
| SER | 967 | A | ALA | 570 | C |
| ARG | 355 | A | TYR | 200 | B |
| ALA | 570 | A | SER | 975 | B |
| GLN | 762 | A | THR | 1006 | C |
| PHE | 759 | A | THR | 1006 | C |
| LEU | 864 | A | GLY | 669 | C |
| PRO | 863 | A | ALA | 668 | C |
| LEU | 390 | A | ARG | 983 | B |
| ARG | 765 | A | ALA | 958 | C |
| GLY | 891 | A | GLY | 1046 | C |
| LEU | 699 | A | MET | 869 | B |
| SER | 383 | A | LEU | 984 | B |
| ASP | 571 | A | SER | 968 | B |
| GLN | 115 | A | ASP | 467 | C |
| GLN | 1010 | A | GLN | 762 | B |
| LYS | 558 | A | ASN | 282 | B |
| PHE | 43 | A | GLN | 563 | C |
| ARG | 567 | A | VAL | 42 | B |
| VAL | 963 | A | ILE | 569 | C |
| HIS | 519 | A | LYS | 41 | B |
| GLU | 1144 | A | LEU | 1145 | C |
| SER | 698 | A | TYR | 873 | B |
| MET | 697 | A | LEU | 864 | B |
| ARG | 983 | A | LYS | 386 | C |
| ASP | 985 | A | SER | 383 | C |
| VAL | 382 | A | ARG | 983 | B |
| ILE | 569 | A | VAL | 963 | B |
| SER | 982 | A | ASP | 389 | C |
| THR | 768 | A | GLN | 314 | C |
| LEU | 560 | A | ASN | 282 | B |
| TYR | 789 | A | ASN | 703 | C |
| ALA | 890 | A | TYR | 1047 | C |
| VAL | 42 | A | HIS | 519 | C |
| ILE | 569 | A | SER | 967 | B |
| ASP | 571 | A | LEU | 966 | B |
| ARG | 1107 | A | ASN | 907 | B |
| GLN | 872 | A | LEU | 699 | C |
| PHE | 1089 | A | ASN | 914 | B |
| LYS | 41 | A | PHE | 565 | C |
| SER | 758 | A | GLN | 965 | C |
| TYR | 917 | A | PRO | 1079 | C |
| GLU | 918 | A | VAL | 1129 | C |
| ILE | 1013 | A | ILE | 1013 | C |
| PRO | 792 | A | VAL | 705 | C |
| GLN | 1002 | A | PHE | 759 | B |
| GLY | 669 | A | MET | 869 | B |
| ALA | 668 | A | PRO | 862 | B |
| PHE | 1121 | A | GLN | 913 | B |
| GLN | 1010 | A | VAL | 1008 | B |
| SER | 1030 | A | ASP | 1041 | C |
| ASP | 198 | A | ASP | 428 | C |
| SER | 704 | A | ILE | 788 | B |
| ASN | 703 | A | GLN | 787 | B |
| ILE | 1013 | A | THR | 1009 | B |
| ALA | 892 | A | GLU | 1072 | C |
| GLU | 702 | A | ILE | 788 | B |
| GLY | 545 | A | ASN | 978 | B |
| PHE | 559 | A | PHE | 43 | B |
| ASN | 907 | A | ARG | 1091 | C |
| GLN | 1002 | A | GLN | 1002 | B |
| PRO | 715 | A | LEU | 894 | B |
| ALA | 890 | A | ASP | 1041 | C |
| ALA | 890 | A | PRO | 1069 | C |
| THR | 1077 | A | PRO | 897 | B |
| ASP | 427 | A | LYS | 986 | C |
| LEU | 546 | A | ASN | 978 | B |
| ASN | 919 | A | VAL | 1128 | C |
| PHE | 562 | A | VAL | 42 | B |
| SER | 975 | A | ASP | 571 | C |
| ASN | 1074 | A | GLN | 895 | B |
| ASN | 1125 | A | GLU | 918 | B |
| THR | 549 | A | ASP | 745 | B |
| LEU | 984 | A | LYS | 386 | C |
| ILE | 788 | A | GLU | 702 | C |
| ASN | 914 | A | PHE | 1121 | C |
| TYR | 38 | A | LEU | 560 | C |
| GLN | 895 | A | VAL | 705 | C |
| PHE | 562 | A | GLU | 224 | B |
| GLY | 545 | A | SER | 982 | B |
| VAL | 1129 | A | GLU | 918 | B |
| MET | 740 | A | PRO | 589 | C |
| LYS | 1038 | A | LYS | 1038 | B |
| LEU | 1141 | A | PRO | 1140 | B |
| LEU | 864 | A | CYS | 671 | C |
| THR | 791 | A | VAL | 705 | C |
| LEU | 1141 | A | GLU | 1144 | B |
| ASN | 709 | A | PRO | 897 | B |
| LEU | 981 | A | THR | 547 | C |
| TYR | 917 | A | PHE | 1089 | C |
| ILE | 896 | A | TYR | 707 | C |
| GLY | 566 | A | ARG | 44 | B |
| VAL | 1128 | A | GLU | 918 | B |
| GLN | 1005 | A | THR | 1006 | C |
| ASP | 985 | A | LYS | 386 | C |
| VAL | 1040 | A | LEU | 1034 | B |
| VAL | 1122 | A | GLN | 1113 | B |
| ILE | 712 | A | TYR | 904 | B |
| VAL | 976 | A | ASP | 571 | C |
| MET | 900 | A | VAL | 1094 | C |
| THR | 302 | A | THR | 761 | B |
| LEU | 861 | A | GLN | 613 | C |
| GLY | 889 | A | LYS | 1045 | C |
| GLN | 779 | A | MET | 697 | C |
| GLY | 1035 | A | VAL | 1040 | C |
| GLN | 414 | A | ASP | 985 | C |
| ARG | 646 | A | PRO | 862 | B |
| VAL | 1040 | A | GLY | 889 | B |
| ASP | 389 | A | SER | 982 | B |
| THR | 866 | A | GLY | 669 | C |
| LEU | 560 | A | GLU | 224 | B |
| LEU | 864 | A | MET | 697 | C |
| GLN | 1036 | A | VAL | 1040 | C |
| SER | 735 | A | GLN | 314 | C |
| ARG | 319 | A | ASP | 745 | B |
| PHE | 1042 | A | SER | 1030 | B |
| ASN | 282 | A | LYS | 558 | C |
| PHE | 1095 | A | MET | 900 | B |
| PRO | 561 | A | GLU | 224 | B |
| VAL | 1040 | A | GLN | 1036 | B |
| PHE | 759 | A | SER | 1003 | C |
| GLN | 965 | A | SER | 758 | B |
| TYR | 917 | A | VAL | 1129 | C |
| SER | 704 | A | LYS | 790 | B |
| LYS | 1045 | A | LYS | 786 | B |
| ASN | 317 | A | ASP | 737 | B |
| ASP | 1041 | A | ALA | 890 | B |
| GLY | 669 | A | PRO | 863 | B |
| GLU | 702 | A | TYR | 789 | B |
| ARG | 357 | A | PRO | 230 | B |
| GLN | 115 | A | ILE | 468 | C |
| PRO | 1090 | A | ASN | 907 | B |
| ASN | 914 | A | PHE | 1089 | C |
| SER | 711 | A | ILE | 896 | B |
| VAL | 705 | A | SER | 884 | B |
| LEU | 864 | A | CYS | 662 | C |
| PHE | 565 | A | LYS | 41 | B |
| PRO | 384 | A | ASP | 985 | B |
| MET | 697 | A | GLN | 779 | B |
| THR | 791 | A | SER | 704 | C |
| GLN | 1002 | A | GLN | 1005 | B |
| MET | 869 | A | LEU | 699 | C |
| GLN | 613 | A | LEU | 861 | B |
| GLY | 971 | A | GLN | 755 | B |
| TRP | 886 | A | ARG | 1107 | C |
| ILE | 666 | A | LEU | 864 | B |
| SER | 711 | A | GLN | 895 | B |
| SER | 1123 | A | GLU | 918 | B |
| PHE | 562 | A | ASP | 40 | B |
| ILE | 788 | A | SER | 704 | C |
| ASN | 969 | A | TYR | 756 | B |
| ALA | 890 | A | TYR | 1067 | C |
| GLU | 1031 | A | VAL | 1040 | C |
| GLN | 1002 | A | LEU | 1001 | B |
| LYS | 786 | A | LEU | 699 | C |
| GLY | 199 | A | PRO | 463 | C |
| ILE | 233 | A | ARG | 466 | C |
| GLN | 1002 | A | GLN | 1002 | C |
| PHE | 43 | A | ARG | 567 | C |
| LYS | 921 | A | ILE | 1130 | C |
| ILE | 896 | A | ALA | 713 | C |
| PRO | 862 | A | ILE | 666 | C |
| GLN | 755 | A | ALA | 972 | C |
| THR | 385 | A | LEU | 984 | B |
| PRO | 863 | A | GLY | 667 | C |
| LEU | 984 | A | SER | 383 | C |
| GLN | 563 | A | LYS | 41 | B |
| ARG | 983 | A | THR | 430 | C |
| LEU | 699 | A | GLN | 872 | B |
| VAL | 42 | A | ARG | 567 | C |
| LYS | 558 | A | PHE | 43 | B |
| ALA | 893 | A | GLU | 1072 | C |
| GLY | 381 | A | ILE | 973 | B |
| PHE | 592 | A | ASN | 856 | B |
| TYR | 396 | A | TYR | 200 | B |
| PRO | 862 | A | ALA | 668 | C |
| PRO | 589 | A | PHE | 855 | B |
| PRO | 897 | A | ASN | 709 | C |
| LEU | 390 | A | SER | 982 | B |
| LYS | 386 | A | SER | 982 | B |

**Table S3.** The list of the inter-protomer contacts of the protomer A in the structure of SARS-CoV-2 S-GSAS/G614 in the closed state (pdb id 7KDK).

| **S protein residue name** | **S protein residue number** | **S protein chain** | **S protein residue name** | **S protein residue number** | **S protein chain** |
| --- | --- | --- | --- | --- | --- |
| PHE | 562 | A | GLU | 224 | B |
| PHE | 562 | A | PRO | 225 | B |
| GLN | 755 | A | PHE | 970 | C |
| GLY | 757 | A | SER | 968 | C |
| ARG | 1039 | A | THR | 1027 | B |
| LEU | 1034 | A | VAL | 1040 | C |
| TYR | 707 | A | ILE | 794 | B |
| GLY | 700 | A | LYS | 786 | B |
| GLY | 798 | A | TYR | 707 | C |
| ALA | 713 | A | GLN | 895 | B |
| TYR | 200 | A | GLU | 516 | C |
| GLN | 895 | A | ASN | 1074 | C |
| GLY | 857 | A | ASN | 317 | C |
| ASN | 856 | A | ALA | 570 | C |
| PHE | 592 | A | THR | 859 | B |
| ASP | 994 | A | PHE | 970 | C |
| VAL | 1008 | A | GLN | 1010 | C |
| GLN | 563 | A | PHE | 43 | B |
| LYS | 1045 | A | GLY | 889 | B |
| ARG | 567 | A | VAL | 976 | B |
| CYS | 379 | A | GLU | 988 | B |
| LEU | 699 | A | ILE | 870 | B |
| ALA | 890 | A | TYR | 1047 | C |
| ILE | 1013 | A | ALA | 1016 | B |
| TYR | 707 | A | PHE | 797 | B |
| PRO | 863 | A | GLY | 667 | C |
| GLU | 702 | A | TYR | 789 | B |
| LEU | 699 | A | VAL | 785 | B |
| ALA | 1080 | A | TYR | 917 | B |
| PHE | 562 | A | VAL | 42 | B |
| SER | 711 | A | ILE | 896 | B |
| TYR | 904 | A | VAL | 1094 | C |
| GLY | 283 | A | LEU | 560 | C |
| GLU | 918 | A | PHE | 1089 | C |
| ARG | 567 | A | ARG | 44 | B |
| ARG | 1039 | A | ARG | 1039 | C |
| GLU | 988 | A | SER | 383 | C |
| LEU | 699 | A | LYS | 786 | B |
| PRO | 384 | A | THR | 415 | C |
| VAL | 987 | A | PRO | 412 | B |
| ILE | 1130 | A | GLY | 798 | B |
| LEU | 1145 | A | GLU | 1144 | B |
| ILE | 896 | A | ILE | 712 | C |
| GLN | 913 | A | PHE | 1089 | C |
| SER | 1003 | A | PHE | 759 | B |
| PRO | 230 | A | ARG | 357 | C |
| GLY | 700 | A | GLN | 787 | B |
| THR | 961 | A | ARG | 765 | B |
| LEU | 864 | A | PRO | 665 | C |
| VAL | 1068 | A | GLY | 891 | B |
| ALA | 892 | A | PRO | 1069 | C |
| SER | 982 | A | GLY | 545 | C |
| PHE | 464 | A | TYR | 200 | B |
| GLN | 1010 | A | GLN | 762 | B |
| PRO | 897 | A | THR | 1077 | C |
| TYR | 789 | A | SER | 704 | C |
| ASP | 1041 | A | GLY | 889 | B |
| TYR | 707 | A | ILE | 882 | B |
| GLY | 545 | A | SER | 982 | B |
| SER | 1147 | A | SER | 1147 | B |
| TYR | 707 | A | ALA | 899 | B |
| ALA | 701 | A | ILE | 788 | B |
| ASP | 979 | A | ARG | 567 | C |
| MET | 697 | A | MET | 869 | B |
| TYR | 789 | A | VAL | 705 | C |
| ILE | 468 | A | GLN | 115 | B |
| GLY | 669 | A | THR | 866 | B |
| MET | 697 | A | LEU | 865 | B |
| VAL | 382 | A | LEU | 984 | B |
| GLN | 115 | A | ILE | 468 | C |
| GLN | 1010 | A | ALA | 766 | B |
| GLY | 669 | A | MET | 869 | B |
| GLY | 971 | A | GLN | 755 | B |
| LYS | 795 | A | TYR | 707 | C |
| GLY | 232 | A | GLU | 465 | C |
| THR | 859 | A | PHE | 592 | C |
| GLN | 1002 | A | LEU | 1001 | B |
| ARG | 983 | A | LEU | 390 | C |
| ALA | 713 | A | LEU | 894 | B |
| ASP | 1041 | A | GLN | 784 | B |
| ASP | 985 | A | THR | 415 | B |
| GLY | 889 | A | ASP | 1041 | C |
| PRO | 589 | A | PHE | 855 | B |
| PRO | 863 | A | ALA | 668 | C |
| GLN | 1005 | A | THR | 1006 | C |
| LEU | 560 | A | THR | 284 | B |
| LEU | 861 | A | GLN | 613 | C |
| ALA | 890 | A | PRO | 1069 | C |
| ARG | 567 | A | PHE | 43 | B |
| SER | 383 | A | LEU | 984 | B |
| GLU | 918 | A | SER | 1123 | C |
| THR | 415 | A | TYR | 369 | B |
| VAL | 705 | A | SER | 884 | B |
| GLN | 1002 | A | GLN | 1002 | C |
| ASN | 914 | A | PHE | 1121 | C |
| LEU | 1145 | A | LEU | 1145 | B |
| PHE | 592 | A | LEU | 858 | B |
| ALA | 766 | A | GLN | 1010 | C |
| MET | 869 | A | ALA | 668 | C |
| ASP | 198 | A | PHE | 464 | C |
| SER | 968 | A | GLY | 757 | B |
| LYS | 786 | A | LYS | 1045 | C |
| ASP | 985 | A | LYS | 386 | C |
| MET | 869 | A | GLY | 669 | C |
| TYR | 917 | A | ILE | 1130 | C |
| TYR | 789 | A | ASN | 703 | C |
| THR | 572 | A | ASN | 856 | B |
| ASP | 198 | A | ASP | 428 | C |
| MET | 869 | A | THR | 696 | C |
| SER | 704 | A | ILE | 788 | B |
| LYS | 776 | A | LYS | 947 | C |
| ILE | 788 | A | ASN | 703 | C |
| GLN | 895 | A | VAL | 705 | C |
| ARG | 646 | A | THR | 866 | B |
| ASP | 568 | A | PHE | 855 | B |
| LEU | 864 | A | CYS | 662 | C |
| GLN | 895 | A | ALA | 706 | C |
| ARG | 44 | A | GLN | 563 | C |
| GLN | 895 | A | SER | 708 | C |
| ILE | 569 | A | ASN | 960 | B |
| ASP | 428 | A | ASP | 198 | B |
| GLY | 1035 | A | VAL | 1040 | C |
| ALA | 570 | A | SER | 975 | B |
| SER | 711 | A | GLN | 895 | B |
| LEU | 984 | A | VAL | 382 | C |
| GLY | 757 | A | GLN | 965 | C |
| ARG | 319 | A | ASP | 745 | B |
| GLN | 564 | A | LYS | 41 | B |
| ASP | 571 | A | SER | 967 | B |
| LEU | 966 | A | ASP | 571 | C |
| GLU | 1031 | A | PHE | 1042 | C |
| ARG | 466 | A | GLN | 115 | B |
| VAL | 705 | A | PRO | 792 | B |
| PHE | 855 | A | ASP | 568 | C |
| ALA | 570 | A | ASN | 856 | B |
| THR | 430 | A | ARG | 983 | B |
| ASP | 796 | A | TYR | 707 | C |
| PRO | 230 | A | TYR | 396 | C |
| ASP | 571 | A | ARG | 44 | B |
| ILE | 231 | A | ARG | 466 | C |
| ALA | 668 | A | PRO | 862 | B |
| ASN | 703 | A | GLN | 787 | B |
| ARG | 1107 | A | ASN | 907 | B |
| VAL | 1129 | A | TYR | 917 | B |
| LYS | 41 | A | ALA | 520 | C |
| GLN | 1002 | A | TYR | 756 | B |
| GLN | 920 | A | ILE | 1130 | C |
| LEU | 560 | A | GLY | 283 | B |
| SER | 982 | A | THR | 547 | C |
| GLU | 516 | A | ASP | 228 | B |
| GLN | 563 | A | TYR | 38 | B |
| TYR | 917 | A | VAL | 1128 | C |
| SER | 704 | A | LYS | 790 | B |
| ALA | 701 | A | GLN | 787 | B |
| ASN | 764 | A | GLN | 314 | C |
| ASN | 856 | A | THR | 572 | C |
| MET | 869 | A | LEU | 699 | C |
| ARG | 983 | A | LYS | 386 | C |
| MET | 869 | A | ILE | 670 | C |
| GLN | 755 | A | ALA | 972 | C |
| GLU | 224 | A | PRO | 561 | C |
| LYS | 790 | A | ASN | 703 | C |
| TYR | 396 | A | PRO | 230 | B |
| ILE | 712 | A | ILE | 896 | B |
| LEU | 1012 | A | ILE | 1013 | C |
| ILE | 788 | A | LEU | 699 | C |
| GLY | 1093 | A | GLN | 913 | B |
| PHE | 43 | A | PHE | 565 | C |
| ARG | 983 | A | GLY | 381 | C |
| GLU | 1111 | A | SER | 1123 | C |
| TRP | 886 | A | ASN | 1108 | C |
| ASN | 914 | A | PHE | 1089 | C |
| SER | 698 | A | TYR | 873 | B |
| ALA | 713 | A | ILE | 896 | B |
| ARG | 319 | A | MET | 740 | B |
| TYR | 1047 | A | THR | 887 | B |
| ASP | 571 | A | VAL | 976 | B |
| THR | 1077 | A | MET | 900 | B |
| GLN | 314 | A | ASN | 764 | B |
| HIS | 519 | A | ARG | 983 | B |
| GLY | 199 | A | GLU | 465 | C |
| GLY | 667 | A | PRO | 863 | B |
| LEU | 1141 | A | LEU | 1141 | B |
| SER | 708 | A | PRO | 897 | B |
| ILE | 714 | A | LEU | 894 | B |
| LEU | 699 | A | TYR | 873 | B |
| PRO | 862 | A | GLY | 667 | C |
| TYR | 756 | A | GLN | 1002 | C |
| GLY | 667 | A | LEU | 864 | B |
| LEU | 865 | A | LEU | 699 | C |
| GLY | 669 | A | PRO | 863 | B |
| TYR | 789 | A | GLU | 702 | C |
| VAL | 42 | A | GLN | 563 | C |
| GLN | 913 | A | PRO | 1079 | C |
| LYS | 41 | A | PHE | 565 | C |
| ALA | 1078 | A | TYR | 917 | B |
| LYS | 386 | A | ARG | 983 | B |
| GLY | 548 | A | ASN | 978 | B |
| LYS | 378 | A | GLU | 988 | B |
| MET | 697 | A | TYR | 873 | B |
| PHE | 1089 | A | GLU | 918 | B |
| LEU | 966 | A | ALA | 570 | C |
| ALA | 890 | A | ASP | 1041 | C |
| SER | 1123 | A | GLU | 1111 | B |
| LEU | 560 | A | TYR | 38 | B |
| LYS | 202 | A | LEU | 518 | C |
| ARG | 983 | A | VAL | 382 | C |
| PRO | 897 | A | ILE | 712 | C |
| THR | 547 | A | ASN | 978 | B |
| GLU | 516 | A | TYR | 200 | B |
| SER | 968 | A | TYR | 756 | B |
| VAL | 1094 | A | MET | 900 | B |
| SER | 975 | A | ARG | 567 | C |
| TYR | 917 | A | VAL | 1129 | C |
| GLN | 895 | A | ALA | 713 | C |
| TYR | 369 | A | ARG | 408 | C |
| ASP | 745 | A | ARG | 319 | C |
| ARG | 466 | A | ILE | 233 | B |
| PHE | 562 | A | LYS | 41 | B |
| GLU | 988 | A | CYS | 379 | C |
| GLU | 1072 | A | ALA | 893 | B |
| SER | 704 | A | TYR | 789 | B |
| ASN | 234 | A | ARG | 466 | C |
| SER | 1123 | A | GLU | 918 | B |
| GLN | 1010 | A | LEU | 1012 | B |
| LYS | 558 | A | PHE | 43 | B |
| GLN | 563 | A | ARG | 44 | B |
| LEU | 864 | A | ILE | 670 | C |
| ARG | 319 | A | ASP | 737 | B |
| VAL | 1040 | A | GLN | 1036 | B |
| MET | 900 | A | VAL | 1094 | C |
| PHE | 592 | A | MET | 740 | B |
| ARG | 355 | A | TYR | 200 | B |
| LYS | 790 | A | GLU | 702 | C |
| VAL | 42 | A | PHE | 565 | C |
| ALA | 570 | A | LYS | 964 | B |
| LEU | 894 | A | ILE | 714 | C |
| PHE | 592 | A | LYS | 854 | B |
| ARG | 1107 | A | TRP | 886 | B |
| VAL | 963 | A | ALA | 570 | C |
| GLN | 762 | A | TYR | 1007 | C |
| TYR | 707 | A | GLN | 895 | B |
| GLN | 787 | A | GLY | 700 | C |
| ASN | 317 | A | MET | 740 | B |
| TYR | 707 | A | GLY | 798 | B |
| TYR | 1007 | A | GLN | 762 | B |
| ILE | 569 | A | VAL | 47 | B |
| LYS | 1038 | A | GLN | 1036 | B |
| THR | 961 | A | SER | 758 | B |
| PRO | 1090 | A | GLN | 913 | B |
| ALA | 893 | A | VAL | 705 | C |
| TYR | 756 | A | PHE | 970 | C |
| GLY | 857 | A | PHE | 592 | C |
| ALA | 1016 | A | ILE | 1013 | C |
| PHE | 970 | A | PHE | 759 | B |
| ASN | 978 | A | GLY | 548 | C |
| PHE | 464 | A | GLY | 199 | B |
| ILE | 788 | A | SER | 704 | C |
| GLN | 965 | A | PHE | 759 | B |
| VAL | 705 | A | ALA | 879 | B |
| ASN | 709 | A | PRO | 897 | B |
| THR | 385 | A | ASP | 985 | B |
| VAL | 382 | A | GLU | 988 | B |
| PRO | 521 | A | LYS | 41 | B |
| ASN | 914 | A | SER | 1123 | C |
| THR | 866 | A | ALA | 668 | C |
| PRO | 1069 | A | LEU | 894 | B |
| ILE | 788 | A | ALA | 701 | C |
| VAL | 963 | A | ILE | 569 | C |
| LEU | 390 | A | SER | 982 | B |
| TYR | 873 | A | MET | 697 | C |
| CYS | 662 | A | LEU | 864 | B |
| ASN | 1074 | A | GLN | 895 | B |
| THR | 415 | A | THR | 385 | B |
| ALA | 890 | A | LYS | 1045 | C |
| SER | 383 | A | GLU | 988 | B |
| ALA | 520 | A | LYS | 41 | B |
| GLN | 1002 | A | GLN | 1005 | B |
| GLN | 755 | A | GLY | 971 | C |
| ASP | 1041 | A | LEU | 1034 | B |
| LYS | 921 | A | ILE | 1130 | C |
| GLU | 702 | A | ILE | 788 | B |
| LEU | 699 | A | ILE | 788 | B |
| PHE | 759 | A | THR | 1006 | C |
| ILE | 1130 | A | TYR | 917 | B |
| ASP | 427 | A | LYS | 986 | C |
| SER | 975 | A | ASP | 571 | C |
| TRP | 886 | A | ARG | 1107 | C |
| PHE | 759 | A | GLN | 1002 | C |
| ASN | 317 | A | ASP | 737 | B |
| GLY | 891 | A | LYS | 1045 | C |
| LEU | 1001 | A | GLN | 1002 | C |
| ALA | 890 | A | VAL | 1068 | C |
| VAL | 1094 | A | ILE | 896 | B |
| LEU | 546 | A | ASN | 978 | B |
| TYR | 38 | A | LEU | 560 | C |
| ARG | 983 | A | SER | 383 | C |
| VAL | 1128 | A | ASN | 919 | B |
| TYR | 873 | A | LEU | 699 | C |
| GLN | 957 | A | ARG | 765 | B |
| ARG | 1019 | A | GLU | 1017 | C |
| ALA | 372 | A | ARG | 403 | C |
| GLY | 880 | A | VAL | 705 | C |
| GLU | 988 | A | PRO | 384 | C |
| ALA | 1016 | A | ARG | 1019 | B |
| TYR | 756 | A | GLN | 965 | C |
| GLN | 762 | A | GLN | 965 | C |
| TYR | 707 | A | PRO | 897 | B |
| MET | 900 | A | PRO | 1079 | C |
| LYS | 790 | A | VAL | 705 | C |
| GLN | 613 | A | LEU | 861 | B |
| ILE | 896 | A | TYR | 707 | C |
| PHE | 565 | A | PHE | 43 | B |
| PHE | 592 | A | PHE | 855 | B |
| GLY | 798 | A | ILE | 1130 | C |
| PRO | 863 | A | GLY | 669 | C |
| ASN | 1108 | A | TRP | 886 | B |
| LYS | 790 | A | SER | 704 | C |
| ILE | 666 | A | LEU | 864 | B |
| LEU | 1141 | A | LEU | 1141 | C |
| VAL | 1129 | A | GLU | 918 | B |
| LYS | 41 | A | PHE | 562 | C |
| THR | 1006 | A | GLN | 762 | B |
| ASP | 571 | A | SER | 975 | B |
| ILE | 896 | A | SER | 711 | C |
| ASP | 427 | A | VAL | 987 | C |
| TYR | 1067 | A | ALA | 890 | B |
| THR | 883 | A | TYR | 707 | C |
| PRO | 862 | A | ALA | 647 | C |
| ILE | 1013 | A | THR | 1009 | B |
| VAL | 1040 | A | LEU | 1034 | B |
| ASN | 907 | A | ARG | 1107 | C |
| PRO | 715 | A | LEU | 894 | B |
| GLN | 1010 | A | VAL | 1008 | B |
| ILE | 870 | A | LEU | 699 | C |
| TYR | 200 | A | ARG | 355 | C |
| TYR | 873 | A | GLY | 700 | C |
| GLN | 563 | A | TYR | 279 | B |
| SER | 1030 | A | ASP | 1041 | C |
| ALA | 713 | A | ALA | 893 | B |
| PRO | 1079 | A | MET | 900 | B |
| SER | 968 | A | GLN | 755 | B |
| ILE | 712 | A | PRO | 897 | B |
| ASP | 985 | A | THR | 385 | C |
| GLU | 1072 | A | ALA | 892 | B |
| GLN | 755 | A | ASN | 969 | C |
| LEU | 984 | A | LYS | 386 | C |
| GLY | 232 | A | ARG | 466 | C |
| GLY | 566 | A | ARG | 44 | B |
| PHE | 797 | A | TYR | 707 | C |
| LEU | 894 | A | PRO | 1069 | C |
| LEU | 699 | A | MET | 869 | B |
| PRO | 897 | A | SER | 708 | C |
| LEU | 894 | A | ARG | 1107 | C |
| VAL | 1128 | A | TYR | 917 | B |
| PHE | 559 | A | PHE | 43 | B |
| VAL | 1040 | A | GLU | 1031 | B |
| SER | 758 | A | SER | 968 | C |
| ILE | 1013 | A | ILE | 1013 | C |
| PRO | 463 | A | GLY | 199 | B |
| THR | 1077 | A | PRO | 897 | B |
| ARG | 646 | A | PRO | 863 | B |
| GLN | 913 | A | GLY | 1093 | C |
| GLN | 613 | A | THR | 859 | B |
| ARG | 1091 | A | ASN | 907 | B |
| PRO | 230 | A | ARG | 355 | C |
| ILE | 569 | A | LYS | 964 | B |
| PRO | 589 | A | MET | 740 | B |
| MET | 900 | A | PHE | 1095 | C |
| ALA | 570 | A | LEU | 966 | B |
| PHE | 759 | A | GLY | 999 | C |
| LEU | 517 | A | ARG | 983 | B |
| GLN | 1002 | A | GLN | 1002 | B |
| GLY | 889 | A | VAL | 1040 | C |
| ARG | 1039 | A | GLU | 1031 | B |
| VAL | 1068 | A | ALA | 890 | B |
| GLN | 965 | A | GLN | 762 | B |
| LEU | 865 | A | GLY | 669 | C |
| GLY | 413 | A | VAL | 987 | C |
| GLY | 700 | A | TYR | 873 | B |
| MET | 740 | A | ASN | 317 | C |
| GLN | 314 | A | THR | 768 | B |
| GLY | 891 | A | GLY | 1046 | C |
| LEU | 699 | A | LEU | 865 | B |
| THR | 547 | A | SER | 982 | B |
| GLU | 1017 | A | ARG | 1019 | B |
| TYR | 1047 | A | ALA | 890 | B |
| GLN | 965 | A | SER | 758 | B |
| VAL | 705 | A | LEU | 894 | B |
| GLY | 700 | A | ILE | 788 | B |
| ALA | 701 | A | TYR | 789 | B |
| LYS | 964 | A | ILE | 569 | C |
| PHE | 1121 | A | THR | 912 | B |
| LEU | 865 | A | ALA | 668 | C |
| SER | 982 | A | LYS | 386 | C |
| SER | 982 | A | LEU | 390 | C |
| ALA | 893 | A | ALA | 713 | C |
| PRO | 862 | A | ARG | 646 | C |
| ASN | 282 | A | LYS | 558 | C |
| PHE | 1089 | A | TYR | 917 | B |
| LEU | 864 | A | ALA | 668 | C |
| GLN | 1036 | A | LYS | 1038 | C |
| ARG | 567 | A | VAL | 42 | B |
| MET | 697 | A | GLN | 779 | B |
| ILE | 670 | A | LEU | 864 | B |
| SER | 735 | A | GLN | 314 | C |
| PHE | 970 | A | ASP | 994 | B |
| LYS | 558 | A | ASN | 282 | B |
| GLN | 895 | A | ILE | 712 | C |
| VAL | 42 | A | HIS | 519 | C |
| ILE | 233 | A | ARG | 466 | C |
| ARG | 408 | A | TYR | 369 | B |
| GLN | 762 | A | THR | 1006 | C |
| ILE | 666 | A | PRO | 862 | B |
| PRO | 463 | A | ASP | 198 | B |
| HIS | 519 | A | ASP | 979 | B |
| GLY | 999 | A | PHE | 759 | B |
| VAL | 1040 | A | GLY | 1035 | B |
| MET | 869 | A | MET | 697 | C |
| PRO | 862 | A | ALA | 668 | C |
| LYS | 986 | A | ASP | 427 | B |
| ASP | 796 | A | ASN | 709 | C |
| PHE | 43 | A | PHE | 559 | C |
| ILE | 712 | A | GLN | 895 | B |
| VAL | 47 | A | ILE | 569 | C |
| SER | 758 | A | LYS | 964 | C |
| VAL | 1094 | A | TYR | 904 | B |
| ARG | 357 | A | PRO | 230 | B |
| ARG | 1107 | A | LEU | 894 | B |
| PHE | 43 | A | GLY | 566 | C |
| ALA | 647 | A | PRO | 862 | B |
| LEU | 858 | A | PHE | 592 | C |
| THR | 1009 | A | THR | 1009 | C |
| GLU | 702 | A | LYS | 790 | B |
| ASN | 978 | A | LEU | 546 | C |
| PHE | 898 | A | TYR | 707 | C |
| TYR | 38 | A | GLN | 563 | C |
| PRO | 412 | A | VAL | 987 | C |
| GLU | 465 | A | ILE | 233 | B |
| SER | 514 | A | TYR | 200 | B |
| GLN | 784 | A | ASP | 1041 | C |
| LEU | 1141 | A | PRO | 1140 | B |
| GLY | 669 | A | LEU | 864 | B |
| ARG | 1107 | A | TYR | 904 | B |
| VAL | 705 | A | TYR | 789 | B |
| LEU | 1145 | A | LEU | 1141 | B |
| THR | 883 | A | VAL | 705 | C |
| ALA | 879 | A | VAL | 705 | C |
| PRO | 897 | A | ASN | 709 | C |
| THR | 696 | A | MET | 869 | B |
| LEU | 560 | A | PHE | 43 | B |
| GLN | 563 | A | ASP | 40 | B |
| ASP | 796 | A | SER | 708 | C |
| LYS | 41 | A | GLN | 564 | C |
| PRO | 665 | A | LEU | 864 | B |
| ASP | 228 | A | GLU | 516 | C |
| PRO | 384 | A | GLU | 988 | B |
| GLU | 1017 | A | GLU | 773 | B |
| ASP | 979 | A | LEU | 546 | C |
| VAL | 42 | A | PHE | 562 | C |
| SER | 758 | A | GLN | 965 | C |
| LYS | 386 | A | SER | 982 | B |
| ALA | 668 | A | MET | 869 | B |
| VAL | 382 | A | ARG | 983 | B |
| GLU | 1072 | A | LEU | 894 | B |
| TYR | 396 | A | TYR | 200 | B |
| ASP | 40 | A | HIS | 519 | C |
| THR | 1009 | A | THR | 1009 | B |
| LEU | 865 | A | MET | 697 | C |
| TYR | 369 | A | GLY | 416 | C |
| PHE | 970 | A | GLN | 755 | B |
| GLU | 1031 | A | VAL | 1040 | C |
| CYS | 671 | A | LEU | 864 | B |
| TYR | 917 | A | PHE | 1089 | C |
| VAL | 976 | A | THR | 572 | C |
| ARG | 1019 | A | ILE | 1013 | C |
| PHE | 970 | A | GLY | 757 | B |
| ARG | 646 | A | PRO | 862 | B |
| ILE | 788 | A | GLY | 700 | C |
| PRO | 1069 | A | ALA | 892 | B |
| ILE | 712 | A | TYR | 904 | B |
| GLN | 779 | A | MET | 697 | C |
| PHE | 592 | A | ASN | 856 | B |
| ARG | 319 | A | THR | 739 | B |
| GLN | 563 | A | GLY | 283 | B |
| VAL | 976 | A | ASP | 571 | C |
| PRO | 426 | A | ASP | 198 | B |
| THR | 385 | A | THR | 415 | C |
| ALA | 890 | A | GLY | 1046 | C |
| SER | 708 | A | GLN | 895 | B |
| ASN | 703 | A | TYR | 789 | B |
| GLY | 667 | A | PRO | 862 | B |
| GLY | 1124 | A | GLU | 918 | B |
| GLY | 669 | A | LEU | 865 | B |
| ALA | 892 | A | GLU | 1072 | C |
| PHE | 759 | A | GLN | 965 | C |
| TYR | 904 | A | ARG | 1107 | C |
| VAL | 976 | A | ARG | 567 | C |
| ARG | 466 | A | THR | 167 | B |
| PHE | 43 | A | LYS | 558 | C |
| THR | 1006 | A | PHE | 759 | B |
| GLY | 566 | A | PHE | 43 | B |
| THR | 1027 | A | ARG | 1039 | C |
| GLN | 1036 | A | VAL | 1040 | C |
| ILE | 896 | A | SER | 708 | C |
| VAL | 705 | A | LYS | 790 | B |
| LYS | 557 | A | PHE | 43 | B |
| THR | 887 | A | ASN | 1108 | C |
| SER | 967 | A | ASP | 571 | C |
| SER | 968 | A | SER | 758 | B |
| GLU | 988 | A | VAL | 382 | C |
| TYR | 789 | A | ALA | 701 | C |
| LEU | 984 | A | SER | 383 | C |
| GLY | 891 | A | PRO | 1069 | C |
| LEU | 894 | A | PRO | 715 | C |
| LYS | 964 | A | SER | 758 | B |
| LEU | 864 | A | MET | 697 | C |
| ILE | 569 | A | SER | 967 | B |
| ASP | 198 | A | PRO | 463 | C |
| SER | 967 | A | ALA | 570 | C |
| PHE | 464 | A | ASP | 198 | B |
| ILE | 1130 | A | LYS | 921 | B |
| ARG | 1107 | A | GLN | 913 | B |
| PHE | 592 | A | GLY | 857 | B |
| TYR | 707 | A | PHE | 898 | B |
| GLN | 895 | A | TYR | 707 | C |
| VAL | 705 | A | GLN | 895 | B |
| GLY | 1046 | A | ALA | 890 | B |
| LEU | 390 | A | ARG | 983 | B |
| PHE | 1042 | A | GLU | 1031 | B |
| GLU | 918 | A | VAL | 1128 | C |
| VAL | 705 | A | THR | 883 | B |
| GLY | 889 | A | GLY | 1046 | C |
| THR | 912 | A | PHE | 1121 | C |
| VAL | 987 | A | ASP | 427 | B |
| MET | 900 | A | THR | 1077 | C |
| LEU | 894 | A | ASN | 1108 | C |
| PRO | 897 | A | SER | 711 | C |
| ASP | 40 | A | GLN | 563 | C |
| VAL | 1128 | A | GLU | 918 | B |
| GLN | 613 | A | SER | 735 | B |
| ILE | 1013 | A | ILE | 1013 | B |
| GLN | 563 | A | VAL | 42 | B |
| HIS | 519 | A | VAL | 42 | B |
| VAL | 1040 | A | GLY | 889 | B |
| THR | 1006 | A | GLN | 1005 | B |
| ILE | 896 | A | VAL | 1094 | C |
| TYR | 279 | A | GLN | 563 | C |
| GLN | 755 | A | SER | 968 | C |
| LYS | 557 | A | SER | 45 | B |
| GLN | 563 | A | LYS | 41 | B |
| ILE | 1013 | A | ARG | 1019 | B |
| ALA | 890 | A | TYR | 1067 | C |
| SER | 758 | A | THR | 961 | C |
| ASP | 198 | A | PRO | 426 | C |
| PRO | 792 | A | VAL | 705 | C |
| LEU | 1145 | A | SER | 1147 | B |
| TYR | 707 | A | PRO | 792 | B |
| GLN | 913 | A | PRO | 1090 | C |
| TYR | 1047 | A | TRP | 886 | B |
| GLY | 1093 | A | TYR | 904 | B |
| GLN | 314 | A | SER | 735 | B |
| GLY | 1046 | A | GLY | 891 | B |
| PHE | 970 | A | TYR | 756 | B |
| PHE | 43 | A | ARG | 567 | C |
| PRO | 792 | A | TYR | 707 | C |
| ARG | 1039 | A | ARG | 1039 | B |
| LYS | 386 | A | LEU | 981 | B |
| GLN | 872 | A | LEU | 699 | C |
| ASN | 856 | A | ASP | 568 | C |
| GLN | 1002 | A | THR | 998 | B |
| ASN | 710 | A | PRO | 897 | B |
| ASP | 1041 | A | ALA | 890 | B |
| ARG | 466 | A | GLY | 232 | B |
| GLN | 314 | A | LEU | 861 | B |
| PRO | 1079 | A | TYR | 917 | B |
| GLY | 381 | A | ARG | 983 | B |
| TYR | 917 | A | ALA | 1080 | C |
| LEU | 864 | A | GLY | 669 | C |
| ARG | 983 | A | THR | 430 | C |
| SER | 383 | A | ARG | 983 | B |
| GLY | 1046 | A | GLY | 889 | B |
| LEU | 864 | A | ILE | 666 | C |
| ASP | 1146 | A | SER | 1147 | B |
| LEU | 864 | A | GLY | 667 | C |
| ILE | 882 | A | TYR | 707 | C |
| THR | 739 | A | ARG | 319 | C |
| LEU | 1012 | A | GLN | 1010 | C |
| GLN | 787 | A | ASN | 703 | C |
| LEU | 1141 | A | GLU | 1144 | B |
| ASN | 978 | A | THR | 547 | C |
| GLY | 381 | A | LEU | 984 | B |
| LYS | 41 | A | GLN | 563 | C |
| THR | 866 | A | GLY | 669 | C |
| MET | 900 | A | ALA | 1078 | C |
| GLN | 965 | A | TYR | 756 | B |
| PHE | 565 | A | LYS | 41 | B |
| ALA | 668 | A | THR | 866 | B |
| LYS | 41 | A | HIS | 519 | C |
| TYR | 170 | A | ARG | 357 | C |
| PRO | 225 | A | PHE | 562 | C |
| ALA | 668 | A | PRO | 863 | B |
| ARG | 983 | A | LEU | 517 | C |
| ASN | 907 | A | ARG | 1091 | C |
| ARG | 44 | A | GLY | 566 | C |
| ILE | 794 | A | TYR | 707 | C |
| SER | 1123 | A | ASN | 914 | B |
| TYR | 200 | A | PHE | 464 | C |
| ARG | 403 | A | ALA | 372 | B |
| GLU | 918 | A | GLY | 1124 | C |
| PHE | 43 | A | LYS | 557 | C |
| SER | 884 | A | VAL | 705 | C |
| PHE | 464 | A | ILE | 233 | B |
| ASN | 703 | A | ILE | 788 | B |
| LEU | 894 | A | GLU | 1072 | C |
| PHE | 1121 | A | ASN | 914 | B |
| GLN | 762 | A | THR | 961 | C |
| VAL | 705 | A | GLY | 880 | B |
| LYS | 854 | A | PHE | 592 | C |
| ILE | 712 | A | MET | 900 | B |
| PHE | 1089 | A | GLN | 913 | B |
| GLU | 702 | A | GLN | 787 | B |
| LYS | 1045 | A | LEU | 1034 | B |
| THR | 887 | A | TYR | 1047 | C |
| TYR | 200 | A | SER | 514 | C |
| TYR | 369 | A | THR | 415 | C |
| GLU | 224 | A | PHE | 562 | C |
| THR | 1009 | A | ILE | 1013 | C |
| ASN | 969 | A | GLN | 755 | B |
| PHE | 464 | A | GLY | 232 | B |
| VAL | 1040 | A | SER | 1030 | B |
| SER | 1030 | A | VAL | 1040 | C |
| GLY | 891 | A | VAL | 1068 | C |
| ALA | 570 | A | SER | 967 | B |
| THR | 961 | A | GLN | 762 | B |
| GLN | 913 | A | PHE | 1121 | C |
| PHE | 855 | A | PRO | 589 | C |
| GLU | 1031 | A | ARG | 1039 | C |
| LEU | 894 | A | VAL | 705 | C |
| LEU | 894 | A | ALA | 713 | C |
| ASN | 317 | A | ASP | 745 | B |
| SER | 708 | A | ASP | 796 | B |
| LYS | 964 | A | ALA | 570 | C |
| ARG | 765 | A | GLN | 957 | C |
| ILE | 569 | A | VAL | 963 | B |
| GLU | 465 | A | GLY | 232 | B |
| PHE | 1121 | A | GLN | 913 | B |
| MET | 697 | A | LEU | 864 | B |
| PHE | 855 | A | PHE | 592 | C |
| GLN | 1005 | A | GLN | 1002 | C |
| ASN | 703 | A | LYS | 790 | B |
| ASP | 571 | A | HIS | 49 | B |
| VAL | 42 | A | ARG | 567 | C |
| ARG | 765 | A | THR | 961 | C |
| ARG | 357 | A | TYR | 170 | B |
| LYS | 1045 | A | GLN | 784 | B |
| GLY | 232 | A | PHE | 464 | C |
| ILE | 1013 | A | LEU | 1012 | B |
| LEU | 984 | A | GLY | 381 | C |
| PHE | 759 | A | SER | 1003 | C |
| GLU | 988 | A | LYS | 378 | C |
| THR | 415 | A | ASP | 985 | C |
| PRO | 1069 | A | GLY | 891 | B |
| GLU | 773 | A | GLU | 1017 | C |
| PHE | 562 | A | ASP | 40 | B |
| LEU | 1034 | A | ASP | 1041 | C |
| SER | 708 | A | ILE | 896 | B |
| ALA | 570 | A | VAL | 963 | B |
| VAL | 987 | A | GLY | 413 | B |
| LYS | 786 | A | ALA | 701 | C |
| PHE | 565 | A | VAL | 42 | B |
| VAL | 42 | A | GLY | 566 | C |
| PRO | 1079 | A | GLN | 913 | B |
| MET | 740 | A | PHE | 592 | C |
| ASN | 919 | A | VAL | 1128 | C |
| PRO | 897 | A | ASN | 710 | C |
| TYR | 756 | A | SER | 968 | C |
| PRO | 1069 | A | ALA | 890 | B |
| TYR | 707 | A | ASP | 796 | B |
| ALA | 1078 | A | MET | 900 | B |
| GLY | 757 | A | PHE | 970 | C |
| ASN | 709 | A | ASP | 796 | B |
| VAL | 785 | A | LEU | 699 | C |
| ASN | 370 | A | LYS | 417 | C |
| GLN | 895 | A | SER | 711 | C |
| ASP | 228 | A | LEU | 518 | C |
| PHE | 43 | A | GLN | 563 | C |
| GLU | 1144 | A | LEU | 1141 | C |
| TYR | 200 | A | TYR | 396 | C |
| LEU | 560 | A | ASN | 282 | B |
| GLU | 918 | A | VAL | 1129 | C |
| LYS | 417 | A | ASN | 370 | B |
| TYR | 505 | A | ALA | 372 | B |
| GLN | 965 | A | GLY | 757 | B |
| VAL | 705 | A | ALA | 893 | B |
| THR | 167 | A | ARG | 466 | C |
| ASP | 737 | A | ARG | 319 | C |
| GLU | 224 | A | LEU | 560 | C |
| ALA | 706 | A | GLN | 895 | B |
| TYR | 707 | A | ILE | 896 | B |
| GLY | 889 | A | LYS | 1045 | C |
| LEU | 546 | A | ASP | 979 | B |
| PHE | 1089 | A | ASN | 914 | B |
| GLY | 283 | A | GLN | 563 | C |
| ASP | 568 | A | ASN | 856 | B |
| ASN | 856 | A | PHE | 592 | C |
| LYS | 786 | A | GLY | 700 | C |
| LYS | 386 | A | LEU | 984 | B |
| ILE | 1130 | A | GLN | 920 | B |
| SER | 711 | A | PRO | 897 | B |
| ARG | 355 | A | PRO | 230 | B |
| ASP | 737 | A | ASN | 317 | C |
| GLN | 787 | A | ALA | 701 | C |
| GLY | 381 | A | ILE | 973 | B |
| ALA | 899 | A | TYR | 707 | C |
| TRP | 886 | A | TYR | 1047 | C |
| GLU | 1144 | A | LEU | 1145 | C |
| PRO | 897 | A | TYR | 707 | C |
| LEU | 981 | A | LYS | 386 | C |
| GLY | 566 | A | VAL | 42 | B |
| ALA | 972 | A | GLN | 755 | B |
| ASP | 40 | A | PHE | 562 | C |
| ILE | 468 | A | GLU | 132 | B |
| ALA | 701 | A | LYS | 786 | B |
| ASP | 985 | A | SER | 383 | C |
| TYR | 873 | A | SER | 698 | C |
| TYR | 707 | A | THR | 883 | B |
| LEU | 699 | A | GLN | 872 | B |
| GLY | 199 | A | PRO | 463 | C |
| LEU | 546 | A | VAL | 976 | B |
| SER | 383 | A | ASP | 985 | B |
| ILE | 788 | A | GLU | 702 | C |
| THR | 768 | A | GLN | 314 | C |
| THR | 572 | A | VAL | 976 | B |
| SER | 45 | A | LYS | 557 | C |
| ALA | 668 | A | LEU | 864 | B |
| GLN | 787 | A | GLU | 702 | C |
| TYR | 917 | A | PRO | 1079 | C |
| ASP | 389 | A | SER | 982 | B |
| ASN | 960 | A | ILE | 569 | C |
| ARG | 466 | A | ILE | 231 | B |
| HIS | 519 | A | LYS | 41 | B |
| MET | 740 | A | PRO | 589 | C |
| MET | 740 | A | ARG | 319 | C |
| ASP | 1041 | A | SER | 1030 | B |
| GLY | 199 | A | PHE | 464 | C |
| MET | 900 | A | ILE | 712 | C |
| THR | 998 | A | GLN | 1002 | C |
| TRP | 353 | A | GLY | 232 | B |
| PHE | 759 | A | PHE | 970 | C |
| GLU | 1017 | A | LYS | 776 | B |
| PHE | 43 | A | LEU | 560 | C |
| ILE | 896 | A | ALA | 713 | C |
| LYS | 1045 | A | ALA | 890 | B |
| PHE | 562 | A | TYR | 38 | B |
| GLN | 1002 | A | PHE | 759 | B |
| ILE | 973 | A | GLY | 381 | C |
| TYR | 904 | A | ILE | 712 | C |
| GLN | 762 | A | GLN | 1010 | C |
| TYR | 38 | A | PHE | 562 | C |
| SER | 967 | A | ILE | 569 | C |
| TYR | 904 | A | GLY | 1093 | C |
| ILE | 233 | A | PHE | 464 | C |
| LYS | 386 | A | ASP | 985 | B |
| LEU | 864 | A | CYS | 671 | C |
| LYS | 786 | A | LEU | 699 | C |
| ARG | 44 | A | ARG | 567 | C |
| PHE | 1095 | A | MET | 900 | B |

**Table S4.** The list of the inter-protomer contacts of the protomer A in the structure of SARS-CoV-2 S-GSAS/G614 in the open state (pdb id 7KDL).

| **S protein residue name** | **S protein residue number** | **S protein chain** | **S protein residue name** | **S protein residue number** | **S protein chain** |
| --- | --- | --- | --- | --- | --- |
| VAL | 1040 | A | GLN | 1036 | B |
| VAL | 1068 | A | GLY | 891 | B |
| ILE | 788 | A | SER | 704 | C |
| LEU | 966 | A | ALA | 570 | C |
| VAL | 785 | A | LEU | 699 | C |
| ILE | 870 | A | LEU | 699 | C |
| SER | 758 | A | LYS | 964 | C |
| TYR | 380 | A | ILE | 973 | B |
| GLN | 762 | A | THR | 1006 | C |
| ASN | 710 | A | PRO | 897 | B |
| VAL | 42 | A | PHE | 562 | C |
| THR | 430 | A | ARG | 983 | B |
| LYS | 557 | A | SER | 45 | B |
| SER | 383 | A | LEU | 984 | B |
| GLU | 918 | A | SER | 1123 | C |
| GLN | 1036 | A | VAL | 1040 | C |
| PHE | 759 | A | GLN | 965 | C |
| THR | 696 | A | MET | 869 | B |
| PHE | 759 | A | GLN | 1002 | C |
| GLN | 314 | A | SER | 735 | B |
| GLY | 999 | A | PHE | 759 | B |
| GLN | 755 | A | SER | 968 | C |
| ASP | 1041 | A | GLN | 784 | B |
| GLN | 414 | A | VAL | 987 | C |
| SER | 968 | A | GLN | 755 | B |
| TYR | 707 | A | ILE | 896 | B |
| THR | 859 | A | GLN | 613 | C |
| THR | 883 | A | TYR | 707 | C |
| SER | 1030 | A | ASP | 1041 | C |
| ASP | 1041 | A | GLY | 889 | B |
| GLN | 314 | A | LEU | 861 | B |
| MET | 869 | A | GLY | 669 | C |
| GLN | 895 | A | ALA | 713 | C |
| ALA | 890 | A | VAL | 1068 | C |
| GLY | 232 | A | GLU | 465 | C |
| TYR | 38 | A | PHE | 562 | C |
| ARG | 1091 | A | ASP | 1118 | B |
| LEU | 560 | A | TYR | 38 | B |
| GLN | 784 | A | ASP | 1041 | C |
| ILE | 569 | A | VAL | 47 | B |
| GLU | 702 | A | GLN | 787 | B |
| GLN | 1002 | A | GLN | 1002 | C |
| PHE | 43 | A | LEU | 560 | C |
| GLN | 913 | A | PRO | 1090 | C |
| ALA | 892 | A | GLU | 1072 | C |
| ARG | 44 | A | ARG | 567 | C |
| GLU | 1144 | A | LEU | 1145 | C |
| GLU | 988 | A | SER | 383 | C |
| TYR | 279 | A | GLN | 563 | C |
| GLY | 669 | A | LEU | 864 | B |
| ALA | 890 | A | TYR | 1067 | C |
| LEU | 864 | A | CYS | 662 | C |
| PRO | 521 | A | LYS | 41 | B |
| GLY | 757 | A | GLN | 965 | C |
| SER | 758 | A | THR | 961 | C |
| ASP | 979 | A | ARG | 567 | C |
| TYR | 904 | A | VAL | 1094 | C |
| TYR | 707 | A | GLY | 798 | B |
| PRO | 384 | A | GLU | 988 | B |
| PRO | 897 | A | ASN | 709 | C |
| GLY | 1046 | A | GLY | 889 | B |
| TYR | 917 | A | VAL | 1129 | C |
| VAL | 705 | A | PRO | 792 | B |
| THR | 887 | A | TYR | 1047 | C |
| PHE | 970 | A | GLN | 755 | B |
| ASN | 703 | A | TYR | 789 | B |
| SER | 383 | A | GLU | 988 | B |
| ALA | 892 | A | PRO | 1069 | C |
| PHE | 559 | A | PHE | 43 | B |
| PRO | 230 | A | ARG | 355 | C |
| ALA | 570 | A | LEU | 966 | B |
| ILE | 569 | A | VAL | 963 | B |
| GLN | 564 | A | PHE | 43 | B |
| PRO | 862 | A | ALA | 668 | C |
| ALA | 899 | A | TYR | 707 | C |
| THR | 1009 | A | THR | 1009 | C |
| ASN | 978 | A | THR | 547 | C |
| CYS | 379 | A | GLU | 988 | B |
| GLY | 594 | A | THR | 859 | B |
| LEU | 699 | A | LYS | 786 | B |
| LEU | 517 | A | ARG | 983 | B |
| GLN | 787 | A | ALA | 701 | C |
| TYR | 707 | A | ALA | 899 | B |
| ILE | 896 | A | ALA | 713 | C |
| ILE | 1013 | A | LEU | 1012 | B |
| GLY | 199 | A | ARG | 355 | C |
| TYR | 1047 | A | TRP | 886 | B |
| ILE | 233 | A | GLU | 465 | C |
| VAL | 1094 | A | ILE | 896 | B |
| VAL | 976 | A | LEU | 546 | C |
| GLN | 965 | A | TYR | 756 | B |
| ASP | 979 | A | LEU | 546 | C |
| LEU | 1141 | A | LEU | 1141 | B |
| LYS | 386 | A | LEU | 981 | B |
| ASN | 703 | A | LYS | 790 | B |
| LEU | 1145 | A | LEU | 1145 | B |
| ASP | 1139 | A | GLU | 1144 | B |
| GLY | 413 | A | LYS | 986 | C |
| ARG | 983 | A | LYS | 386 | C |
| LEU | 865 | A | MET | 697 | C |
| ALA | 570 | A | ASN | 856 | B |
| THR | 547 | A | SER | 982 | B |
| GLU | 1072 | A | LEU | 894 | B |
| TRP | 886 | A | TYR | 1047 | C |
| GLN | 913 | A | ARG | 1107 | C |
| GLN | 1142 | A | GLU | 1144 | B |
| GLN | 895 | A | SER | 708 | C |
| PHE | 855 | A | PRO | 589 | C |
| ASP | 796 | A | TYR | 707 | C |
| THR | 385 | A | THR | 415 | C |
| GLY | 1093 | A | GLN | 913 | B |
| LEU | 894 | A | GLU | 1072 | C |
| ASN | 856 | A | ASP | 568 | C |
| MET | 900 | A | PRO | 1079 | C |
| PRO | 863 | A | ALA | 668 | C |
| GLY | 857 | A | PHE | 592 | C |
| ASP | 427 | A | GLU | 990 | C |
| GLU | 773 | A | GLU | 1017 | C |
| ASP | 985 | A | THR | 385 | C |
| ALA | 570 | A | LYS | 964 | B |
| GLN | 1002 | A | TYR | 756 | B |
| LEU | 894 | A | ALA | 713 | C |
| LEU | 981 | A | LYS | 386 | C |
| GLY | 669 | A | MET | 869 | B |
| TYR | 904 | A | ARG | 1107 | C |
| GLY | 798 | A | TYR | 707 | C |
| VAL | 705 | A | THR | 883 | B |
| GLY | 798 | A | ILE | 1130 | C |
| LEU | 894 | A | ILE | 714 | C |
| VAL | 1128 | A | ASN | 919 | B |
| VAL | 963 | A | ASP | 568 | C |
| LYS | 790 | A | ASN | 703 | C |
| GLN | 314 | A | THR | 768 | B |
| VAL | 705 | A | ALA | 879 | B |
| ASP | 198 | A | ASP | 428 | C |
| PHE | 1089 | A | ASN | 914 | B |
| HIS | 519 | A | VAL | 42 | B |
| THR | 588 | A | PHE | 855 | B |
| VAL | 705 | A | GLN | 895 | B |
| MET | 740 | A | CYS | 590 | C |
| PRO | 412 | A | VAL | 987 | C |
| ARG | 1039 | A | ARG | 1039 | C |
| ASP | 228 | A | TYR | 396 | C |
| ARG | 1107 | A | TRP | 886 | B |
| SER | 704 | A | ILE | 788 | B |
| TYR | 1047 | A | ALA | 890 | B |
| VAL | 1094 | A | TYR | 904 | B |
| ARG | 765 | A | THR | 961 | C |
| ARG | 765 | A | GLN | 957 | C |
| ILE | 231 | A | ARG | 355 | C |
| SER | 383 | A | ASP | 985 | B |
| THR | 961 | A | GLN | 762 | B |
| ILE | 1013 | A | ALA | 1016 | B |
| PRO | 665 | A | LEU | 864 | B |
| GLN | 613 | A | THR | 859 | B |
| LEU | 1141 | A | LEU | 1141 | C |
| ASP | 994 | A | PHE | 970 | C |
| LEU | 560 | A | ASN | 282 | B |
| TYR | 200 | A | SER | 514 | C |
| ASN | 856 | A | PHE | 592 | C |
| SER | 1003 | A | PHE | 759 | B |
| ASN | 282 | A | LYS | 558 | C |
| GLU | 1072 | A | ALA | 893 | B |
| GLY | 889 | A | LYS | 1045 | C |
| GLN | 787 | A | ASN | 703 | C |
| GLN | 787 | A | GLY | 700 | C |
| ILE | 712 | A | MET | 900 | B |
| TYR | 873 | A | LEU | 699 | C |
| GLN | 762 | A | THR | 961 | C |
| LEU | 966 | A | ASP | 571 | C |
| GLU | 918 | A | VAL | 1128 | C |
| TYR | 200 | A | ARG | 355 | C |
| PHE | 592 | A | THR | 859 | B |
| ARG | 1019 | A | GLU | 1017 | C |
| LYS | 1045 | A | GLY | 889 | B |
| GLN | 762 | A | TYR | 1007 | C |
| PRO | 1069 | A | ALA | 890 | B |
| VAL | 963 | A | ILE | 569 | C |
| LYS | 558 | A | PHE | 43 | B |
| TYR | 707 | A | ILE | 882 | B |
| GLY | 889 | A | ASP | 1041 | C |
| ASN | 317 | A | GLY | 857 | B |
| PHE | 43 | A | LYS | 557 | C |
| PHE | 562 | A | LYS | 41 | B |
| ILE | 1013 | A | ILE | 1013 | C |
| VAL | 382 | A | GLU | 988 | B |
| GLU | 1111 | A | VAL | 1122 | C |
| PHE | 562 | A | PRO | 225 | B |
| VAL | 1094 | A | MET | 900 | B |
| PRO | 862 | A | GLY | 667 | C |
| GLY | 566 | A | VAL | 42 | B |
| LEU | 560 | A | PHE | 43 | B |
| ALA | 701 | A | ILE | 788 | B |
| THR | 1027 | A | ARG | 1039 | C |
| THR | 961 | A | SER | 758 | B |
| ARG | 983 | A | GLY | 381 | C |
| THR | 167 | A | ARG | 466 | C |
| SER | 982 | A | ASP | 389 | C |
| THR | 547 | A | ASP | 979 | B |
| MET | 900 | A | THR | 1077 | C |
| PRO | 1079 | A | GLN | 913 | B |
| LYS | 1045 | A | ALA | 890 | B |
| GLN | 1010 | A | LEU | 1012 | B |
| GLY | 667 | A | PRO | 863 | B |
| PHE | 898 | A | TYR | 707 | C |
| SER | 1123 | A | ASN | 914 | B |
| ILE | 712 | A | TYR | 904 | B |
| ILE | 973 | A | GLY | 381 | C |
| PHE | 759 | A | GLY | 999 | C |
| LEU | 1145 | A | LEU | 1141 | B |
| ASP | 1041 | A | SER | 1030 | B |
| GLY | 880 | A | VAL | 705 | C |
| PHE | 43 | A | LYS | 558 | C |
| THR | 1006 | A | GLN | 1005 | B |
| ASN | 914 | A | PHE | 1121 | C |
| ARG | 983 | A | VAL | 382 | C |
| THR | 883 | A | VAL | 705 | C |
| LEU | 1034 | A | ASP | 1041 | C |
| GLY | 1046 | A | ALA | 890 | B |
| LYS | 790 | A | VAL | 705 | C |
| GLU | 918 | A | VAL | 1129 | C |
| GLY | 566 | A | PHE | 43 | B |
| LYS | 786 | A | GLY | 700 | C |
| LEU | 560 | A | GLY | 283 | B |
| ASP | 40 | A | PHE | 562 | C |
| LYS | 41 | A | ALA | 520 | C |
| PHE | 970 | A | PHE | 759 | B |
| LEU | 894 | A | ARG | 1107 | C |
| PRO | 1069 | A | ALA | 892 | B |
| MET | 869 | A | THR | 696 | C |
| TYR | 200 | A | TYR | 396 | C |
| LEU | 1012 | A | GLN | 1010 | C |
| PHE | 1121 | A | GLN | 913 | B |
| GLN | 564 | A | VAL | 42 | B |
| ASP | 40 | A | GLN | 563 | C |
| ARG | 1039 | A | GLU | 1031 | B |
| SER | 698 | A | TYR | 873 | B |
| GLN | 563 | A | LYS | 41 | B |
| ALA | 1078 | A | MET | 900 | B |
| GLU | 1144 | A | LEU | 1141 | C |
| SER | 967 | A | ILE | 569 | C |
| SER | 975 | A | ALA | 570 | C |
| ALA | 701 | A | GLN | 787 | B |
| GLN | 957 | A | ARG | 765 | B |
| TYR | 707 | A | LYS | 795 | B |
| ALA | 570 | A | VAL | 963 | B |
| GLU | 1031 | A | VAL | 1040 | C |
| LYS | 557 | A | PHE | 43 | B |
| GLU | 1072 | A | ALA | 892 | B |
| PHE | 43 | A | GLN | 563 | C |
| THR | 859 | A | GLY | 594 | C |
| ALA | 713 | A | LEU | 894 | B |
| GLY | 413 | A | VAL | 987 | C |
| VAL | 382 | A | LEU | 984 | B |
| VAL | 1040 | A | SER | 1030 | B |
| ALA | 1016 | A | ILE | 1013 | C |
| VAL | 47 | A | ILE | 569 | C |
| VAL | 382 | A | ARG | 983 | B |
| ILE | 882 | A | TYR | 707 | C |
| PRO | 863 | A | GLY | 669 | C |
| GLY | 891 | A | GLY | 1046 | C |
| SER | 967 | A | ALA | 570 | C |
| THR | 302 | A | THR | 761 | B |
| GLY | 891 | A | PRO | 1069 | C |
| PHE | 43 | A | PHE | 565 | C |
| ASN | 969 | A | GLN | 755 | B |
| ALA | 668 | A | LEU | 864 | B |
| PHE | 562 | A | TYR | 38 | B |
| ILE | 788 | A | LEU | 699 | C |
| GLU | 1092 | A | GLN | 913 | B |
| GLY | 283 | A | GLN | 563 | C |
| ARG | 1107 | A | LEU | 894 | B |
| GLN | 965 | A | GLN | 762 | B |
| TYR | 396 | A | PRO | 230 | B |
| TYR | 707 | A | GLN | 895 | B |
| SER | 1030 | A | VAL | 1040 | C |
| GLN | 895 | A | SER | 711 | C |
| MET | 697 | A | LEU | 865 | B |
| PHE | 43 | A | PHE | 559 | C |
| ALA | 668 | A | LEU | 865 | B |
| ILE | 670 | A | LEU | 864 | B |
| LEU | 699 | A | ILE | 788 | B |
| LEU | 699 | A | VAL | 785 | B |
| THR | 1009 | A | ILE | 1013 | C |
| LEU | 984 | A | SER | 383 | C |
| LEU | 861 | A | GLN | 613 | C |
| ASP | 994 | A | ARG | 995 | C |
| LEU | 894 | A | VAL | 705 | C |
| LYS | 921 | A | ILE | 1130 | C |
| MET | 900 | A | PHE | 1095 | C |
| TYR | 707 | A | THR | 883 | B |
| ALA | 899 | A | PRO | 1079 | C |
| SER | 758 | A | GLN | 965 | C |
| LEU | 699 | A | GLN | 872 | B |
| LYS | 41 | A | GLN | 564 | C |
| LEU | 1034 | A | VAL | 1040 | C |
| ILE | 794 | A | TYR | 707 | C |
| MET | 697 | A | TYR | 873 | B |
| SER | 982 | A | THR | 547 | C |
| TYR | 917 | A | ILE | 1130 | C |
| LEU | 858 | A | PHE | 592 | C |
| LEU | 984 | A | GLY | 381 | C |
| ALA | 701 | A | TYR | 789 | B |
| GLU | 1031 | A | ARG | 1039 | C |
| TYR | 756 | A | GLN | 1002 | C |
| GLY | 669 | A | PRO | 863 | B |
| TYR | 707 | A | ILE | 794 | B |
| GLN | 1005 | A | THR | 1006 | C |
| GLY | 971 | A | TYR | 756 | B |
| GLU | 1111 | A | SER | 1123 | C |
| PRO | 792 | A | TYR | 707 | C |
| ASP | 1041 | A | ALA | 890 | B |
| GLU | 1017 | A | GLU | 773 | B |
| PHE | 565 | A | PHE | 43 | B |
| GLN | 1002 | A | GLN | 1005 | B |
| ALA | 668 | A | PRO | 863 | B |
| ASP | 796 | A | SER | 708 | C |
| ARG | 357 | A | PRO | 230 | B |
| SER | 735 | A | GLN | 314 | C |
| VAL | 963 | A | ALA | 570 | C |
| ALA | 890 | A | GLY | 1046 | C |
| LEU | 894 | A | PRO | 1069 | C |
| ARG | 983 | A | LEU | 390 | C |
| GLY | 669 | A | THR | 866 | B |
| ASN | 914 | A | SER | 1123 | C |
| GLN | 564 | A | LYS | 41 | B |
| PHE | 1089 | A | TYR | 917 | B |
| PRO | 897 | A | SER | 711 | C |
| TYR | 707 | A | PHE | 898 | B |
| GLY | 381 | A | ILE | 973 | B |
| ARG | 567 | A | ARG | 44 | B |
| GLY | 1046 | A | GLY | 891 | B |
| ILE | 231 | A | PHE | 464 | C |
| ASN | 919 | A | VAL | 1128 | C |
| ASP | 1041 | A | LEU | 1034 | B |
| LEU | 1145 | A | GLU | 1144 | B |
| ARG | 1107 | A | GLN | 913 | B |
| SER | 982 | A | LEU | 390 | C |
| GLY | 1124 | A | GLU | 918 | B |
| ALA | 893 | A | VAL | 705 | C |
| THR | 1006 | A | GLN | 762 | B |
| GLN | 755 | A | ASN | 969 | C |
| GLY | 566 | A | ARG | 44 | B |
| THR | 547 | A | LEU | 981 | B |
| VAL | 42 | A | GLN | 563 | C |
| THR | 859 | A | PHE | 592 | C |
| ALA | 570 | A | SER | 967 | B |
| ASP | 198 | A | PHE | 464 | C |
| PRO | 897 | A | TYR | 707 | C |
| ARG | 983 | A | LEU | 517 | C |
| MET | 900 | A | VAL | 1094 | C |
| VAL | 1040 | A | GLY | 889 | B |
| VAL | 42 | A | GLY | 566 | C |
| ASP | 571 | A | SER | 967 | B |
| ARG | 567 | A | PHE | 43 | B |
| THR | 549 | A | ASP | 745 | B |
| LYS | 786 | A | LEU | 699 | C |
| GLY | 381 | A | LEU | 984 | B |
| TYR | 369 | A | THR | 415 | C |
| THR | 1009 | A | THR | 1009 | B |
| ILE | 712 | A | PRO | 897 | B |
| GLN | 779 | A | MET | 697 | C |
| LYS | 790 | A | SER | 704 | C |
| SER | 1123 | A | GLU | 1111 | B |
| SER | 982 | A | LYS | 386 | C |
| THR | 547 | A | ASN | 978 | B |
| ILE | 1130 | A | LYS | 921 | B |
| LEU | 984 | A | LYS | 386 | C |
| VAL | 860 | A | ARG | 646 | C |
| PRO | 1079 | A | TYR | 917 | B |
| ASN | 709 | A | PRO | 897 | B |
| TYR | 756 | A | SER | 968 | C |
| LEU | 699 | A | LEU | 865 | B |
| TYR | 904 | A | ILE | 712 | C |
| SER | 708 | A | GLN | 895 | B |
| GLN | 913 | A | PHE | 1089 | C |
| ILE | 896 | A | ILE | 712 | C |
| SER | 704 | A | TYR | 789 | B |
| ALA | 706 | A | GLN | 895 | B |
| GLN | 1002 | A | THR | 998 | B |
| THR | 572 | A | ASN | 856 | B |
| LEU | 546 | A | ASP | 979 | B |
| VAL | 1040 | A | GLY | 1035 | B |
| ILE | 712 | A | ILE | 896 | B |
| ASN | 856 | A | ALA | 570 | C |
| LEU | 699 | A | ILE | 870 | B |
| GLN | 762 | A | GLN | 1010 | C |
| SER | 708 | A | PRO | 897 | B |
| GLY | 757 | A | PHE | 970 | C |
| GLY | 889 | A | GLY | 1046 | C |
| PHE | 592 | A | MET | 740 | B |
| VAL | 1128 | A | TYR | 917 | B |
| PHE | 970 | A | TYR | 756 | B |
| PRO | 897 | A | SER | 708 | C |
| ARG | 567 | A | VAL | 976 | B |
| LEU | 984 | A | VAL | 382 | C |
| ASN | 978 | A | GLY | 545 | C |
| ARG | 983 | A | SER | 383 | C |
| LYS | 41 | A | GLN | 563 | C |
| GLN | 895 | A | VAL | 705 | C |
| VAL | 42 | A | ARG | 567 | C |
| THR | 284 | A | LEU | 560 | C |
| ASP | 571 | A | VAL | 976 | B |
| MET | 869 | A | MET | 697 | C |
| GLN | 563 | A | TYR | 279 | B |
| GLY | 199 | A | PHE | 464 | C |
| TYR | 904 | A | GLY | 1093 | C |
| LYS | 964 | A | ILE | 569 | C |
| PRO | 1069 | A | LEU | 894 | B |
| VAL | 976 | A | THR | 572 | C |
| ALA | 890 | A | ASP | 1041 | C |
| GLN | 895 | A | ASN | 1074 | C |
| ARG | 1039 | A | ARG | 1039 | B |
| TYR | 756 | A | GLY | 971 | C |
| SER | 968 | A | GLY | 757 | B |
| PHE | 1089 | A | GLN | 913 | B |
| VAL | 1040 | A | GLU | 1031 | B |
| ILE | 1130 | A | GLY | 798 | B |
| TYR | 756 | A | ARG | 995 | C |
| SER | 375 | A | ARG | 408 | C |
| LEU | 864 | A | ILE | 666 | C |
| ASN | 317 | A | MET | 740 | B |
| ARG | 1019 | A | ALA | 1016 | C |
| SER | 758 | A | SER | 968 | C |
| VAL | 705 | A | LYS | 790 | B |
| ILE | 973 | A | TYR | 380 | C |
| LEU | 864 | A | CYS | 671 | C |
| ASN | 317 | A | ASP | 737 | B |
| PHE | 43 | A | GLY | 566 | C |
| ILE | 666 | A | LEU | 864 | B |
| ALA | 713 | A | GLN | 895 | B |
| ALA | 713 | A | ILE | 896 | B |
| LEU | 560 | A | THR | 284 | B |
| GLN | 755 | A | GLY | 971 | C |
| GLY | 700 | A | ILE | 788 | B |
| ARG | 1107 | A | TYR | 904 | B |
| THR | 859 | A | GLY | 593 | C |
| VAL | 705 | A | GLY | 880 | B |
| GLY | 757 | A | SER | 968 | C |
| LEU | 48 | A | ILE | 569 | C |
| VAL | 1128 | A | GLU | 918 | B |
| ALA | 890 | A | LYS | 1045 | C |
| ARG | 1091 | A | ASN | 907 | B |
| GLY | 413 | A | ASP | 985 | C |
| GLN | 895 | A | TYR | 707 | C |
| CYS | 166 | A | ILE | 468 | C |
| GLN | 563 | A | PHE | 43 | B |
| PRO | 792 | A | VAL | 705 | C |
| LEU | 864 | A | PRO | 665 | C |
| GLY | 700 | A | LYS | 786 | B |
| SER | 982 | A | LEU | 546 | C |
| LEU | 1145 | A | LEU | 1141 | C |
| GLN | 563 | A | ARG | 44 | B |
| THR | 1077 | A | MET | 900 | B |
| SER | 704 | A | LYS | 790 | B |
| GLU | 224 | A | PHE | 562 | C |
| PRO | 1090 | A | ASN | 907 | B |
| ILE | 233 | A | ARG | 466 | C |
| SER | 711 | A | PRO | 897 | B |
| ALA | 668 | A | MET | 869 | B |
| PHE | 797 | A | TYR | 707 | C |
| GLN | 321 | A | ASP | 745 | B |
| PRO | 1090 | A | GLN | 913 | B |
| GLN | 755 | A | ALA | 972 | C |
| ALA | 893 | A | GLU | 1072 | C |
| ILE | 1130 | A | TYR | 917 | B |
| LYS | 41 | A | HIS | 519 | C |
| ASN | 394 | A | PRO | 230 | B |
| PHE | 562 | A | GLU | 224 | B |
| GLN | 1010 | A | ALA | 766 | B |
| LYS | 41 | A | PHE | 565 | C |
| PHE | 1042 | A | GLU | 1031 | B |
| GLY | 971 | A | GLN | 755 | B |
| TYR | 707 | A | ASP | 796 | B |
| GLN | 965 | A | SER | 758 | B |
| ASP | 571 | A | SER | 975 | B |
| VAL | 42 | A | HIS | 519 | C |
| ALA | 893 | A | ASN | 703 | C |
| SER | 982 | A | GLY | 545 | C |
| VAL | 42 | A | PHE | 565 | C |
| VAL | 1129 | A | TYR | 917 | B |
| SER | 968 | A | SER | 758 | B |
| MET | 697 | A | MET | 869 | B |
| ARG | 319 | A | MET | 740 | B |
| PHE | 759 | A | THR | 1006 | C |
| TYR | 1007 | A | GLN | 762 | B |
| GLU | 224 | A | LEU | 560 | C |
| GLN | 563 | A | GLY | 283 | B |
| ILE | 788 | A | ALA | 701 | C |
| SER | 884 | A | VAL | 705 | C |
| THR | 866 | A | ALA | 668 | C |
| CYS | 662 | A | LEU | 864 | B |
| GLY | 667 | A | PRO | 862 | B |
| PHE | 592 | A | GLY | 857 | B |
| ALA | 1020 | A | ARG | 1019 | B |
| PRO | 1079 | A | ALA | 899 | B |
| GLY | 700 | A | TYR | 873 | B |
| ASP | 985 | A | LYS | 386 | C |
| PRO | 715 | A | LEU | 894 | B |
| GLN | 913 | A | GLY | 1093 | C |
| ASP | 389 | A | SER | 982 | B |
| ASP | 198 | A | PRO | 426 | C |
| ALA | 668 | A | PRO | 862 | B |
| VAL | 705 | A | SER | 884 | B |
| ASP | 985 | A | SER | 383 | C |
| LYS | 386 | A | LEU | 984 | B |
| PRO | 897 | A | ASN | 710 | C |
| ALA | 668 | A | THR | 866 | B |
| ARG | 44 | A | GLN | 563 | C |
| GLY | 889 | A | VAL | 1040 | C |
| SER | 975 | A | ASP | 571 | C |
| PRO | 862 | A | ALA | 647 | C |
| LEU | 864 | A | GLY | 667 | C |
| PHE | 1121 | A | ASN | 914 | B |
| PRO | 589 | A | PHE | 855 | B |
| ALA | 701 | A | LYS | 786 | B |
| SER | 967 | A | ASP | 571 | C |
| TRP | 886 | A | ARG | 1107 | C |
| ASP | 796 | A | ASN | 709 | C |
| LEU | 1145 | A | LEU | 1145 | C |
| ILE | 569 | A | SER | 967 | B |
| ASN | 1108 | A | TRP | 886 | B |
| GLN | 314 | A | ASP | 737 | B |
| ILE | 569 | A | LYS | 964 | B |
| ASN | 709 | A | ASP | 796 | B |
| ILE | 712 | A | GLN | 895 | B |
| PHE | 970 | A | GLY | 757 | B |
| LYS | 558 | A | ASN | 282 | B |
| TYR | 789 | A | SER | 704 | C |
| PRO | 589 | A | MET | 740 | B |
| GLN | 920 | A | ILE | 1130 | C |
| ILE | 788 | A | GLY | 700 | C |
| TYR | 789 | A | ALA | 701 | C |
| LEU | 1001 | A | GLN | 1002 | C |
| CYS | 671 | A | LEU | 864 | B |
| SER | 708 | A | ILE | 896 | B |
| HIS | 519 | A | ASP | 40 | B |
| MET | 740 | A | PHE | 592 | C |
| ASP | 40 | A | HIS | 519 | C |
| ASN | 1074 | A | GLN | 895 | B |
| LYS | 790 | A | GLU | 702 | C |
| PHE | 759 | A | SER | 1003 | C |
| TRP | 886 | A | ASN | 1108 | C |
| ASP | 737 | A | ASN | 317 | C |
| ASP | 745 | A | ARG | 319 | C |
| GLY | 283 | A | LEU | 560 | C |
| GLY | 232 | A | ARG | 466 | C |
| SER | 1123 | A | GLU | 918 | B |
| GLU | 918 | A | GLY | 1124 | C |
| GLN | 563 | A | VAL | 42 | B |
| GLY | 199 | A | PRO | 463 | C |
| VAL | 705 | A | TYR | 789 | B |
| GLN | 1010 | A | VAL | 1008 | B |
| TYR | 917 | A | PHE | 1089 | C |
| TYR | 873 | A | SER | 698 | C |
| TYR | 1067 | A | ALA | 890 | B |
| SER | 711 | A | GLN | 895 | B |
| ALA | 890 | A | PRO | 1069 | C |
| TYR | 917 | A | PRO | 1079 | C |
| PRO | 1079 | A | MET | 900 | B |
| TYR | 789 | A | ASN | 703 | C |
| LEU | 390 | A | SER | 982 | B |
| GLY | 199 | A | GLU | 465 | C |
| LYS | 964 | A | SER | 758 | B |
| TYR | 707 | A | PRO | 897 | B |
| GLN | 965 | A | GLY | 757 | B |
| SER | 1147 | A | LEU | 1145 | C |
| GLY | 232 | A | PHE | 464 | C |
| ARG | 44 | A | GLY | 566 | C |
| GLN | 1002 | A | LEU | 1001 | B |
| ILE | 788 | A | GLU | 702 | C |
| VAL | 1040 | A | LEU | 1034 | B |
| THR | 768 | A | GLN | 314 | C |
| PHE | 1095 | A | MET | 900 | B |
| GLN | 787 | A | GLU | 702 | C |
| LEU | 981 | A | THR | 547 | C |
| GLU | 918 | A | PHE | 1089 | C |
| GLY | 548 | A | ASN | 978 | B |
| PHE | 562 | A | ASP | 40 | B |
| TYR | 756 | A | GLN | 965 | C |
| GLU | 702 | A | ILE | 788 | B |
| MET | 740 | A | ARG | 319 | C |
| GLU | 702 | A | TYR | 789 | B |
| GLY | 381 | A | ARG | 983 | B |
| ILE | 666 | A | PRO | 862 | B |
| LYS | 386 | A | ASP | 985 | B |
| GLN | 563 | A | ASP | 40 | B |
| PHE | 1121 | A | THR | 912 | B |
| GLY | 669 | A | LEU | 865 | B |
| VAL | 1129 | A | GLU | 918 | B |
| ARG | 1107 | A | ASN | 907 | B |
| LEU | 894 | A | PRO | 715 | C |
| TYR | 707 | A | PHE | 797 | B |
| LEU | 699 | A | MET | 869 | B |
| LYS | 386 | A | ARG | 983 | B |
| LEU | 865 | A | ALA | 668 | C |
| HIS | 519 | A | LYS | 41 | B |
| LEU | 1141 | A | GLU | 1144 | B |
| ALA | 890 | A | TYR | 1047 | C |
| GLN | 965 | A | PHE | 759 | B |
| GLN | 613 | A | LEU | 861 | B |
| ASN | 282 | A | LEU | 560 | C |
| GLN | 895 | A | ILE | 712 | C |
| ILE | 231 | A | ARG | 466 | C |
| LYS | 786 | A | ALA | 701 | C |
| GLN | 1002 | A | GLN | 1002 | B |
| TYR | 789 | A | VAL | 705 | C |
| GLY | 700 | A | GLN | 787 | B |
| MET | 869 | A | ALA | 668 | C |
| PRO | 863 | A | GLY | 667 | C |
| TYR | 200 | A | PHE | 464 | C |
| ASN | 703 | A | GLN | 787 | B |
| VAL | 705 | A | LEU | 894 | B |
| ASP | 198 | A | PRO | 463 | C |
| LYS | 41 | A | PHE | 562 | C |
| GLY | 891 | A | VAL | 1068 | C |
| ILE | 896 | A | SER | 708 | C |
| GLN | 1005 | A | GLN | 1002 | C |
| MET | 697 | A | LEU | 864 | B |
| VAL | 976 | A | ARG | 567 | C |
| TYR | 38 | A | GLN | 563 | C |
| PHE | 759 | A | PHE | 970 | C |
| VAL | 976 | A | ASP | 571 | C |
| ILE | 714 | A | LEU | 894 | B |
| ASN | 703 | A | ILE | 788 | B |
| TYR | 873 | A | GLY | 700 | C |
| ILE | 896 | A | SER | 711 | C |
| ARG | 567 | A | VAL | 42 | B |
| ILE | 1013 | A | THR | 1009 | B |
| LEU | 865 | A | GLY | 669 | C |
| ASN | 978 | A | GLY | 548 | C |
| LYS | 1038 | A | LYS | 1038 | C |
| PRO | 230 | A | TYR | 396 | C |
| ALA | 647 | A | PRO | 862 | B |
| LEU | 865 | A | LEU | 699 | C |
| LYS | 558 | A | GLY | 283 | B |
| LEU | 864 | A | ILE | 670 | C |
| LEU | 390 | A | ARG | 983 | B |
| ALA | 879 | A | VAL | 705 | C |
| ARG | 1091 | A | GLN | 913 | B |
| TYR | 756 | A | PHE | 970 | C |
| ILE | 1013 | A | ILE | 1013 | B |
| THR | 961 | A | ARG | 765 | B |
| TYR | 917 | A | ALA | 1080 | C |
| THR | 572 | A | VAL | 976 | B |
| ILE | 788 | A | ASN | 703 | C |
| THR | 866 | A | GLY | 669 | C |
| GLY | 1093 | A | TYR | 904 | B |
| GLN | 872 | A | LEU | 699 | C |
| GLN | 913 | A | PHE | 1121 | C |
| MET | 900 | A | ALA | 1078 | C |
| THR | 385 | A | ASP | 985 | B |
| PHE | 592 | A | ASN | 856 | B |
| SER | 383 | A | ARG | 983 | B |
| PRO | 897 | A | THR | 1077 | C |
| VAL | 976 | A | ALA | 570 | C |
| MET | 869 | A | LEU | 699 | C |
| ILE | 569 | A | LEU | 48 | B |
| TYR | 38 | A | LEU | 560 | C |
| GLN | 762 | A | GLN | 965 | C |
| GLN | 1002 | A | PHE | 759 | B |
| PRO | 1069 | A | GLY | 891 | B |
| MET | 697 | A | GLN | 779 | B |
| PRO | 862 | A | ARG | 646 | C |
| ARG | 646 | A | PRO | 862 | B |
| VAL | 1068 | A | ALA | 890 | B |
| PRO | 897 | A | ILE | 712 | C |
| GLN | 913 | A | PRO | 1079 | C |
| TYR | 917 | A | VAL | 1128 | C |
| TYR | 789 | A | GLU | 702 | C |
| PHE | 1089 | A | GLU | 918 | B |
| MET | 900 | A | ASN | 709 | C |
| ILE | 896 | A | TYR | 707 | C |
| GLU | 868 | A | LEU | 699 | C |
| THR | 912 | A | PHE | 1121 | C |
| PHE | 592 | A | PHE | 855 | B |
| ASN | 907 | A | ARG | 1107 | C |
| SER | 968 | A | TYR | 756 | B |
| TYR | 873 | A | MET | 697 | C |
| SER | 708 | A | ASP | 796 | B |
| PHE | 43 | A | ARG | 567 | C |
| GLY | 545 | A | SER | 982 | B |
| LEU | 699 | A | GLN | 787 | B |
| LYS | 964 | A | ALA | 570 | C |
| ILE | 1130 | A | GLN | 920 | B |
| GLU | 1017 | A | ARG | 1019 | B |
| ASN | 856 | A | THR | 572 | C |
| ASN | 234 | A | ARG | 466 | C |
| GLN | 755 | A | PHE | 970 | C |
| LEU | 864 | A | GLY | 669 | C |
| THR | 998 | A | GLN | 1002 | C |
| MET | 900 | A | ILE | 712 | C |
| SER | 45 | A | LYS | 557 | C |
| VAL | 705 | A | ALA | 893 | B |
| GLN | 895 | A | ALA | 706 | C |
| LEU | 699 | A | TYR | 873 | B |
| ASN | 914 | A | PHE | 1089 | C |
| THR | 1077 | A | PRO | 897 | B |
| TYR | 707 | A | PRO | 792 | B |
| ARG | 319 | A | ASP | 745 | B |
| LEU | 1012 | A | ILE | 1013 | C |
| LEU | 864 | A | ALA | 668 | C |
| GLN | 563 | A | TYR | 38 | B |
| ARG | 1019 | A | ILE | 1013 | C |
| SER | 711 | A | ILE | 896 | B |
| PHE | 565 | A | LYS | 41 | B |
| LYS | 1038 | A | LYS | 1038 | B |
| GLY | 667 | A | LEU | 864 | B |
| GLU | 702 | A | LYS | 790 | B |
| TYR | 1047 | A | THR | 887 | B |
| THR | 1006 | A | PHE | 759 | B |
| PHE | 565 | A | VAL | 42 | B |
| GLN | 1010 | A | GLN | 762 | B |
| LEU | 864 | A | MET | 697 | C |
| ARG | 1039 | A | THR | 1027 | B |
| PHE | 855 | A | PHE | 592 | C |
| GLY | 1035 | A | VAL | 1040 | C |
| ALA | 520 | A | LYS | 41 | B |
| ARG | 983 | A | THR | 430 | C |
| PRO | 225 | A | PHE | 562 | C |
| LYS | 386 | A | SER | 982 | B |

**Table S5.** The list of the inter-protomer contacts of the protomer A in the structure of SARS-CoV-2 S-RRAR/G614 in the closed state (pdb id 7KDI).

| **S protein residue name** | **S protein residue number** | **S protein chain** | **S protein residue name** | **S protein residue number** | **S protein chain** |
| --- | --- | --- | --- | --- | --- |
| THR | 1027 | A | ARG | 1039 | C |
| GLN | 913 | A | GLY | 1093 | C |
| PHE | 855 | A | ASP | 574 | C |
| ASP | 796 | A | TYR | 707 | C |
| GLU | 988 | A | PRO | 384 | C |
| GLY | 1046 | A | ALA | 890 | B |
| THR | 385 | A | ASP | 985 | B |
| PHE | 855 | A | THR | 588 | C |
| VAL | 1068 | A | ALA | 892 | B |
| LEU | 894 | A | GLU | 1072 | C |
| ASP | 571 | A | ARG | 44 | B |
| PHE | 592 | A | THR | 859 | B |
| PHE | 1042 | A | SER | 1030 | B |
| VAL | 1040 | A | GLN | 1036 | B |
| GLN | 755 | A | PHE | 970 | C |
| GLY | 381 | A | ARG | 983 | B |
| TYR | 1047 | A | TRP | 886 | B |
| ARG | 567 | A | ARG | 44 | B |
| ARG | 646 | A | PRO | 862 | B |
| ARG | 44 | A | GLN | 563 | C |
| LEU | 984 | A | LYS | 386 | C |
| THR | 549 | A | ASP | 745 | B |
| ILE | 712 | A | GLN | 895 | B |
| GLN | 895 | A | TYR | 707 | C |
| ASP | 985 | A | THR | 415 | B |
| ASP | 994 | A | PHE | 970 | C |
| MET | 900 | A | PHE | 1095 | C |
| LEU | 546 | A | VAL | 976 | B |
| TYR | 917 | A | VAL | 1129 | C |
| PRO | 897 | A | THR | 1077 | C |
| THR | 961 | A | ARG | 765 | B |
| THR | 1009 | A | THR | 1009 | C |
| GLU | 702 | A | GLN | 787 | B |
| ASP | 571 | A | SER | 967 | B |
| ASN | 856 | A | ALA | 570 | C |
| LEU | 1141 | A | LEU | 1141 | C |
| ILE | 1130 | A | TYR | 917 | B |
| LYS | 786 | A | ALA | 701 | C |
| SER | 967 | A | ALA | 570 | C |
| PHE | 464 | A | GLY | 232 | B |
| ALA | 890 | A | PRO | 1069 | C |
| VAL | 1128 | A | GLU | 918 | B |
| ILE | 666 | A | LEU | 864 | B |
| VAL | 736 | A | GLN | 314 | C |
| ASN | 914 | A | PHE | 1121 | C |
| GLY | 1035 | A | VAL | 1040 | C |
| ARG | 1107 | A | LEU | 894 | B |
| GLU | 465 | A | GLY | 232 | B |
| PHE | 592 | A | GLY | 857 | B |
| LYS | 786 | A | LEU | 699 | C |
| LEU | 390 | A | ARG | 983 | B |
| MET | 697 | A | LEU | 865 | B |
| SER | 735 | A | GLN | 314 | C |
| PRO | 589 | A | PHE | 855 | B |
| GLN | 895 | A | ILE | 712 | C |
| VAL | 1128 | A | TYR | 917 | B |
| PHE | 1042 | A | GLU | 1031 | B |
| GLN | 613 | A | LEU | 861 | B |
| THR | 588 | A | PHE | 855 | B |
| GLY | 889 | A | ASP | 1041 | C |
| VAL | 705 | A | GLN | 895 | B |
| THR | 1077 | A | MET | 900 | B |
| LEU | 861 | A | GLN | 613 | C |
| LEU | 894 | A | ARG | 1107 | C |
| PRO | 792 | A | VAL | 705 | C |
| SER | 704 | A | ILE | 788 | B |
| PRO | 863 | A | GLY | 667 | C |
| GLN | 965 | A | PHE | 759 | B |
| TYR | 279 | A | GLN | 563 | C |
| GLY | 891 | A | LYS | 1045 | C |
| GLN | 1005 | A | GLN | 1002 | C |
| ILE | 712 | A | TYR | 904 | B |
| ILE | 569 | A | ASN | 960 | B |
| ILE | 712 | A | PRO | 897 | B |
| GLU | 1144 | A | LEU | 1141 | C |
| SER | 967 | A | ILE | 569 | C |
| VAL | 1040 | A | ALA | 890 | B |
| GLY | 548 | A | ASN | 978 | B |
| THR | 572 | A | ASN | 856 | B |
| ILE | 788 | A | SER | 704 | C |
| PHE | 562 | A | VAL | 42 | B |
| GLN | 1010 | A | VAL | 1008 | B |
| LEU | 754 | A | SER | 968 | C |
| GLN | 1002 | A | GLN | 1002 | B |
| GLY | 1124 | A | GLU | 918 | B |
| LYS | 964 | A | ILE | 569 | C |
| PRO | 230 | A | GLU | 516 | C |
| TYR | 707 | A | ILE | 896 | B |
| ASP | 737 | A | ARG | 319 | C |
| PRO | 897 | A | ILE | 712 | C |
| LEU | 894 | A | PRO | 715 | C |
| GLN | 872 | A | LEU | 699 | C |
| TYR | 707 | A | PRO | 792 | B |
| ASP | 737 | A | GLN | 314 | C |
| GLN | 1002 | A | GLN | 1005 | B |
| ARG | 567 | A | VAL | 976 | B |
| VAL | 785 | A | LEU | 699 | C |
| GLY | 971 | A | TYR | 756 | B |
| VAL | 705 | A | ALA | 893 | B |
| ASP | 389 | A | SER | 982 | B |
| SER | 982 | A | ASP | 389 | C |
| PHE | 559 | A | PHE | 43 | B |
| THR | 883 | A | VAL | 705 | C |
| LEU | 984 | A | GLY | 381 | C |
| GLY | 891 | A | VAL | 1068 | C |
| LEU | 1145 | A | LEU | 1145 | B |
| GLN | 895 | A | VAL | 705 | C |
| ARG | 1019 | A | GLU | 1017 | C |
| ASN | 856 | A | PHE | 592 | C |
| LYS | 41 | A | PHE | 565 | C |
| LYS | 386 | A | LEU | 984 | B |
| PHE | 592 | A | LEU | 858 | B |
| MET | 869 | A | LEU | 699 | C |
| MET | 900 | A | PRO | 1079 | C |
| GLN | 965 | A | GLN | 762 | B |
| GLY | 669 | A | PRO | 863 | B |
| VAL | 705 | A | ALA | 879 | B |
| TYR | 756 | A | PHE | 970 | C |
| PHE | 592 | A | MET | 740 | B |
| GLN | 779 | A | MET | 697 | C |
| LYS | 41 | A | GLN | 563 | C |
| THR | 415 | A | THR | 385 | B |
| THR | 1006 | A | PHE | 759 | B |
| ILE | 1013 | A | LEU | 1012 | B |
| THR | 415 | A | TYR | 369 | B |
| GLY | 798 | A | TYR | 707 | C |
| LEU | 699 | A | ILE | 788 | B |
| MET | 900 | A | ILE | 712 | C |
| ILE | 712 | A | ILE | 896 | B |
| PRO | 862 | A | ALA | 647 | C |
| GLU | 988 | A | TYR | 380 | C |
| THR | 1006 | A | GLN | 1005 | B |
| LEU | 699 | A | GLN | 787 | B |
| TYR | 873 | A | MET | 697 | C |
| ILE | 788 | A | ASN | 703 | C |
| LEU | 864 | A | GLY | 669 | C |
| GLU | 988 | A | VAL | 382 | C |
| GLN | 1005 | A | THR | 1006 | C |
| VAL | 47 | A | ILE | 569 | C |
| SER | 711 | A | GLN | 895 | B |
| ALA | 701 | A | GLN | 787 | B |
| ARG | 765 | A | ALA | 958 | C |
| PRO | 230 | A | ASN | 394 | C |
| LEU | 518 | A | TYR | 200 | B |
| ILE | 1130 | A | LYS | 921 | B |
| VAL | 705 | A | THR | 883 | B |
| MET | 697 | A | MET | 869 | B |
| LEU | 984 | A | TYR | 380 | C |
| SER | 758 | A | LYS | 964 | C |
| PRO | 225 | A | PHE | 562 | C |
| GLU | 224 | A | PRO | 561 | C |
| PHE | 43 | A | GLN | 563 | C |
| VAL | 963 | A | ASP | 568 | C |
| VAL | 382 | A | ARG | 983 | B |
| ASP | 198 | A | ARG | 355 | C |
| ASP | 40 | A | PHE | 562 | C |
| PHE | 1121 | A | THR | 912 | B |
| VAL | 705 | A | GLY | 880 | B |
| SER | 704 | A | TYR | 789 | B |
| ASN | 914 | A | PHE | 1089 | C |
| THR | 883 | A | TYR | 707 | C |
| ALA | 668 | A | MET | 869 | B |
| ILE | 569 | A | SER | 967 | B |
| ASN | 703 | A | LYS | 790 | B |
| ASP | 985 | A | SER | 383 | C |
| PHE | 759 | A | GLN | 965 | C |
| LEU | 699 | A | GLN | 872 | B |
| PRO | 463 | A | GLY | 199 | B |
| PHE | 565 | A | PHE | 43 | B |
| LYS | 1045 | A | LYS | 786 | B |
| PHE | 565 | A | LYS | 41 | B |
| ALA | 706 | A | GLN | 895 | B |
| PHE | 1121 | A | GLN | 913 | B |
| TYR | 1067 | A | ALA | 890 | B |
| ILE | 896 | A | VAL | 1094 | C |
| GLU | 516 | A | TYR | 200 | B |
| VAL | 705 | A | THR | 791 | B |
| SER | 982 | A | GLY | 545 | C |
| TYR | 756 | A | GLN | 1002 | C |
| THR | 912 | A | PHE | 1121 | C |
| THR | 1077 | A | PRO | 897 | B |
| PHE | 1089 | A | ASN | 914 | B |
| GLY | 1093 | A | GLN | 913 | B |
| GLY | 880 | A | VAL | 705 | C |
| ASP | 40 | A | HIS | 519 | C |
| GLN | 563 | A | LYS | 41 | B |
| ARG | 466 | A | THR | 167 | B |
| ASN | 703 | A | ILE | 788 | B |
| LYS | 41 | A | HIS | 519 | C |
| VAL | 1040 | A | TRP | 886 | B |
| PHE | 43 | A | ARG | 567 | C |
| ILE | 569 | A | VAL | 47 | B |
| SER | 1030 | A | VAL | 1040 | C |
| PHE | 1095 | A | MET | 900 | B |
| PHE | 759 | A | THR | 1006 | C |
| ASN | 317 | A | ASP | 737 | B |
| ALA | 713 | A | ALA | 893 | B |
| ASN | 317 | A | ASN | 764 | B |
| LEU | 560 | A | TYR | 38 | B |
| PRO | 230 | A | TYR | 396 | C |
| GLY | 669 | A | LEU | 865 | B |
| ILE | 788 | A | ALA | 701 | C |
| ASN | 710 | A | PRO | 897 | B |
| ILE | 1013 | A | ARG | 1019 | B |
| TRP | 886 | A | ASN | 1108 | C |
| PHE | 1089 | A | GLN | 913 | B |
| PHE | 464 | A | ASP | 198 | B |
| ASN | 751 | A | GLN | 52 | C |
| SER | 1030 | A | ASP | 1041 | C |
| GLN | 895 | A | ALA | 713 | C |
| ALA | 1078 | A | MET | 900 | B |
| VAL | 705 | A | PRO | 792 | B |
| PRO | 1090 | A | GLN | 913 | B |
| ARG | 765 | A | GLN | 957 | C |
| LEU | 754 | A | GLN | 52 | C |
| LEU | 864 | A | PRO | 665 | C |
| GLN | 762 | A | THR | 961 | C |
| THR | 572 | A | VAL | 976 | B |
| THR | 1009 | A | THR | 1009 | B |
| GLN | 1036 | A | LYS | 1038 | C |
| SER | 711 | A | PRO | 897 | B |
| SER | 698 | A | TYR | 873 | B |
| LYS | 790 | A | VAL | 705 | C |
| LEU | 699 | A | LEU | 865 | B |
| LEU | 1141 | A | LEU | 1141 | B |
| LEU | 560 | A | GLY | 283 | B |
| ARG | 319 | A | MET | 740 | B |
| ARG | 1019 | A | ILE | 1013 | C |
| LEU | 1141 | A | GLU | 1144 | B |
| GLU | 773 | A | GLU | 1017 | C |
| THR | 430 | A | ARG | 983 | B |
| GLY | 798 | A | ILE | 1130 | C |
| SER | 383 | A | ARG | 983 | B |
| TYR | 200 | A | TYR | 396 | C |
| ASP | 428 | A | ASP | 198 | B |
| PHE | 970 | A | GLN | 755 | B |
| ALA | 713 | A | ILE | 896 | B |
| LYS | 1045 | A | ALA | 890 | B |
| GLU | 1017 | A | GLU | 773 | B |
| LEU | 1141 | A | LEU | 1145 | B |
| GLN | 1010 | A | GLN | 762 | B |
| GLY | 283 | A | GLN | 563 | C |
| ARG | 983 | A | LYS | 386 | C |
| ILE | 1130 | A | GLN | 920 | B |
| ARG | 1091 | A | ASN | 907 | B |
| GLY | 545 | A | SER | 982 | B |
| TYR | 707 | A | GLN | 895 | B |
| PRO | 897 | A | ASN | 710 | C |
| VAL | 1068 | A | ALA | 890 | B |
| MET | 900 | A | VAL | 1094 | C |
| TYR | 200 | A | ARG | 355 | C |
| ARG | 983 | A | LEU | 517 | C |
| CYS | 671 | A | LEU | 864 | B |
| ARG | 466 | A | ILE | 231 | B |
| GLU | 1072 | A | LEU | 894 | B |
| LEU | 390 | A | SER | 982 | B |
| HIS | 49 | A | ASP | 571 | C |
| GLY | 667 | A | PRO | 862 | B |
| ILE | 712 | A | MET | 900 | B |
| GLN | 1002 | A | LEU | 1001 | B |
| GLU | 1031 | A | VAL | 1040 | C |
| TYR | 200 | A | LEU | 518 | C |
| THR | 866 | A | ALA | 668 | C |
| ARG | 567 | A | PHE | 43 | B |
| THR | 859 | A | GLN | 613 | C |
| ARG | 403 | A | ALA | 372 | B |
| LYS | 386 | A | ARG | 983 | B |
| GLY | 1046 | A | GLY | 891 | B |
| LEU | 865 | A | LEU | 699 | C |
| ILE | 569 | A | VAL | 963 | B |
| ALA | 713 | A | LEU | 894 | B |
| GLN | 564 | A | VAL | 42 | B |
| ALA | 890 | A | ASP | 1041 | C |
| PRO | 1069 | A | LEU | 894 | B |
| GLY | 889 | A | GLY | 1046 | C |
| LEU | 518 | A | LYS | 202 | B |
| GLU | 918 | A | PHE | 1089 | C |
| ALA | 892 | A | GLU | 1072 | C |
| PHE | 970 | A | TYR | 756 | B |
| THR | 547 | A | ASN | 978 | B |
| ASN | 703 | A | GLN | 787 | B |
| GLN | 913 | A | PHE | 1121 | C |
| ALA | 892 | A | PRO | 1069 | C |
| ASP | 985 | A | LYS | 386 | C |
| ALA | 668 | A | PRO | 863 | B |
| SER | 383 | A | LEU | 984 | B |
| GLN | 895 | A | SER | 708 | C |
| PHE | 592 | A | PHE | 855 | B |
| TYR | 707 | A | ASP | 796 | B |
| LEU | 865 | A | MET | 697 | C |
| SER | 1123 | A | GLU | 1111 | B |
| LEU | 861 | A | GLN | 314 | C |
| ALA | 1016 | A | ILE | 1013 | C |
| ALA | 570 | A | VAL | 976 | B |
| LEU | 1141 | A | LEU | 1145 | C |
| GLY | 669 | A | THR | 866 | B |
| ASP | 571 | A | VAL | 976 | B |
| GLN | 895 | A | ASN | 1074 | C |
| TYR | 789 | A | VAL | 705 | C |
| ASP | 1041 | A | GLN | 784 | B |
| PRO | 230 | A | ARG | 357 | C |
| VAL | 1068 | A | GLY | 891 | B |
| ALA | 890 | A | VAL | 1068 | C |
| ASN | 978 | A | GLY | 545 | C |
| ASP | 796 | A | SER | 708 | C |
| GLN | 913 | A | ARG | 1107 | C |
| VAL | 1040 | A | LEU | 1034 | B |
| LYS | 386 | A | LEU | 981 | B |
| ALA | 701 | A | LYS | 786 | B |
| PHE | 898 | A | TYR | 707 | C |
| PHE | 592 | A | ASN | 856 | B |
| GLU | 918 | A | VAL | 1128 | C |
| ARG | 466 | A | GLN | 115 | B |
| TYR | 904 | A | ILE | 712 | C |
| GLN | 564 | A | LYS | 41 | B |
| VAL | 42 | A | GLN | 563 | C |
| LEU | 894 | A | VAL | 705 | C |
| ASP | 979 | A | LEU | 546 | C |
| ALA | 570 | A | ASN | 856 | B |
| THR | 768 | A | GLN | 314 | C |
| THR | 761 | A | THR | 302 | C |
| GLN | 1036 | A | VAL | 1040 | C |
| LYS | 790 | A | GLU | 702 | C |
| ARG | 983 | A | SER | 383 | C |
| ILE | 882 | A | TYR | 707 | C |
| TYR | 917 | A | VAL | 1128 | C |
| LEU | 1034 | A | ASP | 1041 | C |
| GLY | 891 | A | GLY | 1046 | C |
| TYR | 1047 | A | THR | 887 | B |
| LYS | 558 | A | GLY | 283 | B |
| ALA | 890 | A | GLY | 1046 | C |
| GLY | 999 | A | PHE | 759 | B |
| SER | 708 | A | GLN | 895 | B |
| ALA | 668 | A | LEU | 865 | B |
| TYR | 917 | A | ILE | 1130 | C |
| PRO | 384 | A | GLU | 988 | B |
| ARG | 1107 | A | TRP | 886 | B |
| PHE | 592 | A | LYS | 854 | B |
| ILE | 896 | A | TYR | 707 | C |
| SER | 968 | A | GLN | 755 | B |
| VAL | 976 | A | ARG | 567 | C |
| LEU | 1034 | A | VAL | 1040 | C |
| PHE | 43 | A | PHE | 565 | C |
| GLN | 755 | A | GLY | 971 | C |
| THR | 1009 | A | GLN | 1010 | C |
| ALA | 890 | A | LYS | 1045 | C |
| VAL | 1128 | A | ASN | 919 | B |
| PHE | 43 | A | GLY | 566 | C |
| ASP | 796 | A | ASN | 709 | C |
| SER | 968 | A | GLY | 757 | B |
| ASN | 919 | A | VAL | 1128 | C |
| LYS | 947 | A | LYS | 776 | B |
| GLN | 1002 | A | PHE | 759 | B |
| TYR | 38 | A | LEU | 560 | C |
| PRO | 589 | A | MET | 740 | B |
| ASN | 370 | A | LYS | 417 | C |
| LYS | 202 | A | LEU | 518 | C |
| GLY | 566 | A | PHE | 43 | B |
| ASP | 571 | A | SER | 975 | B |
| TYR | 904 | A | GLY | 1093 | C |
| PHE | 855 | A | PRO | 589 | C |
| PHE | 797 | A | TYR | 707 | C |
| GLN | 957 | A | ARG | 765 | B |
| VAL | 963 | A | ALA | 570 | C |
| SER | 1030 | A | PHE | 1042 | C |
| MET | 900 | A | THR | 1077 | C |
| GLU | 702 | A | LYS | 790 | B |
| ARG | 357 | A | PRO | 230 | B |
| VAL | 1094 | A | TYR | 904 | B |
| ASN | 969 | A | GLN | 755 | B |
| GLY | 669 | A | MET | 869 | B |
| MET | 900 | A | ALA | 1078 | C |
| TYR | 917 | A | PRO | 1079 | C |
| ALA | 893 | A | VAL | 705 | C |
| PHE | 1089 | A | TYR | 917 | B |
| MET | 869 | A | THR | 696 | C |
| TYR | 707 | A | ALA | 899 | B |
| GLN | 314 | A | SER | 735 | B |
| LYS | 558 | A | PHE | 43 | B |
| ALA | 879 | A | VAL | 705 | C |
| ARG | 765 | A | THR | 961 | C |
| TRP | 886 | A | TYR | 1047 | C |
| PRO | 863 | A | ALA | 668 | C |
| ILE | 896 | A | ALA | 713 | C |
| LEU | 984 | A | SER | 383 | C |
| VAL | 1094 | A | ILE | 896 | B |
| PHE | 562 | A | TYR | 38 | B |
| LYS | 790 | A | ASN | 703 | C |
| ASP | 1041 | A | ALA | 890 | B |
| GLN | 895 | A | ALA | 706 | C |
| SER | 758 | A | GLN | 965 | C |
| PRO | 1079 | A | MET | 900 | B |
| PRO | 897 | A | SER | 708 | C |
| VAL | 705 | A | LEU | 894 | B |
| ARG | 983 | A | VAL | 382 | C |
| SER | 704 | A | LYS | 790 | B |
| ALA | 668 | A | PRO | 862 | B |
| TYR | 873 | A | SER | 698 | C |
| GLY | 1046 | A | GLY | 889 | B |
| GLN | 913 | A | PHE | 1089 | C |
| ALA | 701 | A | ILE | 788 | B |
| PHE | 759 | A | GLY | 999 | C |
| SER | 967 | A | ASP | 571 | C |
| THR | 998 | A | GLN | 1002 | C |
| TYR | 756 | A | GLN | 965 | C |
| VAL | 987 | A | GLY | 413 | B |
| ARG | 355 | A | ASP | 198 | B |
| PHE | 855 | A | ASP | 568 | C |
| GLU | 918 | A | VAL | 1129 | C |
| ILE | 468 | A | THR | 114 | B |
| PRO | 862 | A | ILE | 666 | C |
| PHE | 562 | A | LYS | 41 | B |
| GLN | 913 | A | PRO | 1090 | C |
| TYR | 917 | A | PHE | 1089 | C |
| PRO | 665 | A | LEU | 864 | B |
| ILE | 670 | A | LEU | 864 | B |
| ALA | 766 | A | GLN | 1010 | C |
| SER | 708 | A | PRO | 897 | B |
| LEU | 864 | A | GLY | 667 | C |
| TYR | 707 | A | PHE | 898 | B |
| ILE | 1013 | A | ALA | 1016 | B |
| ALA | 890 | A | TYR | 1067 | C |
| ARG | 466 | A | GLY | 232 | B |
| GLY | 889 | A | LYS | 1045 | C |
| ARG | 408 | A | SER | 375 | B |
| MET | 740 | A | ARG | 319 | C |
| ASP | 568 | A | LYS | 854 | B |
| VAL | 1040 | A | GLU | 1031 | B |
| ILE | 788 | A | LEU | 699 | C |
| MET | 697 | A | LEU | 864 | B |
| LEU | 1001 | A | GLN | 1002 | C |
| VAL | 1008 | A | GLN | 1010 | C |
| GLN | 787 | A | ASN | 703 | C |
| ARG | 44 | A | GLY | 566 | C |
| VAL | 976 | A | THR | 572 | C |
| SER | 758 | A | THR | 961 | C |
| ASP | 1041 | A | GLY | 889 | B |
| ILE | 468 | A | LYS | 113 | B |
| GLN | 563 | A | GLY | 283 | B |
| PHE | 43 | A | LYS | 558 | C |
| PRO | 1069 | A | GLY | 891 | B |
| ARG | 319 | A | ASP | 745 | B |
| GLN | 762 | A | GLN | 965 | C |
| PRO | 897 | A | ASN | 709 | C |
| ALA | 1080 | A | TYR | 917 | B |
| ASP | 745 | A | THR | 549 | C |
| GLY | 971 | A | GLN | 755 | B |
| PRO | 863 | A | GLY | 669 | C |
| ASP | 568 | A | PHE | 855 | B |
| ARG | 1039 | A | GLU | 1031 | B |
| LYS | 964 | A | ALA | 570 | C |
| PHE | 562 | A | PRO | 225 | B |
| TYR | 904 | A | VAL | 1094 | C |
| ARG | 567 | A | VAL | 42 | B |
| VAL | 976 | A | ASP | 571 | C |
| GLN | 755 | A | SER | 968 | C |
| CYS | 379 | A | GLU | 988 | B |
| GLY | 757 | A | PHE | 970 | C |
| PHE | 565 | A | VAL | 42 | B |
| ASN | 914 | A | SER | 1123 | C |
| LYS | 41 | A | GLN | 564 | C |
| LYS | 558 | A | ASN | 282 | B |
| GLU | 281 | A | ASN | 556 | C |
| MET | 869 | A | MET | 697 | C |
| LYS | 557 | A | PHE | 43 | B |
| ARG | 983 | A | LEU | 390 | C |
| PHE | 970 | A | PHE | 759 | B |
| TYR | 789 | A | ASN | 703 | C |
| LEU | 699 | A | LYS | 786 | B |
| VAL | 382 | A | LEU | 984 | B |
| SER | 982 | A | LYS | 386 | C |
| LYS | 41 | A | PRO | 521 | C |
| SER | 45 | A | LYS | 557 | C |
| GLU | 1072 | A | ALA | 892 | B |
| VAL | 963 | A | ILE | 569 | C |
| ILE | 468 | A | GLN | 115 | B |
| PRO | 897 | A | TYR | 707 | C |
| GLY | 700 | A | ILE | 788 | B |
| SER | 1123 | A | GLU | 918 | B |
| THR | 887 | A | TYR | 1047 | C |
| ILE | 896 | A | ILE | 712 | C |
| MET | 869 | A | GLY | 669 | C |
| ALA | 520 | A | LYS | 41 | B |
| PHE | 1121 | A | ASN | 914 | B |
| TYR | 756 | A | SER | 968 | C |
| ASN | 907 | A | ARG | 1091 | C |
| LEU | 699 | A | ILE | 870 | B |
| ASP | 40 | A | GLN | 563 | C |
| LEU | 864 | A | CYS | 662 | C |
| MET | 740 | A | PHE | 592 | C |
| ARG | 983 | A | THR | 430 | C |
| TYR | 873 | A | GLY | 700 | C |
| SER | 383 | A | ASP | 985 | B |
| TYR | 38 | A | PHE | 562 | C |
| ARG | 355 | A | TYR | 200 | B |
| GLN | 563 | A | PHE | 43 | B |
| ALA | 575 | A | PHE | 43 | B |
| SER | 1123 | A | ASN | 914 | B |
| ARG | 1039 | A | THR | 1027 | B |
| SER | 383 | A | GLU | 988 | B |
| LYS | 386 | A | SER | 982 | B |
| ASN | 703 | A | TYR | 789 | B |
| THR | 1009 | A | ILE | 1013 | C |
| GLY | 667 | A | LEU | 864 | B |
| ALA | 893 | A | ALA | 713 | C |
| ASP | 745 | A | ARG | 319 | C |
| THR | 859 | A | GLN | 314 | C |
| GLN | 1002 | A | TYR | 756 | B |
| CYS | 590 | A | MET | 740 | B |
| GLY | 889 | A | VAL | 1040 | C |
| LYS | 986 | A | ASP | 427 | B |
| PHE | 464 | A | GLY | 199 | B |
| ILE | 896 | A | SER | 708 | C |
| ASP | 737 | A | ASN | 317 | C |
| ALA | 713 | A | GLN | 895 | B |
| GLN | 314 | A | THR | 768 | B |
| PRO | 862 | A | GLY | 667 | C |
| LEU | 699 | A | TYR | 873 | B |
| LYS | 790 | A | SER | 704 | C |
| LYS | 1038 | A | LYS | 1038 | C |
| THR | 1006 | A | GLN | 762 | B |
| GLN | 965 | A | GLY | 757 | B |
| GLN | 784 | A | ASP | 1041 | C |
| PHE | 562 | A | ASP | 40 | B |
| TYR | 904 | A | ARG | 1107 | C |
| PHE | 759 | A | SER | 1003 | C |
| ALA | 570 | A | SER | 975 | B |
| VAL | 42 | A | PHE | 562 | C |
| GLN | 787 | A | ALA | 701 | C |
| SER | 968 | A | SER | 758 | B |
| LEU | 894 | A | ALA | 713 | C |
| GLN | 1002 | A | THR | 998 | B |
| PRO | 426 | A | ASP | 198 | B |
| VAL | 987 | A | PRO | 412 | B |
| VAL | 1040 | A | SER | 1030 | B |
| GLU | 1144 | A | LEU | 1145 | C |
| GLN | 563 | A | ASP | 40 | B |
| LYS | 1038 | A | LYS | 1038 | B |
| TYR | 756 | A | GLY | 971 | C |
| TYR | 789 | A | SER | 704 | C |
| THR | 961 | A | SER | 758 | B |
| GLY | 566 | A | VAL | 42 | B |
| TYR | 789 | A | GLU | 702 | C |
| TRP | 886 | A | ARG | 1107 | C |
| VAL | 42 | A | PHE | 565 | C |
| VAL | 705 | A | SER | 884 | B |
| LEU | 865 | A | GLY | 669 | C |
| ASN | 282 | A | LYS | 558 | C |
| ILE | 794 | A | TYR | 707 | C |
| PRO | 897 | A | SER | 711 | C |
| ASN | 978 | A | THR | 547 | C |
| GLU | 1017 | A | ARG | 1019 | B |
| GLN | 762 | A | THR | 1006 | C |
| LEU | 1145 | A | GLU | 1144 | B |
| ARG | 319 | A | ASP | 737 | B |
| GLY | 593 | A | THR | 859 | B |
| LEU | 865 | A | ALA | 668 | C |
| LEU | 546 | A | ASP | 979 | B |
| GLY | 283 | A | LEU | 560 | C |
| ASN | 856 | A | THR | 572 | C |
| LYS | 1038 | A | GLN | 1036 | B |
| ARG | 355 | A | PRO | 230 | B |
| TYR | 200 | A | GLU | 516 | C |
| ASN | 709 | A | PRO | 897 | B |
| THR | 866 | A | GLY | 669 | C |
| MET | 740 | A | ASN | 317 | C |
| LEU | 966 | A | ALA | 570 | C |
| ARG | 466 | A | ASN | 234 | B |
| GLY | 757 | A | GLN | 965 | C |
| LEU | 1145 | A | LEU | 1145 | C |
| LEU | 981 | A | THR | 547 | C |
| HIS | 519 | A | VAL | 42 | B |
| GLU | 224 | A | PHE | 562 | C |
| THR | 547 | A | SER | 982 | B |
| GLN | 755 | A | ASN | 969 | C |
| ASN | 856 | A | ASP | 568 | C |
| GLU | 702 | A | TYR | 789 | B |
| TYR | 707 | A | ILE | 794 | B |
| TRP | 886 | A | VAL | 1040 | C |
| GLN | 1010 | A | LEU | 1012 | B |
| ASN | 907 | A | ARG | 1107 | C |
| PHE | 855 | A | PHE | 592 | C |
| VAL | 42 | A | GLN | 564 | C |
| LEU | 864 | A | ALA | 668 | C |
| GLY | 1093 | A | TYR | 904 | B |
| ASN | 978 | A | GLY | 548 | C |
| ILE | 468 | A | GLU | 132 | B |
| GLN | 965 | A | SER | 758 | B |
| GLY | 891 | A | PRO | 1069 | C |
| VAL | 42 | A | ARG | 567 | C |
| ASN | 1074 | A | GLN | 895 | B |
| GLU | 1072 | A | ALA | 893 | B |
| HIS | 519 | A | LYS | 41 | B |
| VAL | 1094 | A | MET | 900 | B |
| VAL | 705 | A | LYS | 790 | B |
| ILE | 973 | A | GLY | 381 | C |
| TYR | 707 | A | THR | 883 | B |
| MET | 869 | A | ALA | 668 | C |
| GLN | 895 | A | SER | 711 | C |
| VAL | 1040 | A | GLY | 1035 | B |
| LEU | 560 | A | GLU | 224 | B |
| GLN | 913 | A | PRO | 1079 | C |
| TYR | 369 | A | THR | 415 | C |
| LEU | 864 | A | MET | 697 | C |
| ALA | 372 | A | TYR | 505 | C |
| VAL | 1129 | A | TYR | 917 | B |
| GLY | 413 | A | VAL | 987 | C |
| SER | 514 | A | TYR | 200 | B |
| GLN | 563 | A | TYR | 38 | B |
| SER | 982 | A | LEU | 546 | C |
| VAL | 42 | A | GLY | 566 | C |
| ARG | 44 | A | ASP | 571 | C |
| SER | 968 | A | TYR | 756 | B |
| PHE | 759 | A | PHE | 970 | C |
| ARG | 44 | A | ARG | 567 | C |
| GLY | 566 | A | ARG | 44 | B |
| TYR | 200 | A | ASN | 394 | C |
| MET | 697 | A | TYR | 873 | B |
| ALA | 647 | A | PRO | 862 | B |
| SER | 982 | A | THR | 547 | C |
| GLN | 613 | A | THR | 859 | B |
| LEU | 699 | A | VAL | 785 | B |
| GLU | 702 | A | ILE | 788 | B |
| LEU | 518 | A | LYS | 41 | B |
| TYR | 1047 | A | ALA | 890 | B |
| GLN | 762 | A | GLN | 1010 | C |
| SER | 982 | A | LEU | 390 | C |
| PRO | 1069 | A | ALA | 890 | B |
| ALA | 570 | A | VAL | 963 | B |
| GLY | 667 | A | PRO | 863 | B |
| LEU | 699 | A | MET | 869 | B |
| ASN | 960 | A | ILE | 569 | C |
| LYS | 41 | A | ALA | 520 | C |
| THR | 961 | A | GLN | 762 | B |
| VAL | 42 | A | HIS | 519 | C |
| GLN | 913 | A | ARG | 1091 | C |
| LYS | 921 | A | ILE | 1130 | C |
| PRO | 1079 | A | TYR | 917 | B |
| TYR | 396 | A | TYR | 200 | B |
| THR | 739 | A | ARG | 319 | C |
| PHE | 970 | A | GLY | 757 | B |
| ALA | 890 | A | TYR | 1047 | C |
| GLN | 920 | A | ILE | 1130 | C |
| LYS | 386 | A | ASP | 985 | B |
| ASN | 1108 | A | TRP | 886 | B |
| PRO | 715 | A | LEU | 894 | B |
| VAL | 987 | A | ASP | 427 | B |
| ALA | 372 | A | LYS | 417 | C |
| SER | 758 | A | SER | 968 | C |
| GLN | 1002 | A | GLN | 1002 | C |
| TYR | 707 | A | PRO | 897 | B |
| ARG | 983 | A | GLY | 381 | C |
| PHE | 43 | A | LYS | 557 | C |
| GLU | 516 | A | ASP | 228 | B |
| ASP | 1041 | A | SER | 1030 | B |
| ARG | 319 | A | THR | 739 | B |
| PRO | 1079 | A | GLN | 913 | B |
| LEU | 560 | A | PHE | 43 | B |
| TYR | 789 | A | ALA | 701 | C |
| ARG | 1039 | A | ARG | 1039 | C |
| ILE | 1013 | A | THR | 1009 | B |
| GLY | 700 | A | LYS | 786 | B |
| LEU | 517 | A | ARG | 983 | B |
| ARG | 1107 | A | TYR | 904 | B |
| PHE | 562 | A | GLU | 224 | B |
| GLY | 381 | A | ILE | 973 | B |
| SER | 371 | A | LYS | 417 | C |
| ILE | 788 | A | GLY | 700 | C |
| PRO | 463 | A | ASP | 198 | B |
| ALA | 668 | A | THR | 866 | B |
| VAL | 976 | A | LEU | 546 | C |
| ALA | 570 | A | SER | 967 | B |
| LYS | 964 | A | SER | 758 | B |
| LEU | 864 | A | CYS | 671 | C |
| TYR | 707 | A | ILE | 882 | B |
| ILE | 788 | A | GLU | 702 | C |
| LEU | 894 | A | PRO | 1069 | C |
| GLU | 988 | A | SER | 383 | C |
| TYR | 38 | A | GLN | 563 | C |
| SER | 1003 | A | GLN | 762 | B |
| GLY | 669 | A | LEU | 864 | B |
| GLU | 988 | A | CYS | 379 | C |
| GLY | 700 | A | GLN | 787 | B |
| ILE | 714 | A | LEU | 894 | B |
| PHE | 168 | A | ARG | 357 | C |
| GLY | 757 | A | SER | 968 | C |
| LYS | 854 | A | ASP | 568 | C |
| TYR | 396 | A | PRO | 230 | B |
| GLU | 224 | A | LEU | 560 | C |
| CYS | 662 | A | LEU | 864 | B |
| SER | 1003 | A | PHE | 759 | B |
| VAL | 382 | A | GLU | 988 | B |
| SER | 884 | A | VAL | 705 | C |
| TYR | 873 | A | LEU | 699 | C |
| MET | 740 | A | PRO | 589 | C |
| GLU | 918 | A | SER | 1123 | C |
| GLN | 563 | A | VAL | 42 | B |
| GLU | 1111 | A | SER | 1123 | C |
| SER | 708 | A | ASP | 796 | B |
| ALA | 899 | A | TYR | 707 | C |
| THR | 696 | A | MET | 869 | B |
| GLN | 1010 | A | ALA | 766 | B |
| ILE | 1013 | A | ILE | 1013 | C |
| PHE | 43 | A | LEU | 560 | C |
| TYR | 707 | A | PHE | 797 | B |
| MET | 697 | A | GLN | 779 | B |
| ILE | 896 | A | SER | 711 | C |
| LEU | 1012 | A | ILE | 1013 | C |
| PRO | 862 | A | ALA | 668 | C |
| GLN | 965 | A | TYR | 756 | B |
| VAL | 1129 | A | GLU | 918 | B |
| PHE | 759 | A | GLN | 1002 | C |
| ILE | 1013 | A | ILE | 1013 | B |
| LYS | 378 | A | GLU | 988 | B |
| ASP | 228 | A | ALA | 520 | C |
| LEU | 894 | A | ILE | 714 | C |
| GLN | 787 | A | GLY | 700 | C |
| LEU | 560 | A | ASN | 282 | B |
| PHE | 1089 | A | GLU | 918 | B |
| ALA | 570 | A | LEU | 966 | B |
| LEU | 560 | A | THR | 284 | B |
| ALA | 701 | A | TYR | 789 | B |
| GLN | 787 | A | GLU | 702 | C |
| LEU | 864 | A | ILE | 670 | C |
| VAL | 705 | A | TYR | 789 | B |
| GLU | 1031 | A | PHE | 1042 | C |
| ASP | 985 | A | THR | 385 | C |
| PHE | 464 | A | TYR | 200 | B |
| ASN | 709 | A | ASP | 796 | B |
| LYS | 776 | A | LYS | 947 | C |
| ALA | 1016 | A | ARG | 1019 | B |
| ASP | 568 | A | ASN | 856 | B |
| ILE | 666 | A | PRO | 862 | B |
| ALA | 570 | A | LYS | 964 | B |
| LEU | 984 | A | VAL | 382 | C |
| SER | 708 | A | ILE | 896 | B |
| ARG | 1039 | A | ARG | 1039 | B |
| LYS | 786 | A | GLY | 700 | C |
| ARG | 1091 | A | GLN | 913 | B |
| GLU | 1031 | A | ARG | 1039 | C |
| SER | 711 | A | ILE | 896 | B |
| VAL | 1040 | A | GLY | 889 | B |
| PRO | 792 | A | TYR | 707 | C |
| LYS | 1045 | A | GLY | 889 | B |
| ALA | 668 | A | LEU | 864 | B |
| LYS | 41 | A | PHE | 562 | C |
| PHE | 43 | A | PHE | 559 | C |
| GLN | 563 | A | ARG | 44 | B |
| ILE | 1130 | A | GLY | 798 | B |
| PRO | 862 | A | ARG | 646 | C |
| LEU | 864 | A | ILE | 666 | C |
| ILE | 569 | A | LYS | 964 | B |
| GLY | 857 | A | PHE | 592 | C |
| ARG | 1107 | A | ASN | 907 | B |
| ASN | 978 | A | LEU | 546 | C |
| LEU | 1012 | A | GLN | 1010 | C |
| THR | 385 | A | THR | 415 | C |
| TYR | 707 | A | GLY | 798 | B |
| LYS | 786 | A | LYS | 1045 | C |
| PRO | 1069 | A | ALA | 892 | B |
| GLU | 868 | A | LEU | 699 | C |
| GLU | 918 | A | GLY | 1124 | C |
| GLY | 381 | A | LEU | 984 | B |
| GLN | 563 | A | TYR | 279 | B |
| LEU | 981 | A | LYS | 386 | C |
| ASP | 1041 | A | LEU | 1034 | B |

**Table S6.** The list of the inter-protomer contacts of the protomer A in the structure of SARS-CoV-2 S-RRAR/G614 in the open state (pdb id 7KDJ).

| **S protein residue name** | **S protein residue number** | **S protein chain** | **S protein residue name** | **S protein residue number** | **S protein chain** |
| --- | --- | --- | --- | --- | --- |
| HIS | 519 | A | ASP | 228 | B |
| GLN | 1002 | A | TYR | 756 | B |
| ILE | 1130 | A | TYR | 917 | B |
| LEU | 390 | A | ARG | 983 | B |
| GLN | 564 | A | VAL | 42 | B |
| GLU | 1092 | A | GLN | 913 | B |
| GLY | 1046 | A | ALA | 890 | B |
| ARG | 567 | A | PHE | 43 | B |
| TYR | 707 | A | ILE | 882 | B |
| TYR | 789 | A | VAL | 705 | C |
| GLU | 516 | A | PRO | 230 | B |
| ILE | 1013 | A | LEU | 1012 | B |
| GLN | 314 | A | ASP | 737 | B |
| GLN | 913 | A | ARG | 1107 | C |
| LYS | 113 | A | SER | 469 | C |
| ALA | 520 | A | ASP | 228 | B |
| HIS | 519 | A | LYS | 41 | B |
| SER | 708 | A | ILE | 896 | B |
| SER | 383 | A | ARG | 983 | B |
| SER | 1030 | A | VAL | 1040 | C |
| ALA | 701 | A | LYS | 786 | B |
| ARG | 1107 | A | TRP | 886 | B |
| GLN | 1005 | A | THR | 1006 | C |
| TYR | 1067 | A | ALA | 890 | B |
| ARG | 1091 | A | ASN | 907 | B |
| ASN | 317 | A | MET | 740 | B |
| PHE | 759 | A | GLN | 965 | C |
| THR | 167 | A | ASN | 354 | C |
| ILE | 1130 | A | GLN | 920 | B |
| LEU | 864 | A | CYS | 671 | C |
| LEU | 894 | A | PRO | 715 | C |
| LEU | 1141 | A | GLU | 1144 | B |
| ILE | 973 | A | GLY | 381 | C |
| GLY | 889 | A | ASP | 1041 | C |
| LEU | 864 | A | ILE | 670 | C |
| VAL | 963 | A | ALA | 570 | C |
| LEU | 1012 | A | GLN | 1010 | C |
| LYS | 921 | A | ILE | 1130 | C |
| PRO | 230 | A | TYR | 396 | C |
| GLU | 1017 | A | GLU | 773 | B |
| SER | 45 | A | LYS | 557 | C |
| THR | 1009 | A | ILE | 1013 | C |
| ALA | 892 | A | GLU | 1072 | C |
| GLY | 667 | A | PRO | 863 | B |
| PRO | 862 | A | GLY | 667 | C |
| GLU | 1031 | A | PHE | 1042 | C |
| GLN | 895 | A | ALA | 706 | C |
| VAL | 705 | A | ALA | 879 | B |
| GLY | 1046 | A | GLY | 889 | B |
| VAL | 705 | A | LYS | 790 | B |
| TRP | 886 | A | ARG | 1107 | C |
| PHE | 592 | A | MET | 740 | B |
| LEU | 984 | A | SER | 383 | C |
| PRO | 897 | A | TYR | 707 | C |
| LYS | 786 | A | LEU | 699 | C |
| PHE | 43 | A | GLN | 563 | C |
| ASP | 1041 | A | ALA | 890 | B |
| TYR | 873 | A | SER | 698 | C |
| GLN | 613 | A | THR | 859 | B |
| TYR | 38 | A | PHE | 562 | C |
| TYR | 904 | A | ARG | 1107 | C |
| ASN | 709 | A | ASP | 796 | B |
| ILE | 1013 | A | THR | 1009 | B |
| TYR | 707 | A | ILE | 794 | B |
| THR | 549 | A | ASP | 745 | B |
| ASN | 710 | A | PRO | 897 | B |
| ASP | 979 | A | LEU | 546 | C |
| VAL | 705 | A | THR | 883 | B |
| GLU | 224 | A | PRO | 561 | C |
| TYR | 789 | A | SER | 704 | C |
| TYR | 1047 | A | TRP | 886 | B |
| ASN | 978 | A | GLY | 545 | C |
| THR | 859 | A | PHE | 592 | C |
| MET | 697 | A | LEU | 864 | B |
| THR | 1006 | A | GLN | 1005 | B |
| LEU | 864 | A | GLY | 667 | C |
| GLN | 314 | A | THR | 768 | B |
| GLY | 232 | A | GLU | 465 | C |
| ILE | 233 | A | ARG | 466 | C |
| LEU | 894 | A | ALA | 713 | C |
| LYS | 790 | A | SER | 704 | C |
| GLY | 891 | A | PRO | 1069 | C |
| TYR | 369 | A | ARG | 408 | C |
| ILE | 896 | A | ALA | 713 | C |
| MET | 740 | A | PRO | 589 | C |
| ASN | 856 | A | ALA | 570 | C |
| PRO | 225 | A | PHE | 562 | C |
| ASP | 571 | A | SER | 967 | B |
| VAL | 1040 | A | GLN | 1036 | B |
| SER | 884 | A | VAL | 705 | C |
| LYS | 558 | A | GLU | 281 | B |
| ASP | 979 | A | THR | 547 | C |
| PHE | 759 | A | GLN | 1002 | C |
| GLN | 957 | A | ARG | 765 | B |
| LEU | 1145 | A | LEU | 1141 | B |
| ARG | 983 | A | LYS | 386 | C |
| ALA | 890 | A | VAL | 1040 | C |
| ILE | 670 | A | LEU | 864 | B |
| PHE | 592 | A | LEU | 858 | B |
| SER | 758 | A | GLN | 965 | C |
| VAL | 785 | A | LEU | 699 | C |
| SER | 982 | A | LEU | 390 | C |
| GLN | 1036 | A | VAL | 1040 | C |
| GLN | 613 | A | LEU | 861 | B |
| VAL | 1068 | A | ALA | 890 | B |
| SER | 975 | A | ASP | 571 | C |
| GLU | 918 | A | VAL | 1128 | C |
| GLY | 889 | A | VAL | 1040 | C |
| THR | 167 | A | ARG | 466 | C |
| LEU | 864 | A | PRO | 665 | C |
| LEU | 1145 | A | GLU | 1144 | B |
| THR | 887 | A | TYR | 1047 | C |
| ARG | 44 | A | GLN | 563 | C |
| ASN | 907 | A | ARG | 1107 | C |
| SER | 968 | A | SER | 758 | B |
| THR | 1027 | A | ARG | 1039 | C |
| TYR | 756 | A | GLY | 971 | C |
| TYR | 707 | A | ASP | 796 | B |
| GLY | 757 | A | PHE | 970 | C |
| GLY | 1093 | A | GLN | 913 | B |
| ILE | 712 | A | MET | 900 | B |
| PRO | 863 | A | ALA | 668 | C |
| PHE | 1095 | A | MET | 900 | B |
| ALA | 1080 | A | TYR | 917 | B |
| PHE | 592 | A | GLY | 857 | B |
| ASP | 745 | A | ARG | 319 | C |
| ARG | 1039 | A | GLU | 1031 | B |
| THR | 572 | A | ASN | 856 | B |
| ALA | 570 | A | LYS | 964 | B |
| LEU | 894 | A | PRO | 1069 | C |
| ARG | 567 | A | ARG | 44 | B |
| TYR | 707 | A | ALA | 899 | B |
| GLU | 988 | A | SER | 383 | C |
| TYR | 707 | A | ILE | 896 | B |
| LYS | 790 | A | ASN | 703 | C |
| THR | 883 | A | VAL | 705 | C |
| ILE | 712 | A | GLN | 895 | B |
| ASP | 1041 | A | LEU | 1034 | B |
| TYR | 873 | A | MET | 697 | C |
| GLN | 787 | A | ALA | 701 | C |
| GLY | 700 | A | GLN | 787 | B |
| GLY | 798 | A | ILE | 1130 | C |
| PRO | 1069 | A | ALA | 892 | B |
| LYS | 947 | A | LYS | 776 | B |
| VAL | 705 | A | SER | 884 | B |
| ASP | 994 | A | ARG | 995 | C |
| ASN | 394 | A | PRO | 230 | B |
| GLN | 1002 | A | LEU | 1001 | B |
| VAL | 1094 | A | MET | 900 | B |
| GLY | 1035 | A | VAL | 1040 | C |
| VAL | 705 | A | LEU | 894 | B |
| SER | 698 | A | TYR | 873 | B |
| TYR | 917 | A | VAL | 1128 | C |
| LEU | 560 | A | GLY | 283 | B |
| MET | 869 | A | MET | 697 | C |
| SER | 1123 | A | GLU | 1111 | B |
| ILE | 788 | A | ASN | 703 | C |
| ALA | 713 | A | ILE | 896 | B |
| THR | 385 | A | THR | 415 | C |
| SER | 967 | A | ILE | 569 | C |
| GLN | 895 | A | ASN | 1074 | C |
| ILE | 666 | A | PRO | 862 | B |
| GLN | 762 | A | SER | 1003 | C |
| PRO | 792 | A | TYR | 707 | C |
| GLN | 787 | A | ASN | 703 | C |
| PHE | 1121 | A | THR | 912 | B |
| TYR | 904 | A | ILE | 712 | C |
| ASP | 985 | A | LYS | 386 | C |
| GLN | 314 | A | LEU | 861 | B |
| GLN | 314 | A | ASN | 764 | B |
| ALA | 713 | A | LEU | 894 | B |
| ASN | 978 | A | THR | 547 | C |
| TYR | 756 | A | SER | 968 | C |
| THR | 791 | A | VAL | 705 | C |
| PHE | 1089 | A | GLU | 918 | B |
| LYS | 790 | A | VAL | 705 | C |
| LEU | 560 | A | TYR | 38 | B |
| PRO | 589 | A | ASN | 856 | B |
| THR | 768 | A | GLN | 314 | C |
| PRO | 862 | A | ALA | 668 | C |
| THR | 791 | A | SER | 704 | C |
| PHE | 592 | A | PHE | 855 | B |
| GLU | 516 | A | TYR | 200 | B |
| TYR | 904 | A | VAL | 1094 | C |
| VAL | 963 | A | ILE | 569 | C |
| GLN | 563 | A | PHE | 43 | B |
| TYR | 917 | A | ILE | 1130 | C |
| ASN | 709 | A | PRO | 897 | B |
| PHE | 759 | A | THR | 1006 | C |
| TYR | 1007 | A | GLN | 762 | B |
| TYR | 917 | A | VAL | 1129 | C |
| LEU | 1145 | A | LEU | 1145 | C |
| GLU | 224 | A | PHE | 562 | C |
| SER | 704 | A | ILE | 788 | B |
| GLY | 1093 | A | TYR | 904 | B |
| PHE | 43 | A | LYS | 558 | C |
| PRO | 230 | A | ARG | 357 | C |
| LEU | 981 | A | LYS | 386 | C |
| ASN | 907 | A | ARG | 1091 | C |
| VAL | 1008 | A | GLN | 1010 | C |
| GLY | 566 | A | ARG | 44 | B |
| SER | 711 | A | ILE | 896 | B |
| THR | 547 | A | SER | 982 | B |
| LEU | 894 | A | VAL | 705 | C |
| GLN | 1113 | A | VAL | 1122 | C |
| TYR | 789 | A | ALA | 701 | C |
| GLN | 913 | A | PRO | 1090 | C |
| PHE | 797 | A | TYR | 707 | C |
| PRO | 863 | A | GLY | 669 | C |
| ARG | 983 | A | LEU | 390 | C |
| PRO | 412 | A | VAL | 987 | C |
| MET | 900 | A | ALA | 1078 | C |
| ILE | 1013 | A | ALA | 1016 | B |
| ILE | 666 | A | LEU | 864 | B |
| SER | 982 | A | LYS | 386 | C |
| GLY | 566 | A | VAL | 42 | B |
| ILE | 714 | A | LEU | 894 | B |
| THR | 1006 | A | PHE | 759 | B |
| LEU | 699 | A | GLN | 872 | B |
| GLN | 1002 | A | THR | 998 | B |
| LEU | 699 | A | ILE | 788 | B |
| GLN | 1010 | A | VAL | 1008 | B |
| THR | 961 | A | SER | 758 | B |
| GLY | 413 | A | ASP | 985 | C |
| GLY | 891 | A | VAL | 1068 | C |
| GLN | 965 | A | TYR | 756 | B |
| ASP | 198 | A | PRO | 426 | C |
| GLN | 1010 | A | ALA | 766 | B |
| GLY | 283 | A | LEU | 560 | C |
| LEU | 1141 | A | LEU | 1141 | C |
| MET | 740 | A | CYS | 590 | C |
| LEU | 864 | A | CYS | 662 | C |
| VAL | 705 | A | PRO | 792 | B |
| PHE | 759 | A | GLY | 999 | C |
| TYR | 396 | A | TYR | 200 | B |
| ASP | 40 | A | PHE | 562 | C |
| LYS | 386 | A | ASP | 985 | B |
| LEU | 1141 | A | LEU | 1145 | C |
| GLY | 199 | A | PHE | 464 | C |
| GLY | 700 | A | LYS | 786 | B |
| GLY | 669 | A | LEU | 865 | B |
| PHE | 1089 | A | GLN | 913 | B |
| TYR | 396 | A | PRO | 230 | B |
| PHE | 970 | A | GLY | 757 | B |
| PHE | 168 | A | ARG | 357 | C |
| PHE | 855 | A | ASP | 568 | C |
| GLN | 1002 | A | GLN | 1002 | C |
| ILE | 569 | A | VAL | 47 | B |
| GLN | 965 | A | GLN | 762 | B |
| ASP | 1041 | A | SER | 1030 | B |
| GLU | 918 | A | GLY | 1124 | C |
| SER | 968 | A | TYR | 756 | B |
| ILE | 1130 | A | LYS | 921 | B |
| TYR | 707 | A | GLY | 798 | B |
| THR | 866 | A | GLY | 669 | C |
| GLN | 1002 | A | PHE | 759 | B |
| TYR | 917 | A | PRO | 1079 | C |
| GLN | 563 | A | TYR | 279 | B |
| ARG | 646 | A | PRO | 862 | B |
| GLU | 702 | A | LYS | 790 | B |
| PHE | 562 | A | ASP | 40 | B |
| TYR | 904 | A | GLY | 1093 | C |
| SER | 968 | A | GLN | 755 | B |
| PRO | 715 | A | LEU | 894 | B |
| ASP | 198 | A | PRO | 463 | C |
| PRO | 230 | A | ARG | 355 | C |
| PRO | 1090 | A | ASN | 907 | B |
| VAL | 705 | A | GLN | 895 | B |
| VAL | 1128 | A | GLU | 918 | B |
| GLU | 1144 | A | LEU | 1145 | C |
| ASN | 914 | A | PHE | 1121 | C |
| GLY | 669 | A | MET | 869 | B |
| VAL | 1040 | A | GLY | 889 | B |
| ASN | 703 | A | ALA | 893 | B |
| PHE | 592 | A | THR | 859 | B |
| GLN | 895 | A | ILE | 712 | C |
| LYS | 41 | A | GLN | 563 | C |
| LEU | 699 | A | TYR | 873 | B |
| ASP | 40 | A | GLN | 563 | C |
| PHE | 1042 | A | SER | 1030 | B |
| THR | 572 | A | VAL | 976 | B |
| LEU | 699 | A | MET | 869 | B |
| PHE | 565 | A | PHE | 43 | B |
| SER | 735 | A | GLN | 314 | C |
| ARG | 355 | A | PRO | 230 | B |
| GLU | 1031 | A | VAL | 1040 | C |
| ALA | 520 | A | LYS | 41 | B |
| VAL | 382 | A | LEU | 984 | B |
| GLN | 895 | A | TYR | 707 | C |
| ALA | 1078 | A | MET | 900 | B |
| ILE | 712 | A | TYR | 904 | B |
| ALA | 879 | A | VAL | 705 | C |
| ALA | 668 | A | LEU | 864 | B |
| LEU | 517 | A | ARG | 983 | B |
| PHE | 562 | A | VAL | 42 | B |
| LYS | 386 | A | ARG | 983 | B |
| LEU | 864 | A | GLY | 669 | C |
| GLY | 232 | A | TRP | 353 | C |
| ARG | 1039 | A | THR | 1027 | B |
| ASN | 703 | A | GLN | 787 | B |
| ASP | 228 | A | ASN | 394 | C |
| THR | 1009 | A | THR | 1009 | C |
| LEU | 560 | A | ASN | 282 | B |
| GLN | 563 | A | LYS | 41 | B |
| PRO | 863 | A | GLY | 667 | C |
| GLN | 965 | A | GLY | 757 | B |
| PRO | 384 | A | ASP | 985 | B |
| LEU | 1034 | A | VAL | 1040 | C |
| ARG | 567 | A | VAL | 976 | B |
| ALA | 899 | A | TYR | 707 | C |
| GLN | 762 | A | GLN | 965 | C |
| ILE | 231 | A | ARG | 466 | C |
| ASN | 978 | A | GLY | 548 | C |
| ARG | 1019 | A | GLU | 1017 | C |
| PRO | 792 | A | VAL | 705 | C |
| LYS | 786 | A | GLY | 700 | C |
| PHE | 1121 | A | ASN | 914 | B |
| ASP | 979 | A | GLY | 545 | C |
| ALA | 1070 | A | ALA | 892 | B |
| MET | 869 | A | LEU | 699 | C |
| LYS | 386 | A | LEU | 984 | B |
| ASP | 571 | A | VAL | 976 | B |
| GLY | 1124 | A | GLU | 918 | B |
| PRO | 897 | A | SER | 708 | C |
| MET | 900 | A | VAL | 1094 | C |
| ASP | 228 | A | TYR | 396 | C |
| GLY | 891 | A | GLY | 1046 | C |
| THR | 547 | A | ASN | 978 | B |
| VAL | 1068 | A | GLY | 891 | B |
| GLN | 895 | A | ALA | 713 | C |
| GLY | 757 | A | GLN | 965 | C |
| MET | 900 | A | PHE | 1095 | C |
| LEU | 858 | A | PHE | 592 | C |
| LEU | 865 | A | MET | 697 | C |
| GLY | 857 | A | PHE | 592 | C |
| VAL | 705 | A | THR | 791 | B |
| THR | 1077 | A | PRO | 897 | B |
| ALA | 668 | A | THR | 866 | B |
| SER | 982 | A | THR | 547 | C |
| MET | 697 | A | MET | 869 | B |
| ILE | 1013 | A | ILE | 1013 | C |
| PHE | 970 | A | GLN | 755 | B |
| ARG | 44 | A | GLY | 566 | C |
| PHE | 562 | A | LYS | 41 | B |
| PRO | 862 | A | ALA | 647 | C |
| GLN | 762 | A | THR | 1006 | C |
| VAL | 1040 | A | TRP | 886 | B |
| ASP | 198 | A | PHE | 464 | C |
| ASN | 1108 | A | TRP | 886 | B |
| ARG | 983 | A | VAL | 382 | C |
| LEU | 966 | A | ALA | 570 | C |
| PHE | 970 | A | TYR | 756 | B |
| THR | 1009 | A | THR | 1009 | B |
| ARG | 995 | A | ASP | 994 | B |
| ILE | 896 | A | SER | 708 | C |
| PHE | 1089 | A | TYR | 917 | B |
| LEU | 546 | A | ASP | 979 | B |
| PRO | 1079 | A | TYR | 917 | B |
| SER | 1003 | A | PHE | 759 | B |
| MET | 740 | A | PHE | 592 | C |
| GLY | 413 | A | LYS | 986 | C |
| ALA | 701 | A | ILE | 788 | B |
| GLY | 667 | A | LEU | 864 | B |
| ARG | 567 | A | VAL | 42 | B |
| GLN | 755 | A | PHE | 970 | C |
| ILE | 666 | A | LYS | 733 | B |
| MET | 740 | A | ARG | 319 | C |
| GLY | 594 | A | THR | 859 | B |
| VAL | 705 | A | TYR | 789 | B |
| SER | 758 | A | LYS | 964 | C |
| GLY | 744 | A | ARG | 319 | C |
| SER | 982 | A | GLY | 545 | C |
| GLN | 563 | A | VAL | 42 | B |
| VAL | 976 | A | ASP | 571 | C |
| VAL | 382 | A | GLU | 988 | B |
| SER | 383 | A | ASP | 985 | B |
| GLN | 787 | A | GLY | 700 | C |
| HIS | 519 | A | ASP | 40 | B |
| GLU | 702 | A | TYR | 789 | B |
| ARG | 319 | A | MET | 740 | B |
| PRO | 1079 | A | GLN | 913 | B |
| GLN | 563 | A | ARG | 44 | B |
| LEU | 560 | A | THR | 284 | B |
| ASN | 282 | A | LEU | 560 | C |
| PHE | 565 | A | VAL | 42 | B |
| ASP | 796 | A | ASN | 709 | C |
| THR | 859 | A | GLY | 593 | C |
| ARG | 1039 | A | ARG | 1039 | B |
| TYR | 200 | A | PHE | 464 | C |
| PHE | 43 | A | LYS | 557 | C |
| LEU | 560 | A | PHE | 43 | B |
| ARG | 765 | A | THR | 961 | C |
| SER | 982 | A | ASP | 389 | C |
| GLN | 965 | A | PHE | 759 | B |
| TYR | 917 | A | PHE | 1089 | C |
| SER | 708 | A | PHE | 898 | B |
| GLU | 1144 | A | LEU | 1141 | C |
| ALA | 893 | A | GLU | 1072 | C |
| GLN | 563 | A | TYR | 38 | B |
| TYR | 707 | A | PHE | 898 | B |
| LYS | 386 | A | SER | 982 | B |
| ARG | 1039 | A | SER | 1037 | B |
| TYR | 707 | A | PRO | 792 | B |
| GLU | 918 | A | PHE | 1089 | C |
| LYS | 41 | A | GLN | 564 | C |
| PHE | 759 | A | SER | 1003 | C |
| ILE | 1013 | A | ILE | 1013 | B |
| TYR | 1047 | A | THR | 887 | B |
| LEU | 865 | A | ALA | 668 | C |
| LYS | 964 | A | ILE | 569 | C |
| GLN | 755 | A | ALA | 972 | C |
| VAL | 47 | A | ILE | 569 | C |
| GLN | 779 | A | MET | 697 | C |
| GLN | 895 | A | SER | 708 | C |
| ILE | 233 | A | PHE | 464 | C |
| LEU | 754 | A | SER | 968 | C |
| SER | 968 | A | GLY | 757 | B |
| SER | 383 | A | GLU | 988 | B |
| GLY | 381 | A | ILE | 973 | B |
| PHE | 759 | A | PHE | 970 | C |
| ALA | 668 | A | PRO | 862 | B |
| PHE | 562 | A | TYR | 38 | B |
| TYR | 369 | A | GLY | 416 | C |
| ASN | 914 | A | PHE | 1089 | C |
| ALA | 890 | A | VAL | 1068 | C |
| LYS | 558 | A | ASN | 282 | B |
| LEU | 1034 | A | ASP | 1041 | C |
| ARG | 319 | A | ASP | 745 | B |
| LEU | 864 | A | ILE | 666 | C |
| TYR | 1047 | A | ALA | 890 | B |
| TYR | 873 | A | GLY | 700 | C |
| GLN | 895 | A | SER | 711 | C |
| LEU | 1001 | A | GLN | 1002 | C |
| THR | 961 | A | ARG | 765 | B |
| PRO | 862 | A | ARG | 646 | C |
| GLY | 381 | A | ARG | 983 | B |
| GLY | 548 | A | ASN | 978 | B |
| GLY | 700 | A | ILE | 788 | B |
| SER | 982 | A | LEU | 546 | C |
| VAL | 382 | A | ARG | 983 | B |
| LEU | 699 | A | VAL | 785 | B |
| VAL | 705 | A | GLY | 880 | B |
| THR | 547 | A | ASP | 979 | B |
| GLN | 913 | A | PHE | 1121 | C |
| PRO | 897 | A | THR | 1077 | C |
| LYS | 1038 | A | LYS | 1038 | B |
| VAL | 976 | A | ARG | 567 | C |
| GLU | 773 | A | GLU | 1017 | C |
| SER | 1003 | A | GLN | 762 | B |
| CYS | 671 | A | LEU | 864 | B |
| GLN | 755 | A | SER | 968 | C |
| MET | 900 | A | THR | 1077 | C |
| GLN | 895 | A | VAL | 705 | C |
| PHE | 1089 | A | ASN | 914 | B |
| MET | 697 | A | GLN | 779 | B |
| LEU | 699 | A | LEU | 865 | B |
| LEU | 865 | A | GLY | 669 | C |
| THR | 859 | A | GLY | 594 | C |
| CYS | 379 | A | GLU | 988 | B |
| GLY | 669 | A | LEU | 864 | B |
| VAL | 1128 | A | TYR | 917 | B |
| THR | 912 | A | PHE | 1121 | C |
| ASP | 1118 | A | ARG | 1091 | C |
| ARG | 357 | A | ASP | 228 | B |
| GLY | 545 | A | ASN | 978 | B |
| ASN | 703 | A | ILE | 788 | B |
| PHE | 565 | A | LYS | 41 | B |
| ARG | 357 | A | LEU | 229 | B |
| GLU | 224 | A | LEU | 560 | C |
| VAL | 1129 | A | GLU | 918 | B |
| ALA | 668 | A | MET | 869 | B |
| ASP | 1041 | A | GLY | 889 | B |
| ASP | 796 | A | SER | 708 | C |
| ILE | 788 | A | GLU | 702 | C |
| GLN | 762 | A | GLN | 1010 | C |
| SER | 967 | A | ALA | 570 | C |
| ALA | 766 | A | GLN | 1010 | C |
| TRP | 886 | A | TYR | 1047 | C |
| PHE | 43 | A | LEU | 560 | C |
| SER | 708 | A | GLN | 895 | B |
| TYR | 873 | A | LEU | 699 | C |
| GLU | 1072 | A | ALA | 892 | B |
| PRO | 1069 | A | GLY | 891 | B |
| LYS | 1038 | A | LYS | 1038 | C |
| SER | 967 | A | ASP | 571 | C |
| ARG | 1091 | A | GLN | 913 | B |
| LYS | 41 | A | HIS | 519 | C |
| ASN | 282 | A | LYS | 558 | C |
| GLN | 872 | A | LEU | 699 | C |
| THR | 430 | A | ARG | 983 | B |
| ALA | 890 | A | TYR | 1047 | C |
| GLN | 913 | A | GLY | 1093 | C |
| ASP | 389 | A | SER | 982 | B |
| ASN | 856 | A | PRO | 589 | C |
| PHE | 43 | A | PHE | 565 | C |
| VAL | 1040 | A | LEU | 1034 | B |
| LEU | 1012 | A | ILE | 1013 | C |
| SER | 1123 | A | GLU | 918 | B |
| ASP | 979 | A | ARG | 567 | C |
| ALA | 890 | A | PRO | 1069 | C |
| GLY | 199 | A | PRO | 463 | C |
| ALA | 647 | A | PRO | 862 | B |
| ARG | 44 | A | ARG | 567 | C |
| ALA | 668 | A | LEU | 865 | B |
| ASP | 985 | A | THR | 385 | C |
| VAL | 42 | A | PHE | 565 | C |
| PHE | 43 | A | ARG | 567 | C |
| GLY | 413 | A | VAL | 987 | C |
| SER | 383 | A | LEU | 984 | B |
| TYR | 200 | A | ARG | 355 | C |
| LEU | 546 | A | VAL | 976 | B |
| PHE | 898 | A | TYR | 707 | C |
| ARG | 983 | A | SER | 383 | C |
| ASN | 317 | A | ASP | 737 | B |
| ASP | 198 | A | ASP | 428 | C |
| PHE | 43 | A | PHE | 559 | C |
| LYS | 786 | A | ALA | 701 | C |
| ILE | 788 | A | LEU | 699 | C |
| ALA | 1016 | A | ILE | 1013 | C |
| LEU | 864 | A | MET | 697 | C |
| GLU | 1031 | A | ARG | 1039 | C |
| PRO | 1140 | A | LEU | 1141 | C |
| ARG | 319 | A | ASP | 737 | B |
| PHE | 592 | A | ASN | 856 | B |
| GLY | 700 | A | TYR | 873 | B |
| PRO | 665 | A | LEU | 864 | B |
| ALA | 570 | A | VAL | 963 | B |
| GLN | 564 | A | LYS | 41 | B |
| LYS | 1045 | A | ALA | 890 | B |
| LYS | 964 | A | SER | 758 | B |
| ARG | 357 | A | PHE | 168 | B |
| LYS | 558 | A | PHE | 43 | B |
| GLY | 232 | A | PHE | 464 | C |
| PRO | 1079 | A | MET | 900 | B |
| ALA | 972 | A | GLN | 755 | B |
| TYR | 38 | A | LEU | 560 | C |
| GLN | 563 | A | GLY | 283 | B |
| ASN | 856 | A | PHE | 592 | C |
| ALA | 570 | A | LEU | 966 | B |
| TYR | 707 | A | PHE | 797 | B |
| LEU | 699 | A | LYS | 786 | B |
| GLY | 798 | A | TYR | 707 | C |
| PRO | 897 | A | ASN | 710 | C |
| GLY | 669 | A | PRO | 863 | B |
| ALA | 701 | A | GLN | 787 | B |
| THR | 866 | A | ALA | 668 | C |
| GLN | 965 | A | SER | 758 | B |
| ASN | 856 | A | ASP | 568 | C |
| SER | 704 | A | TYR | 789 | B |
| ASP | 985 | A | SER | 383 | C |
| VAL | 1040 | A | GLY | 1035 | B |
| TYR | 756 | A | ARG | 995 | C |
| GLY | 757 | A | SER | 968 | C |
| LYS | 557 | A | SER | 45 | B |
| GLY | 880 | A | VAL | 705 | C |
| LYS | 41 | A | PHE | 565 | C |
| ARG | 983 | A | THR | 430 | C |
| LYS | 41 | A | PHE | 562 | C |
| ARG | 1107 | A | ASN | 907 | B |
| LEU | 894 | A | GLU | 1072 | C |
| ILE | 788 | A | ALA | 701 | C |
| LYS | 790 | A | GLU | 702 | C |
| LYS | 964 | A | ALA | 570 | C |
| GLY | 1046 | A | GLY | 891 | B |
| ILE | 794 | A | TYR | 707 | C |
| TYR | 38 | A | GLN | 563 | C |
| PRO | 1069 | A | ALA | 890 | B |
| GLU | 1017 | A | ARG | 1019 | B |
| ALA | 701 | A | TYR | 789 | B |
| LYS | 386 | A | LEU | 981 | B |
| TYR | 756 | A | GLN | 965 | C |
| ILE | 1130 | A | GLY | 798 | B |
| ASN | 969 | A | GLN | 755 | B |
| ILE | 870 | A | LEU | 699 | C |
| GLY | 889 | A | GLY | 1046 | C |
| LYS | 1045 | A | GLY | 889 | B |
| GLU | 702 | A | GLN | 787 | B |
| VAL | 1122 | A | GLN | 1113 | B |
| ASN | 394 | A | TYR | 200 | B |
| GLN | 1002 | A | GLN | 1005 | B |
| ALA | 668 | A | PRO | 863 | B |
| PRO | 521 | A | LYS | 41 | B |
| PHE | 1042 | A | GLU | 1031 | B |
| LEU | 864 | A | ALA | 668 | C |
| GLY | 971 | A | GLN | 755 | B |
| TYR | 707 | A | PRO | 897 | B |
| GLY | 667 | A | PRO | 862 | B |
| THR | 1077 | A | MET | 900 | B |
| ARG | 357 | A | PRO | 230 | B |
| THR | 859 | A | GLN | 613 | C |
| ARG | 1039 | A | ARG | 1039 | C |
| VAL | 1040 | A | SER | 1030 | B |
| LYS | 558 | A | GLY | 283 | B |
| GLY | 889 | A | LYS | 1045 | C |
| SER | 711 | A | PRO | 897 | B |
| ALA | 570 | A | SER | 967 | B |
| ALA | 706 | A | GLN | 895 | B |
| SER | 1030 | A | PHE | 1042 | C |
| ASN | 914 | A | SER | 1123 | C |
| MET | 900 | A | PRO | 1079 | C |
| GLN | 755 | A | GLY | 971 | C |
| TYR | 707 | A | GLN | 895 | B |
| ALA | 890 | A | LYS | 1045 | C |
| ASP | 737 | A | ASN | 317 | C |
| PHE | 1121 | A | GLN | 913 | B |
| TRP | 886 | A | VAL | 1040 | C |
| MET | 869 | A | THR | 696 | C |
| VAL | 1040 | A | GLU | 1031 | B |
| ALA | 892 | A | PRO | 1069 | C |
| SER | 1123 | A | ASN | 914 | B |
| ILE | 788 | A | GLY | 700 | C |
| HIS | 519 | A | VAL | 42 | B |
| GLY | 566 | A | PHE | 43 | B |
| GLN | 913 | A | PHE | 1089 | C |
| ILE | 896 | A | SER | 711 | C |
| THR | 1006 | A | GLN | 762 | B |
| LEU | 894 | A | ILE | 714 | C |
| ILE | 896 | A | TYR | 707 | C |
| TYR | 380 | A | ILE | 973 | B |
| LYS | 557 | A | PHE | 43 | B |
| ILE | 569 | A | LYS | 854 | B |
| VAL | 1129 | A | TYR | 917 | B |
| PHE | 855 | A | PHE | 592 | C |
| TYR | 707 | A | THR | 883 | B |
| GLU | 702 | A | ILE | 788 | B |
| GLN | 755 | A | ASN | 969 | C |
| ILE | 569 | A | VAL | 963 | B |
| SER | 1037 | A | ARG | 1039 | C |
| ALA | 893 | A | VAL | 705 | C |
| THR | 385 | A | ASP | 985 | B |
| VAL | 42 | A | ARG | 567 | C |
| ASP | 198 | A | ARG | 355 | C |
| TYR | 369 | A | THR | 415 | C |
| THR | 883 | A | TYR | 707 | C |
| ASN | 703 | A | TYR | 789 | B |
| ASN | 856 | A | THR | 572 | C |
| VAL | 42 | A | PHE | 562 | C |
| PHE | 562 | A | GLU | 224 | B |
| ASP | 796 | A | TYR | 707 | C |
| MET | 900 | A | ILE | 712 | C |
| MET | 869 | A | ALA | 668 | C |
| TYR | 756 | A | PHE | 970 | C |
| ASN | 919 | A | VAL | 1128 | C |
| GLN | 784 | A | ASP | 1041 | C |
| LEU | 984 | A | GLY | 381 | C |
| ASN | 1074 | A | GLN | 895 | B |
| GLU | 516 | A | ASP | 228 | B |
| VAL | 1040 | A | ALA | 890 | B |
| GLN | 1002 | A | GLN | 1002 | B |
| GLN | 762 | A | TYR | 1007 | C |
| PRO | 897 | A | SER | 711 | C |
| ASP | 1041 | A | GLN | 784 | B |
| VAL | 42 | A | GLY | 566 | C |
| PHE | 855 | A | PRO | 589 | C |
| GLN | 1010 | A | LEU | 1012 | B |
| SER | 758 | A | SER | 968 | C |
| GLN | 1005 | A | GLN | 1002 | C |
| PHE | 562 | A | PRO | 225 | B |
| TRP | 886 | A | ASN | 1108 | C |
| PHE | 43 | A | GLY | 566 | C |
| THR | 302 | A | THR | 761 | B |
| TYR | 200 | A | SER | 514 | C |
| VAL | 976 | A | THR | 572 | C |
| GLU | 918 | A | VAL | 1129 | C |
| VAL | 1094 | A | TYR | 904 | B |
| ALA | 570 | A | ASN | 856 | B |
| LYS | 41 | A | ALA | 520 | C |
| ILE | 882 | A | TYR | 707 | C |
| PRO | 1090 | A | GLN | 913 | B |
| CYS | 662 | A | LEU | 864 | B |
| PRO | 897 | A | ILE | 712 | C |
| LEU | 390 | A | SER | 982 | B |
| ALA | 890 | A | GLY | 1046 | C |
| GLN | 314 | A | SER | 735 | B |
| LEU | 984 | A | VAL | 382 | C |
| SER | 708 | A | PRO | 897 | B |
| LEU | 984 | A | LYS | 386 | C |
| SER | 711 | A | GLN | 895 | B |
| MET | 869 | A | GLY | 669 | C |
| THR | 1027 | A | PHE | 1042 | C |
| GLY | 232 | A | ARG | 466 | C |
| GLU | 1111 | A | SER | 1123 | C |
| GLY | 669 | A | THR | 866 | B |
| PHE | 559 | A | PHE | 43 | B |
| SER | 704 | A | LYS | 790 | B |
| ILE | 712 | A | PRO | 897 | B |
| GLN | 762 | A | THR | 961 | C |
| VAL | 42 | A | GLN | 563 | C |
| ARG | 1107 | A | GLN | 913 | B |
| LEU | 861 | A | GLN | 613 | C |
| ASN | 764 | A | ASN | 317 | C |
| ARG | 983 | A | GLY | 381 | C |
| ALA | 570 | A | LYS | 854 | B |
| TYR | 789 | A | ASN | 703 | C |
| ARG | 765 | A | GLN | 957 | C |
| GLY | 381 | A | LEU | 984 | B |
| ASP | 574 | A | PHE | 43 | B |
| SER | 758 | A | THR | 961 | C |
| PRO | 384 | A | GLU | 988 | B |
| PRO | 862 | A | ILE | 666 | C |
| MET | 697 | A | LEU | 865 | B |
| LEU | 1145 | A | LEU | 1145 | B |
| ILE | 896 | A | ILE | 712 | C |
| ILE | 569 | A | LYS | 964 | B |
| GLU | 918 | A | SER | 1123 | C |
| PRO | 897 | A | ASN | 709 | C |
| GLY | 283 | A | GLN | 563 | C |
| MET | 697 | A | TYR | 873 | B |
| ILE | 788 | A | SER | 704 | C |
| ARG | 319 | A | THR | 739 | B |
| ARG | 765 | A | ALA | 958 | C |
| ILE | 712 | A | ILE | 896 | B |
| GLU | 1072 | A | LEU | 894 | B |
| ARG | 1107 | A | TYR | 904 | B |
| TYR | 200 | A | TYR | 396 | C |
| GLN | 1010 | A | GLN | 762 | B |
| ILE | 569 | A | ASN | 960 | B |
| ASP | 994 | A | PHE | 970 | C |
| PRO | 665 | A | ASP | 775 | B |
| ARG | 983 | A | LEU | 517 | C |
| GLU | 1072 | A | ALA | 893 | B |
| VAL | 705 | A | ALA | 893 | B |
| GLY | 999 | A | PHE | 759 | B |
| THR | 961 | A | GLN | 762 | B |
| GLY | 545 | A | SER | 982 | B |
| ASN | 703 | A | LYS | 790 | B |
| GLN | 920 | A | ILE | 1130 | C |
| GLN | 563 | A | ASP | 40 | B |
| LEU | 1141 | A | LEU | 1141 | B |
| ALA | 713 | A | GLN | 895 | B |
| SER | 1030 | A | ASP | 1041 | C |
| PRO | 1069 | A | LEU | 894 | B |

**Table S7.** The description of local interacting communities of the protomer A in the structure of SARS-CoV-2 S-GSAS/D614 in the closed state (pdb id 7KDG).

| Number | Contributing structural domains | Local Community Interacting Residues |
| --- | --- | --- |
| 1 | HR1-HR2 | A_1042_PHE A_1031_GLU A_1039_ARG |
| 2 | S2-HR1-HR2 | A_1067_TYR A_1049_LEU A_906_PHE A_909_ILE A_911_VAL A_915_VAL |
| 3 | S2-HR1-HR2 | A_1054_GLN A_816_SER A_819_GLU |
| 4 | NTD | A_215_ASP A_266_TYR A_64_TRP |
| 5 | NTD | A_275_PHE A_290_ASP A_58_PHE |
| 6 | NTD | A_279_TYR A_44_ARG A_47_VAL A_49_HIS |
| 7 | CTD2-NTD | A_664_ILE A_312_ILE A_598_ILE |
| 8 | RBD | A_464_PHE A_355_ARG A_514_SER A_429_PHE A_425_LEU A_426_PRO |
| 9 | RBD | A_509_ARG A_442_ASP A_438_SER |
| 10 | CTD2 | A_693_ILE A_656_VAL A_660_TYR |
| 11 | S2 | A_888_PHE A_789_TYR A_880_GLY |
| 12 | S2-HR1 | A_1001_LEU A_1005_GLN A_759_PHE |
| 13 | HR1-HR2 | A_1024_LEU A_1028_LYS A_1042_PHE A_1032_CYS |
| 14 | S2-HR1-HR2 | A_1062_PHE A_1029_MET A_1033_VAL A_1053_PRO A_877_LEU |
| 15 | HR1-HR2 | A_1032_CYS A_1043_CYS A_1048_HIS A_1051_SER A_1064_HIS |
| 16 | S2-HR1-HR2 | A_819_GLU A_1055_SER A_874_THR |
| 17 | HR1-HR2 | A_1075_PHE A_1096_VAL A_1110_TYR A_714_ILE |
| 18 | HR1-HR2 | A_1106_GLN A_1109_PHE A_915_VAL |
| 19 | NTD | A_231_ILE A_130_VAL A_168_PHE |
| 20 | NTD | A_193_VAL A_204_TYR A_37_TYR |
| 21 | NTD | A_285_ILE A_279_TYR A_38_TYR |
| 22 | NTD-CTD1-CTD2 | A_299_THR A_315_THR A_597_VAL |
| 23 | RBD | A_338_PHE A_342_PHE A_368_LEU |
| 24 | RBD | A_451_TYR A_401_VAL A_442_ASP |
| 25 | RBD | A_353_TRP A_398_ASP A_464_PHE |
| 26 | RBD-CTD1 | A_543_PHE A_585_LEU A_576_VAL |
| 27 | CTD2 | A_666_ILE A_650_LEU A_670_ILE A_645_THR |
| 28 | HR1 | A_977_LEU A_749_CYS A_993_ILE A_997_ILE |
| 29 | S2 | A_878_LEU A_806_LEU A_882_ILE |
| 30 | S2 | A_906_PHE A_923_ILE A_916_LEU |
| 31 | HR1 | A_1000_ARG A_977_LEU A_996_LEU |
| 32 | S2-HR1-HR2 | A_741_TYR A_1004_LEU A_962_LEU A_858_LEU |
| 33 | S2-HR1-HR2 | A_902_MET A_1050_MET A_898_PHE |
| 34 | RBD | A_342_PHE A_511_VAL A_374_PHE A_436_TRP |
| 35 | RBD | A_353_TRP A_400_PHE A_423_TYR A_512_VAL A_410_ILE |
| 36 | RBD | A_392_PHE A_515_PHE A_395_VAL |
| 37 | S2-HR1-HR2 | A_1065_VAL A_1052_PHE A_802_PHE A_927_PHE |
| 38 | RBD-CTD2 | A_314_GLN A_596_SER A_613_GLN |

**Table S8.** The description of	local interacting communities of the protomer A in the structure of	SARS-CoV-2 S-GSAS/D614 in the open state (pdb id 7KDH)

| Number | Contributing structural domains | Local Community Interacting Residues |
| --- | --- | --- |
| 1 | S2-HR1 | A_731_MET A_1014_ARG A_955_ASN |
| 2 | NTD | A_101_ILE A_240_THR A_265_TYR A_92_PHE |
| 3 | S2-HR1-HR2 | A_1028_LYS A_1062_PHE A_727_LEU |
| 4 | HR1-HR2 | A_1042_PHE A_1031_GLU A_1039_ARG |
| 5 |  | A_1032_CYS A_1048_HIS A_1051_SER A_1064_HIS |
| 6 | S2-HR1-HR2 | A_1067_TYR A_1049_LEU A_909_ILE |
| 7 | S2-HR1-HR2 | A_1054_GLN A_816_SER A_819_GLU |
| 8 | HR1-HR2 | A_1102_TRP A_1081_ILE A_1135_ASN |
| 9 | NTD | A_238_PHE A_92_PHE A_267_VAL |
| 10 | NTD | A_275_PHE A_290_ASP A_58_PHE |
| 11 | NTKD | A_279_TYR A_44_ARG A_47_VAL A_49_HIS |
| 12 | NTD-CTD2 | A_299_THR A_315_THR A_597_VAL |
| 13 | CTD2 | A_664_ILE A_312_ILE A_598_ILE |
| 14 | RBD | A_418_ILE A_495_TYR A_453_TYR |
| 15 | RBD | A_464_PHE A_429_PHE A_425_LEU A_426_PRO A_514_SER |
| 16 | RBD | A_454_ARG A_457_ARG A_467_ASP |
| 17 | HR1-HR2 | A_1062_PHE A_1029_MET A_1033_VAL |
| 18 | S2-HR1-  HR2 | A_819_GLU A_1055_SER A_874_THR |
| 19 | HR1-HR2 | A_1106_GLN A_1109_PHE A_915_VAL |
| 20 | NTD | A_193_VAL A_204_TYR A_37_TYR |
| 21 | NTD | A_285_ILE A_279_TYR A_38_TYR |
| 22 | RBD-CTD1 | A_579_PRO A_330_PRO A_544_ASN |
| 23 | RBD | A_351_TYR A_454_ARG A_492_LEU |
| 24 | RBD-CTD1 | A_365_TYR A_387_LEU A_515_PHE A_432_CYS |
| 25 | CTD1 | A_541_PHE A_552_LEU A_587_ILE |
| 26 | S2-HR1 | A_997_ILE A_749_CYS A_993_ILE |
| 27 | S2 | A_898_PHE A_802_PHE A_797_PHE |
| 28 | S2 | A_878_LEU A_806_LEU A_882_ILE |
| 29 | S2 | A_818_ILE A_804_GLN A_935_GLN |
| 30 | S2-HR1 | A_923_ILE A_916_LEU A_906_PHE |
| 31 | S2-HR1 | A_905_ARG A_1050_MET A_898_PHE A_902_MET |
| 32 | S2-HR1-HR2 | A_1065_VAL A_1052_PHE A_802_PHE A_805_ILE A_878_LEU A_927_PHE |
| 33 | RBD-CTD1 | A_328_ARG A_543_PHE A_579_PRO |
| 34 | RBD | A_342_PHE A_511_VAL A_374_PHE A_436_TRP |
| 35 | S2-HR1-HR2 | A_805_ILE A_1054_GLN A_818_ILE |

**Table S9.** The description of local interacting communities of the protomer A in the structure of SARS-CoV-2 S-GSAS/G614 in the closed state (pdb id 7KDK)

| Number | Contributing structural domains | Local Community Interacting Residues |
| --- | --- | --- |
| 1 | NTD | A_101_ILE A_190_ARG A_94_SER |
| 2 | HR1-HR2 | A_1042_PHE A_1031_GLU A_1037_SER A_1039_ARG A_1032_CYS |
| 3 | S2-HR1-HR2 | A_1067_TYR A_1049_LEU A_906_PHE A_909_ILE A_911_VAL A_915_VAL |
| 4 | S2-HR1-HR2 | A_1054_GLN A_816_SER A_819_GLU |
| 5 | S2-HR1-HR2 | A_1063_LEU A_724_THR A_934_ILE |
| 6 | S2-HR1-HR2 | A_1075_PHE A_1110_TYR A_714_ILE |
| 7 | NTD | A_265_TYR A_92_PHE A_240_THR |
| 8 | NTD | A_275_PHE A_290_ASP A_58_PHE |
| 9 | NTD | A_279_TYR A_44_ARG A_47_VAL A_49_HIS |
| 10 | NTD-CTD2 | A_315_THR A_299_THR A_597_VAL |
| 11 | NTD-RBD-CTD1 | A_328_ARG A_530_SER A_580_GLN |
| 12 | CTD1 | A_579_PRO A_544_ASN A_564_GLN |
| 13 | S2 | A_888_PHE A_789_TYR A_880_GLY |
| 14 | S2-HR1 | A_1001_LEU A_1005_GLN A_759_PHE |
| 15 | HR1-HR2 | A_1024_LEU A_1028_LYS A_1042_PHE |
| 16 | S2-HR1-HR2 | A_1062_PHE A_1029_MET A_1033_VAL A_1053_PRO A_877_LEU |
| 17 | S2-HR1-HR2 | A_905_ARG A_1050_MET A_898_PHE A_901_GLN |
| 18 | S2-HR1-HR2 | A_819_GLU A_1055_SER A_874_THR |
| 19 | NTD | A_194_PHE A_238_PHE A_106_PHE A_117_LEU A_201_PHE A_235_ILE A_86_PHE A_231_ILE A_90_VAL |
| 20 | NTD | A_193_VAL A_204_TYR A_37_TYR |
| 21 | NTD-RBD-CTD1 | A_328_ARG A_533_LEU A_578_ASP |
| 22 | RBD | A_338_PHE A_342_PHE A_368_LEU |
| 23 | RBD | A_353_TRP A_398_ASP A_464_PHE |
| 24 | RBD | A_365_TYR A_387_LEU A_515_PHE |
| 25 | RBD | A_406_GLU A_403_ARG A_495_TYR |
| 26 | RBD | A_509_ARG A_401_VAL A_451_TYR A_442_ASP A_438_SER A_507_PRO |
| 27 | CTD1 | A_543_PHE A_585_LEU A_576_VAL |
| 28 | S2-HR1 | A_977_LEU A_749_CYS A_993_ILE A_997_ILE |
| 29 | S2 | A_878_LEU A_806_LEU A_882_ILE |
| 30 | HR1 | A_973_ILE A_984_LEU A_992_GLN |
| 31 | S2-HR1 | A_1000_ARG A_977_LEU A_996_LEU |
| 32 | S2-HR1 | A_741_TYR A_1004_LEU A_962_LEU A_858_LEU |
| 33 | HR1-HR2 | A_1065_VAL A_1052_PHE A_802_PHE A_927_PHE |
| 34 | NTD-RBD-CTD1 | A_328_ARG A_543_PHE A_579_PRO |
| 35 | RBD | A_342_PHE A_511_VAL A_374_PHE A_436_TRP A_347_PHE |
| 36 | RBD | A_353_TRP A_400_PHE A_423_TYR A_512_VAL A_410_ILE |
| 37 | NTD | A_104_TRP A_194_PHE A_238_PHE A_92_PHE |
| 38 | NTD-CTD2 | A_664_ILE A_312_ILE A_598_ILE |
| 39 | CTD2 | A_611_LEU A_666_ILE A_650_LEU |
| 40 | S2 | A_898_PHE A_802_PHE A_797_PHE |

**Table S10.** The description of local interacting communities of the protomer A in the structure of SARS-CoV-2 S-GSAS/G614 in the open state (pdb id 7KDL)

| Number | Contributing structural domains | Local Community Interacting Residues |
| --- | --- | --- |
| 1 | S2-HR1 | A_1001_LEU A_1005_GLN A_759_PHE |
| 2 | HR1-HR2 | A_1042_PHE A_1031_GLU A_1037_SER A_1039_ARG |
| 3 | S2-HR1-HR2 | A_1067_TYR A_1049_LEU A_906_PHE A_909_ILE A_911_VAL |
| 4 | S2-HR1-HR2 | A_1054_GLN A_816_SER A_819_GLU |
| 5 | S2-HR1-HR2 | A_1075_PHE A_1110_TYR A_714_ILE |
| 6 | HR1-HR2 | A_1102_TRP A_1081_ILE A_1135_ASN |
| 7 | NTD | A_186_PHE A_264_ALA A_66_HIS |
| 8 | NTD | A_189_LEU A_210_ILE A_217_PRO |
| 9 | NTD | A_220_PHE A_288_ALA A_36_VAL |
| 10 | NTD | A_265_TYR A_92_PHE A_240_THR |
| 11 | NTD | A_265_TYR A_65_PHE A_82_PRO |
| 12 | NTD | A_275_PHE A_290_ASP A_58_PHE |
| 13 | NTD | A_279_TYR A_44_ARG A_49_HIS |
| 14 | NTD | A_299_THR A_315_THR A_597_VAL |
| 15 | CTD2 | A_664_ILE A_312_ILE A_598_ILE |
| 16 | RBD | A_342_PHE A_511_VAL A_374_PHE A_436_TRP A_347_PHE A_399_SER A_509_ARG |
| 17 | RBD | A_351_TYR A_452_LEU A_454_ARG A_492_LEU |
| 18 | NTD | A_91_TYR A_35_GLY A_56_LEU |
| 19 | RBD | A_464_PHE A_429_PHE A_425_LEU A_426_PRO A_514_SER |
| 20 | RBD | A_451_TYR A_401_VAL A_497_PHE A_448_ASN |
| 21 | RBD | A_454_ARG A_457_ARG A_467_ASP |
| 22 | CTD2 | A_611_LEU A_666_ILE A_650_LEU |
| 23 | S2 | A_888_PHE A_789_TYR A_880_GLY |
| 24 | HR1-HR2 | A_1000_ARG A_977_LEU A_996_LEU |
| 25 | HR1-HR2 | A_1024_LEU A_1028_LYS A_1042_PHE A_1032_CYS |
| 26 | S2-HR1-HR2 | A_1062_PHE A_1029_MET A_1033_VAL A_1053_PRO A_877_LEU |
| 27 | HR1-HR2 | A_1032_CYS A_1043_CYS A_1048_HIS A_1051_SER A_1064_HIS |
| 28 | S2-HR1-HR2 | A_819_GLU A_1055_SER A_874_THR |
| 29 | HR1-HR2 | A_1115_ILE A_1104_VAL A_1119_ASN |
| 30 | S2-HR1-HR2 | A_1106_GLN A_1109_PHE A_915_VAL |
| 31 | NTD | A_193_VAL A_204_TYR A_37_TYR |
| 32 | NTD | A_104_TRP A_194_PHE A_238_PHE A_92_PHE A_267_VAL A_84_LEU |
| 33 | NTD | A_238_PHE A_106_PHE A_117_LEU A_201_PHE A_235_ILE A_86_PHE A_90_VAL |
| 34 | RBD-CTD1 | A_579_PRO A_330_PRO A_544_ASN |
| 35 | RBD | A_338_PHE A_342_PHE A_368_LEU |
| 36 | RBD | A_353_TRP A_398_ASP A_423_TYR A_464_PHE |
| 37 | RBD | A_365_TYR A_387_LEU A_515_PHE |
| 38 | CTD1 | A_543_PHE A_585_LEU A_576_VAL |
| 39 | S2-HR1 | A_997_ILE A_749_CYS A_993_ILE |
| 40 |  | A_878_LEU A_806_LEU A_882_ILE |
| 41 | S2 | A_906_PHE A_902_MET A_923_ILE A_916_LEU |
| 42 | S2-HR1 | A_741_TYR A_1004_LEU A_962_LEU A_858_LEU |
| 43 | S2-HR1-HR2 | A_905_ARG A_1050_MET A_898_PHE A_902_MET |
| 44 | HR1-HR2 | A_1065_VAL A_1052_PHE A_802_PHE A_927_PHE |
| 45 | HR1-HR2 | A_1081_ILE A_1115_ILE A_1137_VAL |
| 46 | NTD | A_285_ILE A_38_TYR A_279_TYR |
| 47 | NTD-RBD-CTD1 | A_328_ARG A_543_PHE A_579_PRO |
| 48 | S2 | A_898_PHE A_802_PHE A_797_PHE |
| 49 | RBD-CTD2 | A_314_GLN A_596_SER A_613_GLN |

**Table S11.** The description of	local interacting communities of the protomer A in the structure of	SARS-CoV-2 S-RRAR/G614 in the closed state (pdb id 7KDI)

| Number | Contributing structural domains | Local Community Interacting Residues |
| --- | --- | --- |
| 1 | HR1-HR2 | A_1042_PHE A_1031_GLU A_1039_ARG |
| 2 | HR1-HR2 | A_1032_CYS A_1051_SER A_1064_HIS |
| 3 | S2-HR1-HR2 | A_1067_TYR A_1049_LEU A_906_PHE |
| 4 | S2-HR1-HR2 | A_1054_GLN A_816_SER A_819_GLU |
| 5 | NTD | A_34_ARG A_191_GLU A_221_SER |
| 6 | NTD | A_265_TYR A_65_PHE A_82_PRO |
| 7 | NTD | A_275_PHE A_290_ASP A_58_PHE |
| 8 | NTD | A_279_TYR A_44_ARG A_49_HIS |
| 9 | NTD-RBD-CTD1 | A_326_ILE A_534_VAL A_539_VAL |
| 10 | NTD-RBD-CTD1 | A_328_ARG A_530_SER A_580_GLN |
| 11 | RBD | A_353_TRP A_398_ASP A_464_PHE |
| 12 | S2 | A_888_PHE A_789_TYR A_880_GLY |
| 13 | S2-HR1 | A_1001_LEU A_1005_GLN A_759_PHE |
| 14 | S2-HR1-HR2 | A_1024_LEU A_1028_LYS A_1042_PHE A_727_LEU A_1062_PHE |
| 15 | S2-HR1-HR2 | A_1062_PHE A_1029_MET A_1033_VAL A_1053_PRO A_877_LEU |
| 16 | NTD | A_104_TRP A_194_PHE A_240_THR A_92_PHE A_265_TYR |
| 17 | S2-HR1-HR2 | A_819_GLU A_1055_SER A_874_THR |
| 18 | S2-HR1-HR2 | A_1106_GLN A_1109_PHE A_915_VAL |
| 19 | NTD | A_193_VAL A_204_TYR A_37_TYR |
| 20 | NTD | A_106_PHE A_201_PHE A_235_ILE A_238_PHE A_86_PHE A_90_VAL |
| 21 | NTD-RBD-CTD1 | A_299_THR A_315_THR A_597_VAL |
| 22 | NBD-RBD-CTD1 | A_328_ARG A_533_LEU A_578_ASP |
| 23 | RBD | A_365_TYR A_387_LEU A_515_PHE |
| 24 | RBD | A_418_ILE A_406_GLU A_409_GLN |
| 25 | RBD | A_509_ARG A_438_SER A_442_ASP |
| 26 | CTD1 | A_543_PHE A_585_LEU A_576_VAL |
| 27 | CTD-CTD2 | A_552_LEU A_541_PHE A_587_ILE |
| 28 | CTD1 | A_577_ARG A_584_ILE A_559_PHE |
| 29 | S2-HR1 | A_977_LEU A_749_CYS A_993_ILE A_997_ILE |
| 30 | S2 | A_878_LEU A_806_LEU A_882_ILE |
| 31 | S2 | A_906_PHE A_923_ILE A_916_LEU |
| 32 | S2-HR1 | A_741_TYR A_1004_LEU A_962_LEU A_858_LEU |
| 33 | S2-HR1-HR2 | A_1050_MET A_898_PHE A_902_MET A_905_ARG |
| 34 | S2-HR1-HR2 | A_1065_VAL A_1052_PHE A_802_PHE A_927_PHE |
| 35 | NTD | A_92_PHE A_238_PHE A_267_VAL A_84_LEU A_65_PHE |
| 36 | RBD | A_353_TRP A_400_PHE A_423_TYR A_410_ILE |
| 37 | S2 | A_898_PHE A_802_PHE A_797_PHE |
| 38 | NTD-CTD2 | A_664_ILE A_312_ILE A_598_ILE |
| 39 | CTD2 | A_611_LEU A_666_ILE A_650_LEU |

**Table S12.** The description of	local interacting communities of the protomer A in the structure of	SARS-CoV-2 S-RRAR/G614 in the open state (pdb id 7KDJ)

| Number | Contributing structural domains | Local Community Interacting Residues |
| --- | --- | --- |
| 1 | S2-HR1 | A_1001_LEU A_1005_GLN A_759_PHE |
| 2 | HR1-HR2 | A_1024_LEU A_1028_LYS A_1042_PHE |
| 3 | HR1-HR2 | A_1042_PHE A_1031_GLU A_1039_ARG |
| 4 | HR1-HR2 | A_1032_CYS A_1051_SER A_1064_HIS |
| 5 | S2-HR1-HR2 | A_1067_TYR A_1049_LEU A_906_PHE A_909_ILE A_911_VAL |
| 6 | S2-HR1-HR2 | A_1054_GLN A_816_SER A_819_GLU |
| 7 | S2-HR1-HR2 | A_819_GLU A_1055_SER A_874_THR |
| 8 | NTD | A_201_PHE A_117_LEU A_231_ILE |
| 9 | NTD | A_220_PHE A_288_ALA A_36_VAL |
| 10 | NTD | A_275_PHE A_290_ASP A_58_PHE |
| 11 | NTD | A_276_LEU A_289_VAL A_306_PHE |
| 12 | NTD | A_279_TYR A_44_ARG A_49_HIS |
| 13 | S2-N2R-CTD2 | A_664_ILE A_312_ILE A_598_ILE |
| 14 | NTD | A_91_TYR A_35_GLY A_56_LEU |
| 15 | RBD | A_464_PHE A_429_PHE A_512_VAL A_425_LEU A_426_PRO A_514_SER |
| 16 | RBD | A_454_ARG A_457_ARG A_467_ASP |
| 17 | S2-HR1 | A_997_ILE A_749_CYS A_977_LEU A_993_ILE |
| 18 | S2 | A_888_PHE A_789_TYR A_880_GLY |
| 19 | S2-HR1 | A_1007_TYR A_1011_GLN A_767_LEU |
| 20 | NTD | A_101_ILE A_240_THR A_265_TYR A_92_PHE |
| 21 | S2-HR1-HR2 | A_1029_MET A_1053_PRO A_877_LEU |
| 22 | S2-HR1-HR2 | A_1065_VAL A_1052_PHE A_802_PHE A_927_PHE |
| 23 | NTD | A_238_PHE A_106_PHE A_201_PHE A_235_ILE A_86_PHE A_90_VAL |
| 24 | S2-HR1-HR2 | A_1106_GLN A_1109_PHE A_915_VAL |
| 25 | NTD | A_193_VAL A_204_TYR A_37_TYR |
| 26 | NTD | A_238_PHE A_267_VAL A_84_LEU A_65_PHE |
| 27 | N2R-RBD-CTD1 | A_328_ARG A_543_PHE A_579_PRO |
| 28 | RBD | A_338_PHE A_342_PHE A_368_LEU |
| 29 | RBD | A_350_VAL A_402_ILE A_495_TYR |
| 30 | RBD | A_353_TRP A_398_ASP A_464_PHE |
| 31 | RBD | A_365_TYR A_387_LEU A_515_PHE |
| 32 | S2 | A_878_LEU A_806_LEU A_882_ILE |
| 33 | S2 | A_898_PHE A_802_PHE A_797_PHE |
| 34 | S2 | A_906_PHE A_923_ILE A_916_LEU |
| 35 | NTD | A_104_TRP A_194_PHE A_92_PHE |
| 36 | S2-HR1-HR2 | A_905_ARG A_1050_MET A_898_PHE A_902_MET |
| 37 | NTD | A_223_LEU A_193_VAL A_91_TYR |
| 38 | NTD-RBD-CTD1 | A_597_VAL A_315_THR A_299_THR |
| 39 | RBD | A_342_PHE A_511_VAL A_374_PHE A_436_TRP |
| 40 | RBD | A_353_TRP A_400_PHE A_423_TYR A_512_VAL A_410_ILE |
| 41 | CTD1 | A_543_PHE A_585_LEU A_576_VAL |
| 42 | CTD-CTD2 | A_552_LEU A_541_PHE A_587_ILE |
| 43 | CTD1 | A_577_ARG A_584_ILE A_559_PHE |
| 44 | NTD-RBD-CTD1 | A_326_ILE A_534_VAL A_539_VAL |
| 45 | NTD-RBD-CTD1 | A_328_ARG A_530_SER A_580_GLN |
